# The control of transcriptional memory by stable mitotic bookmarking

**DOI:** 10.1101/2021.08.30.458146

**Authors:** Maëlle Bellec, Jérémy Dufourt, George Hunt, Hélène Lenden-Hasse, Antonio Trullo, Amal Zine El Aabidine, Marie Lamarque, Marissa M Gaskill, Heloïse Faure-Gautron, Mattias Mannervik, Melissa M Harrison, Jean-Christophe Andrau, Cyril Favard, Ovidiu Radulescu, Mounia Lagha

**Affiliations:** Institut de Génétique Moléculaire de Montpellier, University of Montpellier, CNRS-UMR 5535, 1919 Route de Mende, Montpellier 34293 cedex 5, France; Department of Molecular Biosciences, The Wenner-Gren Institute, Stockholm University, 10691 Stockholm, Sweden; Department of Biomolecular Chemistry, School of Medicine and Public Health, University of Wisconsin-Madison, Madison, Wisconsin 53706, USA; Institut de Recherche en Infectiologie de Montpellier, CNRS UMR 9004, University of Montpellier, 1919 Route de Mende, Montpellier 34293 cedex 5, France; LPHI, UMR CNRS 5235, University of Montpellier, Place E. Bataillon – Bât. 24 cc 107, Montpellier 34095 cedex 5, France

## Abstract

To maintain cellular identities during development, gene expression profiles must be faithfully propagated through cell generations. The reestablishment of gene expression patterns upon mitotic exit is thought to be mediated, in part, by mitotic bookmarking by transcription factors (TF). However, the mechanisms and functions of TF mitotic bookmarking during early embryogenesis remain poorly understood. In this study, taking advantage of the naturally synchronized mitoses of *Drosophila* early embryos, we provide evidence that the pioneer-like transcription factor GAF acts as stable mitotic bookmarker during zygotic genome activation. We report that GAF remains associated to a large fraction of its interphase targets including at *cis*-regulatory sequences of key developmental genes, with both active and repressive chromatin signatures. GAF mitotic targets are globally accessible during mitosis and are bookmarked via histone acetylation (H4K8ac). By monitoring the kinetics of transcriptional activation in living embryos, we provide evidence that GAF binding establishes competence for rapid activation upon mitotic exit.

## Introduction

Cellular identities are determined by the precise spatio-temporal control of gene expression programs. These programs must be faithfully transmitted during each cellular division. However, with its drastic nuclear reorganization, mitosis represents a major challenge to the propagation of gene expression programs. How cells overcome this mitotic challenge to transmit information to their progeny remains relatively unexplored during embryogenesis (Bellec et al. 2018; Festuccia et al. 2017; Elsherbiny and Dobreva 2021).

Based on live imaging studies and genome-wide profiling experiments on drug-synchronized mitotic cells, it is now well established that a subset of transcription factors (TF), chromatin regulators and histone modifications are retained on their targets during mitosis (Festuccia et al. 2017; Raccaud and Suter 2017; Raccaud et al. 2019) *via* specific, non-specific DNA binding or a combination of both (Cirillo et al. 2002; Raccaud et al. 2019).

When the persistence of TF binding during mitosis is associated with a regulatory role in transcriptional activation upon mitotic exit, TFs can be envisaged as mitotic bookmarkers. The kinetics of post-mitotic re-activation are often examined by whole-genome profiling experiments of nascent transcription in early G1 (Palozola et al. 2017; Zhang et al. 2019). Combining such approaches with the mitotic depletion of candidate bookmarkers, it was established that some mitotically retained TFs/General TFs/histone marks act as *bona fide* mitotic bookmarkers (Kadauke et al. 2012; Teves et al. 2018; Festuccia et al. 2017).

Parallel to these multi-omics approaches, imaging of transcription in live cells with signal amplifying systems as the MS2/MCP (Pichon et al. 2018) allows for the direct quantification of the kinetics of transcriptional activation upon mitotic exit. With such approaches, mitotic bookmarking has been associated with an accelerated transcriptional reactivation after mitosis in cultured cells (Zhao et al. 2011). Moreover, this method enabled the visualization of the transmission of active states, referred to as ‘transcriptional memory’ in *Dictyostellium* and in *Drosophila* embryos (Ferraro et al. 2016; Muramoto et al. 2010). However, how mitotic bookmarking is associated with the transmission of states across mitosis in the context of a developing embryo remains unclear.

This question is particularly important during the first hours of development of all metazoans, when cellular divisions are rapid and frequent. During this period, there is a substantial chromatin reprogramming and transcriptional activation, called Zygotic Genome Activation (ZGA)(Vallot and Tachibana 2020; Schulz and Harrison 2019). The control of this major developmental transition is supervised by key TFs, a subset of which are capable of engaging inaccessible chromatin and foster nucleosome eviction, a defining property of pioneer factors (Iwafuchi-Doi and Zaret 2016). Remarkably, many mitotic bookmarking factors have pioneer factor properties (Zaret 2020).

In *Drosophila melanogaster*, two essential transcription factors with pioneering factor properties, Zelda and GAGA Associated Factor (GAF), orchestrate the reshaping of the genome during ZGA (Liang et al. 2008; Sun et al. 2015; Gaskill et al. 2021; Moshe and Kaplan 2017; Schulz et al. 2015). Contrary to Zelda, which is not retained during mitosis and is dispensable for transcriptional memory (Dufourt et al. 2018), GAF is known to decorate mitotic chromosomes (Raff et al. 1994; Gaskill et al. 2021; Dufourt et al. 2018). Here we asked whether GAF acts as a mitotic bookmarker during ZGA. GAF, encoded by the *Trithorax-like* gene, binds to repeating (GA)_n_ sequences and displays a broad set of functions including gene activation or silencing, nucleosome remodeling and chromatin organization (Chetverina et al. 2021). In addition, GAF has been shown to be enriched at paused promoters (Li and Gilmour 2013) and its manipulation in *Drosophila* S2 cells demonstrated a capacity to rapidly evict nucleosomes, thereby facilitating the recruitment of Pol II at promoters (Fuda et al. 2015; Judd et al. 2021). Together with its mitotic retention, these properties place GAF as a reasonable candidate for mitotic bookmarking during development.

## Results

### Endogenous GAF is retained during mitosis in early development and stably binds DNA

To investigate the function of GAF during mitosis, we first characterized its distribution during the cell cycle. With immunostaining, we confirmed that GAF is present on chromatin during all stages of mitosis from prophase to telophase (Fig. 1a) (Raff et al. 1994; Dufourt et al. 2018). Next, we examined GAF behavior in living embryos using an endogenously GFP tagged allele of GAF (Gaskill et al. 2021) (Fig. 1b, Movie 1). During mitosis, a large amount of GAF protein is displaced to the cytoplasm, but a clear pool of GAF protein remains associated with mitotic chromosomes (Fig. 1b).

**Figure 1:**
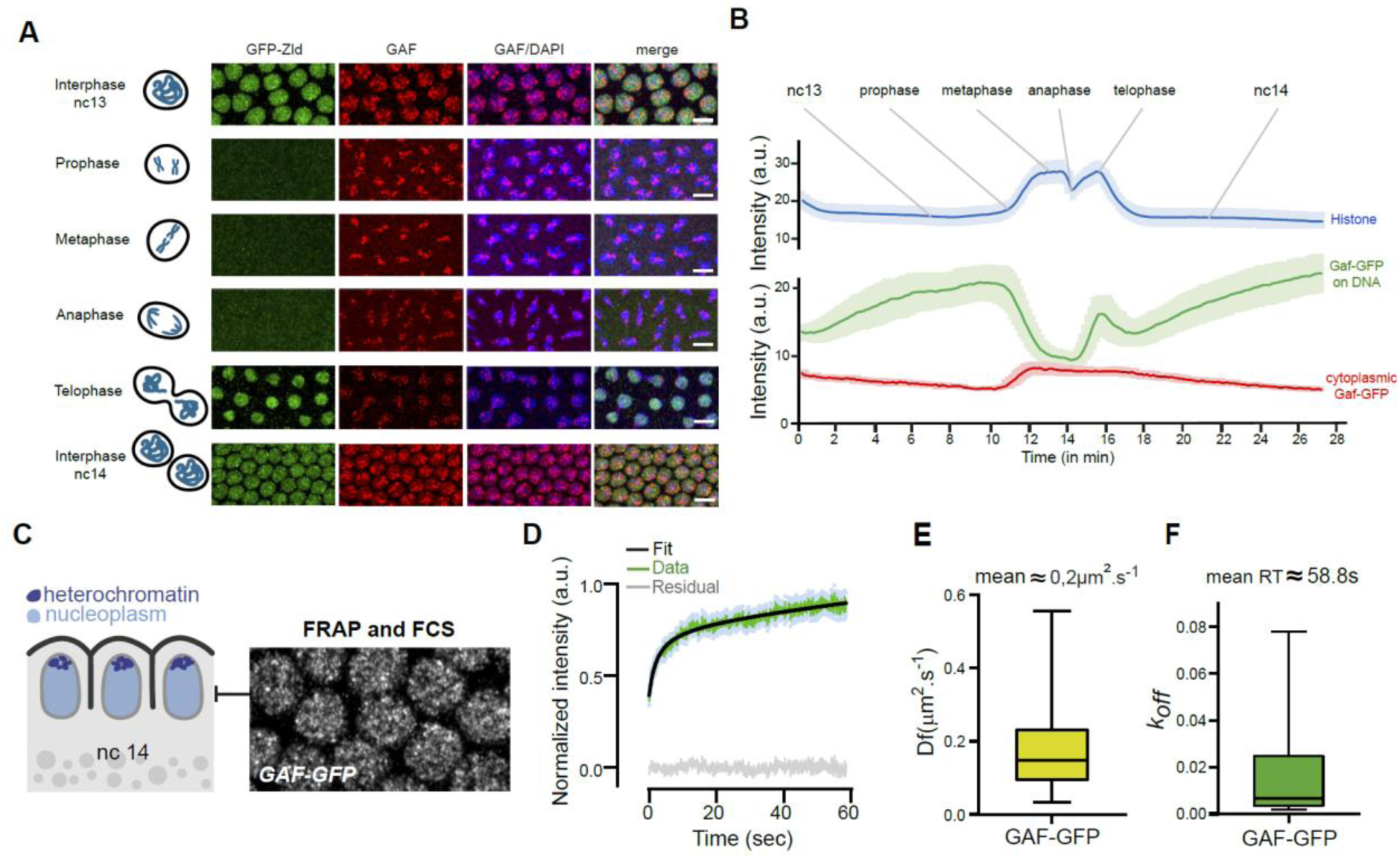
**GAF dynamics during nuclear cycles and its kinetic properties** (A) Maximum intensity projected Z-planes of confocal images from immunostaining of Zelda-GFP (green) and GAF (red) on interphase and mitotic embryos at the indicated stages counterstained with DAPI (blue). Scale bar is 5µm. (B) Mean fluorescent signal quantifications of GAF-GFP in nucleoplasm (green) and cytoplasm (red), and H2Av-RFP in nucleoplasm during nuclear cycle 13 to 14 extracted from time-lapse movies of embryos expressing GAF-GFP and H2Av-RFP (mean from three movies of three independent embryos). (C) Schematic of sagittal view of nc14 embryos. Nuclei are represented in light blue and apical heterochromatin regions in dark blue. Right panel shows regions targeted by FRAP and FCS, performed on GAF-GFP embryos. (D) Mean fluorescence recovery curve (green) from FRAP experiment and fit (black) using a reaction-diffusion model determined at the bleached spot for 23 nuclei from nine nc14 GAF-GFP embryos. Light blue dots represent SEM from different nuclei. Grey curve represents the residual of the fit. (E) Estimated diffusion coefficient of GAF-GFP. Centered line represents the median and whiskers represent min and max values. (F) Estimated koff (RT: residence time = 1/koff) of GAF-GFP. Centered line represents the median and whiskers represent min and max values.

From both live imaging and immunofluorescence data, we observed a strong GAF signal concentrated in large distinct puncta as well as a more diffuse signal within the nucleus. Consistent with previous work ^29^, we found that the majority of large GAF puncta are located at the apical side of the nuclei (Supplementary Fig. 1a, Movie 2), where at this stage, most of centromeric heterochromatin is located (Supplementary Fig. 1b) (Foe et al. 1993). In contrast to GAF apical foci, the rest of the nuclear space contains a homogeneously distributed GAF signal, potentially representing GAF binding to euchromatin (Supplementary Fig. 1a-b, Movie 2). To characterize GAF diffusion and binding kinetics in these regions, we performed Fluorescence Correlation Spectroscopy (FCS) and imaging Fluorescence Recovery After Photobleaching (FRAP) (Auer et al. 2021) on living GAF-GFP embryos during interphase (Fig. 1c, Supplementary Fig. 1c-d). We could not perform FRAP and FCS during mitosis due to their short durations and rapid nuclear movements.

FRAP recovery curves showed two characteristic times, a short one and a surprisingly long one (Fig. 1d). The short recovery time could correspond to fast unbinding or to diffusion. We confirmed that this short recovery time corresponds to diffusion (Fig. 1e) using FCS experiments (Supplementary Fig. 1e-f). The long-lived characteristic time from FRAP data, with a residence time close to a minute (Fig. 1f) is believed to correspond to GAF sequence-specific binding. Such a long-lived binding is an order of magnitude longer than typical TF residence times in *Drosophila* embryos (Mir et al. 2017; Dufourt et al. 2018). We conclude that GAF protein has the intrinsic capacity to stably bind chromatin. This property could be involved in its capacity to associate to mitotic chromosomes during embryonic divisions.

### Capturing GAF mitotic targets genome-wide

Early *Drosophila* embryogenesis provides an ideal system to study mitosis. Indeed, nuclei of the syncytial embryo divide 13 times synchronously before cellularization (Farrell and O’Farrell 2014). To perform mitotic ChIP, we stained early staged embryos with antibodies against the mitotic specific marker H3S10ph (Supplementary Fig. 2a) (Hendzel et al. 1997; Follmer et al. 2012) and sorted them with a flow cytometer (Fig. 2a and Supplementary Fig. 2b). The pools of embryos were further manually sorted to avoid contamination (Supplementary Fig. 2b). We applied this method to map GAF targets during mitosis and interphase. We retrieved GAF peaks genome-wide in interphase and mitotic samples and classified them into three categories: present only in interphase, only during mitosis, or during both interphase and mitosis, referred to as ‘mitotically retained’ (Fig. 2b-c’’). Remarkably, mitotically retained GAF targets represent 37% of interphase targets, corresponding to a group of ∼2000 peaks bound by GAF both in interphase and mitotic embryos (Fig. 2b). The mitotically retained loci comprise many key developmental patterning genes, as exemplified by *snail*, for which the proximal enhancer shows a GAF mitotic peak (Fig. 2c’).

**Figure 2:**
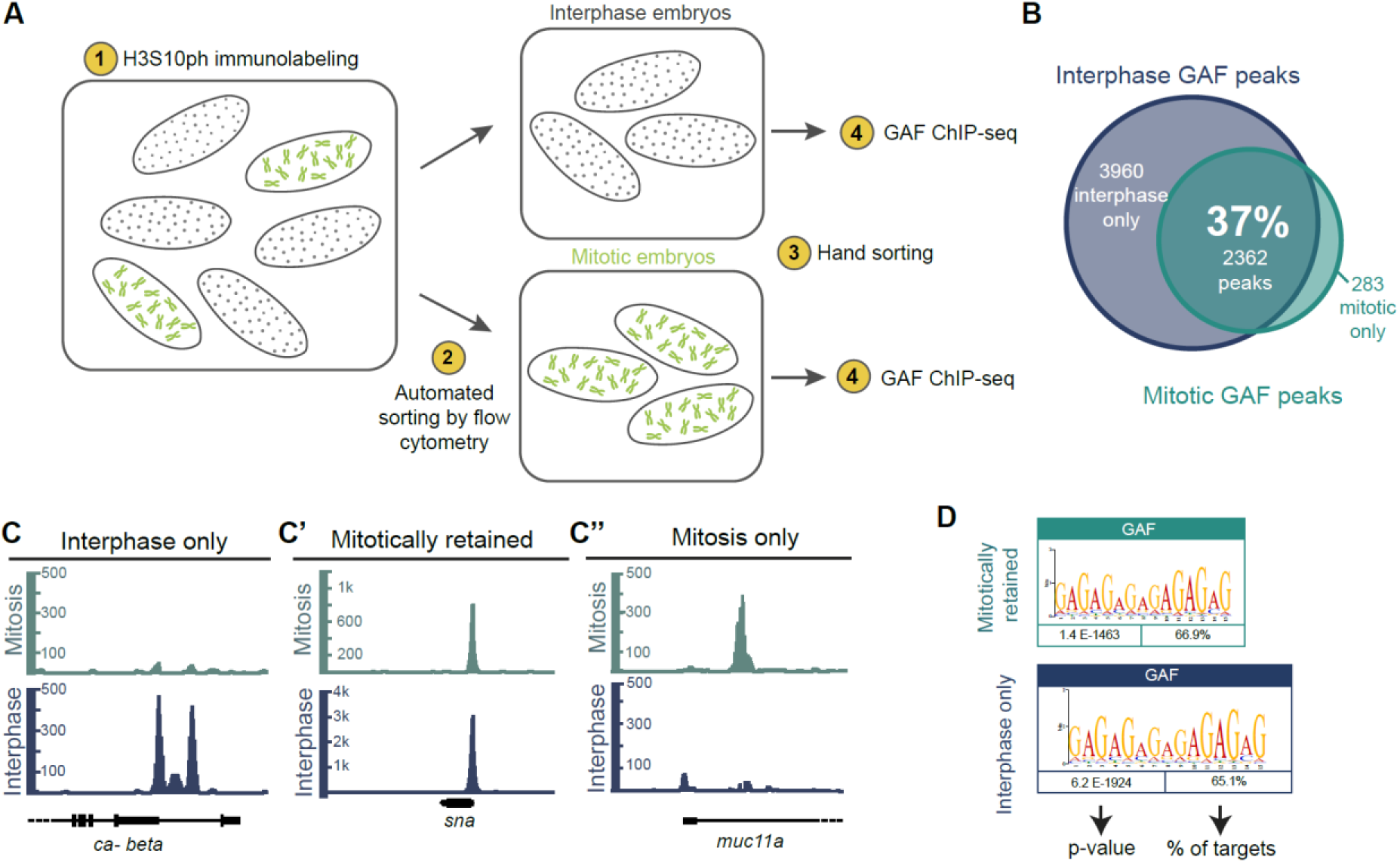
**Identification of thousands of mitotically retained GAF loci** (A) Experimental workflow of mitotic embryo sorting followed by GAF-ChIP-seq. (B) Venn diagram representing the overlap of called GAF-ChIP-seq peaks between interphase and mitotic embryos. (C) (C’) (C’’) Genome browser examples of genes from the identified three categories of GAF-ChIP-seq peaks: interphase only, mitotically retained and mitosis only, respectively. (D) (GA)n motif enrichment within GAF mitotically retained and interphase only peaks, as reported by MEME. (E) Box plot representing the number of GAGAG motifs within three different classes of GAF peaks: mitotically retained (light blue), interphase only (dark blue) and all peaks (grey). Centered horizontal line represents the median, whiskers represent min and max values. Two tailed Welch’s t-test ****p<0.0001. (F) (F’) (F’’) Proportions of GAF-ChIP-seq peaks that overlap diverse cis-regulatory regions in interphase only, mitotically retained and mitosis only GAF-ChIP-seq.

Motif search confirmed that GAF peaks are enriched in GAGAG motifs (Fig. 2d), and are centered inside the reads (Supplementary Fig. 2c). However, this consensus GAF binding site did not emerge as a significantly enriched motif in the small sample of GAF mitotic-only targets. We therefore did not analyze in depth this group of GAF targets. Moreover, there was a substantial degree of overlap (∼93.5%) when comparing our interphase GAF peaks with published GAF-ChIP-seq data from bulk 2-4h embryos (Koenecke et al. 2017). Thus, we established a pipeline, able to profile mitotic nuclei at a genomic scale, for the first time in a multicellular organism, in the absence of drug synchronization.

Interestingly, the number of GAGAG motifs differs between mitotically retained peaks and interphase only peaks. On average, mitotically retained peaks have 6.2 GAGAG repeats while interphase only bound targets show 2.9 number of motifs (Fig. 2e). Therefore, we conclude that loci with significant number of GAF binding sites are more likely to be bound during mitosis.

Moreover, *de-novo* motif search revealed that while some motifs are present on both categories (interphase only and mitotically retained), a combination of consensus binding sites is specifically enriched in mitotically retained peaks (e.g. Dorsal, Supplementary Fig. 2d). GAF mitotically retained targets might therefore be regulated by a distinct *cis*-regulatory logic than those from which GAF dissociates during mitosis.

To better characterize GAF-bound loci, we used existing genomic annotations of *cis*-regulatory modules (enhancers, promoters, and insulators) that were previously obtained from whole-genome profiling of the early *Drosophila* embryo (Hug et al. 2017; Nègre et al. 2010; Koenecke et al. 2017) or validated via reporter transgenes (Kvon et al. 2014) (see Methods, Fig. 2f-f’’). This stringent analysis revealed that the majority of GAF mitotically retained regions (65%) corresponds to *cis*-regulatory sequences (Fig. 2f’). This proportion is higher than the interphase only peaks (40%, Fig. 2f).

### Mitotically retained GAF marks accessible regions during zygotic genome activation

As GAF displays pioneering properties in many contexts (Moshe and Kaplan 2017; Gaskill et al. 2021; Fuda et al. 2015), we hypothesized that GAF could contribute to chromatin accessibility during mitosis. We therefore determined the degree of chromatin accessibility at GAF-bound loci by using available ATAC-seq data (Blythe and Wieschaus 2016). We observed that GAF mitotically retained regions are globally more open than GAF interphase only or mitotic only targets (Fig. 3a).

**Figure 3:**
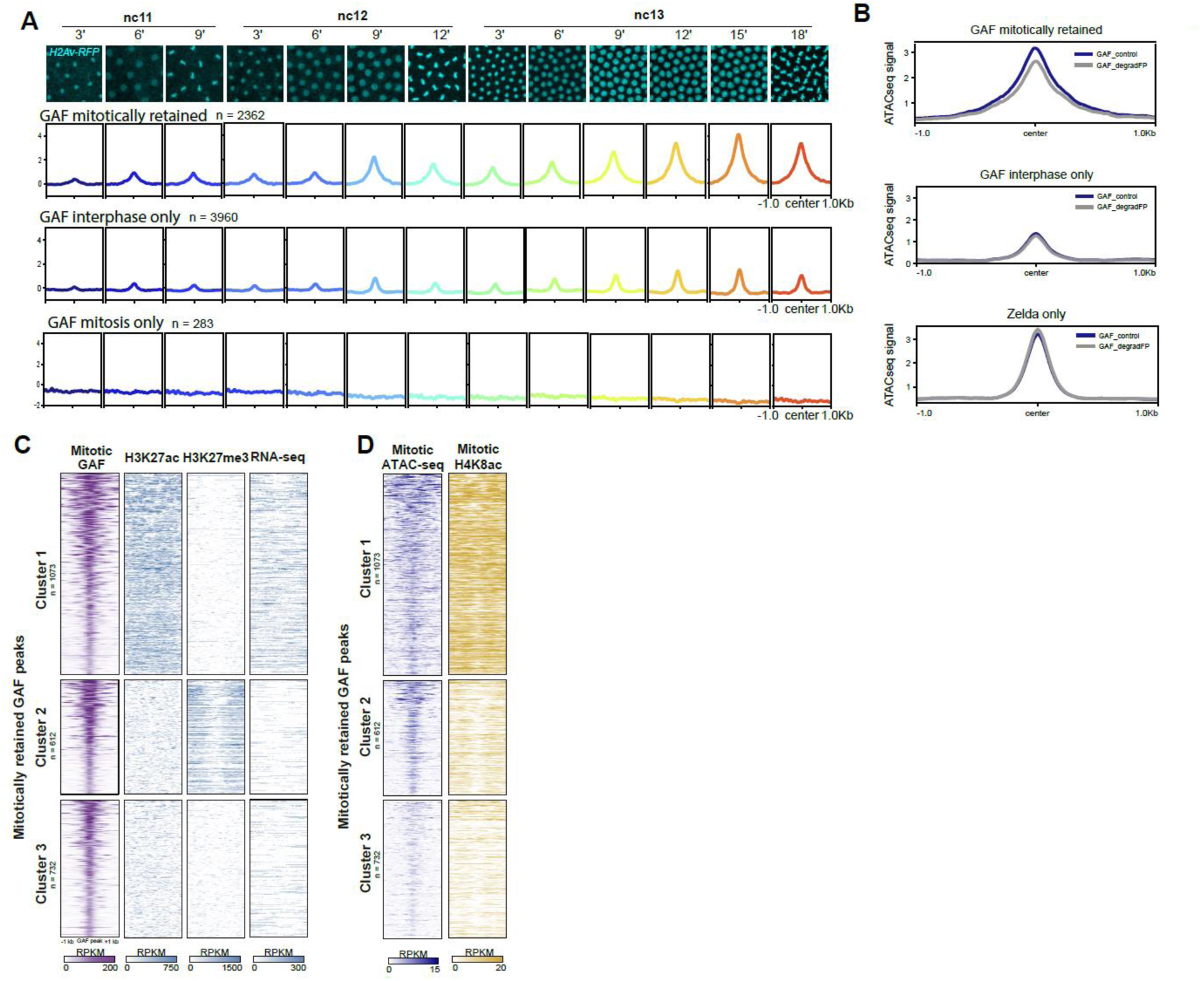
**Mitotically retained GAF loci become progressively accessible during Zygotic Genome Activation** (A) Metagene profiles of ATAC-seq signal (Blythe and Wieschaus 2016) centered at mitotically retained, interphase only and mitosis only GAF-ChIP-seq peaks across the indicated stages and represented by the time lapse images from a movie of H2Av-RFP embryos (cyan). (B) Metagene profiles of ATAC-seq signal in WT (GAF_control, dark blue) and GAF-depleted (GAF_degradFP, grey) embryos (Gaskill et al. 2021) on GAF mitotically retained, GAF interphase only and Zelda only regions. (C) Heatmaps of k-means clustered mitotically retained GAF peaks, based on H3K27ac and H3K27me3 ChIP-seq (Li et al. 2014) and RNA-seq (Lott et al. 2011). (D) Heatmaps representing the mitotic ATAC-seq signal (Blythe and Wieschaus 2016) (dark blue) and the ChIP-seq enrichment of H4K8ac in mitotic embryos at the clustered mitotically retained GAF peaks from (C).

More specifically, chromatin accessibility at mitotically retained regions encompasses larger regions than at loci bound by GAF only during interphase. This is in agreement with mitotically retained regions exhibiting a larger number of GAGA binding sites, potentially reflecting an enhanced number of bound GAF proteins able to foster nucleosome eviction (Supplementary Fig. 3a). Moreover, mitotically retained loci open gradually across developmental time windows and remain accessible during mitosis (Fig. 3a). Global chromatin accessibility at GAF mitotically retained targets is mostly linked to accessibility at *cis*-regulatory regions (Supplementary Fig. 3b).

We then asked whether chromatin accessibility at GAF mitotically retained regions required the presence of GAF. For this, we used ATAC-seq data performed on embryos where GAF levels were significantly reduced (Gaskill et al. 2021). From this dataset, we retrieved GAF bound loci for which accessibility was shown to be dependent on GAF. We found that the vast majority of these GAF-dependent regions (96%) correspond to GAF targets that we identified as mitotically retained (Supplementary Fig. 3d). Interestingly, targets depending on GAF for their accessibility mostly coincide with TSS and enhancer regions but don’t overlap TAD boundaries (Hug et al. 2017) (Supplementary Fig. 3c). Importantly, interphase GAF targets or Zelda-only bound targets (not bound by GAF) did not show such a dependency on GAF for their accessibility (Fig. 3b). Collectively, these results suggest that GAF retention at specific promoters and enhancers during mitosis may foster an accessible chromatin organization, which resists the overall compaction of the genome occurring during mitosis. However, other factors in addition to GAF are likely to foster chromatin accessibility during mitosis.

### GAF mitotic-bound regions are enriched with active and repressive histone marks

GAF is known to be present on both active and repressive chromatin regions (Adkins et al. 2006; Chetverina et al. 2021). We therefore assessed the chromatin landscape of GAF mitotically retained regions. For this purpose, we focused on embryonic ChIP-seq profiles of characteristic chromatin marks: H3K27ac for active chromatin state and H3K27me3 for the repressed chromatin state (Li et al. 2014), as well as RNA-seq signal from nc14 embryos (Lott et al. 2011). By clustering GAF mitotically retained regions, we partitioned GAF targets into three distinct clusters (Fig. 3c and Supplementary Fig. 3e). The first cluster (44% of mitotically retained GAF) corresponds to GAF mitotic peaks with significant enrichment in H3K27ac, depleted in H3K27me3 and with a high RNA-seq signal. In contrast, the second cluster (26% of mitotically retained GAF peaks) displayed enrichment for H3K27me3 concomitant with depletion in H3K27ac and low RNA-seq signal. The remaining GAF mitotic targets fall into a third cluster (30 % of mitotically retained GAF peaks), which displays no particular epigenetic features with our clustering analysis but shows significantly less chromatin accessibility (Fig. 3d). To examine if additional chromatin modifications mark could discriminate between these three GAF clusters we performed ChIP-seq on the acetylation of lysine 8 of histone H4 (H4K8ac). Indeed among the myriad of chromatin marks labeling active regions, H4K8ac is a prominent mark during initial reshaping of the genome during *Drosophila* ZGA (Li et al. 2014). We used our mitotic ChIP-seq method (Fig. 2a) to map H4K8ac in interphase and mitotic embryos genome-wide (Supplementary Fig. 4a-d). We observed that H4K8ac was particularly enriched in cluster 1 (Fig. 3d and Supplementary Fig. 4e).

Together, these results demonstrate that mitotic GAF retention occurs at genomic regions associated with both active or repressive chromatin states. We propose that the combinatorial action of GAF and histone marks, contribute to the selective mitotic bookmarking of active regions to propagate transcriptional programs across cellular divisions.

### GAF mitotic bookmarking is not associated with mitotic loops

Strictly speaking, mitotic occupancy by a TF can be envisaged as a mitotic bookmark only if it leads to a functional ‘advantage’ upon mitotic exit. Because chromatin loops between *cis*-regulatory regions were observed to be re-established by late anaphase/telophase in mammalian cells (Zhang et al. 2019) and since GAF is implicated in loop formation in *Drosophila* (Mahmoudi et al. 2002; Ogiyama et al. 2018), we asked if GAF mitotically bound loci could form loops during mitosis in the embryo. We first focused on a specific genomic region containing two developmental genes, *charybde* (*chrb*) and *scylla* (*scyl*), separated by 235kb and bound by GAF during both interphase and mitosis (Supplementary Fig. 5a-b). These early expressed genes were previously shown to form a long-range chromatin loop during early development (Ghavi-Helm et al. 2014).

We first confirmed that these loci are physically close and form a loop in nc14 by DNA FISH (Supplementary Fig. 5c). Interestingly, this proximity seems to be reinforced during nc14 progression (Supplementary Fig. 5c). However, while there is an overall genome compaction during mitosis, the distance between *scyl* and *chrb* is not different from that of a control locus, in mixed stages of mitosis (Supplementary Fig. 5c). To confirm this result, we examined two other loci using DNA FISH and assessed their potential looping across the cell cycle (Supplementary Fig. 5d). Both *snail* and *escargot* show GAF binding and the H4K8ac mark in interphase and mitosis. While these loci, form a loop in interphase nuclei, this long-range loop is not different from the control locus during mitosis (Supplementary Fig. 5e).

We therefore conclude that, at least for these regions, GAF mitotic binding is not associated with detectable stable mitotic DNA looping.

### The GAF bookmarked *scyl* gene harbors transcriptional memory

To test if GAF fosters rapid post-mitotic reactivation, we employed quantitative imaging on a selected GAF mitotically bound target, the zygotically expressed gene *scylla* (*scyl*). This gene is regulated by a promoter/proximal enhancer containing six GAGAG motifs, bound by GAF during interphase and mitosis (cluster 1 of mitotically retained loci) (Fig. 4a). To follow transcription dynamics with high temporal resolution, we utilized the MS2/MCP signal amplification method (Pichon et al. 2018) and quantitative imaging in living embryos. An array of 24X-MS2 repeats was inserted by CRISPR/Cas9 gene editing into the 3’UTR of *scyl* (Fig. 4a). MS2 reporter expression follows *scyl* endogenous expression (Fig. 4b and Supplementary Fig. 6a). Then, we monitored post-mitotic gene reactivation in nc14 in the ventral (Fig. 4b and Supplementary Fig. 6c) and dorsal side (Supplementary Fig. 6b). In both locations, post-mitotic activation was found to be relatively fast, with a lag time of only 7.5 min and 9 min to reach 50% of the full pattern of activation (t50) in the dorsal ectoderm and mesoderm, respectively (Supplementary Fig. 6d).

**Figure 4:**
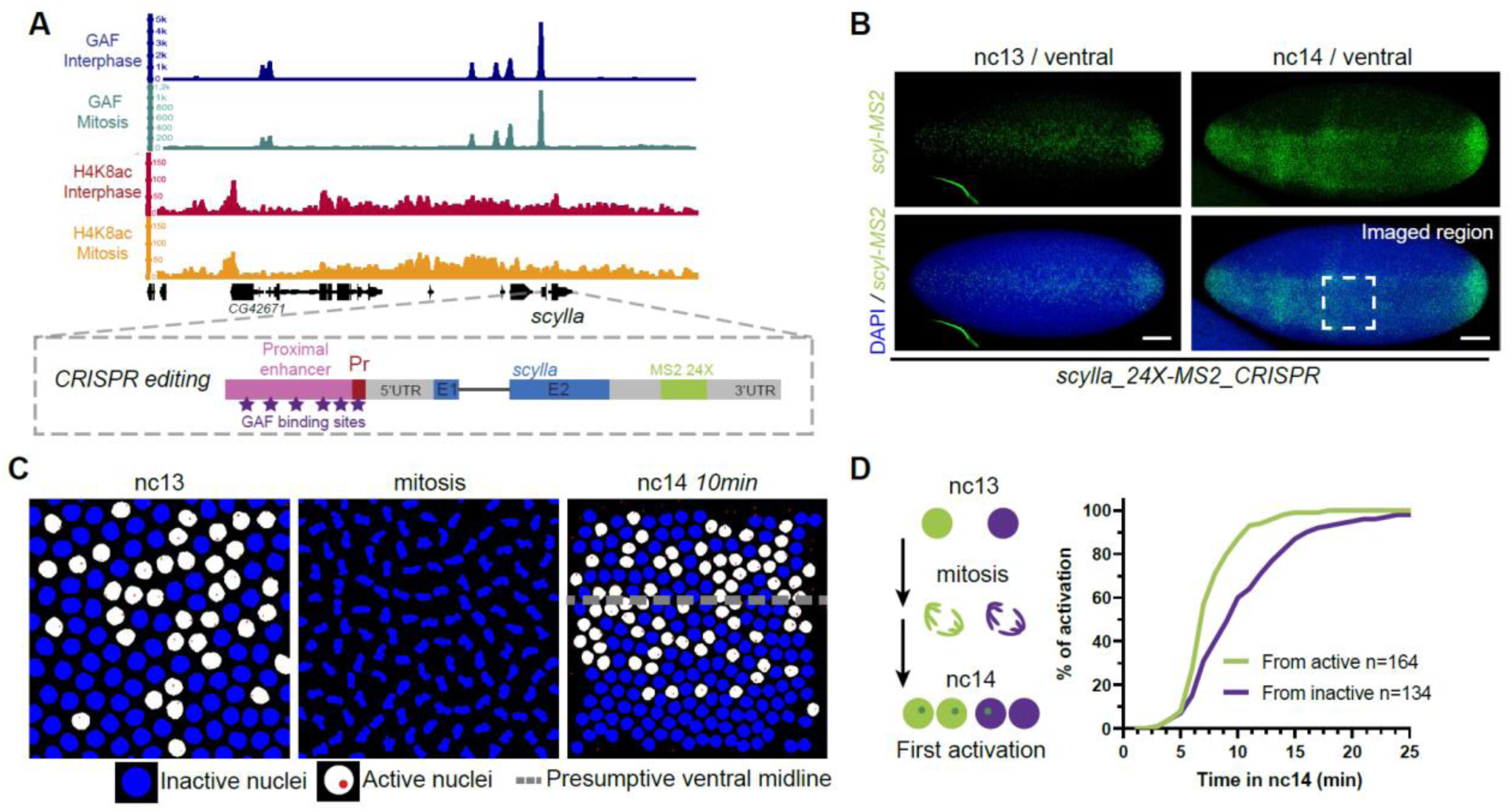
***scylla* gene harbors a transcriptional memory across mitosis.** (A) (Top) Genome browser image of interphase and mitotic GAF (dark blue and turquoise) and H4K8ac (red and orange) ChIP-seq signal at the scyl locus. (Bottom) Schematic of the 24X-MS2 tagging strategy of the scyl locus by CRISPR editing. (B) Maximum intensity projected Z-planes of confocal images from smiFISH with MS2 probes (green) counterstained with DAPI (blue) of scylla_24X-MS2_CRISPR/+ embryos in nc13 and nc14. Scale bars are 50µm. Dashed box represents the region considered for live imaging experiments. (C) Snapshots from a representative false-colored movie of scylla_24X-MS2_CRISPR/+ embryo carrying MCP-eGFP, H2Av-RFP. Active nuclei are represented in white and inactive nuclei in blue. Transcriptional sites are false colored in red. Dashed line represents the presumptive ventral midline. (D) Quantification of transcriptional memory for scyl gene. Left panel: schematic of the two populations of nuclei studied; those derived from active (in green) and those from inactive nuclei (purple). Right panel: cumulative activation of the first activated nuclei coming from active nuclei (green) and from inactive (purple). n=number of analyzed nuclei from 4 movies of 4 independent embryos.

In addition to this temporal information within a given interphase, live imaging of transcription in the context the fast-developing *Drosophila* embryo gives access to nuclei genealogy. We assessed whether the transcriptional status of mother nuclei (prior to division) influences that of their descendants (Trullo et al. 2020). Indeed, we have previously shown that within the mesoderm, descendants of active nuclei in nc13 activate transcription significantly faster than those arising from inactive nuclei, a bias named ‘transcriptional memory’ (Ferraro et al. 2016). However, this was shown in the context of reporter transgenes and has thus far never been demonstrated at an endogenous locus.

To assess the existence of transcriptional memory at an endogenously mitotically bookmarked locus, we imaged *scyl* expression in the mesoderm. Within this domain, the expression was stochastic in nc13 (Fig. 4b and Supplementary Fig. 6c, and Movie 3), allowing unambiguous discrimination between active and inactive mother nuclei prior to mitosis. By tracking the timing of activation for daughters arising from active mother nuclei compared to those coming from inactive mother nuclei (Fig. 4c), we observe a clear transcriptional memory bias (Fig. 4d and Supplementary Fig. 6e).

In order to test if this bias was due to a stronger activity of the *scyl* gene in nuclei coming from active mothers, we examined instantaneous intensities of transcriptional sites as they are directly correlated to the mRNA synthesis efficiency. Global transcriptional site intensities were similar in nuclei coming from active mothers compared to those coming from inactive mothers (Supplementary Fig. 6f).

### GAF knock-down delays post-mitotic transcriptional reactivation and affects transcriptional memory

To test whether GAF was involved in the establishment of transcriptional memory, we employed RNAi knock-down (KD) to reduce the pool of maternal GAF. As previous studies reported difficulties to successfully deplete maternal GAF using a specific set of Gal4 driver (Rieder et al. 2017), we decided to increase the efficiency of our depletion by combining two strong Gal4 drivers (mat-alpha-Gal4 and nanos-Gal4). The level of maternal GAF mRNA KD was estimated to be 88% by qRT-PCR and also confirmed by western blot (Supplementary Fig. 7a), creating a substantial embryonic lethality. However, in this genetic context a few embryos survived until gastrulation, albeit with clear mitotic and patterning defects for GAF targets genes (Movie 4, Supplementary Fig. 7b).

By quantifying post-mitotic reactivation timing of *scyl* in *RNAi-GAF* embryos, we observed a delay of ∼6min for t50 (Fig. 5a). However, transcription site intensities were not affected upon GAF reduction (Supplementary Fig. 7c). This global trend is also observed when following single nuclei behavior during nc14 (Supplementary Fig. 7c). We then compared the kinetics of activation in the two subpopulations (from active and from inactive) and found that the transcriptional memory bias was significantly reduced in *RNAi-GAF* embryos (Fig. 5b-c). Such a memory reduction does not occur upon maternal depletion of the pioneer factor Zelda (Dufourt et al. 2018).

**Figure 5:**
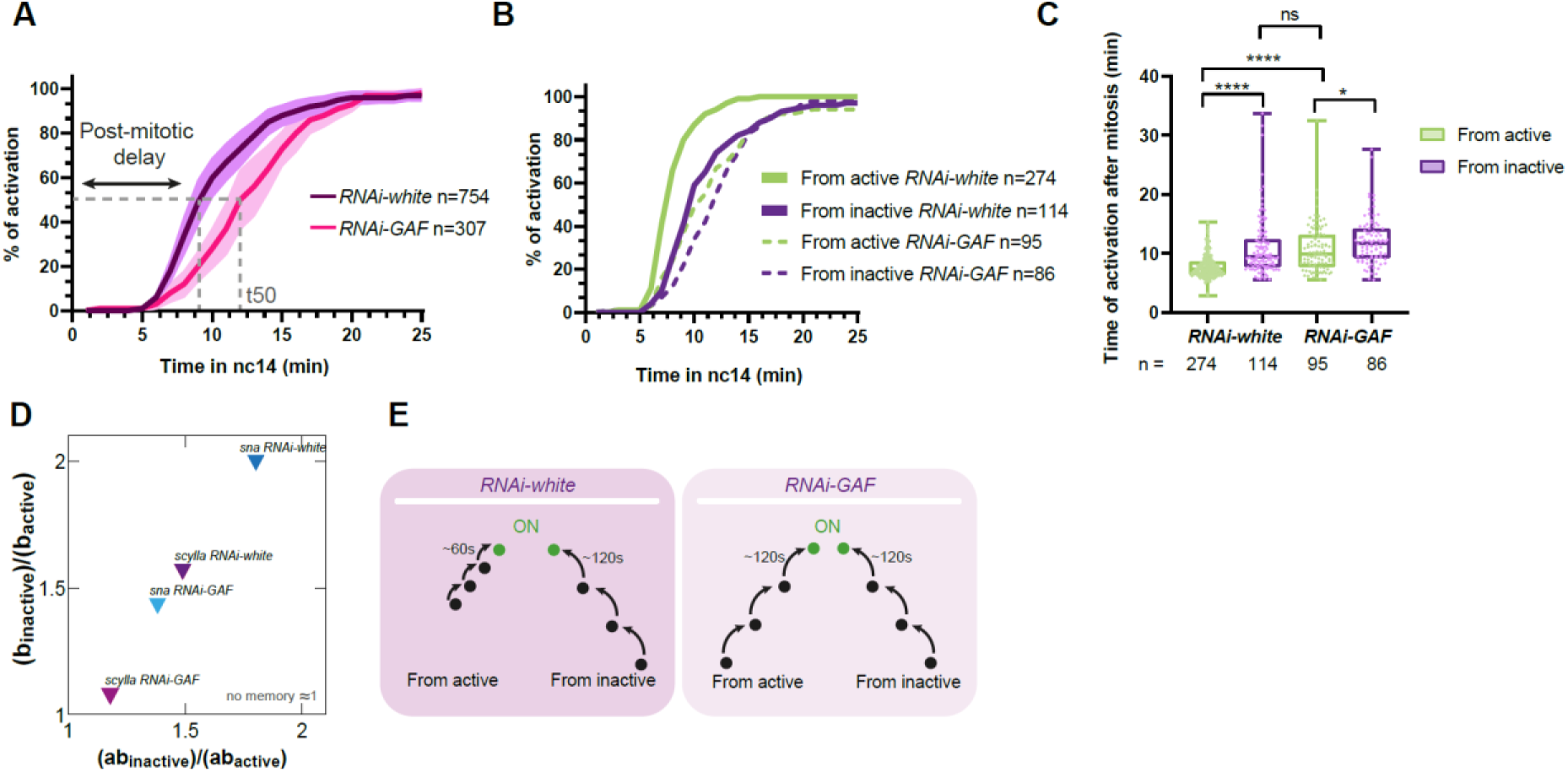
**GAF is required for transcriptional memory of *scylla*** (A) Quantification of transcriptional synchrony of scylla_24X-MS2_CRISPR/+ embryo after mitosis in RNAi-white (control, purple) and mat-alpha-Gal4/+; nos-Gal4/UASp-shRNA-GAF embryos (pink). Dashed line represents the t50 where 50% of the pattern is activated during nc14. Both of the two daughters derived from each nucleus are quantified. SEM are represented in light purple and light pink. n=number of nuclei analyzed from 4 movies of 4 independent embryos for each condition. (B) Cumulative activation of the first activated nuclei coming from active nuclei (green) and from inactive (purple) in RNAi-white embryos (control, solid curves) and RNAi-GAF scylla_24X- MS2_CRISPR/+ embryos (dashed curves). n=number of nuclei analyzed from 4 movies of 4 embryos. (C) Box plot representing the mean time of the first activation after mitosis of nuclei derived from active (green) and inactive (purple) nuclei in RNAi-white embryos and RNAi-GAF scylla_24X-MS2_CRISPR/+ embryos. Centered horizontal line represents the median. Two tailed Welch’s t-test ****p<0.0001, *p<0.05. (D) Ratios of parameter ‘b’ and ‘ab’ in subpopulations from inactive and active nuclei of scylla_24X-MS2_CRISPR/+ (purples) and snail-primary-enhancer_MS2 (blues) in RNAi-white or RNAi-GAF embryos. The parameter ‘a’ corresponds to the average number of transitions (provided by the sum of weighted probabilities) and the parameter ‘b’ to the time of each jump from one state to another. (E) Schematic of the proposed role of GAF in transcriptional memory. In the presence of GAF, nuclei derived from active nuclei have shorter ‘b’ length than those derived from inactive nuclei whereas in the absence of GAF, both have the same transition times.

Collectively, these data demonstrate that GAF controls the timing of transcriptional activation after mitosis and participates in the establishment of transcriptional memory. Moreover, this temporal effect is not due to a differential promoter activity between neighboring nuclei.

### Modeling GAF driven transcriptional memory

We analyzed the statistical distribution of the post-mitotic delay, defined as the lag time between the end of mitosis and the first activation in nc14. We have previously developed a simple mathematical model of memory, where this delay was modeled by a mixed gamma distribution (Dufourt et al. 2018) with two main parameters, the average number of rate-limiting transitions prior to reach the transcription active state (ON) (parameter ‘a’) and their durations (parameter ‘b’). Applying this mathematical model to our live imaging movies of *scyl* transcription dynamics in control (*RNAi-white*) and in GAF depleted embryos (*RNAi-GAF*) revealed that the ‘a’ parameter was comparable across genotypes (Supplementary Fig. 7d). However, upon GAF KD, the ‘b’ parameter significantly increased in nuclei coming from active mother nuclei (Supplementary Fig. 7d). Remarkably, this selective decrease in the ‘b’ parameter within a subpopulation was not observed upon Zelda depletion(Dufourt et al. 2018). In order to be able to compare the effect of various genotypes, subject to distinct *cis*- regulatory codes, we introduced a memory score defined by the ratio (ab_inactive_)/(ab_active_). A memory bias exists when this ratio is higher than 1. Using this metric, we observe that endogenous *scyl* exhibits a clear memory bias that vanishes upon GAF depletion (Fig. 5d).

Interestingly, a GAF-dependent memory bias was also observed with a second GAF-mitotically bound region (*sna-proximal-enhancer*, cluster 1, Fig. 2c’, see Methods) (Fig. 5d and Supplementary Fig. 7e). In all cases, we observe (ab_inactive_)/(ab_active_) ≈ b_inactive_/b_active_ (Fig. 5d), suggesting that the primary contribution to the memory bias comes from the transition duration ‘b’.

Collectively, these results suggest a model where transcriptional memory bias results from distinct epigenetic paths in nuclei where a given locus is bookmarked by GAF and in nuclei where the same locus is not bound by GAF (Fig. 5e). The preferential bookmarking of active nuclei by GAF could be explained by stochastic GAF binding.

## Discussion

We set out to determine how gene regulation by a transcription factor might be propagated through mitosis in a developing embryo. By using a combination of quantitative live imaging and genomics, we provide evidence that the pioneer-like factor GAF acts as a stable mitotic bookmarker during zygotic genome activation in *Drosophila* embryos.

Our results indicate that during mitosis, GAF binds to an important fraction of its interphase targets, largely representing *cis*-regulatory sequences of key developmental genes (Supplementary table 2). We noticed that GAF mitotically retained targets contain a larger number of GAGA repeats than GAF interphase only targets and that this number of GAGA repeats correlates with the broadness of accessibility. Multiple experiments, with model genes *in vitro* (e.g. *hsp70*, *hsp26*) or from genome-wide approaches clearly demonstrated that GAF contributes to generate nucleosome-free regions (Chetverina et al. 2021). The general view is that this capacity is permitted though the interaction of GAF with nucleosome remodeling factors as PBAP (SWI/SNIF), NURF (ISWI) (Judd et al. 2021) or FACT (Orphanides et al. 1998). Although not yet confirmed with live imaging, immuno-staining data suggest that NURF is removed during metaphase but re-engages chromatin by anaphase (Kwon et al. 2021). If the other partners of GAF implicated in chromatin remodeling are evicted during early mitosis, chromatin accessibility at GAF mitotic targets could be established prior to mitosis onset and then maintained through mitosis owing to the remarkable stability of GAF binding. However, we cannot exclude GAF interactions with other chromatin remodelers (e.g. PBAP) during mitosis and a scenario whereby mitotic accessibility at GAF targets would be dynamically established during mitosis thanks to the coordinated action of GAF and its partners.

We propose that the function of GAF as a mitotic bookmarker is possible because GAF has the intrinsic property to remain bound to chromatin for long periods (residence time in the order of minute). This long engagement of GAF to DNA is in sharp contrast with the binding kinetics of many other TF, such as Zelda or Bicoid in *Drosophila* embryos (Dufourt et al. 2018; Mir et al. 2017). Another particularity of GAF binding, contrasting with other TF, resides in the multimerization of its DNA binding sites as GAGAG repeats in a subset of its targets (76% of mitotically retained peaks display four or more repetitions of GAGAG motifs). Given the known oligomerization of GAF (Espinás et al. 1999) and as GAF is able to regulate transcription in a cooperative manner (Van Steensel et al. 2003), it is tempting to speculate that GAF cooperative binding on long stretches of GAGAG motifs may contribute to a long residence time.

Collectively, we propose that the combination of long residence time and the organization of GAF binding sites in the genome may allow the stable bookmarking of a subset of GAF targets during mitosis.

In this study, we also discovered that a combination of GAF and histone modification could be at play to maintain the chromatin state during mitosis. Indeed, mitotic bookmarking may also be supported by the propagation of histone tail modifications from mother to daughter cells. Work from mammalian cultured cells revealed widespread mitotic bookmarking by epigenetic modifications, such as H3K27ac and H4K16ac (Behera et al. 2019; Liu et al. 2017). Moreover, H4K16ac transmission from maternal germline to embryos has recently been established (Samata et al. 2020). In the case of GAF, we propose that the combinatorial action of GAF and epigenetic marks, possibly selected via GAF interacting partners, will contribute to the propagation of various epigenetic programs. It would be therefore interesting to employ our established mitotic ChIP method to survey the extent to which *cis*-regulatory regions exhibit different mitotic histone mark modifications during embryogenesis.

A key aspect of mitotic bookmarking is to relate mitotic binding to the rapid transcriptional activation after mitosis. Here we show that GAF plays a role in the timing of re-activation after mitosis. However, we note that GAF binding during mitosis is not the only means to accelerate gene activation. Indeed, we and others have shown that mechanisms such as enhancer priming by Zelda, paused polymerase or redundant enhancers contribute to fast gene activation (Bentovim et al. 2017). Moreover, a transcriptional memory bias can occur for a transgene not regulated by GAF (Ferraro et al. 2016). By modeling the transcriptional activation of the gene *scylla*, we reveal that GAF accelerates the epigenetic steps prior to activation selectively in the descendants of active nuclei. We propose a model where GAF binding helps in the decision-making of the post-mitotic epigenetic path. In this model, mitotic bookmarking by GAF would favor an epigenetic path with fast transitions after mitosis (Fig. 5e). In the context of embryogenesis, bookmarking would lead to the fast transmission of select epigenetic states and may contribute to gene expression precision.

Interestingly, GAF vertebrate homolog (vGAF/Th-POK) has recently been implicated in the maintenance of chromatin domains during zebrafish development (Matharu et al. 2021). We therefore suspect that GAF action as a stable bookmarking factor controlling transcriptional memory during *Drosophila* ZGA might be conserved in vertebrates.

## Acknowledgments

We are grateful to G.Cavalli, B. Schuttengruber, F. Juge, F.Bantignes and J.Chubb for sharing antibodies, advise on DNA FISH and/or insightful discussions.

We thank member of the Lagha lab, Cyril Esnault, Virginia Pimmett and Etienne Schwob for their critical reading of the manuscript. We acknowledge M. Dejean and M. Goussard for technical assistance. We acknowledge the Montpellier Ressources Imagerie facility (France-BioImaging). Funding: M.B. is a recipient of an FRM fellowship. This work was supported by the ERC SyncDev starting grant to M.L. M.L., J.D., and C.F. are sponsored by CNRS. O.R. acknowledges support from the French National Research Agency (ANR-17-CE40-0036, project SYMBIONT).

## Author contributions

M.L. conceived the project. M.L., M.B. and J.D. designed the experiments. J.D., M.B., H.L-H., M. Lam. and H.F-G. performed experiments. A.T. developed software. C.F., M.B., and J.D. performed kinetic analysis. M.G. and M.H. provided the GAF-GFP strain. O.R. performed mathematical modeling, machine learning and interpreted data. G.H., M.B., A.Z-E-A., M.M. and J-C.A. performed the bioinformatics analysis. M.L. and M.B. wrote the manuscript. All authors discussed, approved, and reviewed the manuscript.

## Material and Methods

### Fly stocks, handling and genetics

The *yw* stock was used as a wild type. The germline driver *nos-Gal4:VP16*(BL4937) was previously recombined with a *MCP-eGFP-His2Av-mRFP* fly line (Dufourt et al. 2018). RNAi were expressed after crossing this recombinant for live imaging (or *nos-Gal4:VP16* for fixed experiments) with *mat-alpha-Gal4* (BL7063), then with *UASp-shRNA-w* (BL35573) or *UASp-shRNA-GAF* (BL41582). Virgin females expressing RNAi, MCP-GFP-His2Av-mRFP and both Gal4 constructs were crossed with MS2 containing CRISPR alleles or transgene-containing males. All experiments were done at 21 °C except RNAi experiments which were done at 25 °C. The C-terminal tagged version of GAF-sfGFP was obtained by CRISPR/Cas9 (Gaskill et al. 2021).

### Cloning and transgenesis

The *snail-primary-enhancer_MS2* transgene was obtained by amplification of the *sna* endogenous promoter and primary enhancer using the primers listed in Supplementary table 1. The 128XMS2 tag was inserted immediately upstream of the yellow reporter gene sequence of the pbphi-yellow plasmid (Ferraro et al. 2016). The transgenic construct was inserted in the VK0033 landing site (BL9750) using PhiC31 targeted insertion (Venken et al. 2006).

The homology arms for the recombination template for CRISPR/Cas9 editing of *scyl* gene to generate *scyl_24X-MS2_CRISPR* were assembled with NEBuilder® HiFi DNA Assembly Master Mix (primers listed in Supplementary table 1) and inserted into pBluescript opened *SpeI/AscI* (for the 5’ homology arm) or *XmaI/NheI* (for the 3’ homology arm) containing the 24X-MS2 (as in (Dufourt et al. 2018)) inserted after *Not1* digestion. Guide RNA (Supplementary table 1) were cloned into pCFD3-dU6:3gRNA (Addgene 49410) digested by BbsI using annealed oligonucleotides (Integrated DNA Technology™). The recombination template and guide RNA plasmids were injected into BDSC#55821 (BestGene Inc.). Transformant flies were screened using a dsRed marker inserted downstream of the 3’UTR of the genes.

### Fluorescence recovery after photobleaching

Fluorescence recovery after photobleaching (FRAP) in embryos at nc14 was performed on a Zeiss LSM880 using a 40 × /1.3 Oil objective and a pinhole of 84 μm. Images (256 × 128 pixels, 16bits/pixel, zoom 6x) were acquired every ≈ 53 ms for 1200 frames. GFP was excited with an Argon laser at 488 nm and detected between 492–534 nm. Laser intensity was kept as low as possible to minimize unintentional photobleaching. A circular ROI (12 × 12 pixels) 0.138 µm/pixel, was bleached using two laser pulses at maximal power during a total of ≈ 110 ms after 10 frames. To discard any source of fluorescence intensity fluctuation other than molecular diffusion, the measured fluorescence recovery in the bleached ROI region (*I*_bl_) was corrected by an unbleached ROI (*I*_unbl_) of a neighbor’s nucleus and another ROI outside of the nucleus (*I*_out_) following the simple equation:

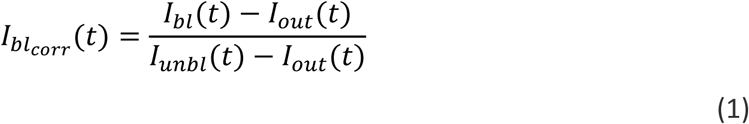

The obtained fluorescence recovery was then normalized to the mean value of fluorescence before the bleaching i.e.

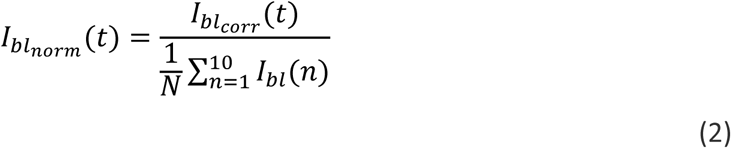

Analytical equations used to fit the fluorescence recovery was chosen with two exchanging population on the first 1100 frames: we started from the analytical expression developed in the Supplementary Equation 35 of (Michelman-Ribeiro et al. 2009).

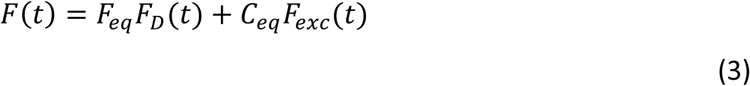

with C_eq_ defined as above and F_eq_ = k_off_ / (k_off_ + k*_on_). F_D_(t) is the fluorescence recovery due to diffusion and F_exc_(t) the fluorescence recovery due to exchange.

Since we used a Gaussian shape illumination profile, F_D_(t) is defined using a slightly modified version of the analytical equation of the 20th order limited development of the Axelrod model for Gaussian profile illumination and diffusion (Escoffre et al. 2014; Axelrod et al. 1976):

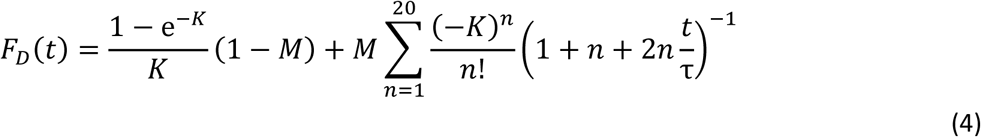

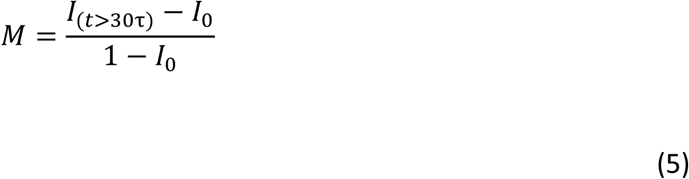

where K is a constant proportional to bleaching deepness, M is the mobile fraction and is the half time of recovery. To minimize the effect of mobile fraction on C_eq_, M was kept between 0.9 and 1.1.

Diffusion coefficients of the different molecules were determined according to

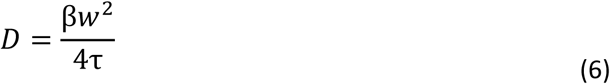

with w the value of the radius at 1/e^2^ of the Gaussian beam (in our case, w=0.83µm) and β a discrete function of K tabulated in (Yguerabide et al. 1982).

F_exc_(t) is defined as in (Michelman-Ribeiro et al. 2009), slightly modified with respect to the Gaussian illumination, leading to the following equation:

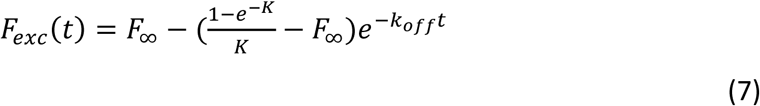

with K defined as previously.

### Fluorescence correlation spectroscopy

Florescence correlation spectroscopy (FCS) experiments were performed on a Zeiss LSM780 microscope using a 40x/1.2 water objective. GFP was excited using the 488 nm line of an Argon laser with a pinhole of 1 airy unit. Intensity fluctuation measured for 10 s were acquired and auto-correlation functions (ACFs) generated by Zen software were loaded in the PyCorrFit program (Müller et al. 2014). Multiple measurements per nucleus in multiple nuclei and embryos at 20 °C were used to generate multiple ACF, used to extract parameters. The FCS measurement volume was calibrated with a Rhodamine6G solution (Dertinger et al. 2008) using *D*_f_ = 414 μm^2^.s^−1^. Each time series was fitted with the following generic equation:

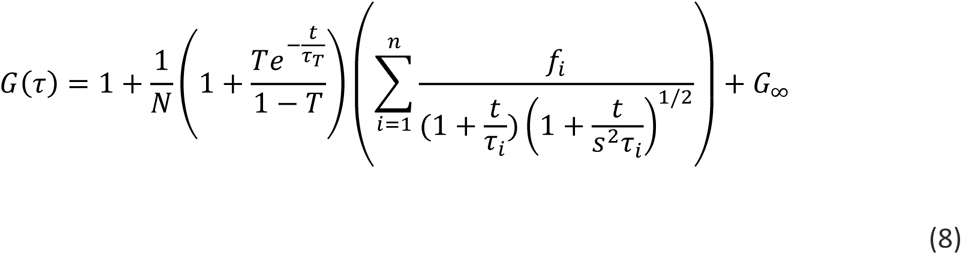

Using n=2 in our fit and where N is the total number of molecules, T is the proportion of the fluorescent molecules N in the triplet state with a triplet state lifetime *τ*_T_(constrained below 10µs in our fit), *f_i_* is the proportion of each different diffusing species 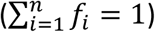 with a diffusion time τ_i_ = *w*^2^*_xy_* / 4 *D* and *s^2^* = *w_z_* / *w_xy_*. We also introduced a G_∞_ value to account for long time persistent correlation during the measurements.

### Immunostaining and RNA in situ hybridization

A pool of 0-4h after egg-laying (AEL) or 2-4h AEL embryos were dechorionated with bleach for 3 min and thoroughly rinsed with H_2_O. They were fixed in 1:1 heptane:formaldehyde-10% for 25 min on a shaker at 450 rpm; formaldehyde was replaced by methanol and embryos were shaken by hand for 1 min. Embryos that sank to the bottom of the tube were rinsed three times with methanol. For immunostaining, embryos were rinsed with methanol two times and washed three times 3 min with PBT (PBS 1x 0.1% triton). Embryos were incubated on a wheel at room temperature for 30 min in PBT, then for 20 min in PBT 1% BSA, and at 4 °C overnight in PBT 1% BSA with primary antibodies. Embryos were rinsed three times, washed twice for 20 min in PBT, then incubated in PBT 1% BSA for 20 min, and in PBT 1% BSA with secondary antibodies for 2 h at room temperature. Embryos were rinsed three times then washed three times in PBT for 10 min. DNA staining was performed using DAPI at 0.5μg/ml. Primary antibody dilutions for immunostaining were mouse anti-GFP (Roche IgG1κclones 7.1 and 13.1) 1:200; rabbit anti-GAF (gift from Dr. G.Cavalli) 1:250. Secondary antibodies (anti-rabbit Alexa 488-conjugated (Life Technologies, A21206); anti-mouse Alexa 488-conjugated (Life Technologies, A21202); anti-rabbit Alexa 555-conjugated (Life Technologies, A31572)) were used at a dilution 1:500. Fluorescent *in situ* hybridization (FISH) was performed as described in (Dufourt et al. 2018). The dixogygenin-MS2 probe was obtained as (Dufourt et al. 2018) by *in vitro* transcription from a pBluescript plasmid containing the 24X-MS2 sequences, isolated with BamH1/BglII enzymes from the original Addgene MS2 plasmid (#31865). *snail* probe generation was described in (Dufourt et al. 2018). Primary and secondary antibody for FISH were sheep anti-digoxigenin (Roche 11333089001) 1:375; mouse anti-biotin (Life technologies, 03–3700) 1:375; anti-mouse Alexa 488-conjugated (Life Technologies, A21202) and anti-sheep Alexa 555-conjugated (Life Technologies, A21436) 1:500. Mounting was performed in Prolong® Gold.

Images in Supplementary Fig. 1a represent a maximum intensity projection of a stack of 3 z-planes (≈1 μm). Images in Supplementary Fig. 1b represent a single Z-plane. Images in Fig. 1a represent a maximum intensity projection of a stack of 9 z-planes (≈4,5 μm).

### Single molecule fluorescence *in situ* hybridization (smFISH)

Embryos were fixed as in the previous section, then washed 5 min in 1:1 methanol:ethanol, rinsed twice with ethanol 100%, washed 5 min twice in ethanol 100%, rinsed twice in methanol, washed 5 min once in methanol, rinsed twice in PBT-RNasin (PBS 1x, 0.1% tween, RNasin® Ribonuclease Inhibitors). Next, embryos were washed 4 times for 15 min in PBT-RNasin supplemented with 0.5% ultrapure BSA and then once 20 min in Wash Buffer (10% 20X SCC, 10% Formamide). They were then incubated overnight at 37 °C in Hybridization Buffer (10% Formamide, 10% 20x SSC, 400 µg/ml *E. coli* tRNA (New England Biolabs), 5% dextran sulfate, 1% vanadyl ribonucleoside complex (VRC) and smFISH Stellaris probes against *sna* coupled to Quasar 670 and/or FLAP probes). FLAP-probes against 24X-MS2 and *scyl* were prepared by duplexing 40 pmol of target-specific probes with 100 pmol FLAP-Cy3 oligonucleotides and 1X NEBuffer™ 3 for 3 min at 85 °C, 3 min at 65 °C and 5 min at 25 °C and kept on ice until use. Probe sequences are listed in Supplementary table 1.

Embryos were washed in Wash Buffer at 37 °C and in 2x SCC, 0.1% Tween at room temperature before being mounted in ProLong® Gold antifade reagent. Images were acquired using a Zeiss LSM880 confocal microscope with an Airyscan detector in SR mode with a 40x Plan-Apochromat (1.3 NA) oil objective lens or a 20x Plan-Apochromat (0.8NA) air objective lens. Images were taken with 1024 x 1024 pixels and Z-planes 0.5μm apart. GFP was excited using a 488 nm laser, Cy3 were excited using a 561 nm laser, Quasar670 was excited using a 633 nm laser.

Images in Fig. 4b and Supplementary Fig. 6a-b represent a maximum intensity projection of a stack of 15 z-planes (≈9.5 μm).

### H3S10ph immunostaining and mitotic embryos sorting

A pool of 1h30-2h30 AEL embryos were fixed as for immunostaining except the fixation was in 1:1 heptane:1.8% formaldehyde/1X PBS (Thermo Scientific 28906) for exactly 10 min shaking at 450 rpm. Then embryos were rapidly quenched with 125 mM glycine PBS-1x and shaken for 1 min by hand. An anti-phospho-Histone H3 (Ser10) antibody (Cell Signalling #9701) was used at a dilution 1:200. Anti-mouse Alexa 488-conjugated (Life technologies, A21202) was used as a secondary antibody at a dilution 1:500. Embryos were kept in PBT until sorting. Sorting was done using a COPAS SelectInstrument (Biometrica) with the following parameters: sorting limit low: 1, high: 256; PMT control: Green 650, Yellow 425 and Red 800. A restricted area of sorting (with the highest green signal) was selected representing ≈ 8% of the total population. A container was placed at the output of the non-selected embryos in order to re-pass them through the sorter to collect non-green embryos corresponding to interphase embryos. Right after the sorting, embryos were manually checked under a Leica Z16 APO macroscope by placing them on a glass cup and using Drummond Microcaps® micropipettes to remove mis-sorted embryos individually. 1000 embryos per tube were then dried by removing the PBT and kept at -80 °C.

### Chromatin Immuno-Precipitation and library preparation

1000 embryos were homogenized in 1 ml of Buffer A (60 mM KCl,15 mM NaCl, 4 mM MgCl2, 15 mM HEPES (pH 7.6), 0.5% Triton X100, 0.5 mM DTT, 10 mM Sodium Butyrate and Protease Inhibitors Roche 04693124001) using a 2 ml Dounce on ice. The solution was then centrifuged 4 min at 2000g at 4 °C. Supernatant was removed and 1 ml of Buffer A was added and this was repeated two times with Buffer A and once with Lysis Buffer without SDS (140 mM NaCl, 15 mM HEPES (pH 7.6), 1 mM EDTA (pH 8), 0.5mM EGTA, 1% Triton X100, 0.5 mM DTT, 0.1% Sodium Deoxycholate, 10 mM Sodium Butyrate and Protease Inhibitors). The pellet was resuspended in 200 µl of Lysis Buffer with 0.1% SDS and 0.5% N-Laurosylsarcosine and incubated 30 min at 4 °C on a rotating wheel. Sonication was done with a Bioruptor® Pico sonication device with 30 sec ON/30 sec OFF cycles for 6-7 min for interphase and 8-9 min for mitotic chromatin. Sonicated chromatin was then centrifuged 5 min at 14000 rpm at 4 °C. The chromatin was then diluted in 1 ml of Lysis Buffer.

Dynabeads® M-270 Epoxy (Invitrogen Life Technologies^TM^, 14301) were prepared in order to directly crosslink antibodies to the beads (anti-GAF, gift from G. Cavalli, or anti-H4K8ac, abcam 15823), avoiding cross reaction with the H3S10ph antibody, following manufacturer protocol. Prior to this, anti-GAF was purified using NAb™ Protein A/G Spin Kit (ThermoScientific). Once the magnetic beads were cross-linked, chromatin was incubated over night at 4 °C on a rotating wheel. Then, beads were washed 7 min at 4 °C once in Lysis Buffer, once in FAT Buffer (1 M TrisHCl pH 8, 0.5 M EDTA pH 8, SDS 10%, 5 M NaCl, 10% Triton), once in FA Buffer (1 M HEPES, pH 7.0-7.6, 5 M NaCl, 0.5 M EDTA pH8, Triton X-100 – 10% NaDeoxycholate) once in LiCL Buffer (1 M Tris-HCl pH 8, 4 M LiCl, 10% Nonidet-P40-Nonidet, 10% NaDeoxycholate and protease inhibitors) and twice in TE (10 mM Tris-HCl pH 8, 0.1 mM EDTA). Elution was done in elution Buffer 1 (10 mM EDTA, 1% SDS, 50 mM Tris-HCl pH 8) for 30 min at 65 °C at 1300 rpm. Eluted chromatin was removed and a second elution step with Elution Buffer 2 (TE, 0.67% SDS) was performed. The two elutions were pooled. Chromatin was then reverse-crosslinked by heating onvernight at 65 °C. Next, chromatin was incubated 3 h at 50 °C with ProteinaseK (Thermo Scientific^TM^ EO0491) and RNAseA (Thermo Scientific^TM^ EN0531). DNA was then extracted with phenol/chloroform purification. Biological duplicates were performed for each sample.

Libraries were then prepared using the NEBNext UltraII DNA Library Prep Kit for Illumina, following the manufacturer’s instructions. Sequencing was performed on Illumina HiSeq 4000 on pair-end 75 bp.

### ChIP-seq analysis

Both reads from ChIP-seq and Input experiments were trimmed for quality using a threshold of 20 and filtered for adapters using Cutadapt (v1.16). Reads shorter than 30 bp after trimming were removed. Reads were mapped to *Drosophila melanogaster* genome (dm6 release) using Bowtie2 (Langmead and Salzberg 2012). Aligned sequences were processed with the R package PASHA to generate the used wiggle files. Pasha elongates *in sillico* the aligned reads using the DNA fragment size estimated from paired-reads. Then, the resulting elongated reads were used to calculate the coverage score at each nucleotide in the genome. Wiggle files representing average enrichment score every 50 bp were generated. In order to normalize the enrichment scores to reads per million, we rescaled the wiggle files using PASHA package. Besides, in order to reduce the over-enrichment of some genomic regions due to biased sonication and DNA sequencing, we subtracted from ChIP sample wiggle files the signal present in Input sample wiggle files. The Rescaled and Input subtracted wiggle files from biological replicate were then used to generate the final wiggle file representing the mean signal.

In order to call the enriched peaks from the final wiggle files, we used *Thresholding* function of the Integrated Genome Browser (IGB) to define the signal value over which we consider a genomic region to be enriched compared to background noise (*Threshold*). We used also the minimum number of consecutive enriched bins to be considered an enriched region (*Min.Run*) as well as the minimum gap above which two enriched regions were considered to be distinct (*Max.Gap*). The three parameters were then used with an in-house script that realizes peak calling by using algorithm employed by *Thresholding* function of IGB.

Peaks calling was done with a threshold of 100 for GAF-ChIP-seq and 22 for H4K8ac-ChIP-seq, minimum run of 50 bp and maximum gap of 200bp. Interphase only peaks correspond to peaks from interphase ChIP-seq with no overlap with peaks from mitotic ChIP-seq. Mitotically retained correspond to interphase peaks with an overlap (min 1 base pair) with peaks from mitotic ChIP-seq. Mitotic only peaks correspond to peaks from mitotic ChIP-seq with no overlap with peaks from interphase ChIPseq.

Motif search was done with the MEME ChIP tool (MEME suite 5.1.1).

Peaks were considered as promoter if overlap with the region defined by 100 bp aroud TSS. Peaks were considered as enhancers if overlapping with identified enhancer (Kvon et al. 2014) and/or overlapping with a H3K27ac peak (Koenecke et al. 2016).

ATAC-seq data are from (Blythe and Wieschaus 2016)(GSE83851). Wig files were converted to BigWig using Wig/BedGraph-to-bigWig converter (Galaxy Version 1.1.1). ATACseq mean signal was then plotted on regions of interest (mitotically retained peak coordinates and Interphase only coordinates) using computeMatrix by centering ATAC-seq signal to the center of the regions (and +/- 1 kb) followed by plotProfile (Galaxy Version 3.3.2.0.0).

Mitotically retained GAF peaks were subdivided by *k*-means clustering based on chromatin state (H3K27ac and H3K27me3 ChIP-seq (Li et al. 2014)) and transcriptional status (nc14 RNA-seq (Lott et al. 2011)) using deepTools (Ramírez et al. 2016). Peaks were partitioned into three clusters: cluster 1, n=1073, cluster 2, n=612 and cluster 3, n=732. To further characterize mitotically retained clusters we plotted heatmaps using deepTools (Ramirez et al., 2016) for publicly available ChIP-seq data for H3K27ac (Li et al. 2014), H3K27me3 (Li et al. 2014) and ATAC-seq (Blythe and Wieschaus 2016).

GAF bound loci for which accessibility are dependent on GAF were taken from (Gaskill et al. 2021).

### Live imaging

Movies of *His2Av-mRFP; sfGFP-GAF_CRISPR* (related to Movie 1, Movie 2 and Fig. 1b) were acquired using a Zeiss LSM880 with confocal microscope in fast Airyscan mode with a Plan-Apochromat 40x/1.3 oil objective lens. GFP and mRFP were excited using a 488 nm and 561 nm laser respectively with the following settings: 256 x 256-pixel images, 15 z-planes 1 μm apart and zoom 4x, resulting in a time resolution of 9.5 sec per Z-stack. Average intensity profiles were measured for histones, nucleoplasmic GAF and cytoplasmic GAF from three movies of embryos transitioning from nc13 into nc14. An automatic tracking of maximum intensity projected images fluorescence was done using a home-made software as in (Dufourt et al. 2018). First a detection of nuclei is made using His2Av-mRFP allowing the monitoring of histone intensity fluctuation, then a mask of His2Av-mRFP detected nuclei was projected on the sfGFP-GAF channel allowing the recovery of sf-GFP-GAF present on histones. Finally, five ROI in each movie corresponding to cytoplasmic regions were tracked for sfGFP-GAF intensity in the cytoplasm.

Movies of *MCP-eGFP-His2Av-mRFP*>*snail-primary-enhancer_MS2/+* embryos (related to Supplementary Fig. 7e) were acquired using a Zeiss LSM780 with confocal microscope with a Plan-Apochromat 40x/1.3 oil objective lens. GFP and mRFP were excited using a 488 nm and 561 nm laser respectively with the following settings: 512 x 512-pixel images, 21 z-planes 0.5μm apart and zoom 2.1x, resulting in a time resolution of 22 sec per frame. Movies were subjected to filtering steps to track transcription foci as 128XMS2 loops result in signal retention during mitosis.

Movies of *MCP-eGFP-His2Av-mRFP*>*scyl_MS2_CRISPR/+* in *RNAi-White* and *RNAi-GAF* background (related to Movie 3 and 4 and to Fig. 4 and 5 and Supplementary Fig. 6c-f and 7c) were acquired using a Zeiss LSM880 with confocal microscope in fast Airyscan mode with a Plan-Apochromat 40x/1.3 oil objective lens. GFP and mRFP were excited using a 488 nm and 561 nm laser respectively with the following settings: 552 x 552-pixel images, 21 z-planes 0.5 μm apart and zoom 2.1x, resulting in a time resolution of 5.45 sec per frame. As we observed that GAF knock-down was not complete (some *RNAi-GAF* embryos gastrulate and develop), movies showing visible developmental defects, such as nuclear dropout, anaphase bridges or failure to gastrulate, were kept for analysis.

### Memory movies analysis

Movies were analyzed using Mitotrack (Trullo et al. 2020) as in (Dufourt et al. 2018) with newly implemented tools to filter mitotic 128XMS2 foci in movies of *MCP-eGFP-His2Av-mRFP*>*snail-primary-enhancer_MS2/+* embryos (mitotic foci are now detected with the 24MS2 array). Briefly, using a custom-made algorithm developed in Python^TM^ and implemented in the MitoTrack software, nuclei were segmented and tracked in 2D, working on the maximum intensity projected stack. Transcription spots were detected and tracked in 3D. All the spots present during mitosis were removed in the successive cycle such that only *de novo* appearing MS2 punctae were analyzed.

For intensity analysis (related to Supplementary Fig. 6f and 7c) the intensity of detected spots was collected for each frame to study the transcriptional intensity behavior throughout nuclear cycle 14. Nuclei coming from inactive and nuclei coming from active were separated for Supplementary Fig. 6f and pooled for Supplementary Fig. 7c.

### Mathematical modeling of mitotic memory

We are interested in the post-mitotic delay, defined as the time needed for post-mitotic transcription (re)activation. We model this time as the sum of two variables as in (Dufourt et al. 2018):

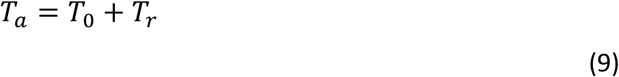

where *T*_0_ is a deterministic incompressible lag time, the same for all nuclei, and *Tr* is a random variable whose value fluctuates from one nucleus to another. The decomposition in Eq. 9 can be justified by the experimental observation that all the reactivation curves (Fig. 5a, Supp7c) start with a nonzero length interval during which no nuclei are activated. Furthermore, *Tr* is defined such that it takes values close to zero with non-zero probability. This property allows us to set *T*_0_ to the instant when the first nucleus initiates transcription, in order to determine *Tr*. The random variable *Tr* is modeled using a finite state, continuous time, Markov chain. The states of the process are *A*_1_, *A*_2_,…, *A*_n−1_, *A*_n_. The states *A*_i_, 1 ≤ *i* ≤ *n* − 1 are OFF, i.e. not transcribing. The state *A*_n_ is ON, i.e. transcribing. Each OFF state has a given lifetime, defined as the waiting time before leaving the state and going elsewhere. Like in (Dufourt et al. 2018), we considered that each of the states has the same lifetime denoted *τ*. Also, the transitions are considered linear and irreversible: in order to go to *A*_i+1_ one has to visit *A*_i_ and once there, no return is possible. The time *Tr* is the time needed to reach *A*_n_ starting from one of the OFF states. The predictions of these models were compared to the empirical survival function *S_exp_(t)* defined as the probability that *Tr>t*, obtained using the Meier-Kaplan method from the values *Tr* for all the analyzed nuclei. Following Occam’s razor principle, we based our analysis on the simplest model that is compatible with the data, which is a model with *n=4* and homogeneous lifetimes. For this model the theoretical survival function is a mixture of Gamma distributions:

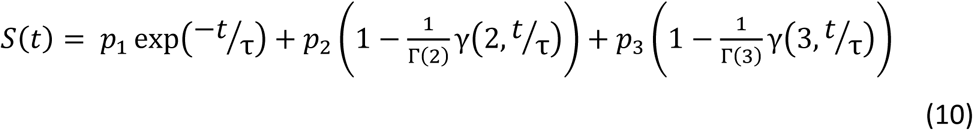

where γ, Γ are the complete and incomplete gamma functions and *p_1_*, *p_2_*, *p_3_* (satisfying *p_1_*+*p_2_*+*p_3_*=1) are the probabilities to reach ON after one, two or three jumps, respectively. We have also tested more complex models, with uneven lifetimes, more states and therefore, more parameters. However, the complex models did not provide a sensibly better fit with data and generated overfitting identified as large parametric uncertainty.

The parameters ‘a’ and ‘b’, summarizing the statistics of the post-mitotic reactivation are defined as

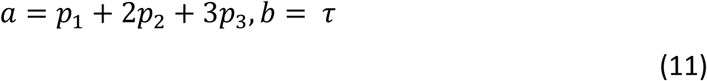

Data and materials availability: GSE180812. These data are private until the revision process.

## Supplementary information

### qRT-PCR in *RNAi* embryos

Total RNA from 0-2h AEL *RNAi-white* or *RNAi-GAF* driven by *nos-Gal4* and *mat-alphaTub-Gal4* embryos was extracted with TRIzol following the manufacturer’s instructions. RNA was DNase-treated. 1 μg of RNA extracted from ∼300 embryos per replicate was reverse transcribed using SuperScript IV and random primers. Quantitative PCR analyses were performed with the LightCycler480 SYBR Green I Master system (primers used listed in Supplementary table 1, targeting both isoforms of *GAF*). RNA levels were calculated using the *RpL32* housekeeping gene as reference and not bound by GAF according to the GAF-ChIP-seq. Each experiment was performed with biological triplicates and technical triplicates.

### Western blot analysis

Fifty embryos from *RNAi-white* or *RNAi-GAF* driven by *nos-Gal4* and *mat-alphaTub-Gal4* 0-2h AEL embryos were collected and crushed in 100μl of NuPAGE™ LDS sample buffer and reducing agent. Samples were heated 10min at 70°C, and the volume-equivalent of 5 embryos was loaded per well on a 4-12% Bis-Tris NuPAGE™ Novex™ gel and ran at 180V. Protein transfer was done for 1h10 at 110V to a nitrocellulose membrane, 0.2 μm (Invitrogen, LC2000). Membrane was blocked in 5% milk-PBT (PBS 1X 0.1% Tween 20) for 40 min and incubated overnight at 4°C with primary antibodies 1/2000 mouse anti-GAF or 1/2000 mouse anti-Tubulin in PBT. Anti-mouse and -rabbit IgG-HRP (Cell Signaling #7076 and #7074) secondary antibody were used at 1/4000 and incubated 1hour at room temperature. Chemiluminescent detection was done using Pierce™ ECL Plus (ThermoFisher) kit.

### DNA probe preparation and DNA-FISH

Probes were generated using 4 to 6 consecutive PCR fragments of 1.2 to 1.5 kb from *Drosophila* genomic DNA, covering approximately a 10 kb region. Primers are listed in Supplementary table 1. Probes were labeled using the FISH Tag DNA Kit (Invitrogen Life Technologies, F32951) with Alexa Fluor 488, 555, and 647 dyes following manufacturer’s protocol. Probes for satellite regions (related to Supplementary Fig. 1b) are from(Garavís et al. 2015).

DNA FISH was performed on 0-4 h AEL *yw* embryos adapted from (Bantignies and Cavalli 2014). Briefly, embryos were fixed as described above and were rehydratated with successive 3-5 min 1 ml washes on a rotating wheel with the following solutions: (1) 90% MeOH, 10% PBT; (2) 70% MeOH, 30% PBT; (3) 50% MeOH, 50% PBT; (4) 30% MeOH, 70% PBT; (5) 100 % PBT. Embryos were subsequently incubated in 200 µg RNase A (Thermo Scientific, EN0531) in 1 ml PBT for 2 h then 1 h at room temperature on a rotating wheel. Embryos were then slowly transferred to 100% pHM buffer (50% Formamide, 4x SSC, 100 mM NaH2PO4, pH 7.0, 0.1% Tween 20) in rotating wheel 20 min per solution 1 (1) 20% pHM, 80% PBS-Triton; (2) 50% pHM, 50% PBS-Triton; 80% pHM, 20% PBS-Triton; 100% pHM. Cellular DNA and probes were respectively denaturized in pHM and FHB (50% Formamide, 10% Dextran sulfate, 2x SSC, 0.05% Salmon Sperm DNA) for 15 min at 80 °C.

Probes and embryos were quickly pooled in the same PCR tube and slowly hybridized together with the temperature decreasing 1 °C every 10 min to reach 37 °C in a thermocycler. Washes were performed in pre-warmed solution (1 to 4) at 37 °C for 20 min under 900 rpm agitation (1) 50% Formamide, 10% CHAPS 3%, 10% SSC; (2) 40% Formamide, 10% CHAPS 3%, 10% SSC; (3) 30% Formamide; (4) 20% Formamide; then 20 min on a rotating wheel at room temperature using (5) 10% Formamide; (6) PBT; (7) PBS-Triton. Embryos were stained with DAPI at 0.5μg/ml, washed in PBT and mounted between slide and coverslip.

Images were acquired using a Zeiss LSM880 with confocal microscope in Airyscan mode with a Plan-Apochromat 63x/1.4 oil objective lens with the following settings: zoom 3.0x, z-planes 0.3 μm apart, 1024x1024 pixels.

### Distance measurements

To measure the distances between probes (*scyl-chrb* and *chrb-ctrl*, or *esg-sna* and *sna-ctrl*), we used a custom-made software developed in Python^TM^. This software is available through this link: https://github.com/ant-trullo/DNA_FishAnalyzer.

All the probes channels were treated with a 3D Laplacian of Gaussian filter (with kernel size 1) and then detected in 3D with manual thresholding on the filtered matrices; for each of the detected spots, the center of mass was determined. DAPI signal was treated with a 3D Gaussian filter (with user-defined kernel size) and the logarithm of the resulting matrix is thresholded with an Otsu algorithm, the threshold value being adjusted separately in each frame. The logarithm was used in order to compensate for non-homogeneous intensity inside nuclei. In order to generate distances, all the spots outside nuclei were removed. Then, nearest mutual neighbor spots were selected by calculating the distances of all the possible couples of spots and picking the smallest set. The distances were calculated with respect to the center of mass and using the Euclidean distance, taking into account the different pixel size on the z axis. A minimum of 10 images from 5 different embryos were analyzed for each condition. Aberrant distances (superior to 1μm) were not considered.

---

**Figure Sup1:**
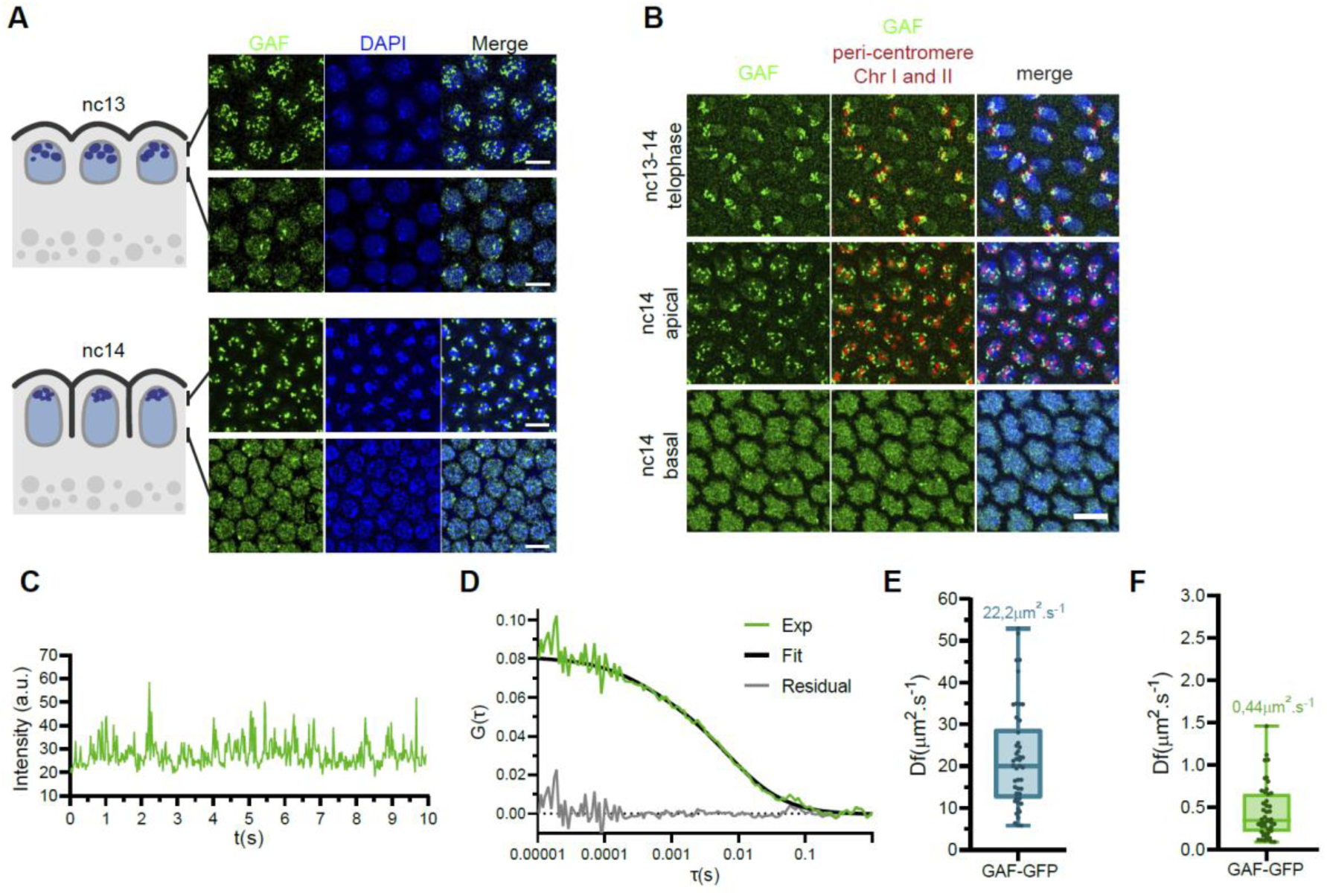
**GAF puncta localize to heterochromatin nuclear regions** (A) Maximum intensity projected Z-planes of confocal images showing GAF immunostaining (green) in wild type embryos at the indicated stages counterstained with DAPI (blue). Upper panels are images taken at the apical side of the nuclei, bottom panels at the basal side. Scale bars are 5µm. (B) Maximum intensity projected Z-planes of confocal images from DNA-immunoFISH with peri-centromeric probes (Garavís et al. 2015) (red) and anti-GAF (green) on wild type embryos and at the indicated nuclear cycles. Scale bar is 5µm. (C) Example of an intensity time trace obtained from FCS in a GAF-GFP nc14 embryo. (D) Example of autocorrelation function (green curve) related to (c) (black curve represents fitting using a double-diffusion model). (E) Estimated fast diffusion time for GAF-GFP extracted after fitting FCS data with a double-diffusion model performed in nucleoplasm. Centered horizontal line of the box plot represents the median. (F) Estimated slow diffusion time for GAF-GFP extracted after fitting FCS data with a double-diffusion model performed in nucleoplasm. Centered horizontal line of the box plot represents the median.

**Figure Sup2:**
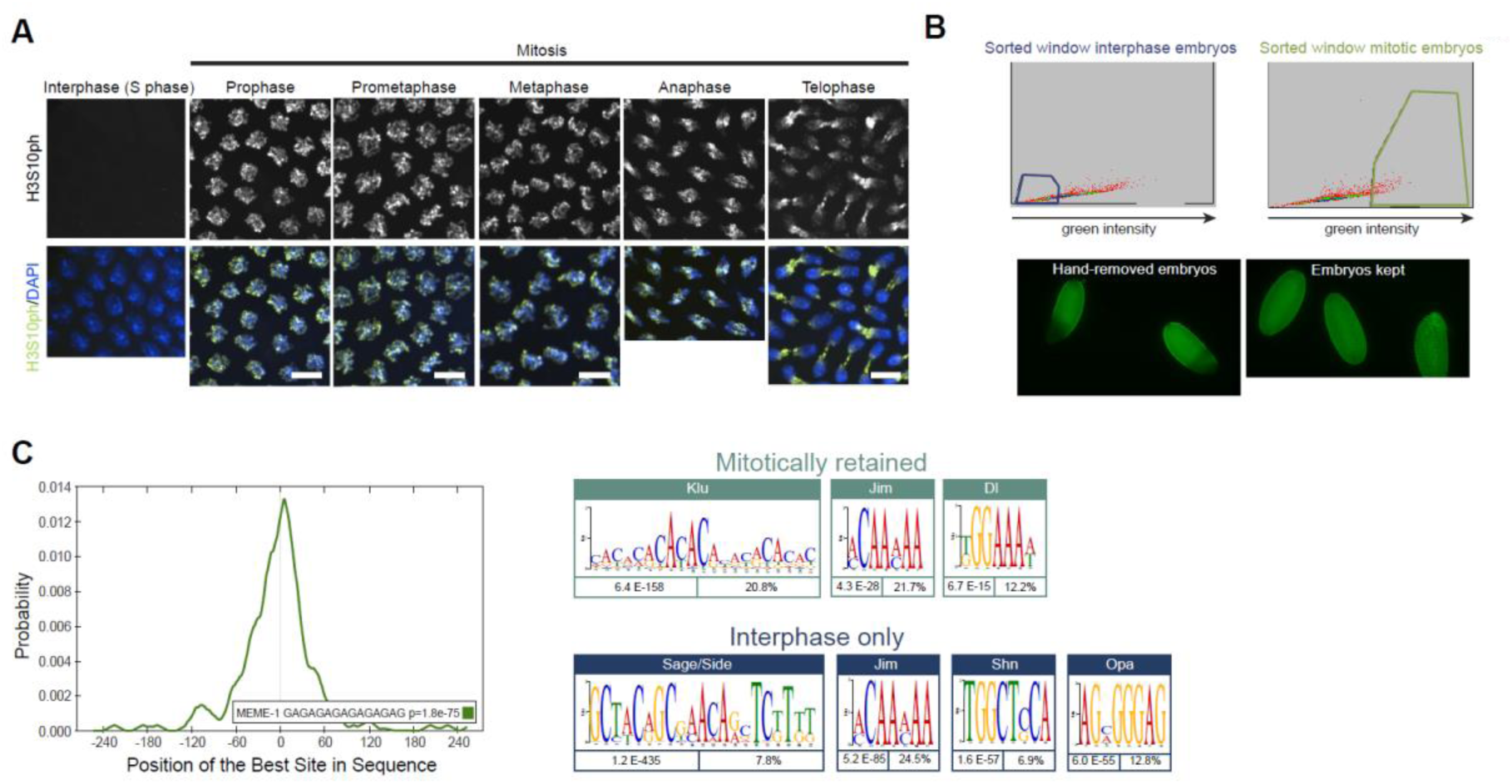
**Characterization of GAF mitotically retained regions** (A) Maximum intensity projected Z-planes of confocal images of H3S10ph immunostaining (grey and green) in wild type embryos at the indicated mitotic stages counterstained with DAPI (blue). Scale bars are 5µm. (B) (Top panels) Isolation of interphase and mitotic embryos previously stained with a H3S10ph antibody with a flow cytometer. Highlighted squares show the range for selected embryos for the interphase population (blue) and the mitotic population (green). (Bottom panels) Representative embryos stained with an H3S10ph antibody, undergoing mitotic waves (left image) (removed during the hand sorting) and fully mitotic embryos (right image). (C) Profile of probability for the GAGAG motif reported by MEME within sequences from GAF-ChIP-seq peaks. (D) De-novo motif enrichment within GAF mitotically retained and interphase only peaks, as reported by MEME.

**Figure Sup3:**
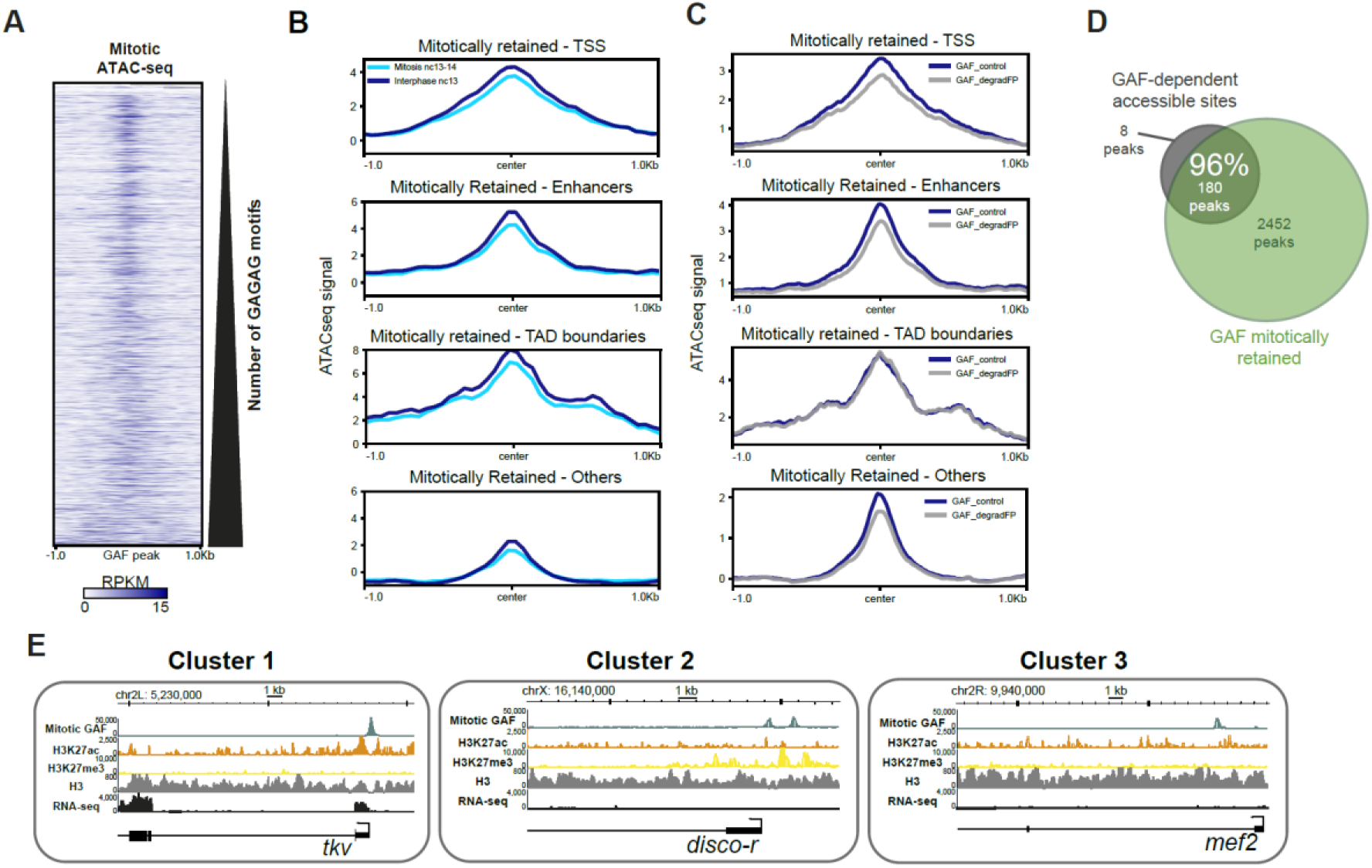
**Chromatin states of GAF mitotically retained regions** (A) Heatmap of ATAC-seq signal (Blythe and Wieschaus 2016) at GAF mitotically retained regions, partitioned by the number of GAGAG motifs they contain. (B) Metagene profiles of the ATAC-seq signal (Blythe and Wieschaus 2016) at GAF mitotically retained regions partitioned by the type of cis-regulatory element they overlap. Plots show peaks located at TSS, TAD boundaries (Hug et al. 2017), enhancers (Li et al. 2014; Kvon et al. 2014) and others that were unassigned. (C) Metagene profiles of the ATAC-seq signal in control (GAF control, blue) or GAF depleted (GAF_degradFP, grey) (Gaskill et al. 2021) embryos in different categories of GAF mitotically retained regions defined in Fig. 2b. (D) Venn diagram of the overlap between GAF dependent accessible loci (Gaskill et al. 2021) (grey) and GAF mitotically retained peaks (green). (E) Genome browser images showing ChIP-seq signals of mitotic GAF, H3K27ac, H3K27me3 and H3 (Li et al. 2014), alongside RNA-seq (Lott et al. 2011) at three loci containing GAF mitotically retained peaks. These three examples represent each of the three clusters defined by distinct chromatin states.

**Figure Sup4:**
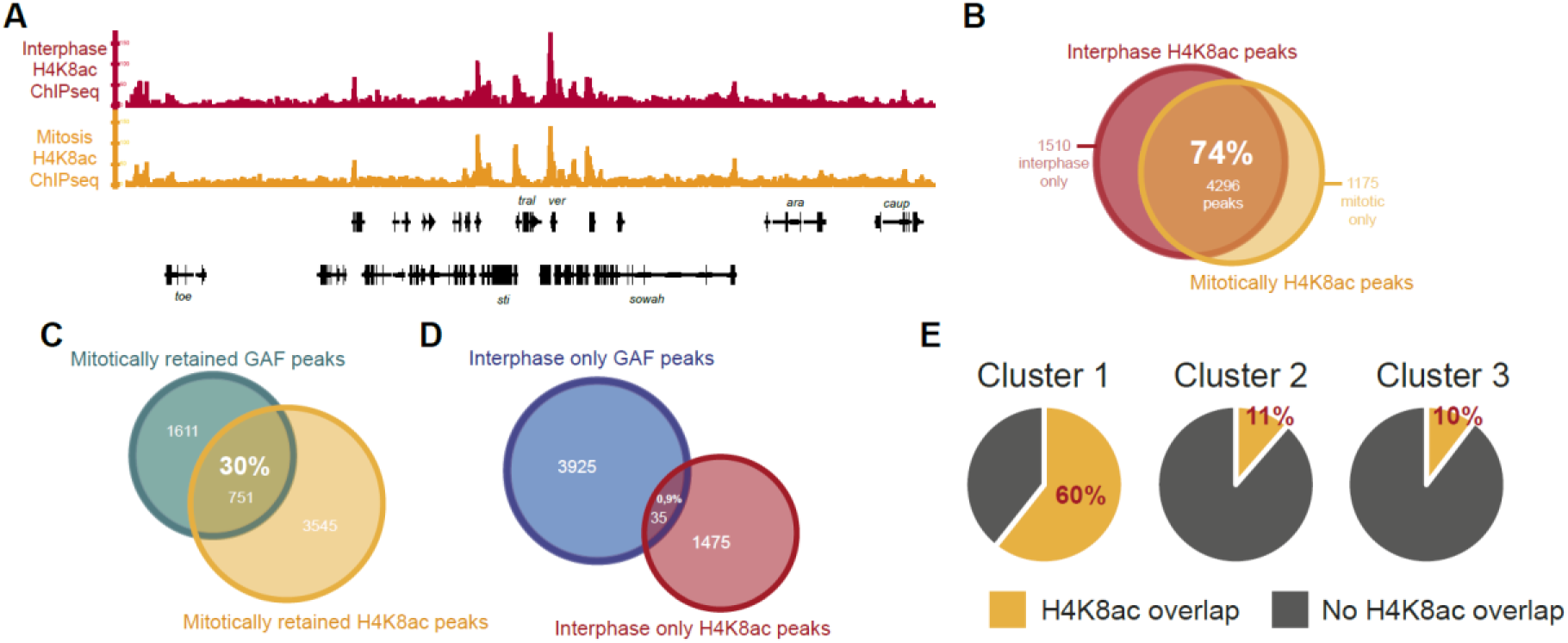
**H4K8ac-ChIP-seq in interphase and mitotic embryos** (A) Genome browser image showing H4K8ac-ChIP-seq profiles of interphase and mitotic embryos at a representative genomic region containing mitotically retained H4K8ac peaks. (B) Venn diagram of the overlap between H4K8ac interphase (red) and mitotic (orange) ChIP-seq peaks. (C-D) Venn diagram of the overlap between mitotically retained and interphase only (c) or H4K8ac and GAF-ChIP-seq peaks (D). (E) Pie charts of the overlap between H4K8ac-ChIP-seq mitotic peaks and GAF mitotically retained peaks, after their partitioning into three clusters.

**Figure Sup5:**
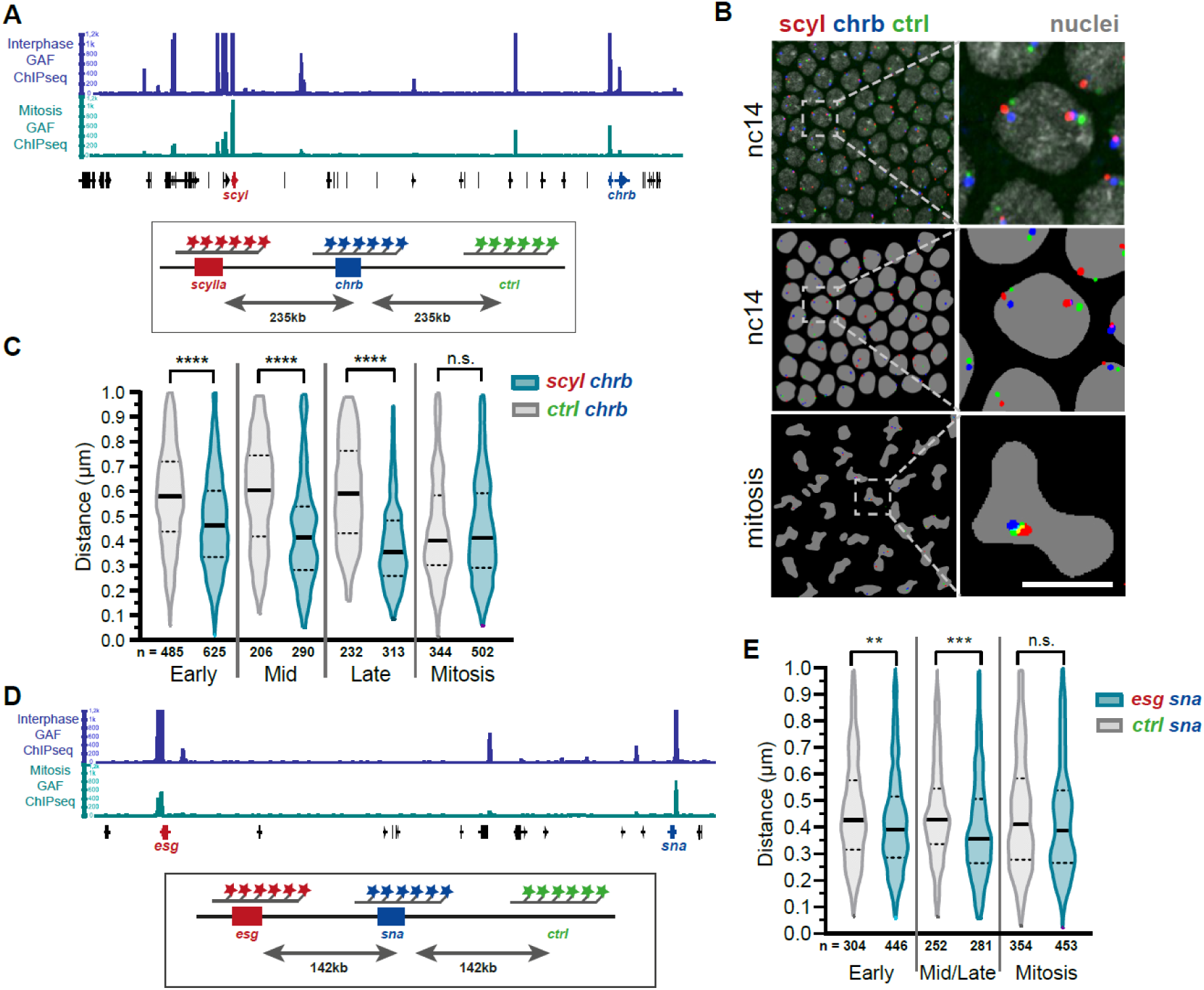
**Absence of mitotic loops for two GAF mitotically retained loci** (A) (Top) Genome browser image of the interphase and mitotic GAF- and H4K8ac-ChIP-seq profiles at the scyl-chrb locus. (Bottom) Schematic of the scyl-chrb locus with the indicated designed probes for DNA-FISH. The spatial distance between scyl and chrb was compared to that of a non-bookmarked region located at an equivalent distance (235kb). (B) DNA-FISH image in nc14 wild type embryo with scyl labeled in red, chrb in blue and control region in green. Bottom images represent the same image false colored after analysis as well as nuclei in mitosis. Scale bar is 5µm. (C) Violin plot representing the distance between scyl-chrb and chrb-ctrl from images taken in early, middle, late n.c.14 interphase or mitotic wild type embryos. Two tailed Welch’s t-test ****p<0.0001. (D) (Top) Genome browser image of the interphase and mitotic GAF-ChIP-seq profiles at the sna-esg locus. (Bottom) schematic of the sna-esg locus with the positions of designed probes for DNA-FISH indicated. (E) Violin plot representing the distance between sna-esg and sna-ctrl from images taken in nc14 interphase or mitotic wild type embryos. Two tailed Welch’s t-test ***p<0.001, **p<0.01.

**Figure Sup6:**
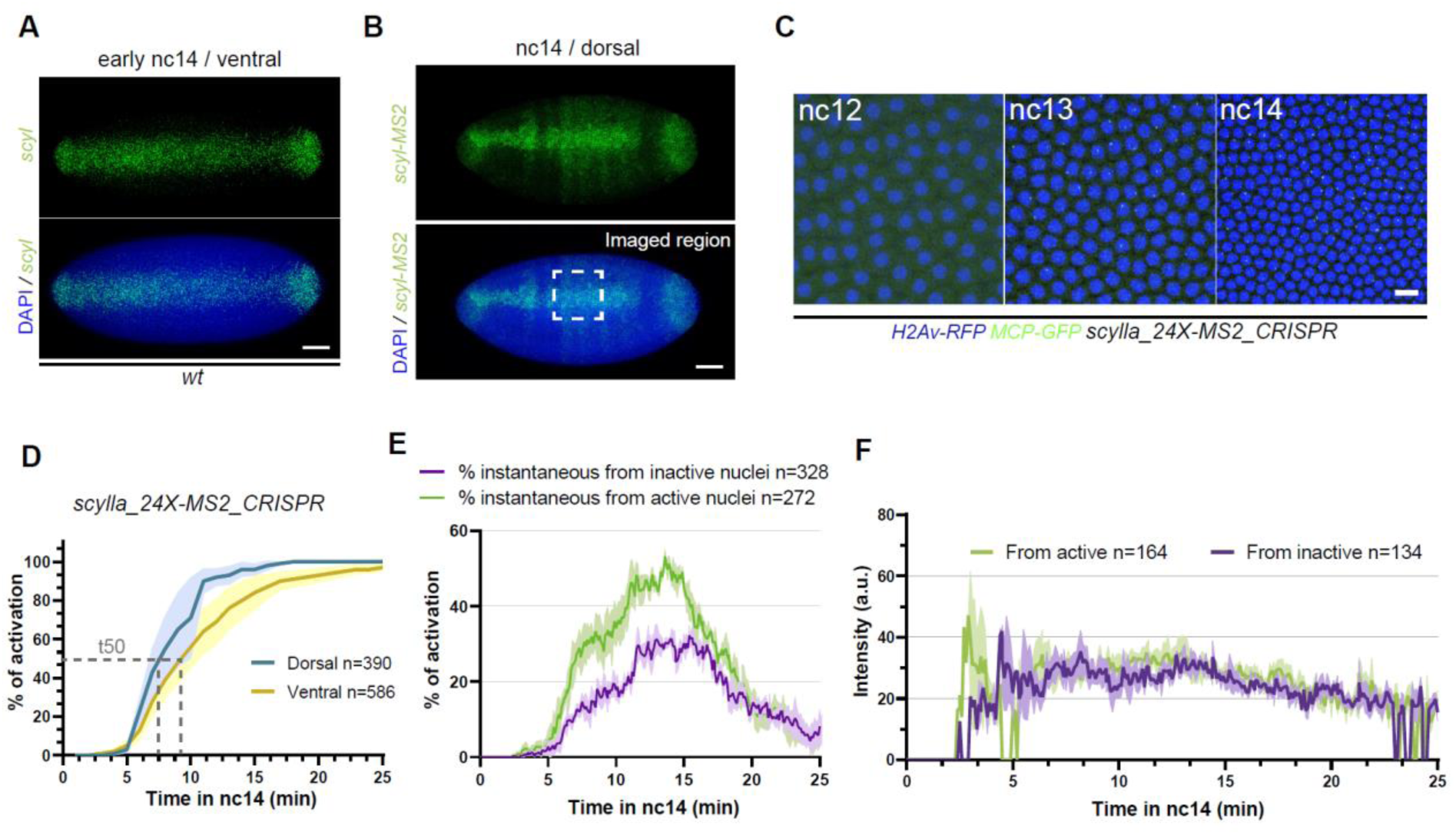
***scylla* transcription in the early embryo** (A) Maximum intensity projected Z-planes of confocal images from smiFISH with scyl probes (green) counterstained with DAPI (blue) on wild type embryos in early nc14 on the ventral side. Scale bars are 50µm. (B) Maximum intensity projected Z-planes of confocal images from smiFISH with scyl probes (green) counterstained with DAPI (blue) on wild type embryos in nc14 on the dorsal side. Dashed square represents the regions imaged for the quantifications shown in (D). Scale bars are 50µm. (C) Snapshots from a representative movie of scylla_24X-MS2_CRISPR/+ embryo carrying MCP-eGFP, H2Av-RFP. Nuclei are visualized in blue and transcription sites in green. Scale bar corresponds to 10µm. (D) Quantification of the transcriptional synchrony after mitosis in scylla_24X-MS2_CRISPR embryos in dorsal (blue) and ventral (yellow) regions. Both daughters of each nucleus are quantified. SEM are represented in light blue and light yellow respectively. n=number of nuclei analyzed from 3 movies of 3 embryos for each condition. (E) Instantaneous percentage of activation after mitosis of nuclei coming from active nuclei (green) and from inactive (purple). SEM are represented in light purple and light green respectively. n=number of nuclei analyzed from 3 movies of 3 embryos. (F) Mean intensity of scyl transcriptional site of nuclei coming from active nuclei (green) and from inactive nuclei (purple). SEM are represented in light green and light purple. Both of the two daughters of each nucleus are quantified. n=number of nuclei analyzed from 4 movies of 4 embryos.

**Figure Sup7:**
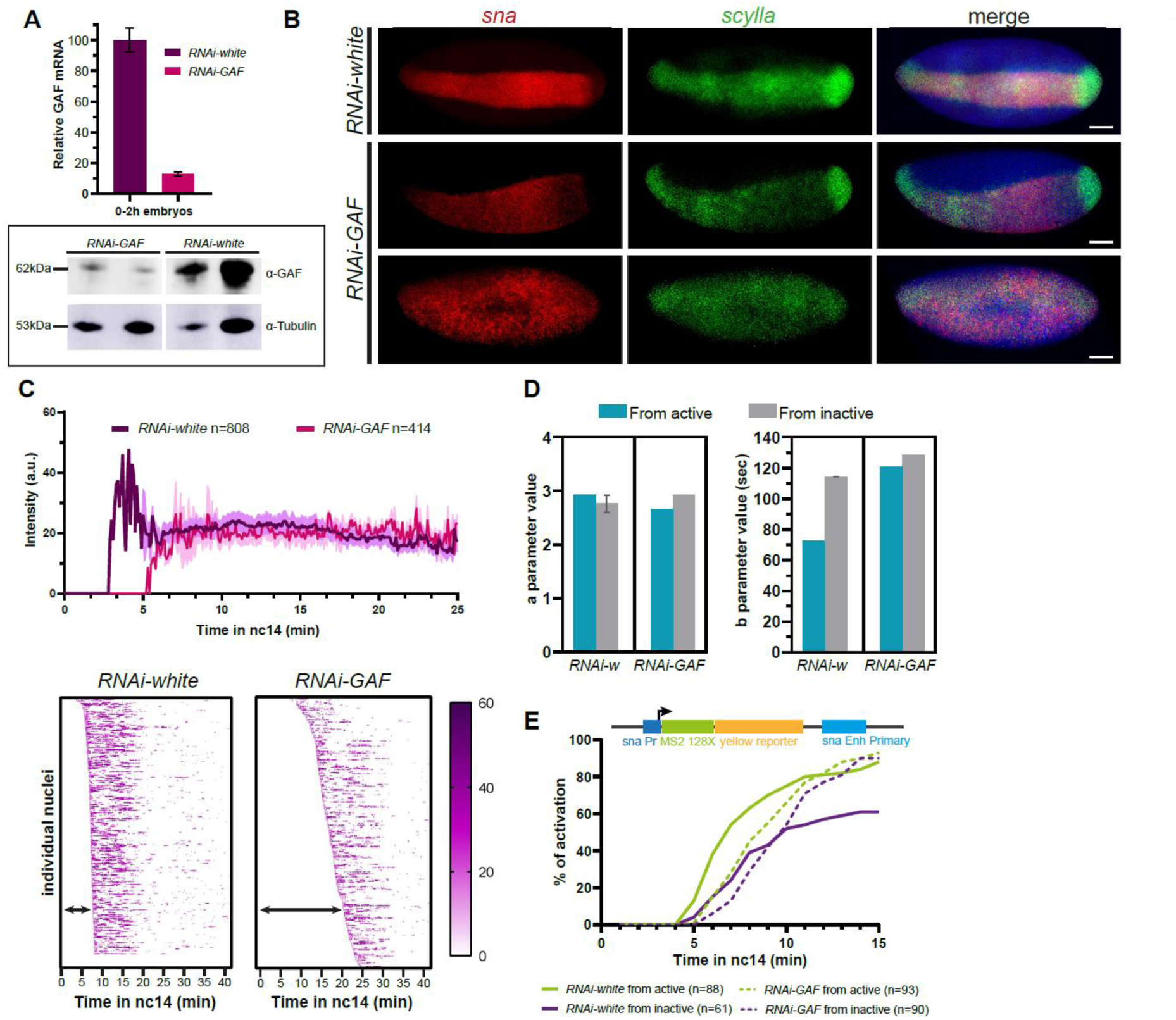
**Effect of GAF reduction on *scylla* transcription** (A) (Top) Histogram of the relative amount of RPL32 transcripts normalized GAF mRNA in RNAi-white and RNAi-GAF 0-2h embryos quantified by RT-qPCR. (Bottom) Two examples of western blot analysis of RNAi-white and RNAi-GAF 0-2h embryos, with the indicated antibodies. Each experiment was performed in biological triplicates. (B) Maximum intensity projected Z-planes from confocal images from a smFISH with scyl probes (red) and sna probes (green) counterstained with DAPI (blue) on RNAi-white and RNAi- GAF embryos, showing different types of phenotypes. Scale bars are 50µm. (C) (Top) Mean intensity of scyl transcriptional site (scylla_24X-MS2_CRISPR) of nuclei in RNAi- white embryos (control, purple) and RNAi-GAF embryos (pink). SD are represented in light purple and light pink. Both of the two daughters of each nucleus are quantified. n=number of nuclei analyzed from 4 movies of 4 embryos. (Bottom) Heatmaps of scyl (scylla_24X- MS2_CRISPR) transcriptional site intensity of individual nuclei in RNAi-white and RNAi-GAF embryos sorted by their first activation time. (D) Histograms of ‘a’ and ‘b’ parameters extracted from mathematical modeling for the scyl (scylla_24X-MS2_CRISPR) gene in RNAi-white and RNAi-GAF scylla_24X-MS2_CRISPR embryos. (E) (Top) Schematic representing the snail-primary-enhancer_MS2 transgene. (Bottom) Cumulative activation of the first activated nuclei coming from active nuclei (green) and from inactive (purple) in RNAi-white embryos (control, solid curves) and RNAi-GAF embryos (dashed curves). n=number of nuclei analyzed from 4 movies of 4 embryos.

## Movie Legends

**Movie1: Imaging GAF behavior during the cell cycle**

Maximum intensity projection of confocal live imaging of a developing *His2Av-mRFP;GAF-GFP* embryo. Scale bar is 5μm.

**Movie2: GAF subnuclear localization**

Maximum intensity projection of confocal live imaging of *His2Av-mRFP;GAF-GFP* embryo. Top movie comprises a six Z-planes projected images at the apical side of nuclei, bottom movie is a six Z-planes projected images at the basal side of nuclei. Time is in minutes. Scale bar is 5μm.

**Movie3: Transcription of *scylla* in a control embryo**

Maximum intensity projection of confocal live imaging of a *mat-alpha-Gal4/+; nos-Gal4, MCP-eGFP, H2Av-RFP/UASp-shRNA-white > scylla_24X-MS2_CRISPR/+* embryo.

Nuclei are visualized in red and transcriptional sites in green. Scale bar is 10μm.

**Movie4: Transcription of *scylla* in a GAF maternally depleted embryo**

Maximum intensity projection of confocal live imaging of a *mat-alpha-Gal4/+; nos-Gal4, MCP-eGFP, H2Av-RFP/UASp-shRNA-GAF > scylla_24X-MS2_CRISPR/+* embryo.

Nuclei are visualized in red and transcriptional sites in green. Scale bar is 10μm.

## Supplementary Tables Legends

**Supplementary table 1:** Primers sequences for cloning, DNA-FISH and smFISH probes sequences used in this study.

**Supplementary table 2:** Identified GAF mitotically retained, interphase only and mitotic only peak coordinates with the nearest gene identified and its distance in base pair from the TSS.

**Supplementary table 3:** Identified GAF mitotically retained, interphase only and mitotic only peak coordinates with their respective features.

**Supplementary table 4:** Whole-genome data used in this study.

