## Supplementary Table 1 for "The control of transcriptional memory by stable mitotic bookmarking"

**Primers for cloning**

| | |
|---|---|
| SnaPr_F | acgcggccgcGACAGCGGCGTCGGCAG |
| SnaPr_R | cgggatccTGGTTGCGTTCTCAACG |
| SnaPrimaryEnh_F | tttctagaGCTCCTCTTCGAACAATGTCAG |
| SnaPrimaryEnh_R | cctctagaGCCATTTGGTGGTGGTTCTTCTTC |
| 3'Homology_scyl_F | aaagatctccatgcataagggcgaaacaaaaaaaacacgacc |
| 3'Homology_scyl_R | cgagctgcagaaggcctagggcagcaagctgcgtctaac |
| 5'Homology_scyl_F | gcaggtggaattcttgcatggtattctacctcgttttattacc |
| 5'Homology_scyl_R | atccatcacactggcggccggcagacacatatacgaac |
| Guide_scyl_F | gtcgGCTCTCGATCATCACAGGGG |
| Guide_scyl_R | aaacCCCCTGTGATGATCGAGAGC |

**FLAPY Probes**

| | |
|---|---|
| ScylCDSandUYR_FLAPY1 | GGC TTC GAA TAC TTG CTT CGT TGT TGC ACT TAC ACT CGG ACC TCG TCG ACA TGC ATT |
| ScylCDSandUYR_FLAPY2 | GAT GGT GAT CGT GCG GGC CAA ACT CTT TTA CAC TCG GAC CTC GTC GAC ATG CAT T |
| ScylCDSandUYR_FLAPY3 | GCG ATG ATC CTG CCT GAG GGT CAG ATT TAC ACT CGG ACC TCG TCG ACA TGC ATT |
| ScylCDSandUYR_FLAPY4 | ACT TCG AAT GTG GAC ACC GTA TCC GAA TCG TTA CAC TCG GAC CTC GTC GAC ATG CAT T |
| ScylCDSandUYR_FLAPY5 | CTT GAT CGA GGC AAT GCG ACG CGA ATT CTT TAC ACT CGG ACC TCG TCG ACA TGC ATT |
| ScylCDSandUYR_FLAPY6 | GGC TCA TCC TCG AAC TCA ATG TAG ATG GTT TAC ACT CGG ACC TCG TCG ACA TGC ATT |
| ScylCDSandUYR_FLAPY7 | AGT TGC TCA GTG CAT AGT CAG CGC TAC TTA CAC TCG GAC CTC GTC GAC ATG CAT T |
| ScylCDSandUYR_FLAPY8 | GGC TTT GTC TTC TTG CTC TGA CTT TGG CTT ACA CTC GGA CCT CGT CGA CAT GCA TT |
| ScylCDSandUYR_FLAPY9 | CAT CAG GTC CAG AGT GTA TGC CGA CGG TTA CAC TCG GAC CTC GTC GAC ATG CAT T |
| ScylCDSandUYR_FLAPY10 | TTT GAT CCC TGC AGG CTA CCG TAG ATG TTA CAC TCG GAC CTC GTC GAC ATG CAT T |
| ScylCDSandUYR_FLAPY11 | ATT TTA TCA ACG TCT TTT GCC GGA CTT TGG TGT TAC ACT CGG ACC TCG TCG ACA TGC ATT |
| ScylCDSandUYR_FLAPY12 | TGA TTC TGG GGA TTG ATT CGT GGG GCT GTT TAC ACT CGG ACC TCG TCG ACA TGC ATT |
| ScylCDSandUYR_FLAPY13 | AGA GTG TGG TTG GCG TAA CCT AAT GTG TAT GTT TAC ACT CGG ACC TCG TCG ACA TGC ATT |
| ScylCDSandUYR_FLAPY14 | ACA GGG GTG GTG TTT TTT TTG TTT CGC GCA GAT TAC ACT CGG ACC TCG TCG ACA TGC ATT |
| ScylCDSandUYR_FLAPY15 | CTG ATC ACC ATT AGT TGG TGG GCG TGG CTT ACA CTC GGA CCT CGT CGA CAT GCA TT |
| ScylCDSandUYR_FLAPY16 | TAG CTT ATT CTT GGT GAT CGT ATA CTC CGG ATT ACA CTC GGA CCT CGT CGA CAT GCA TT |

| | |
|---|---|
| ScylCDSandUYR_FLAPY17 | GCT CCT TCT CCG ACA CGC GGA TTA TTT TAC ACT CGG ACC TCG TCG ACA TGC ATT |
| ScylCDSandUYR_FLAPY18 | CGA CAC TTC CGT ACA GGT CAA ATG ACG CGT TAC ACT CGG ACC TCG TCG ACA TGC ATT |
| ScylCDSandUYR_FLAPY19 | GTA GTT GCT ATT GCT ATT ATA GCT GCC GCT TAC ACT CGG ACC TCG TCG ACA TGC ATT |
| ScylCDSandUYR_FLAPY20 | GTG GCA GTC TGC TGG TCC AAT CTT TAT TTT TTA CAC TCG GAC CTC GTC GAC ATG CAT T |
| ScylCDSandUYR_FLAPY21 | AGT CGA ATC GAA TCG AAA GTC GTT CGT TCG CTT TAC ACT CGG ACC TCG TCG ACA TGC ATT |
| ScylCDSandUYR_FLAPY22 | CTC GCT CGC AAT TTT CCG ACG GCT TTT GTT ACA CTC GGA CCT CGT CGA CAT GCA TT |
| ScylCDSandUYR_FLAPY23 | TGT TGT ATC ACT TAA ATG GCG TTG GTC CGT TGT TAC ACT CGG ACC TCG TCG ACA TGC ATT |
| ScylCDSandUYR_FLAPY24 | CTG CGA GAG TCT GTT TTC TTG TGT ACT GTG TTT TAC ACT CGG ACC TCG TCG ACA TGC ATT |
| ScylCDSandUYR_FLAPY25 | GAC AGC TCC TCG ACA GCC TTC TTA TCT TAC ACT CGG ACC TCG TCG ACA TGC ATT |
| ScylCDSandUYR_FLAPY26 | CCG CCG GCT GCT TGT GCT TAA TAC TGT TAC ACT CGG ACC TCG TCG ACA TGC ATT |

**Primers for qRT-PCR**

| | |
|---|---|
| RpL32_F | CTTCATCCGCCACCAGTC |
| RpL32_R | CGACGCACTCTGTTGTCG |
| GAF_F | ACAAGATAGTCCTGTGCGCC |
| GAF_R | GCCAACATAACCACTGGATGC |

**Primers for DNA-FISH probes**

**Probe 1 (scyl)**

| | |
|---|---|
| SCYL_primer1_F | CTCGCACGTTGATTCGGTCAC |
| SCYL_primer1_R | CCAATAGATAGCTGGTGACGCC |
| SCYL_primer2_F | GATCATTAACCCATGAAAGTCGTCTG |
| SCYL_primer2_R | cgagctggaataggtgtgaaatatc |
| SCYL_primer3_F | cttcgagcgttacaaacaaacatg |
| SCYL_primer3_R | CCCGAGCAATAACATCCATTTTC |
| SCYL_primer4_F | catgtgtttcccaaaatgcagtc |
| SCYL_primer4_R | CCATCAGCGGAATATAGCTTATTC |
| SCYL_primer5_F | GGTGATCAGCTTGATCTGTCAG |
| SCYL_primer5_R | CGTGTGTGTTTGTATGTTGGTG |
| SCYL_primer6_F | gtcgaacatctagcatccacctcaag |
| SCYL_primer6_R | ccacctacgaccgaagttgatgtg |

**Probe 2 (chrb)**

| | |
|---|---|
| CHRB_primer1_F | gtacacgatgcaaacaaatcacc |
| CHRB_primer1_R | gtcacatttggctgtcaatttgtg |
| CHRB_primer2_F | ctctttctgccgcagcagag |
| CHRB_primer2_R | ggacaaatgtgatgcggaatggg |
| CHRB_primer3_F | ggatatgccatgtgggttctcgc |
| CHRB_primer3_R | gtaccgcctccagttctgttggt |
| CHRB_primer4_F | gccaaattacagaaaaccgcctcc |
| CHRB_primer4_R | GCTATAGGCGTTTGATCTCGACCG |
| CHRB_primer5_F | GAAGATGGAAGTGCTCTCAGTAC |
| CHRB_primer5_R | cggtatatctgcgacagtatgtatg |
| CHRB_primer6_F | gtgtagttaaaggggtattccatctg |
| CHRB_primer6_R | gcacttcggtatggcaatcatg |
| CHRB_primer7_F | cgccagctttgtgcgtctcattc |
| CHRB_primer7_R | cgacagcgtcttgtaattagcgcga |

**Probe 3 (ctrl)**

| | |
|---|---|
| Ctrl_scyl_chrb_1F | ccatttccaccggcaagccttg |
| Ctrl_scyl_chrb_1R | gtcagccgatcggcgtgaag |
| Ctrl_scyl_chrb_2F | ggagctcaaataacttgatggattg |
| Ctrl_scyl_chrb_2R | gaatgtgggcaaatataattagcgg |
| Ctrl_scyl_chrb_3F | ggcagaaatcacttagcgcag |
| Ctrl_scyl_chrb_3R | GTTGATGTCCGGAATCGAAACG |
| Ctrl_scyl_chrb_4F | TCATTGCAGGCACCTTTCCTATCG |
| Ctrl_scyl_chrb_4R | CCTCACCCAGGTGGGAATGAAACC |

**Probe 1 (esg)**

| | |
|---|---|
| ESG_primer1_F | CATGGTGGTTAGGAGAGGAG |
| ESG_primer1_R | CAAAAGTTGGCACTGCAAGC |
| ESG_primer2_F | TGGCTTTGGCGCCTCAAAC |
| ESG_primer2_R | GAGTCTGGGAATAACTGAGCGAG |
| ESG_primer3_F | TGTGCCTGTGTGAACATGAC |
| ESG_primer3_R | AAGGGCTTCTCTCCCGTATG |

| | |
|---|---|
| ESG_primer4_F | CCTCCTAACCAAGCATTCAGAG |
| ESG_primer4_R | TTGGGTCGGCTCTTATTTTCCG |
| ESG_primer5_F | ATTTCGGTGCACGTACGAC |
| ESG_primer5_R | AATGCAGCGATCAAGCGC |
| ESG_primer6_F | TTGACTGATTTTGGGGACCCTG |
| ESG_primer6_R | GACAGCAAGTCCCATTTGTGC |
| **Probe 2 (sna)** | |
| Snail_primer1_F | GAACGGATTGCATCCTAGTCAG |
| Snail_primer1_R | CGATTCCAGTTGATGGGATCCAATCC |
| Snail_primer2_F | ATACCTGGATCGTGGCTCTTGG |
| Snail_primer2_R | TTAATCGCAGCGCTGATGTGG |
| Snail_primer3_F | TAGTTGACTTGGCTGGCG |
| Snail_primer3_R | AGCACAGATGCAACTGGTC |
| Snail_primer4_F | CCAGCGGAATGTGAGTTTGC |
| Snail_primer4_R | CATTGTCTTCGTGGAGGAGC |
| Snail_primer5_F | TTGCGTTCTCAACGAGAGCTG |
| Snail_primer5_R | GACGCGGAAAATCGGACTTG |
| Snail_primer6_F | ATTCCAATTCCCCGCGATC |
| Snail_primer6_R | GATCGCAGCTCCTTTGTTC |
| Snail_primer7_F | TAGGTTCGCGGCACAATG |
| Snail_primer7_R | GCCACCCTCGAGTTGACATTATTTCC |
| **Probe 3 (ctrl)** | |
| Ctrl_esg_sna_1F | CAATAAATCCCCTAAACGGTGGCC |
| Ctrl_esg_sna_1R | CGCATTCATGCCACTCGATCTTC |
| Ctrl_esg_sna_2F | CCACGGTCGGTGGAGTAATTTAATCC |
| Ctrl_esg_sna_2R | CGTGTGCACTCATTTGCTGATG |
| Ctrl_esg_sna_3F | GAGCATAATAAGCTCAACGCCG |
| Ctrl_esg_sna_3R | CTACACATGACTCCCCGTAAAG |
| Ctrl_esg_sna_4F | GTGCACTAGCCGCACATTAAACC |
| Ctrl_esg_sna_4R | ACCGACTGACGCGATTAGATC |
| Ctrl_esg_sna_5F | TGTCCTTGGAGCTAAGACCCAC |
| Ctrl_esg_sna_5R | GCCAAGCAAACTCTTGTGTTCAGC |

| | |
|---|---|
| Ctrl_esg_sna_6F | CTGGAAGCTTTGAAAACTCCTGG |
| Ctrl_esg_sna_6R | GATCTTATCTCATTGGCCCTGGG |
| Ctrl_esg_sna_7F | TCTGAGCTCAACACAGCACAC |
| Ctrl_esg_sna_7R | CTGCTTACCTTTCCTACCGTCCTAC |
