## Supplementary Table 2 for "The control of transcriptional memory by stable mitotic bookmarking"

| **Coordinates_Mitotically_Retained** | **nearst_gene** | **distance_nearest_gene** |
|---|---|---|
| chrY:1050951:1051901: | ARY | 348522 |
| chrY:124051:124901: | kl-5 | 12585 |
| chrY:132251:132801: | kl-5 | 4685 |
| chrY:1416751:1417151: | Ppr-Y | 219423 |
| chrY:1550501:1551001: | Ppr-Y | 85573 |
| chrY:200551:201051: | PRY | 8532 |
| chrY:203501:203901: | PRY | 11482 |
| chrY:3652701:3653101: | CCY | 2706 |
| chrY:3667101:3667501: | CCY | 17106 |
| chrY:853601:854851: | ARY | 151172 |
| chrY:860151:860651: | ARY | 157722 |
| chrY:877851:878351: | ARY | 175422 |
| chrY:883751:884301: | ARY | 181322 |
| chrY:885801:886651: | ARY | 183372 |
| chrY:887401:887901: | ARY | 184972 |
| chrY:917651:918151: | ARY | 215222 |
| chr2R:11450551:11451051: | E | 992 |
| chr3L:4406351:4407301: | DOR | 0 |
| chrX:17826901:17827401: | Socs16D | 163 |
| chr2R:22688701:22690101: | Ppa | 0 |
| chrX:7891351:7892501: | Nek2 | 0 |
| chr2R:21474801:21476951: | Sdc | 4225 |
| chr3L:2589801:2590301: | dos | 388 |
| chr2R:12586751:12590351: | Sin3A | 2566 |
| chr3L:15141651:15142351: | Pdi | 489 |
| chrX:2009951:2010851: | east | 9 |
| chr3L:15507301:15508401: | CG7841 | 0 |
| chr3R:24104651:24106151: | Syx1A | 0 |
| chr2L:19574851:19577201: | spi | 114 |
| chr2R:13041551:13042501: | CG13323 | 4755 |
| chr2R:10806351:10807751: | wde | 1187 |
| chr2L:5977351:5981001: | chic | 17 |
| chr2L:4281201:4283901: | tutl | 0 |
| chr3R:14284851:14287151: | trx | 0 |
| chr3R:14282551:14284601: | trx | 2302 |
| chr2L:16325551:16326251: | cact | 0 |
| chr3R:5216101:5216701: | cno | 755 |
| chrX:7324251:7325051: | Dok | 38 |
| chr3R:20948901:20949701: | Atpalpha | 188 |
| chrX:18309051:18310101: | upd1 | 0 |
| chr3L:14409451:14410951: | CG3919 | 0 |
| chr2R:7381501:7382401: | Dscam1 | 0 |
| chr2R:13957351:13958451: | AGO1 | 0 |

| | | |
|---|---|---|
| chrX:8054101:8055501: | fs | 5935 |
| chr2L:9705201:9706401: | CG31710 | 0 |
| chr3L:13924751:13925851: | Rgl | 0 |
| chr2R:25084251:25084751: | gol | 0 |
| chr3L:9614401:9615101: | LanB2 | 0 |
| chr2R:13822551:13823101: | fl | 1452 |
| chr3L:12265651:12266201: | app | 640 |
| chr2R:18799101:18800901: | MFS15 | 2424 |
| chrX:5083451:5084751: | CG32767 | 17 |
| chr2R:13824001:13824701: | fl | 0 |
| chr3L:20517201:20517951: | CG5059 | 1442 |
| chr2R:18802051:18803251: | MFS14 | 476 |
| chrX:8056201:8059351: | fs | 2085 |
| chr3R:4174251:4175551: | CG12581 | 0 |
| chrX:18312301:18313151: | upd1 | 3011 |
| chr3R:10267051:10267601: | CG43143 | 2746 |
| chrX:12202451:12204101: | CG1924 | 23826 |
| chr3R:7084551:7087351: | alphaTub84B | 0 |
| chr2L:18444551:18445101: | RpS26 | 1275 |
| chr3R:10048601:10049101: | CG12811 | 0 |
| chrX:3134601:3135901: | N | 0 |
| chr2R:11892601:11893301: | eEF1alpha1 | 1467 |
| chr3R:4478001:4479301: | CG31523 | 0 |
| chr3L:13922751:13923301: | Rgl | 2034 |
| chrX:616751:617451: | CG13366 | 2744 |
| chrX:13284201:13285101: | HDAC4 | 531 |
| chr2R:9957901:9958951: | Mef2 | 0 |
| chr3R:30563051:30564051: | CG34155 | 2853 |
| chrX:19264201:19264801: | Vav | 2382 |
| chr2R:12591301:12592801: | Sin3A | 116 |
| chr2R:10422951:10424651: | Prx2540-1 | 0 |
| chr3L:2585601:2586651: | msn | 0 |
| chrX:2343351:2343851: | mRpL14 | 798 |
| chrX:18789851:18790351: | Atg101 | 1317 |
| chr3L:4409801:4410301: | DOR | 2859 |
| chr2R:17547901:17548751: | MESR4 | 133 |
| chr3L:5657501:5658051: | sif | 0 |
| chrX:2011951:2012651: | east | 1092 |
| chrX:19321751:19323951: | out | 0 |
| chr3L:4252051:4252601: | ago | 1705 |
| chr2L:9613151:9613801: | Trx-2 | 0 |
| chr3R:6816201:6816851: | Dfd | 24366 |
| chr3R:11956301:11957001: | Hsp70Aa | 0 |
| chrX:7320601:7321351: | Atg5 | 2419 |
| chr3L:14624751:14625301: | CG9628 | 1956 |

| | | |
|---|---|---|
| chr2L:19570951:19573251: | spi | 4064 |
| chr2R:9283951:9285151: | Pkn | 0 |
| chr3L:4252951:4253551: | ago | 755 |
| chr3L:13919701:13921001: | Rgl | 4334 |
| chrX:12760201:12760751: | ade5 | 0 |
| chrX:10729301:10730551: | ras | 13951 |
| chr3R:8119701:8120351: | CG7878 | 2374 |
| chr2R:10420901:10422351: | CG33474 | 0 |
| chr2R:9954751:9956051: | Mef2 | 2807 |
| chr3L:7134001:7135301: | melt | 0 |
| chr2R:10534051:10536651: | lola | 6640 |
| chr2R:16856151:16857551: | Dek | 0 |
| chr3R:11958151:11958851: | Hsp70Ab | 0 |
| chr3R:9502301:9503001: | RhoL | 0 |
| chr2L:594401:595551: | Gsc | 0 |
| chr2L:7048901:7049551: | Mnn1 | 6750 |
| chr3R:7088601:7089351: | alphaTub84B | 2003 |
| chr3L:1794051:1795351: | dlt | 0 |
| chrX:12574651:12575301: | hwt | 6200 |
| chr3R:25880251:25881101: | Tsp96F | 326 |
| chr3R:16449101:16450051: | Pak3 | 3770 |
| chrX:10731151:10732401: | ras | 12101 |
| chr2L:15491801:15492551: | lace | 6576 |
| chr2R:21047601:21048501: | shg | 8696 |
| chr3R:23669301:23670501: | CG31145 | 0 |
| chr3R:11340051:11340501: | pros | 11572 |
| chrX:19259751:19260501: | rictor | 739 |
| chr3L:15825401:15826451: | DCP2 | 0 |
| chr3L:11821651:11822401: | CG11597 | 0 |
| chrX:8409651:8412151: | lawc | 0 |
| chr2L:3803101:3804251: | Reph | 0 |
| chr3R:18169851:18170451: | wrd | 2095 |
| chrX:19631201:19631901: | CG14212 | 5497 |
| chrX:11350501:11351401: | CG1737 | 0 |
| chr3R:28330451:28331251: | larp | 5189 |
| chr3L:16483401:16484501: | aos | 0 |
| chr3L:11819551:11820401: | CG11597 | 1874 |
| chr2R:8241801:8242351: | pnut | 4531 |
| chr2L:4371351:4372301: | Traf4 | 9427 |
| chrX:16441901:16442201: | rngo | 469 |
| chr3R:18171851:18173151: | wrd | 4095 |
| chr3L:19909951:19910451: | Prp3 | 0 |
| chrX:5754951:5755901: | CG15771 | 0 |
| chr2R:12025051:12025551: | otk2 | 0 |
| chr3L:17421251:17421901: | blot | 0 |

| | | |
|---|---|---|
| chrX:6861351:6862051: | CG43736 | 1837 |
| chr3L:14775651:14776051: | mop | 78 |
| chr2L:21899001:21900001: | CG11629 | 713 |
| chrX:6279051:6279901: | Ubi-p5E | 0 |
| chrX:13396551:13397101: | CG1673 | 0 |
| chr2R:9552601:9553501: | CG1888 | 4467 |
| chrX:6668801:6669551: | CG3198 | 4422 |
| chr2R:24525551:24526201: | Brca2 | 0 |
| chr2R:13223501:13224101: | Fsn | 1043 |
| chr3L:12515601:12516051: | tral | 543 |
| chr3L:16023001:16023701: | Notum | 0 |
| chr3L:1334801:1336601: | CG2211 | 0 |
| chrX:2136651:2137351: | ph-p | 1020 |
| chr3L:18839901:18840951: | Indy | 5316 |
| chrX:2135351:2135851: | ph-p | 2520 |
| chr3L:15336151:15337001: | Toll-6 | 0 |
| chr3L:18834551:18835051: | Indy | 11216 |
| chrX:3686901:3689651: | Mnt | 15101 |
| chr2R:15675101:15676951: | fus | 0 |
| chr2R:11603151:11603951: | tou | 12094 |
| chr2R:11604451:11605601: | tou | 10444 |
| chr3R:10886501:10887401: | Leash | 962 |
| chr3R:10889001:10889751: | Adk3 | 0 |
| chr2L:19164651:19166251: | CG17568 | 15451 |
| chrX:13089651:13090451: | hep | 0 |
| chrX:13739351:13740001: | CG11151 | 0 |
| chrX:13087501:13087951: | hep | 2483 |
| chr2L:107701:108351: | Sam-S | 799 |
| chr3R:10355951:10356951: | jumu | 5770 |
| chr2L:7976701:7977151: | CG7231 | 6795 |
| chr2L:8949651:8950701: | C1GalTA | 515 |
| chr2L:8951001:8951901: | C1GalTA | 0 |
| chr2L:18844801:18845451: | CG17321 | 0 |
| chr3R:13644551:13645601: | sqd | 703 |
| chr3R:12719601:12720301: | Men | 2141 |
| chr3L:11214401:11216151: | CG42671 | 0 |
| chrX:5516751:5518701: | CG12730 | 144 |
| chr3R:9229001:9230751: | pum | 6931 |
| chrX:6981401:6982301: | bou | 4185 |
| chrX:6983501:6984201: | bou | 2285 |
| chr2R:11198501:11199251: | Syx6 | 20288 |
| chr2R:11196851:11197701: | Syx6 | 21838 |
| chr3L:3789101:3789551: | CG32266 | 1253 |
| chr3L:3787101:3788251: | CG32266 | 2553 |
| chr3R:20829001:20830601: | AdSS | 0 |

| | | |
|---|---|---|
| chr2R:18972851:18973551: | Topors | 240 |
| chrX:4686101:4686651: | Pp2C1 | 252 |
| chr2L:4028101:4029351: | ed | 2026 |
| chr3R:18658501:18659151: | CG31224 | 158 |
| chr3L:4692001:4693001: | RhoGEF64C | 0 |
| chr3L:21049201:21049951: | siz | 14667 |
| chrX:12660051:12660901: | Tis11 | 637 |
| chrX:8060051:8061701: | mys | 0 |
| chr2L:4375051:4375501: | CG17612 | 7234 |
| chrX:11805051:11805951: | Hsc70-3 | 1166 |
| chr2R:16104051:16104851: | Rho1 | 1365 |
| chr3L:5654201:5654851: | lin-28 | 0 |
| chr3R:8658651:8659801: | bel | 960 |
| chrX:4684201:4684801: | Pp2C1 | 2102 |
| chr2L:4820201:4821751: | CG15628 | 23 |
| chr2R:10540201:10543351: | lola | 0 |
| chrX:16943351:16944701: | RhoGAP15B | 62 |
| chr2R:23203801:23204701: | CG34371 | 0 |
| chr3L:22720301:22721401: | mael | 0 |
| chrX:6354051:6354651: | kdn | 3423 |
| chr2R:13219101:13219651: | CG4630 | 0 |
| chrX:5885351:5886151: | CG16721 | 5483 |
| chr3R:14277651:14279601: | trx | 7302 |
| chr2L:3805401:3806751: | Reph | 1857 |
| chrX:17170401:17171151: | baz | 10396 |
| chr3L:18890401:18891401: | CG3961 | 0 |
| chr3L:13113651:13114551: | trn | 0 |
| chrX:7959051:7959551: | CG15330 | 2988 |
| chrX:10930451:10931751: | Myo10A | 0 |
| chr2R:21480451:21481201: | Sdc | 0 |
| chr2R:16110451:16111501: | Asph | 121 |
| chr3R:22673201:22674501: | Gclm | 12009 |
| chr3R:21198851:21199451: | CG5862 | 15628 |
| chr3L:1485551:1486151: | stet | 3396 |
| chr3L:19908101:19909401: | Su | 9 |
| chr3R:18167751:18169351: | wrd | 0 |
| chr3R:14723501:14724351: | kibra | 0 |
| chr2R:14755651:14756251: | CG10151 | 37 |
| chr3R:30270651:30271801: | hdc | 6131 |
| chrX:6985651:6987651: | bou | 0 |
| chr3R:11763301:11764301: | Lk6 | 156 |
| chr2R:16338751:16339301: | Vha44 | 2115 |
| chr2R:18288401:18289251: | CG14502 | 0 |
| chrX:4690751:4691251: | ctp | 2952 |
| chr2R:13038601:13039201: | CG13323 | 8055 |

| | | |
|---|---|---|
| chr2R:6195801:6196751: | Ptr | 0 |
| chr2L:21753501:21754151: | step | 3315 |
| chrX:13278201:13279151: | HDAC4 | 6481 |
| chr2R:13565851:13566301: | CG42807 | 6458 |
| chrX:7095901:7096351: | Sxl | 1702 |
| chr2R:18805951:18806601: | MFS14 | 2225 |
| chr3L:14780951:14781551: | mop | 4823 |
| chr3R:9235951:9237801: | pum | 0 |
| chr2R:22403151:22404001: | CG6044 | 176 |
| chr2R:13826001:13827101: | CG6329 | 0 |
| chr3L:14201001:14201501: | saturn | 1822 |
| chrX:19866001:19867201: | Hers | 0 |
| chrX:6866051:6866501: | CG14434 | 693 |
| chr2L:4031051:4031551: | ed | 0 |
| chr2R:15931101:15932051: | bdg | 0 |
| chrX:3861101:3861551: | CG2930 | 7422 |
| chr3R:27281101:27282151: | mrt | 510 |
| chrX:3136151:3137051: | N | 1282 |
| chrX:5086201:5087751: | rg | 443 |
| chr2R:16102651:16103751: | Rho1 | 0 |
| chr2R:10417801:10418751: | Prx2540-2 | 0 |
| chrX:9466251:9466951: | CG44815 | 20040 |
| chrX:8051301:8053701: | fs | 7735 |
| chr2R:22798201:22798701: | CG42260 | 0 |
| chrX:3691301:3692801: | Mnt | 19501 |
| chrX:12661351:12662001: | Tis11 | 1937 |
| chrX:6352151:6353601: | kdn | 1523 |
| chr2R:9547951:9548601: | CG1888 | 0 |
| chr3R:11542851:11543601: | dpr4 | 0 |
| chr2L:9607851:9608601: | gcm2 | 0 |
| chr2L:16496401:16496951: | Tpr2 | 4862 |
| chr2R:15372851:15373551: | Pms2 | 1332 |
| chr2R:19141501:19143401: | ena | 0 |
| chr3R:24046501:24048401: | Rox8 | 0 |
| chr2R:14306901:14308451: | Shroom | 0 |
| chr3L:21046951:21048401: | siz | 12417 |
| chr2R:20831601:20832151: | sktl | 3877 |
| chr3L:15146601:15147401: | FucTA | 0 |
| chr3L:1746601:1747151: | CG13921 | 0 |
| chrX:5886601:5887451: | CG16721 | 6733 |
| chr2R:14981651:14982751: | pcs | 389 |
| chr3R:25646951:25648301: | jigr1 | 9109 |
| chr2L:5981701:5983801: | eIF4A | 0 |
| chr2R:11597501:11598201: | tou | 17844 |
| chr2R:8236901:8238151: | pnut | 0 |

| | | |
|---|---|---|
| chr3R:23345351:23348151: | DNApol-epsilon255 | 0 |
| chr2R:17097051:17098101: | GstS1 | 1400 |
| chrX:7966901:7968351: | sws | 0 |
| chrX:11806901:11807801: | Hsc70-3 | 0 |
| chr3R:18007601:18008051: | DNaseII | 11324 |
| chr2R:9291951:9293101: | Drep2 | 96 |
| chrX:7096951:7097701: | Sxl | 352 |
| chr3L:4102401:4103001: | CG14995 | 255 |
| chr2R:22796701:22798001: | CG42260 | 288 |
| chrX:13276901:13278001: | HDAC4 | 7631 |
| chrX:3682401:3682951: | Mnt | 10601 |
| chr2L:7887051:7887901: | CG7102 | 0 |
| chr2R:14432051:14432551: | phyl | 0 |
| chrX:18647101:18647501: | Rip11 | 147 |
| chrX:9042101:9043001: | Nost | 11897 |
| chr2R:11202101:11203601: | Syx6 | 15938 |
| chr3R:14722101:14722851: | kibra | 1410 |
| chr3L:16572251:16572701: | Mipp1 | 633 |
| chr3R:5672301:5673051: | Atg17 | 2589 |
| chr3R:9226201:9227651: | pum | 10031 |
| chr3R:23887001:23887601: | Pli | 2451 |
| chr3L:19906801:19907551: | Su | 1859 |
| chr2R:22401801:22402551: | babos | 0 |
| chr3R:26041601:26042501: | gro | 0 |
| chr3R:28325951:28327501: | larp | 8939 |
| chrX:1232501:1233201: | CG3638 | 3483 |
| chr2L:2752551:2756001: | Pgk | 0 |
| chr3R:30272551:30273501: | hdc | 4431 |
| chr3L:347551:348151: | mth | 595 |
| chrX:5746651:5747351: | IntS6 | 0 |
| chr3R:9052651:9053551: | Kdm2 | 157 |
| chr2L:8957651:8958651: | CG31886 | 400 |
| chr2L:16485501:16487301: | dac | 0 |
| chr3R:14817701:14818451: | CG14857 | 15929 |
| chr2R:10427751:10431001: | RanBPM | 0 |
| chr3L:22402751:22403301: | Ten-m | 4562 |
| chr2R:17592751:17593751: | Smurf | 0 |
| chr2L:5542751:5543351: | Lam | 272 |
| chr3R:18912751:18913201: | CG14291 | 0 |
| chr3R:16452801:16453451: | Pak3 | 370 |
| chrX:6867851:6868601: | CG14434 | 658 |
| chr2R:21712851:21715001: | CG30403 | 1239 |
| chr2L:160651:162101: | spen | 1620 |
| chr2R:22685701:22687101: | Ppa | 1886 |
| chrX:3137901:3138401: | N | 3032 |

| | | |
|---|---|---|
| chr3R:11758301:11762051: | Lk6 | 2406 |
| chr2R:12452951:12454501: | Lac | 0 |
| chr2R:5772951:5773501: | ZnT41F | 9357 |
| chr3R:18298051:18298701: | l | 178 |
| chr3L:16408051:16409251: | fax | 2577 |
| chr2R:20595851:20596901: | HnRNP-K | 8301 |
| chr2L:19161351:19161901: | CG17568 | 19801 |
| chr3L:22723101:22723551: | CG14451 | 1011 |
| chrX:18243201:18244151: | upd2 | 0 |
| chr2L:20638201:20638801: | CG31680 | 1320 |
| chrX:2336301:2336751: | Ilp6 | 1644 |
| chrX:12663251:12663751: | Tis11 | 3837 |
| chr3R:11773251:11775651: | l | 3477 |
| chr3R:22133301:22133901: | SKIP | 0 |
| chrX:6283401:6283901: | CG11700 | 0 |
| chr3L:17658451:17659451: | CG7484 | 3835 |
| chr3L:20398501:20399001: | trbl | 2608 |
| chr3L:17988551:17990351: | CG32192 | 49966 |
| chr3L:9465751:9466401: | CG44838 | 4702 |
| chr2R:8145751:8146401: | kermit | 10253 |
| chr2R:9198601:9198951: | unpg | 3397 |
| chrX:10738651:10739601: | ras | 4901 |
| chrX:6988651:6989451: | ogre | 1867 |
| chr3R:6810451:6811301: | Dfd | 18616 |
| chr2R:16335601:16336301: | Hmgs | 0 |
| chrX:17710651:17711301: | CG42684 | 508 |
| chrX:1233701:1235751: | CG3638 | 933 |
| chr3R:18913701:18914401: | Xrp1 | 567 |
| chr3R:19163701:19165151: | sqz | 0 |
| chr3R:21059801:21061251: | Mvl | 0 |
| chr2L:14489851:14491201: | noc | 0 |
| chr2R:19143801:19144451: | ena | 2170 |
| chr2L:11968801:11969351: | CG16965 | 12240 |
| chr3R:24295451:24296151: | crb | 374 |
| chr2R:22933851:22934701: | CG3788 | 0 |
| chr2R:16585351:16586101: | RpLP2 | 28 |
| chrX:10473901:10474701: | nocte | 0 |
| chr2L:15498951:15499951: | lace | 0 |
| chrX:4693951:4694551: | ctp | 6152 |
| chr3L:20695151:20696001: | kni | 0 |
| chr2R:19275201:19276001: | rib | 5124 |
| chrX:17834001:17834501: | CG6398 | 3174 |
| chr3R:8669001:8669751: | p | 933 |
| chrX:17159951:17160951: | baz | 0 |
| chr2R:13829051:13829651: | CG6329 | 2681 |

| | | |
|---|---|---|
| chr2L:7984051:7984651: | Snoo | 0 |
| chrX:9684101:9687151: | nej | 0 |
| chr3L:14634101:14635001: | dlp | 1501 |
| chr3R:16249901:16250851: | bor | 13264 |
| chr3R:9054151:9054901: | Kdm2 | 1657 |
| chrX:3694151:3696151: | Mnt | 22351 |
| chr2R:8064801:8065801: | CG12769 | 0 |
| chrX:16780201:16780751: | if | 2683 |
| chr2R:13035051:13035751: | CG13323 | 11505 |
| chr3R:30450151:30450751: | Osi23 | 5106 |
| chr3L:20784451:20785751: | CG11399 | 446 |
| chrX:11154251:11155401: | Dlic | 0 |
| chr3L:15514251:15514901: | CG13454 | 2155 |
| chr3R:28634301:28634951: | CG10011 | 148 |
| chr3L:5664351:5665101: | CG46320 | 231 |
| chrX:514351:514751: | arg | 4917 |
| chr2R:21364351:21364901: | CG17974 | 13774 |
| chr2L:15264401:15265051: | yuri | 0 |
| chr2L:11804401:11805201: | crol | 4237 |
| chrX:8050051:8050551: | fs | 10885 |
| chrX:10209451:10210151: | CG15309 | 2223 |
| chrX:7969451:7970001: | sws | 1216 |
| chr2R:8648751:8650501: | ptc | 0 |
| chr2R:14970051:14970501: | ckn | 6760 |
| chr3R:28339501:28340101: | Gfat2 | 0 |
| chrX:3679851:3680451: | Mnt | 8051 |
| chrX:12649601:12650351: | CkIalpha | 3275 |
| chr3R:28324751:28325351: | larp | 11089 |
| chrX:18864301:18865351: | CG34401 | 352 |
| chr3R:30794651:30795101: | wts | 11518 |
| chr2R:7779401:7780251: | CG45093 | 2386 |
| chr3R:9239751:9240301: | D1 | 353 |
| chr3L:12804051:12805201: | Atg1 | 0 |
| chr2L:5879201:5880201: | rau | 0 |
| chrX:18634651:18635201: | wgn | 0 |
| chr2R:12019601:12020201: | otk | 0 |
| chr2R:19423451:19425201: | CalpA | 0 |
| chrX:8674651:8675201: | CG12772 | 1862 |
| chr2L:2454251:2455101: | dpp | 25880 |
| chr3L:15329251:15330101: | Toll-6 | 6591 |
| chrX:11494901:11495551: | Drak | 1767 |
| chr3R:25049901:25050451: | CG11791 | 286 |
| chr3R:9394251:9395051: | FER | 0 |
| chr3R:13074501:13075051: | CG43063 | 11173 |
| chrX:12664951:12665651: | Tis11 | 5537 |

| | | |
|---|---|---|
| chr2L:21104951:21105551: | CG46307 | 384 |
| chrX:19199951:19200351: | CG7990 | 1628 |
| chr3L:14205051:14205551: | saturn | 1729 |
| chr2R:16210051:16210901: | Shark | 7168 |
| chr2L:14234051:14234901: | Dyrk2 | 0 |
| chr3L:16410101:16412051: | fax | 0 |
| chr3L:19559301:19559851: | CG9300 | 6882 |
| chrX:2333851:2334851: | Ilp6 | 0 |
| chr2R:8724151:8724801: | gcl | 0 |
| chrX:5744201:5744801: | IntS6 | 2453 |
| chr3L:14785251:14785851: | bmm | 561 |
| chr2R:21545251:21547351: | CG30283 | 4679 |
| chr2L:11970251:11971651: | CG16965 | 9940 |
| chr3L:5665301:5665901: | CG46320 | 1181 |
| chr3L:6964101:6964651: | sgl | 107 |
| chr2L:7530351:7531351: | Obp28a | 32987 |
| chr3R:12369101:12369601: | GstD1 | 0 |
| chr2R:24743901:24744601: | Usp15-31 | 153 |
| chr3L:20620401:20622501: | knrl | 0 |
| chr3R:19165451:19166251: | sqz | 1241 |
| chrX:19753951:19754501: | Pmp70 | 3182 |
| chrX:4678001:4679501: | fzr | 36 |
| chr2R:11592151:11594451: | tou | 21594 |
| chr2R:9278651:9279451: | Pkn | 5646 |
| chrX:12753801:12754451: | ade5 | 6166 |
| chr3R:30465551:30466301: | PH4alphaEFB | 0 |
| chr3L:16645551:16646051: | Abl | 1831 |
| chr2L:13165551:13166151: | DnaJ-H | 0 |
| chr2R:10100551:10101951: | 14-3-3zeta | 875 |
| chr3R:23248751:23249401: | CG4467 | 0 |
| chr2R:16583151:16584401: | RpLP2 | 1728 |
| chr3R:11948851:11949351: | Cad87A | 0 |
| chr2R:8665651:8666601: | Acsl | 0 |
| chr2R:20835701:20836801: | sktl | 0 |
| chr2R:10798451:10799201: | CG33144 | 0 |
| chr3L:8978601:8979201: | CG5644 | 679 |
| chrX:2635851:2636401: | sgg | 1900 |
| chr3L:3225851:3226701: | CG11505 | 1014 |
| chrX:17650851:17652751: | RhoGAPp190 | 0 |
| chrX:5417751:5419051: | spoon | 1816 |
| chr3L:10225951:10226401: | scramb1 | 712 |
| chr3L:18846001:18846651: | Indy | 0 |
| chr3L:18621101:18621651: | not | 478 |
| chr2L:19157851:19158851: | l | 22619 |
| chr3L:19081151:19081901: | Mkp3 | 4525 |

| | | |
|---|---|---|
| chr3R:25588201:25588801: | CG4582 | 5023 |
| chrX:3676501:3678801: | Mnt | 4701 |
| chr2L:14233201:14233751: | Dyrk2 | 375 |
| chrX:21033201:21033751: | CG1529 | 1252 |
| chr3L:20401251:20402151: | trbl | 0 |
| chr2L:11806251:11809601: | crol | 0 |
| chr3R:9056301:9058251: | Kdm2 | 3807 |
| chrX:10211351:10212351: | CG15309 | 23 |
| chrX:12666351:12666751: | Tis11 | 6937 |
| chr3L:18148051:18148551: | CG7330 | 372 |
| chr3L:17972251:17973501: | CG5290 | 59687 |
| chr3R:11351501:11353801: | KP78a | 0 |
| chr3L:20507901:20508451: | CG5078 | 1870 |
| chr2R:21056551:21057151: | shg | 46 |
| chr3R:19321601:19322651: | Dl | 3566 |
| chr2R:8647201:8648351: | ptc | 1298 |
| chr2R:18156651:18157501: | lolal | 691 |
| chrX:5661951:5663301: | CG42265 | 0 |
| chr3R:16247801:16248301: | bor | 15814 |
| chrX:3696701:3697351: | Rala | 22284 |
| chr2L:5546801:5547501: | Oscillin | 0 |
| chrX:2636801:2637501: | sgg | 2850 |
| chr2L:3631801:3633201: | CG15418 | 10098 |
| chrX:12006601:12008151: | CG1806 | 0 |
| chrX:10741851:10742801: | ras | 1701 |
| chrX:13746851:13747851: | Grip91 | 2169 |
| chrX:3332451:3333101: | CG10793 | 16050 |
| chr3R:17716901:17718001: | osa | 349 |
| chr3R:19147351:19148051: | Ppcs | 0 |
| chrX:3141951:3142651: | N | 7082 |
| chr2R:17672501:17673001: | P32 | 0 |
| chr3L:18167001:18167501: | CG13699 | 4480 |
| chr3L:9461651:9462951: | CG44838 | 602 |
| chr3R:22666351:22667901: | Gclm | 18609 |
| chr3R:18396651:18397851: | Vti1b | 0 |
| chr2R:8667151:8669601: | Acsl | 1339 |
| chr2R:10432151:10432851: | RanBPM | 2444 |
| chr3R:9512151:9513151: | Ras85D | 0 |
| chr2L:2862151:2862801: | CG2991 | 4988 |
| chr2R:8232101:8232701: | dpn | 0 |
| chr3L:1577301:1577851: | CG13917 | 603 |
| chrX:6656851:6657601: | Cdc7 | 2311 |
| chr3L:13227451:13229051: | caps | 0 |
| chr2L:17382501:17383251: | Lrch | 864 |
| chr3L:22407501:22408651: | Ten-m | 0 |

| | | |
|---|---|---|
| chr2L:11066751:11067451: | Samuel | 0 |
| chrX:12750201:12752451: | CG4004 | 5990 |
| chr3L:11832551:11833351: | CycA | 227 |
| chr3L:1882551:1883601: | CG13937 | 0 |
| chrX:15712601:15713451: | CG8184 | 0 |
| chr3R:26036251:26037351: | E | 0 |
| chr3L:21276651:21277301: | AcCoAS | 2942 |
| chr2R:19292701:19293551: | Tab2 | 173 |
| chr3L:22727701:22728401: | CG11367 | 0 |
| chr3R:6966551:6967251: | Antp | 31977 |
| chr3R:25207751:25208701: | Cad96Ca | 0 |
| chr2L:12421701:12422201: | vir-1 | 1238 |
| chrX:17291251:17292201: | CG8661 | 6752 |
| chrX:19850651:19852151: | pico | 0 |
| chr3R:14737851:14739501: | eff | 1818 |
| chr3L:1861251:1862051: | Rap1 | 188 |
| chrX:16091351:16092051: | Tob | 0 |
| chr3L:2466451:2467051: | CG16762 | 8563 |
| chr3R:10132951:10134001: | tws | 5771 |
| chr3R:24031151:24032001: | KrT95D | 0 |
| chrX:15076001:15077001: | Lsd-2 | 0 |
| chr2R:8083051:8083801: | slv | 0 |
| chrX:9666351:9666901: | CG3106 | 3449 |
| chr2R:17851251:17851901: | CG30323 | 1394 |
| chrX:15143101:15143901: | RpL37a | 3735 |
| chr2R:7488151:7490901: | wech | 0 |
| chrX:16798151:16798651: | CG9132 | 2059 |
| chr2R:20110551:20111801: | 18w | 0 |
| chrX:17178201:17179151: | baz | 18196 |
| chr2R:8060301:8061751: | CG12769 | 3803 |
| chr3R:15973251:15974351: | Hel89B | 0 |
| chr2L:7418251:7419101: | chm | 5835 |
| chr3R:8105801:8106701: | puc | 470 |
| chr3L:17563301:17564351: | CycT | 5514 |
| chr3L:4263401:4263851: | CG1273 | 0 |
| chr3L:17971051:17971551: | CG5290 | 58487 |
| chr2L:2450551:2451501: | dpp | 22180 |
| chrX:6266051:6266451: | schlank | 777 |
| chr3R:15665651:15666451: | pxb | 0 |
| chr3R:29214101:29216301: | eEF1gamma | 3871 |
| chrX:19750851:19751301: | Pmp70 | 82 |
| chr3R:31058701:31060051: | dco | 1159 |
| chrX:21053751:21054901: | bves | 0 |
| chrX:3845201:3846201: | CG2901 | 17005 |
| chr2L:14508801:14509351: | CG33648 | 14961 |

| | | |
|---|---|---|
| chr3L:15815351:15816151: | fwe | 259 |
| chr2R:19668851:19669951: | hrg | 0 |
| chr2R:23658851:23659801: | Pde8 | 1285 |
| chrX:6533901:6534651: | dx | 0 |
| chr2R:16579201:16581051: | Cdk4 | 0 |
| chrX:2125451:2126001: | ph-d | 2166 |
| chr3R:9220201:9221001: | pum | 16681 |
| chr3R:24630001:24630851: | Wsck | 0 |
| chr3L:7225201:7225851: | CG14826 | 8661 |
| chr2R:14964201:14965851: | ckn | 910 |
| chr2R:13449151:13450201: | cnn | 274 |
| chr2R:15329151:15329651: | CG8160 | 881 |
| chrX:5094201:5094851: | rg | 8443 |
| chrX:1899301:1899801: | arm | 845 |
| chr3R:10035151:10035601: | Mical | 7294 |
| chrX:2254851:2255601: | CG3091 | 655 |
| chr2R:24389251:24390601: | mAChR-A | 0 |
| chr3L:1349451:1349901: | 312 | 1215 |
| chr3L:18439651:18440451: | skl | 0 |
| chr3R:11779551:11780951: | l | 424 |
| chr3R:8333651:8335351: | Poxm | 0 |
| chr3R:15664551:15665351: | pxb | 832 |
| chr2L:8534651:8536201: | CG17834 | 5511 |
| chr3R:25183301:25185301: | dan | 0 |
| chr3R:11604401:11605301: | CG6791 | 14256 |
| chr2L:19154551:19155301: | l | 19319 |
| chrX:13624701:13625851: | NFAT | 0 |
| chr3L:7344751:7345501: | Cln7 | 0 |
| chr2L:3789451:3790151: | bark | 2285 |
| chr3L:2489851:2490401: | CG1146 | 1048 |
| chr3R:10108951:10110001: | Fmr1 | 120 |
| chr3L:10660051:10660701: | CG32066 | 312 |
| chrX:9544301:9544901: | CG32700 | 5937 |
| chr2R:8670101:8670951: | Acsl | 4289 |
| chr3L:1885101:1885951: | Tmhs | 0 |
| chrX:12470151:12470751: | CG42258 | 6624 |
| chr3R:14714051:14714801: | kibra | 9460 |
| chrX:16089151:16089751: | Tob | 1744 |
| chr2L:6339001:6339751: | Cpr | 166 |
| chr3L:17995251:17995901: | CG32192 | 44416 |
| chr3R:7973951:7974601: | CD98hc | 3827 |
| chr2R:10237951:10239601: | Hdc | 0 |
| chr2R:21645401:21647051: | CG10082 | 0 |
| chr3R:9033551:9034551: | hyx | 4209 |
| chr3R:22898451:22899551: | klg | 0 |

| | | |
|---|---|---|
| chr3L:17245501:17246101: | CG7707 | 2066 |
| chr3R:15280501:15281651: | Tm1 | 0 |
| chr3L:2228301:2229451: | Oseg2 | 195 |
| chr2L:19820551:19821001: | CG13962 | 6020 |
| chrX:8041001:8044401: | CG2233 | 3485 |
| chrX:19505651:19506351: | Pfrx | 10147 |
| chrX:8420651:8421751: | lawc | 8994 |
| chr3R:18670701:18671351: | CG31230 | 328 |
| chr3R:24218501:24219251: | CG17786 | 0 |
| chr3L:17968451:17969251: | CG5290 | 55887 |
| chr2L:2885751:2887401: | lilli | 0 |
| chr3L:6985801:6986251: | CG43439 | 1216 |
| chr3L:7910801:7911301: | pbl | 653 |
| chr2L:430801:431401: | ex | 0 |
| chr2R:11615801:11616551: | tou | 0 |
| chr2L:16718451:16719151: | Cyt-c-p | 781 |
| chr3R:19325951:19326951: | Dl | 0 |
| chrX:2122751:2124001: | ph-d | 0 |
| chr2R:22678201:22678951: | RYBP | 7615 |
| chr3L:19086051:19087501: | Mkp3 | 0 |
| chr2L:8416051:8416901: | Akap200 | 362 |
| chr2L:15478051:15478901: | sna | 0 |
| chr3R:6893151:6893901: | ftz | 28828 |
| chr2R:18673051:18673901: | edl | 0 |
| chrX:521251:524001: | elav | 0 |
| chr2L:7991251:7992251: | pes | 1933 |
| chrX:12471251:12472951: | CG42258 | 7724 |
| chr3R:7973201:7973701: | CD98hc | 4727 |
| chr2R:20587451:20588701: | HnRNP-K | 0 |
| chr3R:29876301:29877351: | CG31038 | 0 |
| chr2R:10078001:10078601: | egr | 0 |
| chr2R:9566401:9567551: | CG1809 | 4764 |
| chrX:21056501:21057101: | bves | 2539 |
| chrX:20411501:20412251: | CG15459 | 1330 |
| chr3R:22041501:22043201: | how | 0 |
| chr3L:15588001:15588451: | CG7656 | 0 |
| chr2L:11516551:11517101: | Dlg5 | 0 |
| chr3R:25996651:25997651: | E | 0 |
| chr2R:10551651:10552451: | psq | 5437 |
| chrX:11816701:11819151: | rudhira | 0 |
| chr3L:18822551:18823251: | Cat | 0 |
| chrX:18297901:18298251: | upd1 | 11040 |
| chrX:7307301:7308201: | brk | 0 |
| chr3R:25472001:25473201: | Fur1 | 0 |
| chr2R:16187201:16188201: | clu | 618 |

| | | |
|---|---|---|
| chrX:7336801:7337551: | CBP | 0 |
| chr3L:19894951:19898151: | Su | 11259 |
| chr2R:14962101:14963151: | aPKC | 0 |
| chr3L:9137401:9138101: | nwk | 0 |
| chr3L:12146901:12147351: | Nrx-IV | 383 |
| chr3R:24092501:24093051: | Syx1A | 12712 |
| chr3R:16242151:16243051: | tara | 16501 |
| chr3R:21186601:21187951: | Snmp1 | 18397 |
| chr3R:9247051:9247601: | CG8420 | 403 |
| chrX:6997051:6999051: | Inx2 | 0 |
| chr3R:25637251:25637901: | jigr1 | 0 |
| chrX:16087351:16087901: | Tob | 3594 |
| chr2R:5782101:5783101: | ZnT41F | 0 |
| chr3L:13476901:13477801: | stv | 0 |
| chr2L:10967101:10967801: | CG33129 | 2474 |
| chr2R:7492201:7492651: | Coop | 0 |
| chr3L:19292201:19293651: | Gbs-76A | 0 |
| chr2L:4997251:4998051: | Rtnl1 | 11669 |
| chrX:2642251:2643151: | sgg | 8300 |
| chrX:3671901:3672701: | Mnt | 101 |
| chrX:17812101:17812701: | CG12986 | 6984 |
| chrX:3451051:3452701: | CG12535 | 2109 |
| chr2R:21522051:21522651: | Egfr | 0 |
| chrX:17657351:17659501: | beta-Spec | 0 |
| chr3R:31222401:31222951: | CG12054 | 1700 |
| chr2L:13512401:13514651: | CG33640 | 13190 |
| chr3R:9417401:9419901: | ps | 0 |
| chrX:11787101:11787551: | Karl | 0 |
| chr3R:21641001:21642451: | E2f1 | 18420 |
| chrX:9692551:9693201: | btd | 985 |
| chrX:13412601:13413401: | Set2 | 0 |
| chr3L:11196801:11197351: | GlcAT-P | 185 |
| chr2R:17027651:17028751: | RhoGEF2 | 462 |
| chr2R:16607651:16608201: | CG5065 | 0 |
| chr2R:22911451:22912301: | CG30187 | 6489 |
| chr2R:10437701:10438551: | Galphao | 0 |
| chr3L:11707701:11708351: | CG11658 | 202 |
| chr2R:13940751:13942251: | shot | 0 |
| chr2R:24986651:24987251: | emp | 582 |
| chr2L:2887801:2889701: | lilli | 1873 |
| chr2R:24736451:24737151: | Mid1 | 0 |
| chr3R:7102851:7103951: | e | 8917 |
| chr2R:20106101:20107101: | 18w | 4410 |
| chr2R:24767901:24768601: | GstE12 | 0 |
| chrX:7305951:7307051: | brk | 888 |

| | | |
|---|---|---|
| chr3R:18922951:18924751: | Mpc1 | 3908 |
| chr2L:9447951:9450151: | CG33723 | 7212 |
| chr2L:3606301:3607001: | odd | 0 |
| chr3R:30803001:30804301: | wts | 2318 |
| chr3R:20753001:20753651: | Syp | 3078 |
| chr3L:14968051:14969701: | Tom | 0 |
| chr2R:24543051:24543901: | ITP | 5631 |
| chr2L:19426251:19426901: | Pax | 0 |
| chr3L:21843101:21843901: | CG14563 | 716 |
| chr3R:18573101:18574201: | fray | 11 |
| chr2L:16715551:16716851: | CG31808 | 0 |
| chr3R:16240951:16241801: | tara | 15301 |
| chr3L:22851101:22851801: | CG32462 | 1711 |
| chr2R:20960601:20961751: | hbn | 0 |
| chr3L:6495751:6496701: | sfl | 72 |
| chrX:15593351:15594501: | CG9220 | 0 |
| chr2R:17458351:17459001: | CG14478 | 689 |
| chrX:2283401:2283901: | Vml | 7580 |
| chr2R:14768401:14769451: | Spred | 0 |
| chr2L:7993401:7994901: | pes | 0 |
| chr2L:3538451:3540401: | drm | 0 |
| chrX:9693501:9694401: | btd | 0 |
| chrX:16085801:16086451: | CG8958 | 2570 |
| chrX:2325501:2326401: | CG14050 | 0 |
| chrX:12673601:12674451: | Tis11 | 14187 |
| chrX:17733651:17734201: | CG42684 | 23508 |
| chr3R:14020451:14021301: | Cyp6d5 | 7927 |
| chr3R:29063701:29065251: | CG11873 | 662 |
| chrX:12640551:12641251: | Tomosyn | 8947 |
| chr3R:31724501:31726251: | ttk | 10635 |
| chr2L:4998751:4999301: | Rtnl1 | 10419 |
| chr2R:10553751:10554551: | NA-psq | 3337 |
| chrX:9163901:9164451: | mei-P26 | 647 |
| chr3R:22368901:22370151: | PyK | 1366 |
| chr3R:22043951:22044401: | how | 1761 |
| chr3L:19929001:19929651: | RhoGDI | 259 |
| chr3R:15250251:15250801: | CG42404 | 158 |
| chr3L:17999251:18000251: | CG32192 | 40066 |
| chr3R:21640051:21640701: | E2f1 | 20170 |
| chr3L:16845101:16845651: | Zcchc7 | 0 |
| chrX:5899351:5900951: | Act5C | 0 |
| chr3L:11084401:11085101: | JIL-1 | 8092 |
| chr3L:14754701:14760551: | CG42507 | 0 |
| chrX:7944751:7945551: | l | 1357 |
| chr3R:13814351:13815551: | tal-1A | 1243 |

| | | |
|---|---|---|
| chr3R:18274851:18275551: | eIF1A | 187 |
| chr3R:16274451:16275151: | gish | 2000 |
| chr3L:10664451:10665851: | simj | 0 |
| chr3L:17964651:17965451: | CG5290 | 52087 |
| chrX:7709251:7710451: | CHES-1-like | 0 |
| chrX:369551:370251: | ac | 0 |
| chr2R:18309551:18310151: | Tango8 | 1671 |
| chr3L:15619551:15620301: | CG7372 | 0 |
| chr3L:19294651:19295351: | fal | 88 |
| chr3L:9004701:9005401: | Doc3 | 0 |
| chr2R:13939501:13940251: | shot | 1859 |
| chr2L:8419751:8420651: | Akap200 | 4062 |
| chr3R:10139751:10140501: | tws | 0 |
| chrX:1209251:1210101: | CG11382 | 0 |
| chr3L:17569901:17570401: | CycT | 12114 |
| chr3R:23334301:23335001: | pnt | 11166 |
| chr3L:13904251:13905001: | dysc | 1016 |
| chr2L:9874451:9875001: | nAChRalpha6 | 11249 |
| chr2L:14688901:14689951: | osp | 0 |
| chr3L:3060051:3060851: | CG16753 | 4973 |
| chr2R:10555051:10555751: | psq | 2137 |
| chrX:11370101:11370601: | dlg1 | 437 |
| chr3R:14624301:14624851: | put | 852 |
| chr2L:5000251:5001501: | Rtnl1 | 8219 |
| chr3L:18138951:18139701: | geko | 0 |
| chr3R:13063751:13064651: | sim | 5997 |
| chrX:5770401:5771251: | CG3097 | 0 |
| chrX:10823301:10824451: | Imp | 0 |
| chrX:6338701:6339401: | CG3842 | 940 |
| chr2R:14958351:14959401: | aPKC | 3568 |
| chr3R:30805601:30806251: | wts | 368 |
| chrX:2323801:2324301: | CG14050 | 1916 |
| chr2L:19423301:19424301: | CG16771 | 0 |
| chrX:2155751:2156451: | wapl | 2532 |
| chr3R:8680751:8680951: | CG33325 | 718 |
| chr3R:14055851:14056401: | foxo | 544 |
| chr3R:15285851:15287301: | CG45218 | 3498 |
| chr3L:1463501:1464101: | rho | 0 |
| chr3L:4233601:4234101: | ImpL2 | 2046 |
| chr3R:21638251:21639101: | InR | 18931 |
| chr2R:16220951:16221551: | mrj | 242 |
| chr3R:13976001:13976601: | Orc2 | 11117 |
| chrX:12596001:12597001: | Sec16 | 0 |
| chr3L:1586201:1587051: | CG13917 | 9503 |
| chr2R:20996201:20998851: | CG10543 | 0 |

| | | |
|---|---|---|
| chr2L:8277401:8278751: | CG8086 | 19900 |
| chr2R:24891251:24891651: | NaCP60E | 0 |
| chrX:8271251:8271851: | Corp | 0 |
| chr2L:3476351:3477001: | Thor | 1433 |
| chr3L:2896351:2896801: | Shab | 1335 |
| chr2L:4301351:4301901: | tutl | 18204 |
| chrX:5636351:5638101: | CG42492 | 10386 |
| chr3L:10666351:10667051: | simj | 1613 |
| chr3L:21807901:21808601: | eg | 102 |
| chr3L:14177251:14178551: | D | 69 |
| chrX:5901451:5902401: | Act5C | 591 |
| chr2L:8541501:8544651: | Sema1a | 0 |
| chr3R:13812451:13813451: | tal-1A | 0 |
| chr2R:23666551:23667351: | Pde8 | 8985 |
| chr3R:31621601:31622401: | kek6 | 0 |
| chr3R:14056601:14058001: | foxo | 0 |
| chr3L:3336651:3337301: | kst | 0 |
| chrX:18131651:18132151: | CG43133 | 1474 |
| chr3R:14017301:14018301: | Cyp6d5 | 10927 |
| chr3L:13236801:13237351: | caps | 8110 |
| chrX:12437651:12438151: | Pkcdelta | 5428 |
| chr2R:20582001:20583101: | qsm | 490 |
| chr3L:13667051:13668101: | bru3 | 135299 |
| chr2L:5906901:5907801: | bchs | 0 |
| chr3L:18867401:18868051: | Dysb | 0 |
| chrX:10822001:10823051: | Imp | 1226 |
| chr3L:492351:493001: | CG34267 | 7450 |
| chr2R:14332001:14332651: | CG8613 | 19580 |
| chr3R:11977051:11977501: | CG10005 | 0 |
| chr2R:17462051:17462701: | CG14478 | 4389 |
| chr2L:5997101:5998301: | lid | 1200 |
| chr2L:867151:868801: | aru | 0 |
| chr2R:21497151:21497901: | Fkbp14 | 28 |
| chr3L:14972201:14972801: | Brd | 0 |
| chr3R:21637051:21637651: | InR | 17731 |
| chr2L:8162351:8163351: | Piezo | 0 |
| chr3L:12832351:12832851: | CG10960 | 10255 |
| chrX:7704551:7707601: | CHES-1-like | 1784 |
| chrX:17396951:17397601: | B-H1 | 0 |
| chr3R:7107401:7110351: | Mlp84B | 7381 |
| chr3R:16071901:16072501: | CG10185 | 4308 |
| chrX:16967501:16968001: | Ubr1 | 678 |
| chr3R:16277551:16278101: | gish | 5100 |
| chrX:11372601:11373301: | dlg1 | 2937 |
| chrX:1991651:1992351: | PIG-K | 17013 |

| | | |
|---|---|---|
| chr3L:11046351:11047201: | CG43693 | 4170 |
| chr3L:1546201:1547201: | Pcyt1 | 110 |
| chrX:19782851:19783501: | zld | 47 |
| chr2R:25061501:25062101: | gsb | 0 |
| chrX:17737901:17739501: | CG42684 | 27758 |
| chr2L:826051:827051: | dock | 83 |
| chr2L:3477951:3478751: | Thor | 0 |
| chrX:11822951:11823651: | rudhira | 6054 |
| chrX:15483051:15483601: | CG6340 | 678 |
| chr3R:17055951:17056901: | Dad | 1945 |
| chr2R:19411051:19411851: | hts | 13052 |
| chr3L:9373301:9374251: | Hsp22 | 371 |
| chr2R:23978351:23978951: | Sox14 | 0 |
| chr2R:10558501:10559201: | NA-psq | 614 |
| chr3L:16043551:16044201: | Diap1 | 6833 |
| chr2L:128601:129651: | CG3164 | 1140 |
| chrX:3835301:3836351: | CG2901 | 7105 |
| chr3R:14748651:14749851: | smp-30 | 1395 |
| chr2L:3888651:3889151: | capu | 13709 |
| chr3R:11568701:11569551: | Mrp4 | 0 |
| chr3R:23278701:23279701: | orb | 1155 |
| chrX:7299551:7301251: | brk | 6688 |
| chr2R:9978751:9979551: | eve | 0 |
| chr3R:30765301:30766201: | zfh1 | 0 |
| chrX:6543851:6544301: | CG34417 | 3081 |
| chrX:18290601:18291101: | upd3 | 13369 |
| chrX:9578951:9579401: | LPCAT | 586 |
| chr3R:6998951:6999801: | Antp | 0 |
| chr2R:16129051:16130701: | spin | 4071 |
| chr3L:15235151:15235901: | Tollo | 0 |
| chr2R:8174101:8175151: | LRP1 | 0 |
| chrX:5904151:5904901: | CG4020 | 845 |
| chr3R:29069151:29070301: | CG11873 | 6112 |
| chr2R:11175201:11175601: | CG13229 | 2996 |
| chr3R:15094401:15095701: | CG6966 | 0 |
| chr2R:10559451:10560201: | NA-psq | 1564 |
| chr3L:14219501:14220151: | CG7768 | 1298 |
| chr3R:13364101:13365451: | Dic1 | 0 |
| chrX:10224551:10225201: | CG34408 | 4442 |
| chrX:15484601:15485101: | Gmap | 1821 |
| chr2L:8424651:8425351: | grk | 8247 |
| chr2L:6939651:6940251: | CG18304 | 2563 |
| chr2R:18514701:18515251: | CG43066 | 128 |
| chr2R:13539551:13540201: | Vmat | 5301 |
| chr3R:16279851:16282001: | gish | 7400 |

| | | |
|---|---|---|
| chr2R:25214901:25215601: | Kr | 11010 |
| chrX:20489601:20490051: | CG15461 | 3422 |
| chr3L:16659951:16661651: | Baldspot | 0 |
| chr2R:14954251:14955001: | aPKC | 7968 |
| chr2R:12903951:12904851: | NAT1 | 11143 |
| chr3R:11980151:11980751: | CG18547 | 0 |
| chr3L:19045201:19045851: | nkd | 0 |
| chr2R:11475201:11476101: | inv | 737 |
| chr3R:30763901:30764751: | zfh1 | 1175 |
| chr3L:5633951:5634751: | Blimp-1 | 2969 |
| chr3L:15234151:15234751: | Tollo | 868 |
| chr3L:18005251:18005751: | CG32192 | 34566 |
| chr3L:4430251:4430851: | nAChRbeta1 | 903 |
| chr2L:3825301:3825901: | slp1 | 0 |
| chr3R:17053351:17054551: | Dad | 0 |
| chr3R:7000451:7001451: | Antp | 1224 |
| chr2R:19455501:19457501: | mei-W68 | 0 |
| chr2L:17735501:17736051: | CG43231 | 30821 |
| chr3R:10863851:10864451: | CG6574 | 864 |
| chr2R:16834051:16834451: | CG34460 | 7180 |
| chrX:10865551:10866001: | CG2157 | 3283 |
| chr3R:15290651:15292251: | CG45218 | 0 |
| chr3R:8094001:8094301: | CG7900 | 0 |
| chrX:5593801:5594251: | Vsx1 | 4 |
| chr3L:20493501:20494201: | ZnT77C | 720 |
| chrX:19297851:19299201: | CG8034 | 0 |
| chr3R:4452101:4454101: | CG31522 | 8782 |
| chr2L:7188351:7189051: | CG4496 | 959 |
| chr3R:21632301:21634001: | InR | 12981 |
| chr2R:16031001:16032151: | SP2353 | 0 |
| chr2R:11173351:11173951: | CG13229 | 1146 |
| chr2L:12678051:12678901: | CG15485 | 8338 |
| chr3L:7156101:7157351: | corn | 0 |
| chr2R:11626151:11627101: | Egm | 7129 |
| chr2L:19548051:19548751: | Tep4 | 0 |
| chr2R:6168151:6168751: | EcR | 336 |
| chrX:4507851:4508701: | CG3556 | 21530 |
| chr2R:18171301:18171951: | Dgp-1 | 656 |
| chr2R:10227801:10228651: | CG46321 | 0 |
| chrX:6642951:6643601: | CG14443 | 8993 |
| chr3R:19656401:19656951: | CG6184 | 915 |
| chr2R:9711401:9712951: | dap | 0 |
| chr3R:18931501:18932201: | CG42613 | 2306 |
| chr3L:16046551:16047751: | Diap1 | 3283 |
| chr2R:22382601:22383401: | CG13506 | 0 |

| | | |
|---|---|---|
| chrX:19886601:19887701: | amn | 2359 |
| chrX:13111701:13112551: | CG43313 | 0 |
| chr3R:18196701:18199501: | Ssdp | 94 |
| chr2R:10781651:10783251: | mthl13 | 8208 |
| chr3L:9126901:9128251: | bol | 6154 |
| chrX:2317351:2318101: | CG14050 | 8116 |
| chrX:7937551:7938101: | Ubr3 | 1656 |
| chr2L:5296901:5297401: | vri | 7958 |
| chrX:7702301:7703001: | CHES-1-like | 6384 |
| chr2R:6631201:6632951: | Vha16-1 | 139 |
| chr2R:24727151:24727951: | Mid1 | 8706 |
| chr2R:21502051:21503351: | MESK2 | 0 |
| chr3L:9377151:9377951: | Hsp26 | 0 |
| chr3L:15532301:15532701: | CG43248 | 829 |
| chr3L:4227151:4227601: | ImpL2 | 8546 |
| chr3L:14977401:14978051: | CG3349 | 1569 |
| chrX:6331851:6332551: | CG42240 | 1010 |
| chr2R:19629951:19632551: | sm | 0 |
| chr3L:17577451:17578251: | CycT | 19664 |
| chr2L:15467051:15467501: | sna | 10759 |
| chrX:8657051:8657501: | oc | 6371 |
| chr3R:24767501:24769501: | CG13631 | 0 |
| chr3L:21024651:21027451: | skd | 0 |
| chr3L:6716901:6717451: | ple | 2074 |
| chr3L:1592601:1593551: | CG12104 | 9123 |
| chrX:2292651:2294151: | CG2865 | 0 |
| chr3R:17680301:17682251: | alt | 264 |
| chrX:10816751:10817251: | Imp | 7026 |
| chr2L:4346651:4347151: | CG15429 | 5824 |
| chr3R:6791351:6792101: | Dfd | 0 |
| chr3R:26020751:26022001: | E | 0 |
| chrX:19788001:19789101: | zld | 4454 |
| chr3R:13983051:13984251: | Orc2 | 18167 |
| chr3R:9428051:9428851: | ps | 10112 |
| chr2R:21193101:21193701: | Lapsyn | 0 |
| chr2R:21250201:21251701: | ASPP | 0 |
| chr3R:18583451:18585001: | qin | 6026 |
| chr2R:20511051:20511501: | CG13426 | 1668 |
| chrX:9583501:9586151: | Hex-A | 0 |
| chrX:9650051:9651401: | Ptpmeg2 | 7744 |
| chrX:8225901:8226401: | CG2147 | 14283 |
| chrX:9283601:9284801: | lz | 0 |
| chrX:11828601:11829101: | CG15741 | 8925 |
| chr2L:2955751:2956351: | Rbp9 | 990 |
| chr3R:29073701:29074701: | CG11873 | 10662 |

| | | |
|---|---|---|
| chr2L:6078701:6079551: | Kr-h1 | 3052 |
| chr3R:21630351:21631251: | InR | 11031 |
| chr3L:7263751:7264251: | CG14830 | 0 |
| chr2L:8708751:8709601: | Hnf4 | 10 |
| chr3L:19938801:19939401: | Ac76E | 0 |
| chr3R:29253801:29254251: | stg | 1549 |
| chr3L:5629851:5631151: | Blimp-1 | 0 |
| chr3L:4133851:4134501: | ens | 49 |
| chrX:9173851:9174401: | mei-P26 | 10597 |
| chrX:6330151:6331101: | CG42240 | 0 |
| chr2R:13015401:13016101: | Su | 18901 |
| chrX:10815501:10816101: | Imp | 8176 |
| chr2L:7495201:7496051: | Ziz | 256 |
| chrX:590251:591051: | vnd | 8131 |
| chr3L:14600551:14601051: | HGTX | 1012 |
| chrX:18285651:18286001: | upd3 | 8419 |
| chr3R:20574001:20575001: | TFAM | 2009 |
| chr3L:10674001:10675351: | simj | 9263 |
| chr3R:10614051:10615251: | hth | 24317 |
| chr3L:9124901:9125851: | bol | 8554 |
| chr3L:539151:540551: | klar | 63 |
| chrX:19889151:19889751: | amn | 309 |
| chr3L:4380201:4380801: | DopEcR | 0 |
| chr3R:21094201:21094851: | RhoGAP93B | 2976 |
| chrX:18534201:18534851: | CG15047 | 5519 |
| chrX:15675101:15675751: | CG12708 | 0 |
| chr3R:24259251:24260951: | beta-PheRS | 6573 |
| chr2R:10224751:10225701: | CG46321 | 2256 |
| chr3L:21800301:21800701: | eg | 8002 |
| chr2L:7855101:7855701: | CG14535 | 0 |
| chr3R:12244301:12244851: | Cyp313a3 | 5036 |
| chr3L:13949351:13949901: | CG32137 | 10102 |
| chr3L:15534351:15535701: | CrebA | 0 |
| chrX:2294401:2295001: | Raf | 465 |
| chr3L:12489851:12490501: | CG10660 | 175 |
| chrX:2469501:2470501: | boi | 0 |
| chr2R:15349301:15350301: | unc-5 | 8830 |
| chr2L:12459701:12460451: | CG5418 | 6460 |
| chr3L:3069751:3071001: | CG32486 | 0 |
| chr2R:9929451:9930201: | Etf-QO | 18559 |
| chr3L:16049901:16051201: | Diap1 | 0 |
| chrX:5339901:5340501: | SK | 0 |
| chr2L:16299301:16300051: | CG17328 | 0 |
| chrX:10174751:10175051: | CG43902 | 16318 |
| chrX:22854951:22855701: | fog | 0 |

| | | |
|---|---|---|
| chrX:23044951:23045651: | CG12061 | 8886 |
| chr2L:16519951:16521801: | CG5953 | 11076 |
| chrX:5194551:5195001: | CG5062 | 12183 |
| chr3L:14599451:14599951: | HGTX | 0 |
| chrX:4940101:4940651: | Ptp4E | 974 |
| chr3L:9019101:9019851: | Doc2 | 0 |
| chr3R:21348801:21349851: | CG16791 | 0 |
| chr2R:6220151:6221251: | tomboy40 | 4383 |
| chr3R:29255151:29255701: | stg | 99 |
| chrX:19220151:19221251: | CG7884 | 0 |
| chrX:19604201:19604801: | CG14207 | 398 |
| chr2L:248251:249751: | kis | 1072 |
| chr3R:18585251:18586051: | qin | 4976 |
| chr3L:18640301:18641451: | MYPT-75D | 14343 |
| chr2R:21075301:21076351: | Treh | 1284 |
| chr3R:25515301:25516351: | msi | 0 |
| chrX:10814051:10814651: | Imp | 9626 |
| chr3L:18185351:18187301: | hid | 0 |
| chr3R:27580401:27582051: | wdb | 190 |
| chr2L:3594101:3594551: | odd | 12205 |
| chr2L:22134051:22134551: | CG42748 | 2831 |
| chr3R:26010501:26010951: | E | 0 |
| chrX:2295601:2297801: | Raf | 136 |
| chr2R:15890651:15891351: | CG33463 | 5076 |
| chr2R:21248751:21249301: | ASPP | 1849 |
| chr2R:18413301:18414301: | imd | 0 |
| chrX:9118551:9119251: | AP-1gamma | 9861 |
| chr3R:14065751:14066101: | foxo | 8807 |
| chr3R:31712751:31714201: | ttk | 0 |
| chr3L:10675801:10676401: | simj | 11063 |
| chr2L:12460851:12461851: | CG5418 | 7610 |
| chrX:14218051:14219101: | l | 6595 |
| chrX:22855901:22856451: | fog | 708 |
| chr3L:15535901:15536351: | CrebA | 507 |
| chr3R:23228201:23229001: | cnc | 1491 |
| chr3L:6942751:6943901: | tow | 0 |
| chr3L:21023251:21023901: | skd | 3533 |
| chr2L:9863501:9863901: | nAChRalpha6 | 22349 |
| chr2R:6628101:6628801: | CG33919 | 3525 |
| chr3L:11096201:11097051: | CG33947 | 0 |
| chr3R:21096351:21097401: | RhoGAP93B | 426 |
| chr3R:29457101:29458601: | CG2321 | 30664 |
| chr3R:20291951:20293551: | CG4367 | 11403 |
| chr3R:24272551:24273501: | Orct2 | 0 |
| chr2L:13877851:13878501: | Smg5 | 26160 |

| | | |
|---|---|---|
| chrX:13641501:13642401: | NFAT | 16757 |
| chr3R:10616551:10617601: | hth | 21967 |
| chrX:16607801:16608401: | mbt | 2284 |
| chrX:5052651:5053351: | ovo | 4379 |
| chr3R:20007751:20008351: | Hs6st | 161 |
| chr3L:1366751:1367401: | ru | 3227 |
| chr3L:3757551:3758201: | ppk27 | 2351 |
| chr3L:16051801:16053251: | Mbs | 0 |
| chr3R:23921801:23923001: | sba | 0 |
| chr3R:11371801:11372901: | KP78a | 18671 |
| chr2L:3772751:3773151: | bowl | 1052 |
| chrX:12546101:12548151: | Cpr11B | 4973 |
| chrX:3161851:3163351: | CG18508 | 9789 |
| chr2L:10756901:10757501: | CG17124 | 577 |
| chr3R:30912051:30913001: | Ptx1 | 0 |
| chr3R:10387001:10387501: | cwo | 757 |
| chr2R:7352301:7352901: | Dscam1 | 28998 |
| chr2L:6647101:6647801: | CG31635 | 1313 |
| chrX:18282301:18282851: | upd3 | 5069 |
| chr3L:7866901:7867851: | Pdp1 | 0 |
| chr3R:11312301:11312801: | pros | 15679 |
| chr3R:30532151:30532751: | tmod | 0 |
| chr2R:25052301:25052751: | gsb-n | 0 |
| chrX:7695551:7697701: | CHES-1-like | 11684 |
| chr2R:13592351:13593401: | CG45088 | 0 |
| chr3R:13051251:13052601: | pic | 0 |
| chr2R:8272451:8273201: | CG30371 | 3118 |
| chr2L:246301:247501: | kis | 3322 |
| chr3L:16451901:16452401: | dsx-c73A | 0 |
| chr3R:14696701:14697401: | Meltrin | 0 |
| chrX:3372601:3373451: | Myc | 0 |
| chr3L:11242601:11243351: | Ir68a | 6384 |
| chr2L:19136451:19137251: | l | 1219 |
| chrX:19662751:19663351: | CG14223 | 620 |
| chr3R:23226451:23227201: | cnc | 0 |
| chr2R:13925151:13927201: | shot | 14909 |
| chr3R:9532801:9534101: | by | 0 |
| chrX:15162801:15163401: | CG9095 | 0 |
| chr2L:9581601:9582151: | gcm | 0 |
| chr3L:542851:544801: | Hipk | 0 |
| chr3R:25517851:25518601: | msi | 1734 |
| chr2L:3771201:3772101: | bowl | 0 |
| chrX:8560601:8562101: | Trf4-1 | 280 |
| chrX:20481501:20482101: | CG15461 | 4079 |
| chr3R:10387901:10388501: | cwo | 0 |

| | | |
|---|---|---|
| chr2R:14457901:14458751: | Oaz | 22003 |
| chr3R:21626151:21627051: | InR | 6831 |
| chr3R:29257951:29258551: | CG45544 | 1467 |
| chr3R:21856001:21857001: | glec | 0 |
| chr2L:3836351:3837001: | slp2 | 0 |
| chrX:19353001:19354051: | kek5 | 0 |
| chr2R:13191251:13191951: | seq | 1166 |
| chr3R:17775351:17776951: | TyrR | 12164 |
| chrX:13251451:13251951: | mew | 6539 |
| chr2R:11633051:11633901: | CG9005 | 7040 |
| chrX:15333051:15333801: | HDAC6 | 0 |
| chr3R:10236301:10236851: | CG42795 | 10612 |
| chr2R:9988151:9988851: | TER94 | 308 |
| chr3R:15993151:15993701: | srp | 7000 |
| chr3L:15988201:15988751: | Hip14 | 0 |
| chr3R:28858251:28859701: | Inx3 | 0 |
| chrX:2658301:2658951: | HLH3B | 22772 |
| chr3R:29555851:29556601: | Dr | 0 |
| chrX:3718401:3719801: | Rala | 0 |
| chr2R:12958401:12959301: | CG13321 | 5290 |
| chr3L:20823401:20823901: | Mst77F | 5375 |
| chr3R:6848451:6849201: | Scr | 780 |
| chr3R:29618451:29619351: | CG1907 | 0 |
| chrX:7930101:7931501: | mahe | 0 |
| chr2L:3425851:3426501: | pgant2 | 218 |
| chrX:8650451:8651351: | oc | 0 |
| chr3R:30818651:30819301: | CG15544 | 8518 |
| chr2R:10123651:10125051: | CG46319 | 0 |
| chr3R:11648701:11649251: | Csk | 1455 |
| chrX:2410701:2411251: | CG12496 | 0 |
| chr2R:15895751:15896251: | CG33463 | 176 |
| chrX:10170651:10171251: | CG43902 | 12218 |
| chr3L:16164051:16165101: | CG5151 | 0 |
| chr3L:22835151:22835901: | jim | 0 |
| chr3R:10079101:10079801: | CG5361 | 3729 |
| chr2L:19134701:19135851: | l | 0 |
| chr2R:25114151:25114701: | CG30430 | 14731 |
| chr2R:13189001:13190751: | seq | 0 |
| chr2L:20890151:20890701: | CG14401 | 20298 |
| chr3L:12434301:12434851: | toe | 1634 |
| chr3R:24894401:24895301: | CG11069 | 0 |
| chr2L:11045051:11045551: | CG18666 | 10257 |
| chr3R:21459801:21460501: | lbe | 13265 |
| chr3R:9534501:9535051: | by | 1491 |
| chr2L:1649551:1650351: | chinmo | 907 |

| | | |
|---|---|---|
| chrX:10619601:10620701: | CG43347 | 0 |
| chr2R:23924901:23925301: | CG2812 | 0 |
| chr3L:3249701:3250751: | CG43389 | 1687 |
| chr3L:15924701:15925251: | sff | 0 |
| chrX:8624751:8625301: | Caf1-180 | 18448 |
| chr2R:24369551:24370151: | CG13578 | 7680 |
| chr2R:10219351:10220101: | CG46321 | 7856 |
| chr3R:16224401:16225101: | tara | 550 |
| chr3R:6849901:6850251: | Scr | 0 |
| chr2R:11634951:11635551: | CG9005 | 5390 |
| chr2R:9519001:9520001: | brp | 15237 |
| chr2L:5943401:5945001: | Gpdh | 0 |
| chr3R:30820001:30820501: | CG15544 | 7318 |
| chr3L:9694251:9694901: | CG6767 | 1799 |
| chr2L:14385101:14385651: | ppk | 3930 |
| chr3L:9505101:9506201: | path | 0 |
| chr2L:7245151:7245901: | Ndae1 | 3442 |
| chrX:6700201:6700951: | CG14441 | 694 |
| chr3R:10089151:10089751: | CG5361 | 13779 |
| chr3L:17619201:17619751: | Eip74EF | 0 |
| chr2L:3655251:3658551: | for | 0 |
| chrX:12925301:12926601: | fne | 7825 |
| chrX:9495351:9497801: | mgl | 2328 |
| chrX:2300351:2301801: | Raf | 4886 |
| chr3R:21853501:21854601: | glec | 1665 |
| chr2R:12118851:12119601: | jeb | 0 |
| chrX:3099101:3099601: | CG4116 | 31540 |
| chr2R:13923951:13924601: | shot | 17509 |
| chrX:8090401:8091651: | UbcE2H | 2413 |
| chr2L:8010451:8011051: | Glyat | 0 |
| chr2R:15010501:15011151: | hbs | 245 |
| chr2R:25225651:25227401: | Kr | 0 |
| chr2L:1650701:1651451: | chinmo | 0 |
| chr3R:28860751:28862001: | Inx3 | 1255 |
| chr3R:4360751:4361351: | CG1090 | 869 |
| chr3L:11245801:11246651: | NA-scyl | 5656 |
| chr3R:5700851:5701601: | CG2104 | 7566 |
| chrX:10808501:10809051: | Imp | 15226 |
| chr2L:16525951:16526751: | CG5953 | 6126 |
| chr2R:17475951:17476751: | insb | 0 |
| chr2R:17716001:17717951: | CG6424 | 1395 |
| chr2R:13991051:13992751: | mam | 0 |
| chr2R:20943251:20943851: | Act57B | 170 |
| chr3L:15223651:15223851: | Tollo | 11768 |
| chrX:16068151:16068801: | dpr18 | 4414 |

| | | |
|---|---|---|
| chr3L:9692301:9693801: | CG6767 | 2899 |
| chr2L:5306301:5307001: | CG14024 | 7894 |
| chr3R:31636351:31636801: | kek6 | 14216 |
| chr2R:5141401:5141901: | Atf6 | 2832 |
| chr3R:9436401:9437001: | ps | 18462 |
| chr3L:18016451:18017101: | CG32192 | 23216 |
| chrX:5046401:5048501: | ovo | 0 |
| chrX:16476501:16477001: | eIF4H1 | 27837 |
| chr3L:10681551:10682351: | CG11811 | 8673 |
| chr3R:5086501:5088401: | corto | 0 |
| chr3R:4206601:4207201: | aux | 4582 |
| chr2R:18652601:18653351: | SP2637 | 4871 |
| chrX:7690501:7693251: | CHES-1-like | 16134 |
| chr2R:11231751:11233251: | metro | 0 |
| chr2L:5237001:5238151: | TpnC25D | 20301 |
| chr3R:13011851:13013351: | CtBP | 203 |
| chrX:8091951:8092901: | UbcE2H | 1163 |
| chr3R:8792351:8793001: | Ir85a | 1683 |
| chr2R:19467001:19468701: | mei-W68 | 10832 |
| chr2R:8862251:8862951: | RyR | 1888 |
| chr2R:11637051:11637951: | CG9005 | 2990 |
| chr3R:10622051:10622551: | hth | 17017 |
| chr2L:10457351:10457901: | LManII | 0 |
| chr3L:16807101:16808101: | CG9674 | 0 |
| chr3R:29287251:29287851: | CG14506 | 9329 |
| chr3L:377151:378601: | CG13884 | 10715 |
| chrX:14211151:14212701: | l | 0 |
| chr3L:12892301:12892801: | CG14118 | 0 |
| chr2R:5142301:5142701: | Atf6 | 2032 |
| chr3R:8697301:8698451: | hb | 0 |
| chr2R:15252301:15252851: | scb | 3518 |
| chr2L:12986751:12987651: | Vha68-3 | 7966 |
| chr3R:10087001:10087651: | CG5361 | 11629 |
| chr3L:11247451:11248501: | NA-scyl | 3806 |
| chr2R:21202501:21203201: | Glycogenin | 2019 |
| chr2R:6156101:6157401: | EcR | 11686 |
| chr3R:5716651:5717301: | cas | 0 |
| chr2L:8717751:8718551: | Hnf4 | 8141 |
| chr3R:21621601:21622151: | InR | 2281 |
| chr3L:4071451:4072151: | wit | 0 |
| chr2R:18406651:18407101: | GstE7 | 0 |
| chrX:3722901:3723501: | Tlk | 2895 |
| chr2R:10056351:10057051: | Def | 1776 |
| chrX:2305601:2306951: | Raf | 10136 |
| chr2L:17336251:17336851: | CG45691 | 15540 |

| | | |
|---|---|---|
| chr2L:6088201:6088951: | CG45075 | 2461 |
| chrX:16751151:16751751: | goe | 159 |
| chr3R:23221101:23221751: | cnc | 4960 |
| chr3L:9763251:9764101: | CG8177 | 0 |
| chr2R:9056201:9056651: | CG13743 | 3957 |
| chr3L:14448351:14449101: | CG3919 | 37726 |
| chrX:1975751:1976601: | PIG-K | 32763 |
| chr2R:7418401:7419551: | so | 0 |
| chrX:8213701:8216501: | CG2147 | 24183 |
| chr3R:11391051:11391501: | mRpL40 | 16417 |
| chr3R:17653501:17654351: | CG14322 | 5823 |
| chrX:2303501:2304701: | Raf | 8036 |
| chrX:5918551:5919301: | CG3726 | 1455 |
| chr3R:13683651:13684951: | flfl | 0 |
| chr2R:14715601:14716301: | mspo | 1980 |
| chr2L:18483851:18484951: | CG10283 | 0 |
| chr2R:17718901:17719901: | CG6424 | 0 |
| chr2L:9928901:9929601: | CG42367 | 979 |
| chr2R:22728951:22729801: | jbug | 10658 |
| chr2L:18629001:18629651: | MESR3 | 11746 |
| chr3R:19889801:19890951: | CG4733 | 0 |
| chr3R:18049051:18050401: | htl | 2583 |
| chrX:2049101:2050701: | moody | 0 |
| chr3R:4610351:4610851: | 5-HT2A | 2019 |
| chr2L:7249151:7249951: | Ndae1 | 0 |
| chr3L:10254151:10254751: | Or67c | 9446 |
| chrX:7285351:7285801: | CG2059 | 557 |
| chrX:654251:655951: | Hmt4-20 | 82 |
| chr3R:23220101:23220651: | cnc | 6060 |
| chrX:2664401:2664851: | HLH3B | 16872 |
| chrX:13319451:13320101: | Jafrac1 | 9723 |
| chr2L:21794451:21795801: | CG31612 | 0 |
| chrX:19153201:19155451: | RhoGAP18B | 11631 |
| chr2R:19394901:19395451: | SdhA | 520 |
| chrX:11389551:11390401: | CG15196 | 19546 |
| chr3R:13794351:13795401: | Dip-B | 12064 |
| chr2R:13004901:13005301: | Su | 8401 |
| chr3L:13884751:13885201: | CG13737 | 5758 |
| chr3R:11389601:11390151: | mRpL40 | 17767 |
| chrX:17788951:17790151: | OdsH | 859 |
| chrX:12929851:12930401: | fne | 12375 |
| chr3L:15104001:15105101: | CG45071 | 781 |
| chr3L:21014301:21015101: | skd | 12333 |
| chr3R:10849201:10850051: | CG42394 | 0 |
| chr3R:16934101:16935051: | Abd-B | 37185 |

| | | |
|---|---|---|
| chr2L:13059951:13060751: | CG9932 | 0 |
| chr3R:4609351:4610001: | 5-HT2A | 2869 |
| chr2L:12854401:12855001: | kek1 | 31615 |
| chrX:10240001:10240601: | Ipod | 6922 |
| chrX:5579401:5579951: | Vsx1 | 14304 |
| chr3R:23314001:23314851: | bb8 | 23813 |
| chr3R:23295151:23295951: | bb8 | 4963 |
| chr3R:30600301:30601051: | CG2246 | 438 |
| chr2L:8103951:8104601: | mon2 | 19409 |
| chrX:9638901:9639551: | Ptpmeg2 | 19594 |
| chr2R:24964201:24964551: | CG2765 | 525 |
| chr2R:10213851:10214501: | CG46321 | 13456 |
| chrX:2588651:2589501: | egh | 149 |
| chr3L:17590501:17591201: | Eip74EF | 28124 |
| chr3R:21617951:21619451: | InR | 0 |
| chr3R:31168401:31169451: | stops | 0 |
| chr2R:9230751:9231701: | Cyp4p2 | 5811 |
| chr3R:11655801:11656401: | CG42327 | 0 |
| chrX:19360901:19361651: | kek5 | 7721 |
| chr2R:14713351:14714051: | mspo | 0 |
| chr2L:17750951:17751751: | CadN | 26589 |
| chr3R:15923451:15924001: | CG34276 | 1628 |
| chr3L:3081051:3081651: | CG11486 | 9919 |
| chr3L:12692651:12693901: | mirr | 0 |
| chr2L:18881151:18881751: | tup | 0 |
| chr3R:16933301:16933751: | Abd-B | 38485 |
| chrX:1623101:1623601: | dor | 41451 |
| chr2L:20821401:20822051: | CheB38c | 535 |
| chrX:13126451:13126851: | CG32638 | 1943 |
| chr3R:11302551:11303351: | pros | 25129 |
| chr3L:11251701:11252601: | scyl | 0 |
| chr2R:24712551:24713251: | ST6Gal | 18166 |
| chr2L:377601:378151: | al | 0 |
| chrX:3381901:3382551: | Myc | 8743 |
| chr3R:4807551:4808051: | CG1129 | 3393 |
| chr3R:8027001:8027901: | ImpE3 | 1645 |
| chr3L:18022051:18023251: | CG32192 | 17066 |
| chr2L:3662051:3662801: | Drgx | 0 |
| chr2L:16285651:16287901: | crp | 0 |
| chr3R:29087151:29089151: | alpha-Man-Ib | 23671 |
| chrX:18832251:18832801: | Pvf1 | 0 |
| chr2L:17752201:17753151: | CadN | 25189 |
| chrX:13322251:13322751: | Jafrac1 | 7073 |
| chrX:10801051:10802601: | sesB | 14094 |
| chr2L:66351:67601: | dbr | 0 |

| | | |
|---|---|---|
| chr3L:17611801:17612551: | Eip74EF | 6774 |
| chr2R:5992501:5993101: | Src42A | 10658 |
| chr2R:5726701:5727401: | ap | 124 |
| chr2L:16532651:16533751: | CG5953 | 0 |
| chr3R:4805901:4807251: | CG1129 | 4193 |
| chrX:10546501:10547201: | CG15296 | 19390 |
| chr2R:12886651:12887151: | vg | 2020 |
| chr3L:21010751:21012151: | skd | 15283 |
| chr3R:21616101:21617101: | InR | 2220 |
| chr3L:3827901:3828701: | Rdh | 2453 |
| chr3R:17665801:17667001: | CG7523 | 3894 |
| chr2R:10210701:10211951: | CG34221 | 10991 |
| chr3L:5726351:5726851: | CG44521 | 13021 |
| chr2R:24793201:24793751: | Reg-5 | 0 |
| chr3R:16051101:16051751: | msps | 301 |
| chr3R:15919651:15921701: | Fbxl7 | 0 |
| chr3R:23173401:23174251: | CG46310 | 0 |
| chrX:10628401:10629351: | X11Lbeta | 665 |
| chr2R:12111051:12111551: | jeb | 7789 |
| chr2L:12055651:12056551: | CG34164 | 0 |
| chr3R:9543451:9543951: | mura | 7855 |
| chr2L:3580401:3581401: | sob | 0 |
| chr2R:19178651:19179501: | hppy | 0 |
| chr3R:14853651:14854401: | CG3610 | 8609 |
| chrX:8640801:8641251: | oc | 9430 |
| chr2R:22270351:22271251: | CG5819 | 22149 |
| chr2R:11493801:11494351: | inv | 19337 |
| chr3R:17118851:17119551: | CG34278 | 914 |
| chr2L:16534001:16534601: | CG5953 | 1125 |
| chrX:9515351:9515951: | CG34028 | 3828 |
| chr2L:17754051:17754601: | CadN | 23739 |
| chr3L:7854301:7855901: | Pdp1 | 11468 |
| chr3R:28585251:28585851: | fkh | 0 |
| chrX:12699201:12699951: | CG15725 | 5903 |
| chr3R:11254251:11254951: | CG5214 | 9950 |
| chrX:6709251:6709701: | CG14441 | 9744 |
| chrX:17124801:17125701: | CG5004 | 17220 |
| chrX:9599301:9600301: | Gga | 729 |
| chrX:12720051:12720551: | CG15725 | 14198 |
| chr3L:21007201:21010401: | CG12984 | 15529 |
| chr2L:16284151:16285351: | crp | 2379 |
| chr3L:18064601:18065301: | Eip75B | 0 |
| chrX:15834351:15835251: | Chc | 239 |
| chr2R:12884301:12885201: | vg | 0 |
| chr3R:13874851:13875601: | E5 | 0 |

| | | |
|---|---|---|
| chr2L:16534851:16536301: | CG5953 | 1975 |
| chr2L:19764451:19765101: | fbp | 316 |
| chrX:8100001:8100701: | CG2258 | 1394 |
| chr3L:22910051:22910751: | CG12768 | 0 |
| chr3L:20348201:20349901: | eRF1 | 0 |
| chr3R:9359351:9359851: | CG18473 | 3454 |
| chr3R:23175151:23175701: | CG46310 | 1279 |
| chrX:8548901:8549801: | IntS4 | 8980 |
| chr3R:24699151:24699801: | CG46316 | 12979 |
| chrX:15345201:15345851: | CG6227 | 33 |
| chr2R:14814251:14814701: | CG10202 | 5286 |
| chrX:8295351:8296151: | CG15343 | 21546 |
| chr3L:15098551:15099601: | CTPsyn | 638 |
| chr2R:6235401:6236501: | Pld | 5503 |
| chrX:5560451:5561151: | Vsx2 | 269 |
| chrX:395501:396351: | sc | 0 |
| chrX:9190551:9191151: | CG12115 | 0 |
| chr2R:17638951:17639401: | CG10936 | 0 |
| chr2R:14000601:14001401: | mam | 9399 |
| chr2L:19397851:19399351: | CG17544 | 0 |
| chrX:13130651:13131801: | MFS10 | 1634 |
| chr3R:29818601:29819251: | neo | 0 |
| chr3R:10638651:10639251: | hth | 317 |
| chrX:3175751:3176651: | dnc | 0 |
| chrX:17345801:17346501: | B-H2 | 31221 |
| chr3L:4363251:4364151: | Cip4 | 0 |
| chr2L:9567401:9569101: | CG3838 | 0 |
| chrX:17870951:17871551: | CG6769 | 9950 |
| chr2L:8478301:8478951: | CheA29a | 953 |
| chr3R:4433351:4433951: | eIF3a | 0 |
| chr3R:16403401:16403951: | ss | 0 |
| chr2R:18846051:18846651: | CG15099 | 0 |
| chrX:19718301:19718851: | CG12531 | 72 |
| chr3R:18742501:18743801: | Mekk1 | 0 |
| chr3R:4802301:4803751: | CG1129 | 7693 |
| chr3R:14598051:14598751: | CG14853 | 10891 |
| chr2R:6612851:6613651: | CG33919 | 10926 |
| chr3L:18063151:18063651: | Eip75B | 1045 |
| chr3R:22626451:22627401: | mRpL45 | 7960 |
| chr3L:7781501:7782201: | Clk | 5899 |
| chr2R:20932951:20933401: | Act57B | 10620 |
| chr3R:21111601:21113301: | Rab11 | 194 |
| chrX:5561601:5562151: | Vsx2 | 1419 |
| chr3L:18792651:18793301: | ftz-f1 | 8008 |
| chr2L:21141701:21142251: | Dap160 | 96 |

| | | |
|---|---|---|
| chrX:13237601:13238151: | CG32639 | 1036 |
| chr2L:19361901:19363451: | dnt | 0 |
| chr3R:20782401:20783051: | Takl1 | 9801 |
| chr3R:13877001:13877501: | E5 | 2086 |
| chr3L:22822351:22822951: | jim | 12243 |
| chrX:13657101:13657451: | Lig4 | 10360 |
| chrX:20352301:20352851: | CG1504 | 21930 |
| chrX:10247301:10247851: | Ipod | 14222 |
| chrX:2387451:2388551: | PsGEF | 0 |
| chr2R:15816701:15817501: | Zasp52 | 0 |
| chr2R:12996001:12997501: | Su | 0 |
| chr3L:4061901:4062451: | dib | 0 |
| chr2L:21051901:21052451: | CG42238 | 0 |
| chr2L:5107551:5108751: | Msp300 | 6675 |
| chr2L:1662601:1663601: | chinmo | 11344 |
| chrX:6231701:6232301: | CG14445 | 5014 |
| chr3L:6217901:6218951: | LanA | 0 |
| chrX:12937901:12938751: | hec | 18711 |
| chr3R:21446001:21447001: | lbe | 0 |
| chr2R:18583051:18583801: | GEFmeso | 27976 |
| chr3R:27635951:27636901: | Mtl | 0 |
| chr2L:18795901:18796851: | ham | 2996 |
| chr3L:1919551:1921551: | Dbx | 0 |
| chrX:9305251:9306451: | c12 | 12772 |
| chr3R:4910751:4911401: | Cdep | 200 |
| chr3R:5413601:5414651: | CG14669 | 0 |
| chr2L:12655601:12656301: | pdm2 | 864 |
| chrX:16148851:16149701: | disco-r | 0 |
| chr2R:14798851:14799951: | kn | 7110 |
| chr3R:5263951:5265251: | Tim17a2 | 1260 |
| chrX:13784051:13784601: | inaE | 338 |
| chr2L:9205001:9205901: | tai | 38224 |
| chrX:3294651:3295901: | CG10793 | 20501 |
| chr2R:6609951:6610801: | CG15233 | 8974 |
| chr3R:18739901:18740701: | Mekk1 | 2893 |
| chr3L:18029301:18030001: | CG32192 | 10316 |
| chr2L:16279851:16280601: | crp | 7129 |
| chr3L:6789501:6790351: | vvl | 0 |
| chr2L:12974651:12975451: | Vha68-2 | 330 |
| chr3R:20594551:20595201: | bon | 1413 |
| chrX:1963701:1965251: | Hr4 | 21762 |
| chrX:20444851:20445351: | CG42578 | 1020 |
| chr2R:15424851:15425401: | CG30472 | 713 |
| chr3R:4599501:4600101: | 5-HT2A | 12769 |
| chr3R:18244351:18245051: | 14-3-3epsilon | 1821 |

| | | |
|---|---|---|
| chr3R:27149951:27151001: | CG6051 | 68 |
| chr3R:10170101:10170901: | Best1 | 0 |
| chrX:13234101:13234851: | CG32639 | 1715 |
| chrX:16150301:16151151: | disco-r | 1325 |
| chr3R:9550401:9551901: | mura | 0 |
| chr3R:24565401:24566151: | REPTOR | 16713 |
| chr2R:17813801:17814351: | grh | 12670 |
| chr2R:11650751:11651701: | CG9003 | 0 |
| chr3R:9352201:9354151: | CG43675 | 1344 |
| chr2L:368701:369151: | al | 8961 |
| chr3L:880851:881401: | CG13907 | 612 |
| chrX:1275901:1276851: | Atf3 | 618 |
| chr3L:9400951:9402601: | eIF4E1 | 0 |
| chrX:4411201:4411851: | bi | 1005 |
| chr3L:7381201:7381851: | smid | 0 |
| chr3L:9678001:9678701: | fry | 402 |
| chr2R:14927901:14928601: | Kank | 0 |
| chr3L:12467851:12468551: | eyg | 0 |
| chr3R:21152401:21153501: | cDIP | 7591 |
| chr2R:14006501:14007551: | mam | 15299 |
| chr3L:9422701:9423451: | aay | 0 |
| chr3R:4797501:4798351: | CG17387 | 12780 |
| chr3L:20441651:20442201: | CG5910 | 218 |
| chr3L:18787701:18788301: | ftz-f1 | 13008 |
| chr3L:881701:882451: | CG13907 | 0 |
| chr2L:6667651:6668251: | Galt | 0 |
| chrX:3292601:3293151: | CG10793 | 23251 |
| chr3R:7037001:7038101: | Sodh-1 | 14234 |
| chr3R:5072401:5073001: | corto | 14071 |
| chrX:15352301:15352751: | CG6227 | 7133 |
| chr3R:23722551:23724251: | CG10365 | 0 |
| chr2R:23697601:23698251: | mRpL43 | 0 |
| chrX:19682651:19683851: | Ubqn | 0 |
| chr2L:12482651:12484701: | t-cup | 12632 |
| chr3R:17566601:17567301: | Hmx | 7672 |
| chr3L:20442701:20443451: | CG5910 | 283 |
| chr2R:22136551:22137251: | a | 0 |
| chr3R:18121301:18122251: | CG14316 | 29869 |
| chr3L:7947751:7948401: | exex | 0 |
| chr3R:21604651:21607201: | InR | 12120 |
| chr3L:12081601:12082151: | Sema5c | 0 |
| chr2R:14007951:14008801: | mam | 16749 |
| chrX:17767951:17768851: | unc-4 | 0 |
| chr2R:14908051:14908651: | Cyp317a1 | 9740 |
| chr3R:18063151:18063701: | htl | 10168 |

| | | |
|---|---|---|
| chr2L:21851451:21851801: | tsh | 22859 |
| chr2L:11221151:11221701: | ab | 10474 |
| chrX:17878301:17879101: | CG12985 | 4617 |
| chr3L:11118401:11119201: | FoxK | 1582 |
| chr2R:7885751:7886501: | Dgk | 0 |
| chr3R:5518601:5519151: | Itp-r83A | 1485 |
| chr2R:6605701:6606351: | CG15233 | 4724 |
| chr3R:8190551:8191351: | grn | 9108 |
| chr3R:10535601:10536301: | Cyp12e1 | 100813 |
| chr2L:2798701:2799401: | Syt1 | 578 |
| chrX:10783701:10786401: | Ant2 | 506 |
| chrX:5363901:5364601: | SK | 23666 |
| chr3L:9675451:9676051: | fry | 3052 |
| chrX:13663951:13664451: | Lig4 | 3360 |
| chr2R:10485201:10485901: | pre-lola-G | 0 |
| chr3L:21879151:21881051: | mub | 34364 |
| chr3L:15090101:15090751: | sstn | 3968 |
| chr3R:19875001:19875651: | CG4562 | 27 |
| chr2R:11144901:11145501: | CG13231 | 788 |
| chr3R:25094651:25095051: | fd96Cb | 0 |
| chr2L:12044501:12045301: | CG6785 | 0 |
| chr2R:12979701:12981501: | Psc | 0 |
| chr2R:19228801:19230201: | cora | 0 |
| chrX:9624351:9625201: | CG42797 | 15406 |
| chr3L:2824601:2825151: | pgant6 | 42347 |
| chr2L:13549951:13551201: | kuz | 0 |
| chr2R:9875001:9875501: | CG46338 | 0 |
| chr2R:15600051:15600851: | CG42524 | 13003 |
| chrX:6224301:6224851: | sqh | 289 |
| chr3R:31688951:31689751: | CG11550 | 9072 |
| chr2L:220401:224651: | CG13693 | 7836 |
| chrX:3740451:3741301: | Tlk | 20445 |
| chr3R:16918801:16919501: | Abd-B | 52735 |
| chr2L:16545501:16546151: | CG42389 | 486 |
| chr2L:8735551:8736151: | raw | 4509 |
| chr2L:7827801:7829401: | mts | 152 |
| chr3R:24588701:24589401: | mld | 3049 |
| chr2R:7530651:7531301: | Drat | 1680 |
| chrX:2083451:2084251: | Unc-76 | 2842 |
| chr2R:8018701:8019201: | CG14762 | 4947 |
| chr3L:4160801:4161551: | nab | 0 |
| chr2L:8030901:8032251: | Cka | 181 |
| chr2L:21150951:21151601: | del | 2406 |
| chrX:19545951:19546951: | CG32532 | 531 |
| chr3L:2823301:2823951: | pgant6 | 41047 |

| | | |
|---|---|---|
| chr2L:8088151:8088851: | Bsg | 4732 |
| chr3L:18398151:18398851: | rpr | 0 |
| chrX:15356201:15357001: | Pp1-13C | 4060 |
| chrX:9623351:9623751: | CG42797 | 14406 |
| chr2R:14481251:14482251: | PRAS40 | 65 |
| chr3L:12056251:12056751: | rols | 8108 |
| chr2R:21101351:21102351: | ktub | 0 |
| chr3R:22732801:22733551: | Takl2 | 300 |
| chrX:1941551:1942201: | Hr4 | 0 |
| chr2L:11447001:11448301: | salm | 1388 |
| chr3R:24571701:24573101: | REPTOR | 9763 |
| chr2R:22241751:22244301: | dve | 0 |
| chrX:3741801:3743051: | Tlk | 21795 |
| chr2R:11397101:11398151: | sprt | 0 |
| chr3R:8187651:8188151: | grn | 6208 |
| chrX:16216401:16218051: | disco | 0 |
| chr3R:5612201:5612901: | Rga | 211 |
| chr3R:29377101:29377651: | Ptp99A | 0 |
| chrX:16497151:16497751: | Cnx14D | 27664 |
| chr3R:30617251:30617751: | CG44954 | 4155 |
| chrX:18587201:18587701: | CG15040 | 29888 |
| chr3R:16502451:16502951: | CG42342 | 0 |
| chr2R:10477601:10478751: | whd | 0 |
| chr2R:11657601:11658401: | CG13198 | 2704 |
| chr3R:11156701:11157251: | Ugt86Da | 0 |
| chr3R:13901451:13902101: | ems | 0 |
| chrX:21261651:21262101: | CG1722 | 10504 |
| chr3R:24586351:24587051: | mld | 699 |
| chr2L:16548001:16548701: | CG5968 | 1258 |
| chrX:1943101:1943801: | Hr4 | 1162 |
| chr2L:11445401:11446801: | salm | 0 |
| chrX:15633251:15633901: | sog | 0 |
| chr2R:22256201:22256701: | dve | 12239 |
| chrX:19548401:19548901: | CG32532 | 920 |
| chr3L:2820901:2821551: | pgant6 | 38647 |
| chr3R:23201001:23201451: | fzo | 2583 |
| chr2R:12984401:12986301: | Psc | 3408 |
| chr3L:12913751:12914351: | CG32115 | 106 |
| chr3L:11123801:11124501: | NaPi-III | 454 |
| chr3L:11010001:11011101: | klu | 814 |
| chr3R:21433951:21434901: | lbl | 0 |
| chr3R:11269051:11269501: | CG5214 | 24750 |
| chrX:3744301:3745201: | Tlk | 24295 |
| chr2L:14409301:14410101: | elB | 0 |
| chr3L:15085001:15085501: | Or71a | 6555 |

| | | |
|---|---|---|
| chr3L:11764901:11765501: | CG6024 | 681 |
| chr2R:15734651:15735801: | CG30089 | 7271 |
| chr2R:13619651:13620701: | CG17716 | 2343 |
| chr2L:8739801:8740751: | raw | 0 |
| chrX:6804151:6805101: | pod1 | 468 |
| chrX:19689901:19690651: | CG14227 | 954 |
| chr2L:16268901:16270051: | CG4935 | 6761 |
| chr2R:22859951:22860751: | inaD | 1158 |
| chr2L:8084601:8085001: | Bsg | 1182 |
| chr2R:24813401:24815001: | Dll | 0 |
| chr3R:21829201:21829901: | Gr93a | 2602 |
| chr3L:13629201:13629901: | CG10710 | 109597 |
| chr3L:11008551:11009801: | klu | 0 |
| chr3R:18849101:18849701: | CG18208 | 0 |
| chr2R:17804001:17804651: | grh | 2870 |
| chr2L:5658301:5659551: | DIP-theta | 0 |
| chr2R:21899051:21899501: | PpN58A | 16519 |
| chrX:17885501:17887301: | mnb | 0 |
| chrX:8533801:8534451: | PIP82 | 4104 |
| chrX:15028951:15029401: | be | 4516 |
| chr2R:6131701:6134351: | CG14589 | 2952 |
| chrX:3190651:3191451: | dnc | 14212 |
| chrX:18370701:18371851: | CrebB | 4345 |
| chr3R:17558451:17559251: | Hmx | 0 |
| chrX:18930801:18931401: | CG43759 | 467 |
| chr2L:8083251:8084151: | Bsg | 0 |
| chr3R:21598401:21599151: | InR | 20170 |
| chr2L:9175901:9177251: | tai | 9124 |
| chr2R:10596151:10596651: | CG11883 | 21011 |
| chr2L:18782501:18783751: | ham | 9155 |
| chr2L:16267801:16268601: | CG4935 | 5661 |
| chr2L:7701401:7701951: | Tep2 | 0 |
| chr2R:19373051:19373501: | CG9416 | 201 |
| chr3R:25137451:25138351: | danr | 0 |
| chr2R:18232001:18233201: | fj | 0 |
| chr2R:22867601:22868101: | fd59A | 58 |
| chr3R:17557301:17558101: | Hmx | 829 |
| chr3L:18946901:18947451: | CG32204 | 864 |
| chr2R:24342551:24343051: | bs | 0 |
| chr2R:13897451:13898051: | CG16935 | 35450 |
| chrX:10262001:10262551: | Hk | 934 |
| chr2R:13501751:13502951: | drk | 0 |
| chr3R:9340651:9342901: | CG45050 | 436 |
| chrX:20982451:20982901: | CG11666 | 1785 |
| chr2R:17800851:17802801: | grh | 0 |

| | | |
|---|---|---|
| chr3L:4167201:4167851: | mas | 0 |
| chr2L:4221551:4222751: | ft | 0 |
| chrX:16332351:16333901: | Dsp1 | 0 |
| chr2R:11662401:11663251: | Tret1-1 | 2517 |
| chr3L:2182051:2182551: | CG13810 | 3379 |
| chr2R:13622501:13623351: | CG17716 | 0 |
| chr3R:10826551:10827301: | Cad86C | 0 |
| chr3R:6402701:6403351: | Dmtn | 2544 |
| chr3R:31662751:31663251: | CG1815 | 6214 |
| chrX:1947751:1948651: | Hr4 | 5812 |
| chr3R:8181251:8182201: | grn | 0 |
| chrX:1951501:1952151: | Hr4 | 9562 |
| chr3R:19096501:19097051: | CG6040 | 0 |
| chr2R:20915901:20917051: | Rx | 0 |
| chr3R:11281401:11282051: | CG5214 | 37100 |
| chr3R:17556151:17556751: | Hmx | 2179 |
| chr2R:18229501:18231701: | fj | 1060 |
| chr3L:3963401:3964651: | CG12605 | 120 |
| chr3R:21948501:21949301: | Eip93F | 0 |
| chr3R:30880951:30881451: | CG12071 | 0 |
| chrX:3748601:3749401: | Tlk | 28595 |
| chrX:20693601:20694601: | run | 2932 |
| chrX:4423651:4425501: | bi | 10796 |
| chr3L:12328701:12329251: | Lmx1a | 0 |
| chr2L:478751:481251: | cbt | 0 |
| chr3R:12278801:12279551: | Sfp87B | 5676 |
| chr2R:17335651:17336151: | CG10939 | 58442 |
| chr3R:30099101:30099701: | CG45072 | 23562 |
| chrX:18374451:18375001: | por | 4853 |
| chrX:12395001:12395501: | CG32651 | 12976 |
| chrX:13219951:13220401: | REG | 7627 |
| chr2L:6114951:6115551: | CG31642 | 2961 |
| chr3L:15079101:15079951: | Or71a | 655 |
| chr3R:6405051:6405901: | Dmtn | 0 |
| chr3L:12580051:12580751: | ara | 0 |
| chr2R:18355101:18358051: | sbb | 0 |
| chr2R:20913551:20914801: | Rx | 1952 |
| chr3L:768951:769801: | hng3 | 13463 |
| chr2R:11665251:11665901: | Tret1-1 | 0 |
| chrX:15023601:15024651: | be | 0 |
| chr3R:16025501:16026351: | pnr | 0 |
| chr3R:12280501:12281351: | Sfp87B | 7376 |
| chr2L:10353751:10354301: | CG34367 | 0 |
| chr2L:17565701:17566251: | CG7094 | 9937 |
| chr2R:15613151:15614051: | CG42524 | 0 |

| | | |
|---|---|---|
| chrX:15811451:15813801: | CG8509 | 1897 |
| chrX:4596251:4597051: | peb | 20513 |
| chr2R:11666451:11667051: | Tret1-1 | 684 |
| chrX:4426551:4427451: | bi | 13696 |
| chr3R:30851701:30852551: | tll | 0 |
| chr3R:6406701:6408101: | gpp | 0 |
| chr3R:14346701:14347451: | HEATR2 | 14335 |
| chr3L:17486751:17487901: | Oatp74D | 0 |
| chr3R:18722451:18723001: | CG34283 | 0 |
| chr3R:21557001:21558301: | slou | 0 |
| chr2R:18377301:18377801: | CG43202 | 4043 |
| chr2R:25257251:25257751: | CG9380 | 1934 |
| chr3R:16702401:16704101: | Ubx | 30325 |
| chr3R:13147501:13148101: | 2mit | 7586 |
| chr3L:22132551:22133251: | olf413 | 2839 |
| chr3R:5282651:5283601: | CG31538 | 4817 |
| chr3L:21202701:21203251: | CG33056 | 0 |
| chrX:6601301:6602251: | pigs | 1240 |
| chr3L:18771351:18772201: | Atg3 | 22665 |
| chr2L:8587801:8588251: | Sema1a | 45655 |
| chr2R:23713051:23713951: | Sesn | 0 |
| chr2R:8798051:8798651: | sns | 0 |
| chr3L:14131351:14131901: | Sox21b | 28 |
| chr2R:12261101:12261851: | Cam | 1704 |
| chr2L:12365751:12366601: | CG17010 | 9311 |
| chr3R:19860801:19861501: | Ire1 | 0 |
| chr2R:10025101:10026451: | Pka-R2 | 0 |
| chr2L:15425701:15426451: | wor | 122 |
| chr3L:4624351:4626451: | Src64B | 0 |
| chr2R:23579951:23581351: | apt | 15061 |
| chr3L:445701:446301: | CG13891 | 11796 |
| chrX:9743801:9744701: | Sp1 | 14154 |
| chr2R:6258801:6259401: | Pld | 16798 |
| chr3R:21558851:21559351: | slou | 1670 |
| chr3R:8413851:8414351: | Obp85a | 6246 |
| chr3L:22134001:22136501: | olf413 | 0 |
| chrX:13464201:13464801: | CG34324 | 6539 |
| chrX:17484301:17484851: | CG8568 | 31759 |
| chr2L:3095001:3095551: | PpD6 | 14121 |
| chr2R:14024451:14025101: | CG18371 | 3519 |
| chrX:7444951:7445501: | CG11368 | 6708 |
| chr3L:7804501:7805851: | CG32369 | 3974 |
| chr2R:10159601:10160351: | mlt | 671 |
| chr3R:21589901:21590351: | CG15498 | 20753 |
| chr3R:14574751:14575301: | CG7987 | 586 |

| | | |
|---|---|---|
| chrX:6599301:6600301: | pigs | 0 |
| chr3R:20254601:20255101: | cic | 1832 |
| chr3L:6903751:6905001: | Prat2 | 9875 |
| chr3R:31810001:31810501: | CG11576 | 0 |
| chr3L:3149051:3149801: | BtbVII | 406 |
| chr3R:6410201:6411201: | gpp | 3347 |
| chr3R:19225251:19226701: | vib | 0 |
| chr3L:19635251:19636051: | wnd | 0 |
| chr3R:30386601:30389701: | Fer1HCH | 0 |
| chr2R:16270301:16270801: | Khc | 1170 |
| chr2R:12259051:12259651: | Cam | 0 |
| chrX:13805351:13806301: | CG11095 | 5383 |
| chrX:840451:841401: | CG14635 | 3422 |
| chr3R:8895551:8896201: | pyd | 35695 |
| chr3R:17943501:17944401: | cpo | 23670 |
| chr3R:20252651:20254401: | cic | 0 |
| chrX:12085601:12086401: | CG15734 | 624 |
| chr3R:26797751:26799301: | Tl | 0 |
| chr3R:9865851:9866601: | Teh1 | 1519 |
| chr2L:523051:524001: | lwr | 18579 |
| chr3L:15021001:15021451: | CG32148 | 2041 |
| chr2R:20896101:20896651: | otp | 6763 |
| chr3R:21561151:21562151: | slou | 3970 |
| chr3L:2808051:2808751: | pgant6 | 25797 |
| chr3R:11886901:11888551: | mthl5 | 0 |
| chr2L:16422401:16423501: | Idgf1 | 21777 |
| chr2L:348001:348401: | Ent1 | 9473 |
| chr2L:6786601:6788151: | nrv1 | 0 |
| chrX:686701:687851: | sdk | 0 |
| chr2L:486801:487351: | cbt | 7114 |
| chrX:4617001:4617851: | peb | 0 |
| chr2R:20907151:20907751: | CG9235 | 1953 |
| chr2L:522051:522751: | lwr | 19829 |
| chr3L:8512251:8513101: | CG6765 | 0 |
| chr3L:9546751:9547401: | CG42673 | 0 |
| chr2L:20561351:20562401: | CG44270 | 30286 |
| chrX:18946501:18947301: | CG43759 | 16167 |
| chr3R:23962701:23963351: | CG31140 | 0 |
| chr3R:19241651:19242151: | unc79 | 3979 |
| chr3L:270701:272151: | miple2 | 0 |
| chr3L:22787851:22788451: | CG14448 | 0 |
| chr3R:30337951:30338751: | CG34300 | 31312 |
| chr2L:21827951:21829151: | tsh | 0 |
| chr3R:5823151:5823701: | CG2023 | 831 |
| chr3R:14108151:14108951: | CG12402 | 0 |

| | | |
|---|---|---|
| chr2R:14498201:14498651: | Cpsf160 | 4837 |
| chr3L:13993251:13993851: | CG17364 | 0 |
| chrX:19393351:19393851: | kek5 | 40171 |
| chr3R:9573401:9574051: | CG9399 | 160 |
| chr3R:25328501:25329551: | CHKov1 | 0 |
| chr2R:14139651:14141351: | Prosap | 0 |
| chr3L:7808651:7810201: | CG32369 | 0 |
| chr3R:31310551:31311251: | CG34347 | 0 |
| chr3L:18765601:18766251: | Atg3 | 16915 |
| chr3R:12448801:12450051: | CG18549 | 0 |
| chr3L:6813851:6814301: | vvl | 23694 |
| chr2R:6120251:6120951: | CG14589 | 16352 |
| chr2L:12504101:12505651: | t-cup | 6769 |
| chr3L:10289201:10289951: | Or67d | 15998 |
| chrX:9749551:9750101: | Sp1 | 19904 |
| chr3R:8899551:8900151: | pyd | 31745 |
| chr2L:7809451:7810301: | cdc14 | 402 |
| chr3R:16898501:16900201: | abd-A | 68453 |
| chr3L:18539651:18540151: | CG32198 | 12676 |
| chr2R:11529301:11530051: | en | 1092 |
| chr3R:23965001:23965701: | CG31140 | 1822 |
| chr2L:490101:491201: | cbt | 10414 |
| chrX:12090251:12090951: | CG15734 | 3227 |
| chr3L:5840301:5840851: | vn | 4813 |
| chrX:15803851:15804651: | sd | 0 |
| chr3R:18480551:18481101: | fru | 64485 |
| chr3R:23818551:23819351: | eIF4G2 | 0 |
| chr3L:8913651:8914251: | mfr | 981 |
| chrX:5008551:5009051: | CG12680 | 11111 |
| chr2R:23572901:23573951: | apt | 8011 |
| chr2R:14031151:14031751: | CG18371 | 2532 |
| chr2R:11527601:11528751: | en | 0 |
| chr2L:517901:518701: | lwr | 23879 |
| chr3R:9157751:9158651: | CG11997 | 18866 |
| chr2R:19356351:19357401: | Rgk1 | 0 |
| chrX:8517951:8518551: | Nrg | 579 |
| chr2R:18611501:18612201: | GEFmeso | 0 |
| chr2R:12252501:12253251: | CG13168 | 741 |
| chr2L:7121851:7122401: | Pvf3 | 35294 |
| chr2L:6791901:6792501: | nrv1 | 5041 |
| chr2L:9781951:9782601: | IP3K1 | 0 |
| chr3R:9872151:9872901: | Glut4EF | 1780 |
| chrX:16562151:16562751: | Pp2B-14D | 2136 |
| chr2L:15332301:15334201: | esg | 0 |
| chr2L:12507401:12508301: | t-cup | 10069 |

| | | |
|---|---|---|
| chr3R:7162401:7163151: | sas | 0 |
| chr3R:21581501:21582451: | CG15498 | 12353 |
| chrX:8757601:8758301: | CG1789 | 37082 |
| chr3L:14272701:14273351: | NA-fz | 992 |
| chr2L:16251501:16252201: | CG46309 | 0 |
| chr3L:5975701:5977001: | CG10479 | 0 |
| chr2L:516151:517001: | lwr | 25579 |
| chr2L:5460901:5461801: | mid | 0 |
| chrX:4611201:4611801: | peb | 5763 |
| chr3R:9873301:9875201: | Glut4EF | 0 |
| chr3L:14550701:14551651: | Gbs-70E | 0 |
| chrX:276001:276401: | CG3777 | 3302 |
| chr2L:6793601:6794951: | nrv2 | 3911 |
| chrX:3775701:3781351: | HIP-R | 8618 |
| chr3L:14273751:14274701: | fz | 0 |
| chr3L:18233951:18234851: | CheA75a | 17237 |
| chr2R:6269651:6270201: | Pld | 27648 |
| chr2R:16514751:16515301: | Sema2a | 14574 |
| chrX:1398851:1400101: | CG11448 | 5798 |
| chrX:16559201:16560051: | Pp2B-14D | 0 |
| chr3L:22945051:22945851: | Mes2 | 0 |
| chr3R:16715501:16716301: | Ubx | 18125 |
| chr3R:21807851:21809101: | CASK | 13573 |
| chrX:3545951:3546651: | ppk8 | 11161 |
| chr2L:11581151:11582201: | kek2 | 0 |
| chr3L:19833151:19833651: | trc | 653 |
| chr3L:10567801:10568301: | CG6527 | 8720 |
| chrX:15007701:15008251: | be | 16185 |
| chr3R:29782251:29782801: | fig | 1800 |
| chr3L:18756951:18757801: | Atg3 | 8265 |
| chr2R:20327251:20328101: | CG8929 | 0 |
| chr3R:21297151:21297651: | SIFaR | 3154 |
| chr3R:21295601:21296851: | SIFaR | 1604 |
| chr3L:3983151:3984151: | scrt | 0 |
| chr2R:10618201:10618851: | CG11883 | 540 |
| chr3L:22948251:22948901: | Mes2 | 2595 |
| chr2L:6536051:6536601: | eya | 10375 |
| chr3R:16718451:16719051: | Ubx | 15375 |
| chrX:8510751:8511201: | Nrg | 6172 |
| chrX:10120651:10121201: | CG12645 | 32411 |
| chr3L:7978851:7980401: | nmo | 0 |
| chrX:2545251:2546051: | trol | 0 |
| chr2L:510301:511051: | cbt | 30614 |
| chr3R:6424001:6424601: | gpp | 17147 |
| chr3L:20310251:20310901: | polo | 598 |

| | | |
|---|---|---|
| chr3R:4565251:4565801: | CG34357 | 7185 |
| chr2L:5799301:5800601: | CG9171 | 0 |
| chr2L:5805001:5805651: | CG11034 | 0 |
| chr2R:15135101:15135601: | Rpb12 | 9918 |
| chr2R:6574651:6575601: | CG15233 | 25377 |
| chrX:6189501:6189951: | Ca-alpha1T | 2532 |
| chr3L:17149601:17150401: | Rbp6 | 87811 |
| chr3R:16719601:16720501: | Ubx | 13925 |
| chr3L:11294601:11295251: | CG6175 | 668 |
| chr3L:748201:749701: | emc | 0 |
| chr3R:27194001:27194601: | Ser | 0 |
| chrX:5965501:5966101: | Grip | 178 |
| chrX:16525751:16526601: | Cnx14D | 337 |
| chr2R:14040851:14041651: | CG18371 | 12232 |
| chr2L:6833451:6834101: | sens-2 | 11518 |
| chr2L:12617801:12619001: | Ref2 | 27355 |
| chrX:3773051:3773901: | HIP-R | 16068 |
| chr4:613001:613651: | Eph | 2318 |
| chr3L:6256451:6257301: | Best2 | 0 |
| chr3L:5276501:5278501: | shep | 0 |
| chr2R:11366501:11367801: | CG13204 | 0 |
| chr2R:8326701:8327651: | pdm3 | 0 |
| chr2L:4197101:4197851: | CG3714 | 220 |
| chr2R:6037451:6037851: | SCAP | 1064 |
| chr2R:18061801:18062351: | CG34386 | 64 |
| chr2R:21871501:21872151: | Fili | 0 |
| chrX:14923201:14923801: | CG9503 | 0 |
| chr2L:11358301:11359101: | salr | 0 |
| chr3R:21289001:21289651: | SIFaR | 4347 |
| chr3L:7434951:7435751: | Rac2 | 0 |
| chrX:11713501:11715351: | Amun | 214 |
| chr3R:29774051:29775101: | fig | 5351 |
| chrX:14748901:14750051: | NetB | 0 |
| chr3R:14369951:14370601: | NK7 | 2350 |
| chr2R:7699151:7699751: | CG30496 | 0 |
| chr3L:17740251:17740851: | CG7460 | 0 |
| chr2R:8985401:8986001: | CG8213 | 0 |
| chr2L:12520601:12521751: | t-cup | 23269 |
| chrX:18023451:18024251: | CG6867 | 0 |
| chrX:1387901:1389001: | CG11448 | 4053 |
| chr3L:21916001:21916801: | SLC22A | 17903 |
| chr2L:9521051:9523451: | Oatp30B | 0 |
| chr3L:10306201:10306701: | Or67d | 32998 |
| chr3R:29771601:29773701: | kay | 5607 |
| chr3L:14108001:14108651: | Sox21a | 0 |

| | | |
|---|---|---|
| chr3R:16726401:16727001: | Ubx | 7425 |
| chr2R:9376401:9377101: | CG13954 | 12366 |
| chr3L:10736751:10737851: | CG43245 | 34074 |
| chrX:17506801:17507901: | CG43658 | 53279 |
| chr2R:11107351:11107951: | luna | 8427 |
| chr2R:6102301:6102901: | Ars2 | 16677 |
| chr3L:6262501:6263301: | ImpL3 | 0 |
| chr3R:12467501:12468401: | Hsp70Ba | 0 |
| chr3R:19836601:19837301: | bnl | 0 |
| chr2R:14526351:14527301: | ttv | 85 |
| chr3L:10307801:10308651: | Or67d | 34598 |
| chr3L:2151551:2152151: | zormin | 0 |
| chr3R:14373051:14374001: | NK7 | 101 |
| chr2L:17621101:17621651: | CG15147 | 4730 |
| chr2R:8566001:8566401: | Cyp6a14 | 801 |
| chr3R:4738651:4741101: | Nep2 | 6774 |
| chr3L:603751:604301: | CG13893 | 0 |
| chr4:788801:789301: | MED26 | 0 |
| chr3R:8955251:8956001: | CG11966 | 7036 |
| chr3L:12609051:12609901: | caup | 0 |
| chr3L:10739101:10740601: | CG43245 | 31324 |
| chrX:4779101:4779651: | CG2871 | 1727 |
| chr3R:17920301:17920701: | cpo | 470 |
| chr2R:16499501:16500701: | Sema2a | 0 |
| chrX:18404351:18405001: | CG34328 | 4287 |
| chrX:7604351:7605251: | ct | 3897 |
| chr3R:9704301:9705551: | Unc-115a | 0 |
| chr3L:8559701:8560451: | rhea | 14701 |
| chr2L:16234701:16235301: | CG42817 | 3227 |
| chrX:1133801:1134951: | CG3655 | 0 |
| chr3L:738851:739601: | emc | 9799 |
| chrX:6012951:6013851: | mab-21 | 0 |
| chrX:1321551:1322401: | DAAM | 0 |
| chr2R:7692651:7693401: | LRR | 5501 |
| chr3R:9952801:9953301: | Art4 | 32759 |
| chr2R:6287301:6287751: | Pld | 45298 |
| chr2L:16587601:16589001: | CG31815 | 16821 |
| chrX:7607701:7608701: | ct | 447 |
| chr2R:23551201:23552201: | Dcp-1 | 0 |
| chr3R:16876651:16877151: | abd-A | 46603 |
| chr3L:11596351:11596951: | CG7368 | 251 |
| chr2L:16231001:16231751: | CG42817 | 0 |
| chrX:7226051:7226401: | CG1958 | 40503 |
| chrX:1380601:1381101: | CG32813 | 0 |
| chr2L:12529151:12530201: | bun | 16410 |

| | | |
|---|---|---|
| chr3R:16734251:16734901: | Ubx | 0 |
| chr3L:20294601:20295651: | gogo | 0 |
| chrX:18409951:18410501: | CG32549 | 0 |
| chr3L:15699551:15700601: | comm2 | 0 |
| chrX:7224951:7225301: | CG1958 | 41603 |
| chr3L:15729001:15729901: | comm | 572 |
| chr3R:29764001:29764851: | kay | 1144 |
| chrX:7629151:7629701: | ct | 20004 |
| chr3L:19170301:19172401: | CG14082 | 31327 |
| chr3L:15728151:15728801: | comm | 0 |
| chr2R:22592651:22593701: | dnr1 | 0 |
| chrX:11703001:11703501: | CG2444 | 1799 |
| chr3R:25782901:25783451: | CG4960 | 3847 |
| chrX:10102451:10103201: | RhoU | 15925 |
| chr2L:6252251:6253201: | Ddr | 0 |
| chr3R:7186851:7188451: | lap | 0 |
| chr3R:14382151:14383151: | soti | 453 |
| chr2R:13337151:13337801: | CG17048 | 16698 |
| chr3R:12477201:12478101: | Hsp70Ba | 9426 |
| chrX:7626601:7627701: | ct | 17454 |
| chrX:10302301:10302901: | alpha-Man-Ia | 23057 |
| chr3L:3417801:3418801: | sty | 6134 |
| chr3R:4071551:4072051: | CG42402 | 5888 |
| chr2L:1:6251: | CG11023 | 1278 |
| chr3L:20103751:20105651: | CG14186 | 0 |
| chrX:1375201:1375751: | AMPKalpha | 0 |
| chr3L:3419301:3419801: | sty | 5134 |
| chr2L:3354601:3355501: | E23 | 0 |
| chr3L:20949251:20950351: | fng | 0 |
| chr2R:9389801:9390351: | l | 0 |
| chr3L:3420051:3420701: | sty | 4234 |
| chrX:8785701:8786251: | Lim1 | 19553 |
| chr3R:29690851:29691601: | dmrt99B | 990 |
| chrX:7195901:7196501: | CG9650 | 58179 |
| chr3L:8546251:8546901: | rhea | 1251 |
| chr3L:616401:617301: | CG43337 | 5015 |
| chr3L:12627551:12628151: | caup | 17921 |
| chr3L:17757251:17757801: | CG32182 | 311 |
| chr3L:11586551:11587351: | rt | 419 |
| chr3R:22476551:22477101: | Nrx-1 | 8040 |
| chrX:6000901:6001651: | CG4766 | 105 |
| chr2R:22513751:22515101: | CG4610 | 21555 |
| chrX:17535701:17536151: | CG43658 | 25029 |
| chrX:10308851:10309501: | alpha-Man-Ia | 16457 |
| chr3R:30160301:30160851: | CG11498 | 0 |

| | | |
|---|---|---|
| chr3R:15374251:15374801: | Rbp | 0 |
| chr3L:10325001:10326001: | Or67d | 51798 |
| chr2R:24293801:24294301: | CG3394 | 0 |
| chr3R:19740701:19741251: | GluClalpha | 9987 |
| chr2L:16601201:16602101: | CG31815 | 3721 |
| chr2L:701401:702501: | ds | 12482 |
| chr3L:17872851:17873401: | CG7408 | 5695 |
| chr3R:23781701:23783051: | mbc | 0 |
| chr4:807551:808201: | MED26 | 18574 |
| chr2R:9440651:9442251: | ced-6 | 2686 |
| chr2L:12542901:12544501: | bun | 2110 |
| chr3R:27441301:27441651: | CG44290 | 358 |
| chr2L:13665851:13666501: | CG18507 | 327 |
| chr3R:15378901:15379551: | h-cup | 3400 |
| chrX:8924401:8925351: | CG43255 | 442 |
| chr2R:12704751:12705401: | CG42663 | 0 |
| chrX:11689501:11690001: | PhKgamma | 5136 |
| chr2L:12545151:12546851: | bun | 0 |
| chrX:14950451:14951501: | pdgy | 2031 |
| chr3L:5881151:5882451: | CG13287 | 0 |
| chr3R:27438351:27438751: | side | 870 |
| chr2R:21842901:21843601: | CG4372 | 4104 |
| chrX:10087901:10088451: | RhoU | 1375 |
| chr3R:16857351:16858151: | abd-A | 27303 |
| chrX:14952051:14952701: | pdgy | 831 |
| chr2L:12586801:12587551: | nub | 74 |
| chr3L:14037801:14038751: | Hsc70Cb | 783 |
| chr2L:5421551:5422101: | H15 | 17272 |
| chrX:11685951:11686801: | bif | 6982 |
| chrX:14953351:14953951: | pdgy | 0 |
| chrX:14970951:14971551: | eag | 15475 |
| chr2L:22018501:22019751: | tio | 0 |
| chr2L:10009751:10010351: | Trp1 | 371 |
| chr3L:629901:630401: | CG43337 | 7586 |
| chr3R:9928401:9929251: | Glut4EF | 53721 |
| chr3L:19793351:19793951: | SREBP | 0 |
| chr2L:1737001:1737701: | CG17646 | 4478 |
| chr2R:7597601:7598151: | scra | 7483 |
| chr3L:19790251:19792251: | Gyc76C | 1318 |
| chrX:11681651:11682251: | bif | 2682 |
| chr2R:9408751:9409851: | wun | 3612 |
| chr2R:7665501:7666051: | CG2064 | 0 |
| chr2L:22023951:22025151: | tio | 4744 |
| chrX:10324201:10324901: | alpha-Man-Ia | 1057 |
| chr2R:22569501:22570601: | CG11362 | 7195 |

| | | |
|---|---|---|
| chr3L:10769401:10769801: | CG43245 | 2124 |
| chrX:10344751:10345401: | CG43740 | 11100 |
| chr2L:714701:715751: | ds | 0 |
| chrX:10325151:10326051: | alpha-Man-Ia | 0 |
| chrX:8805651:8807101: | Lim1 | 0 |
| chr3R:12508651:12509251: | Hsp70Bc | 0 |
| chr3L:707601:708651: | CG13895 | 0 |
| chr2L:21632851:21633601: | Ac3 | 265 |
| chr2L:21221601:21222351: | CG8677 | 290 |
| chr3R:12502001:12502651: | Hsp70Bbb | 0 |
| chr2L:2241951:2242601: | CG4267 | 0 |
| chr3L:2712401:2713001: | Fife | 712 |
| chr2L:16857651:16858251: | CG31809 | 463 |
| chr2R:9426351:9427301: | Pdk | 0 |
| chr3L:5926251:5927051: | CG33523 | 2390 |
| chrX:20916501:20916901: | shakB | 10149 |
| chr2R:9413151:9414651: | wun2 | 0 |
| chr2L:6178451:6179551: | smal | 0 |
| chr3R:16763451:16764551: | CG31275 | 12118 |
| chr3R:27426001:27426501: | side | 10981 |
| chr3R:12505351:12506101: | Hsp70Bb | 0 |
| chr2R:4708951:4709501: | Nipped-B | 19516 |
| chr3L:959251:960151: | CG12038 | 33832 |
| chr3R:27988901:27989551: | betaTub97EF | 518 |
| chrX:900751:901351: | CG11663 | 6270 |
| chrX:11616201:11616851: | Met | 78 |
| chr3L:1208051:1208551: | LysB | 681 |
| chr3L:10517501:10518051: | S-Lap4 | 26641 |
| chr3L:20922101:20922651: | CG10589 | 5175 |
| chr3L:10516351:10517051: | S-Lap4 | 25491 |
| chr2L:5403401:5404051: | H15 | 229 |
| chrX:9809001:9810351: | CG32698 | 25779 |
| chr2R:17264851:17265801: | CG43108 | 6966 |
| chr2R:10675001:10675401: | stan | 1657 |
| chr3R:19498601:19500101: | CG15025 | 7487 |
| chr2L:6186401:6186901: | smal | 7269 |
| chr3L:4541601:4543301: | CG11357 | 0 |
| chrX:11621851:11623201: | Ptp10D | 0 |
| chr3L:2107251:2107951: | sls | 7660 |
| chr3L:11557151:11557801: | CG7394 | 4195 |
| chr3L:3452751:3453251: | eIF5B | 7726 |
| chrX:17963251:17963851: | CG12672 | 43755 |
| chr4:834101:834851: | Sox102F | 0 |
| chr3R:19495001:19495701: | CG15025 | 3887 |
| chr3R:7749851:7750401: | pyd3 | 0 |

| | | |
|---|---|---|
| chr3L:8034801:8035501: | bip1 | 23212 |
| chr2L:4594801:4595501: | HP6 | 16584 |
| chr2L:20494301:20494951: | CG34007 | 79 |
| chrX:2765451:2765901: | Syx4 | 21376 |
| chr2L:21617901:21619151: | CG2201 | 4929 |
| chr3L:11552851:11553901: | CG7394 | 8095 |
| chr2L:21237101:21238301: | Hr39 | 0 |
| chr4:557051:557601: | Thd1 | 253 |
| chr3L:10357801:10358201: | CG12362 | 54617 |
| chr2L:3302951:3303551: | CG9663 | 0 |
| chrX:2769251:2769651: | CG32795 | 20916 |
| chr3R:16829901:16830651: | abd-A | 0 |
| chrX:10059651:10060201: | Gr9a | 1426 |
| chrX:8893601:8895101: | Moe | 3231 |
| chr2L:12288451:12288951: | CG31862 | 49897 |
| chr2L:2221501:2222351: | papi | 754 |
| chrX:8891401:8892301: | Moe | 6031 |
| chr3L:685001:685951: | ebd1 | 0 |
| chr2R:12779901:12780801: | sca | 0 |
| chr2L:15109001:15109551: | CG15269 | 0 |
| chr3L:4975901:4976851: | CG17030 | 3918 |
| chr3L:681901:682851: | CG3386 | 0 |
| chr2R:12742951:12743551: | Mos | 9385 |
| chrX:2795851:2796951: | w | 0 |
| chr3R:19463101:19464701: | CG6255 | 36 |
| chr3R:15419051:15419751: | Atx2 | 10629 |
| chr3R:15420101:15421201: | Atx2 | 11679 |
| chr2L:21250451:21251251: | l | 9917 |
| chrX:2789501:2793401: | CG32795 | 0 |
| chr2R:5261901:5262401: | d4 | 2898 |
| chr2R:6501501:6502051: | jing | 0 |
| chr2L:4614201:4614851: | dpy | 0 |
| chr3L:11534301:11534951: | CG11714 | 11589 |
| chr3L:1176951:1178601: | bab2 | 0 |
| chr3L:8827051:8827951: | dally | 0 |
| chr3R:16811651:16812401: | abd-A | 17648 |
| chr3L:3625801:3626551: | dar1 | 0 |
| chr3L:10810801:10811601: | CG42831 | 3218 |
| chr3L:19731151:19731801: | CG42529 | 2158 |
| chr2L:2201901:2203801: | CG31668 | 2365 |
| chrX:14082551:14083301: | Ste | 20735 |
| chr3R:14447251:14447651: | natalisin | 28659 |
| chrX:20230801:20232001: | sw | 4381 |
| chr2L:2200201:2200751: | CG34172 | 2290 |
| chr2R:19825401:19826851: | Toll-7 | 0 |

| | | |
|---|---|---|
| chr3R:14473001:14473451: | cv-c | 8262 |
| chr2L:2196601:2197451: | CG34172 | 461 |
| chr2R:6827901:6828651: | CG15236 | 0 |
| chr2R:24229651:24230201: | CG30419 | 0 |
| chr3R:14468201:14468701: | cv-c | 13012 |
| chr3L:20217101:20217651: | CG6933 | 2769 |
| chr2L:21582301:21582651: | nompB | 6772 |
| chr3L:5184351:5185551: | Srp54k | 31674 |
| chr3R:26229501:26230501: | CG12290 | 0 |
| chr3L:21644551:21645201: | CG43980 | 0 |
| chrX:14653701:14654401: | NetA | 0 |
| chr3L:20185751:20187051: | Ssk | 1471 |
| chr2L:21572551:21573351: | Lamp1 | 38 |
| chr3R:6157001:6157601: | CG31559 | 0 |
| chr2L:21571501:21572201: | Lamp1 | 1188 |
| chr2L:21570551:21571101: | Lamp1 | 2288 |
| chrX:23205301:23206551: | ATbp | 99299 |
| chr2L:1805701:1806251: | Gr22a | 10785 |
| chr2L:2178251:2179151: | aop | 0 |
| chr4:507151:508851: | zfh2 | 5342 |
| chr3L:21476301:21477351: | croc | 0 |
| chr3L:21479401:21480201: | croc | 2923 |
| chrX:9870101:9870551: | CG45061 | 1151 |
| chrX:23210651:23212101: | ATbp | 104649 |
| chr3L:20198101:20198701: | CG42674 | 0 |
| chr3R:31937401:31938201: | CG2003 | 39920 |
| chrX:23212551:23213801: | ATbp | 106549 |
| chr4:501351:502151: | zfh2 | 0 |
| chrX:23214101:23218701: | ATbp | 108099 |
| chr3L:21485101:21485801: | Neu2 | 4785 |
| chr3L:23066001:23066601: | RpL10 | 28870 |
| chr3L:11493301:11493801: | chrb | 5752 |
| chr2L:15056601:15057601: | ck | 261 |
| chr3R:15476051:15476951: | AdamTS-A | 0 |
| chr3L:140701:141301: | CG6845 | 5132 |
| chr2L:2164601:2165201: | CG15382 | 8849 |
| chr3R:31945551:31946151: | CG2003 | 31970 |
| chrX:9880751:9881501: | CG9689 | 452 |
| chr2L:2163551:2164101: | CG15382 | 7799 |
| chr3L:10851051:10851651: | NA-tna | 6238 |
| chr2L:2162451:2163301: | CG15382 | 6699 |
| chr3L:11487001:11488001: | chrb | 0 |
| chr3L:10852201:10853201: | NA-tna | 4688 |
| chr2L:2161151:2162051: | CG15382 | 5399 |
| chr3L:10856101:10858101: | tna | 0 |

| | | |
|---|---|---|
| chrX:14037901:14038701: | Ste | 8794 |
| chr3L:130751:132801: | Pdk1 | 1302 |
| chr4:907701:908351: | unc-13 | 3980 |
| chr2R:7170751:7171401: | CG11060 | 4485 |
| chr3L:21498801:21499701: | Hr78 | 1308 |
| chr2L:21309351:21309851: | Mondo | 199 |
| chr3L:128201:129801: | Pdk1 | 0 |
| chr2R:6871351:6872901: | coro | 0 |
| chr2L:20281751:20282301: | CG12617 | 12392 |
| chr2L:1062501:1062901: | IA-2 | 14449 |
| chr3L:11428301:11429001: | CG7560 | 3144 |
| chr2L:21540901:21541301: | His3 | 0 |
| chr3L:21509551:21510001: | M6 | 1700 |
| chr3R:7309551:7310201: | rn | 0 |
| chr3L:21604401:21604851: | S1P | 12525 |
| chr3L:23090151:23090701: | RpL10 | 4770 |
| chr2L:21534401:21534701: | His3 | 0 |
| chr3L:3542251:3543901: | Eip63E | 26501 |
| chrX:23242851:23243501: | ATbp | 136849 |
| chr3L:3544801:3545501: | Eip63E | 29051 |
| chrX:9905001:9905751: | CG1986 | 5375 |
| chr2L:21529501:21529901: | His3 | 0 |
| chr2R:6885551:6886101: | Spn42De | 0 |
| chr3L:21598051:21598951: | TfAP-2 | 11224 |
| chrX:23246201:23246901: | ATbp | 140199 |
| chr2R:7150351:7150901: | pk | 0 |
| chrX:23249601:23250301: | ATbp | 143599 |
| chr2L:21524701:21525051: | His3 | 0 |
| chr4:931501:932101: | eIF4G1 | 787 |
| chr3L:1102101:1103001: | bab1 | 1013 |
| chr4:932501:933051: | eIF4G1 | 1787 |
| chr2R:7146751:7147351: | Spn43Aa | 341 |
| chrX:23253001:23253901: | ATbp | 146999 |
| chr4:933951:935001: | mGluR | 1818 |
| chr2L:2130051:2131001: | GlyP | 0 |
| chr2L:21519851:21520201: | His3 | 0 |
| chrX:20158801:20159701: | Rab35 | 171 |
| chr3L:1098451:1099201: | bab1 | 1888 |
| chr4:936301:936901: | mGluR | 0 |
| chr4:457501:458001: | bip2 | 8827 |
| chr2L:16972601:16973101: | beat-IIIb | 0 |
| chr3L:21584501:21585251: | TfAP-2 | 1577 |
| chr2L:21514801:21515151: | His3 | 0 |
| chr2L:20310351:20311451: | spir | 0 |
| chr4:453801:454451: | CaMKI | 8298 |

| | | |
|---|---|---|
| chr2L:15013351:15013951: | GABA-B-R1 | 2380 |
| chr2L:21509751:21510101: | His3 | 0 |
| chrX:13986451:13987101: | mamo | 3426 |
| chr3R:6002901:6003301: | Rm62 | 5278 |
| chrX:13994551:13995551: | ben | 794 |
| chr2L:21504701:21505051: | His3 | 0 |
| chr2L:22505101:22505651: | RpL5 | 31233 |
| chr2L:15745301:15746501: | CycE | 1649 |
| chr2L:15747201:15748501: | CycE | 0 |
| chr3R:6007501:6008151: | Rm62 | 428 |
| chr2L:21499601:21500101: | His3 | 0 |
| chrX:23278001:23281101: | ATbp | 171999 |
| chr2L:14995401:14996051: | mol | 1508 |
| chr2L:21494601:21494951: | His3 | 0 |
| chrX:23282001:23284301: | ATbp | 175999 |
| chr4:431851:432401: | lgs | 11510 |
| chrX:21634001:21634601: | DIP1 | 3854 |
| chr2L:21489551:21489901: | His3 | 0 |
| chr2L:1885951:1886851: | Wdr62 | 1330 |
| chr2L:15762351:15762951: | Gli | 0 |
| chr2L:1118751:1119651: | CG14340 | 5811 |
| chr2L:21484351:21484851: | His3 | 0 |
| chr3R:32015201:32016001: | heph | 0 |
| chr2R:7106301:7106801: | esn | 19986 |
| chr2L:16049551:16050151: | beat-Ia | 0 |
| chr2L:21479351:21479751: | His3 | 0 |
| chr2L:21474301:21474701: | His3 | 0 |
| chrX:66601:67401: | CG17636 | 59313 |
| chr2L:21469251:21469701: | His3 | 0 |
| chr4:404251:404701: | dati | 8647 |
| chr2L:21464251:21464651: | His3 | 0 |
| chr2R:5401601:5402101: | laccase2 | 11643 |
| chr2L:22549751:22550251: | CG17493 | 0 |
| chr2L:21459151:21459601: | His3 | 0 |
| chr3L:40001:40951: | CG43149 | 13800 |
| chr2L:21454051:21454551: | His3 | 0 |
| chr2L:21439851:21440351: | His3 | 0 |
| chr2L:21435001:21435451: | His3 | 0 |
| chr2L:21419801:21420251: | His3 | 0 |
| chr2L:21430151:21430401: | His3 | 0 |
| chr2L:21424801:21425451: | His3 | 0 |
| chr3R:32071001:32072401: | Map205 | 2561 |
| chr3R:26536751:26537451: | scrib | 402 |
| chr2R:7049401:7050001: | lbm | 164 |
| chr2R:6992501:6993101: | CG30159 | 0 |

| | | |
|---|---|---|
| chr2L:1954301:1955151: | CG33673 | 7574 |
| chrX:6551:7351: | CG17636 | 119363 |
| chr2L:1970201:1971101: | erm | 0 |
| chr2L:2013751:2014351: | CG4238 | 3893 |
| chr2L:1971351:1971851: | erm | 990 |
| chr2L:1972451:1973351: | Der-1 | 606 |
| chr3R:3788401:3789601: | CG41128 | 38039 |
| chrX:22438751:22439301: | CG14621 | 4926 |
| chr4:306401:307051: | Syt7 | 0 |
| chr4:303351:304001: | Syt7 | 2480 |
| chr4:302451:303101: | Syt7 | 3380 |
| chr4:295251:295951: | Syt7 | 10530 |
| chr4:294451:295001: | Syt7 | 11480 |
| chr2R:4169201:4169801: | RNASEK | 18326 |
| chrX:23430601:23433801: | ATbp | 324599 |
| chr2L:15911601:15912701: | CG13244 | 4187 |
| chrX:23438701:23440101: | ATbp | 332699 |
| chrX:23440301:23440901: | ATbp | 334299 |
| chrX:22382651:22383701: | CG14621 | 60526 |
| chr4:207401:207951: | CG31998 | 1457 |
| chr4:183351:184001: | Arl4 | 3963 |
| chrX:23535051:23538701: | ATbp | 429049 |
| chr4:1257851:1258501: | Cadps | 27015 |
| chr4:1260001:1260401: | Cadps | 29165 |
| chrX:22260201:22260401: | CG41562 | 8576 |
| chrX:22258251:22259301: | CG41562 | 6626 |
| chr4:1268401:1269001: | Cadps | 37565 |
| chr4:1274751:1275151: | Cadps | 43915 |
| chr3R:3588801:3589801: | Parp | 1748 |
| chr4:1294901:1295201: | Cadps | 64065 |
| chr4:1318551:1319451: | Cadps | 87715 |
| chr4:53601:54201: | ci | 2840 |
| chr4:52701:53251: | ci | 3790 |
| chr2R:3934351:3934801: | uex | 14624 |
| chr4:41301:41901: | PlexB | 1877 |
| chr4:33701:34201: | PlexB | 9577 |
| chr3L:23582601:23583151: | AGO3 | 28009 |
| chr3R:3344651:3345251: | spok | 21842 |
| chr2R:3750401:3751051: | RYa | 103056 |
| chr2L:23093101:23093501: | Spf45 | 24716 |
| chr2L:23096951:23097851: | Spf45 | 28566 |
| chr2L:23099601:23101451: | Spf45 | 31216 |
| chr2R:3731151:3731701: | RYa | 83806 |
| chr2L:23184951:23185401: | CG12567 | 10468 |
| chr2L:23186151:23186551: | CG12567 | 11668 |

| | | |
|---|---|---|
| chr2L:23189351:23189851: | CG12567 | 14868 |
| chr3R:3169651:3169851: | Pzl | 93731 |
| chr2L:23512601:23513851: | Tim23 | 199024 |
| chr2R:3311851:3312301: | CG41242 | 52101 |
| chr3L:24210051:24210701: | nvd | 114165 |
| chr3L:24286851:24287401: | nvd | 37465 |
| chr2R:3062801:3063451: | dpr21 | 3048 |
| chr2R:3062001:3062451: | dpr21 | 4048 |
| chr3L:24392301:24392801: | nvd | 67436 |
| chr3R:2568551:2569051: | Pzl | 694531 |
| chr2R:2904501:2905001: | CG41378 | 57794 |
| chr3R:2449851:2450251: | Pzl | 813331 |
| chr2R:2841851:2842401: | CG41378 | 4307 |
| chr3L:24608801:24610351: | mRpS5 | 71566 |
| chr3L:24610651:24611301: | mRpS5 | 73416 |
| chr3L:24611651:24612201: | mRpS5 | 74416 |
| chr3L:24614101:24614651: | mRpS5 | 76866 |
| chr3L:24618601:24619201: | mRpS5 | 81366 |
| chr3L:24622751:24624001: | mRpS5 | 85516 |
| chr3L:24626901:24627351: | mRpS5 | 89666 |
| chr2R:2805151:2805801: | CG41378 | 40907 |
| chr3L:24656601:24657001: | scro | 67161 |
| chr3L:24674301:24674951: | scro | 49211 |
| chr3L:24684851:24685251: | scro | 38911 |
| chr3R:2116601:2117101: | Pzl | 1146481 |
| chr3L:25085851:25086451: | Set1 | 21143 |
| chr2R:2260251:2260751: | CG17684 | 1364 |
| chr3L:25804051:25804701: | FASN3 | 41273 |
| chr3L:25805301:25805951: | FASN3 | 42523 |
| chr3L:25874401:25875001: | FASN3 | 111623 |
| chr2R:1528651:1529301: | Maf1 | 1557 |
| chr2R:1524101:1524801: | Maf1 | 2294 |
| chr2R:1512451:1513051: | Maf1 | 14044 |
| chr3R:951951:952401: | Myo81F | 384876 |
| chr2R:1233401:1234601: | CG40191 | 29539 |
| chr2R:870051:870601: | CG46302 | 15238 |
| chr2R:748751:749101: | CG46302 | 105713 |
| chr2R:601751:602351: | CG46302 | 252463 |
| chr2R:553601:554001: | CG45781 | 226834 |
| chr2R:552751:553301: | CG45781 | 225984 |
| chr3L:27007451:27011201: | CG40228 | 72705 |
| chr3L:27068551:27073601: | CG40228 | 10305 |
| chr3L:27075001:27075451: | CG40228 | 8455 |
| chr3L:27075951:27077551: | CG40228 | 6355 |
| chr3L:27079151:27079601: | CG40228 | 4305 |

| | | |
|---|---|---|
| chr3L:27081451:27082251: | CG40228 | 1655 |
| chr3L:27129901:27130501: | vtd | 6024 |
| chr2R:318751:319351: | CG45781 | 7417 |
| chr3L:27289601:27290101: | Dbp80 | 10286 |
| chr3L:27356801:27357501: | Dbp80 | 77486 |
| chr2R:1151:2701: | CG45781 | 324067 |

| **Coordinates_Interphase_Only** | **nearst_gene** | **distance_nearest_gene** |
|---|---|---|
| chrY:1050951:1051901: | ARY | 348522 |
| chrY:124051:124901: | kl-5 | 12585 |
| chrY:132251:132801: | kl-5 | 4685 |
| chrY:1416751:1417151: | Ppr-Y | 219423 |
| chrY:1550501:1551001: | Ppr-Y | 85573 |
| chrY:200551:201051: | PRY | 8532 |
| chrY:203501:203901: | PRY | 11482 |
| chrY:3652701:3653101: | CCY | 2706 |
| chrY:3667101:3667501: | CCY | 17106 |
| chrY:853601:854851: | ARY | 151172 |
| chrY:860151:860651: | ARY | 157722 |
| chrY:877851:878351: | ARY | 175422 |
| chrY:883751:884301: | ARY | 181322 |
| chrY:885801:886651: | ARY | 183372 |
| chrY:887401:887901: | ARY | 184972 |
| chrY:917651:918151: | ARY | 215222 |
| chr2R:11450551:11451051: | E | 992 |
| chr3L:4406351:4407301: | DOR | 0 |
| chrX:17826901:17827401: | Socs16D | 163 |
| chr2R:22688701:22690101: | Ppa | 0 |
| chrX:7891351:7892501: | Nek2 | 0 |
| chr2R:21474801:21476951: | Sdc | 4225 |
| chr3L:2589801:2590301: | dos | 388 |
| chr2R:12586751:12590351: | Sin3A | 2566 |
| chr3L:15141651:15142351: | Pdi | 489 |
| chrX:2009951:2010851: | east | 9 |
| chr3L:15507301:15508401: | CG7841 | 0 |
| chr3R:24104651:24106151: | Syx1A | 0 |
| chr2L:19574851:19577201: | spi | 114 |
| chr2R:13041551:13042501: | CG13323 | 4755 |
| chr2R:10806351:10807751: | wde | 1187 |
| chr2L:5977351:5981001: | chic | 17 |
| chr2L:4281201:4283901: | tutl | 0 |
| chr3R:14284851:14287151: | trx | 0 |
| chr3R:14282551:14284601: | trx | 2302 |
| chr2L:16325551:16326251: | cact | 0 |
| chr3R:5216101:5216701: | cno | 755 |

| | | |
|---|---|---|
| chrX:7324251:7325051: | Dok | 38 |
| chr3R:20948901:20949701: | Atpalpha | 188 |
| chrX:18309051:18310101: | upd1 | 0 |
| chr3L:14409451:14410951: | CG3919 | 0 |
| chr2R:7381501:7382401: | Dscam1 | 0 |
| chr2R:13957351:13958451: | AGO1 | 0 |
| chrX:8054101:8055501: | fs | 5935 |
| chr2L:9705201:9706401: | CG31710 | 0 |
| chr3L:13924751:13925851: | Rgl | 0 |
| chr2R:25084251:25084751: | gol | 0 |
| chr3L:9614401:9615101: | LanB2 | 0 |
| chr2R:13822551:13823101: | fl | 1452 |
| chr3L:12265651:12266201: | app | 640 |
| chr2R:18799101:18800901: | MFS15 | 2424 |
| chrX:5083451:5084751: | CG32767 | 17 |
| chr2R:13824001:13824701: | fl | 0 |
| chr3L:20517201:20517951: | CG5059 | 1442 |
| chr2R:18802051:18803251: | MFS14 | 476 |
| chrX:8056201:8059351: | fs | 2085 |
| chr3R:4174251:4175551: | CG12581 | 0 |
| chrX:18312301:18313151: | upd1 | 3011 |
| chr3R:10267051:10267601: | CG43143 | 2746 |
| chrX:12202451:12204101: | CG1924 | 23826 |
| chr3R:7084551:7087351: | alphaTub84B | 0 |
| chr2L:18444551:18445101: | RpS26 | 1275 |
| chr3R:10048601:10049101: | CG12811 | 0 |
| chrX:3134601:3135901: | N | 0 |
| chr2R:11892601:11893301: | eEF1alpha1 | 1467 |
| chr3R:4478001:4479301: | CG31523 | 0 |
| chr3L:13922751:13923301: | Rgl | 2034 |
| chrX:616751:617451: | CG13366 | 2744 |
| chrX:13284201:13285101: | HDAC4 | 531 |
| chr2R:9957901:9958951: | Mef2 | 0 |
| chr3R:30563051:30564051: | CG34155 | 2853 |
| chrX:19264201:19264801: | Vav | 2382 |
| chr2R:12591301:12592801: | Sin3A | 116 |
| chr2R:10422951:10424651: | Prx2540-1 | 0 |
| chr3L:2585601:2586651: | msn | 0 |
| chrX:2343351:2343851: | mRpL14 | 798 |
| chrX:18789851:18790351: | Atg101 | 1317 |
| chr3L:4409801:4410301: | DOR | 2859 |
| chr2R:17547901:17548751: | MESR4 | 133 |
| chr3L:5657501:5658051: | sif | 0 |
| chrX:2011951:2012651: | east | 1092 |
| chrX:19321751:19323951: | out | 0 |

| | | |
|---|---|---|
| chr3L:4252051:4252601: | ago | 1705 |
| chr2L:9613151:9613801: | Trx-2 | 0 |
| chr3R:6816201:6816851: | Dfd | 24366 |
| chr3R:11956301:11957001: | Hsp70Aa | 0 |
| chrX:7320601:7321351: | Atg5 | 2419 |
| chr3L:14624751:14625301: | CG9628 | 1956 |
| chr2L:19570951:19573251: | spi | 4064 |
| chr2R:9283951:9285151: | Pkn | 0 |
| chr3L:4252951:4253551: | ago | 755 |
| chr3L:13919701:13921001: | Rgl | 4334 |
| chrX:12760201:12760751: | ade5 | 0 |
| chrX:10729301:10730551: | ras | 13951 |
| chr3R:8119701:8120351: | CG7878 | 2374 |
| chr2R:10420901:10422351: | CG33474 | 0 |
| chr2R:9954751:9956051: | Mef2 | 2807 |
| chr3L:7134001:7135301: | melt | 0 |
| chr2R:10534051:10536651: | lola | 6640 |
| chr2R:16856151:16857551: | Dek | 0 |
| chr3R:11958151:11958851: | Hsp70Ab | 0 |
| chr3R:9502301:9503001: | RhoL | 0 |
| chr2L:594401:595551: | Gsc | 0 |
| chr2L:7048901:7049551: | Mnn1 | 6750 |
| chr3R:7088601:7089351: | alphaTub84B | 2003 |
| chr3L:1794051:1795351: | dlt | 0 |
| chrX:12574651:12575301: | hwt | 6200 |
| chr3R:25880251:25881101: | Tsp96F | 326 |
| chr3R:16449101:16450051: | Pak3 | 3770 |
| chrX:10731151:10732401: | ras | 12101 |
| chr2L:15491801:15492551: | lace | 6576 |
| chr2R:21047601:21048501: | shg | 8696 |
| chr3R:23669301:23670501: | CG31145 | 0 |
| chr3R:11340051:11340501: | pros | 11572 |
| chrX:19259751:19260501: | rictor | 739 |
| chr3L:15825401:15826451: | DCP2 | 0 |
| chr3L:11821651:11822401: | CG11597 | 0 |
| chrX:8409651:8412151: | lawc | 0 |
| chr2L:3803101:3804251: | Reph | 0 |
| chr3R:18169851:18170451: | wrd | 2095 |
| chrX:19631201:19631901: | CG14212 | 5497 |
| chrX:11350501:11351401: | CG1737 | 0 |
| chr3R:28330451:28331251: | larp | 5189 |
| chr3L:16483401:16484501: | aos | 0 |
| chr3L:11819551:11820401: | CG11597 | 1874 |
| chr2R:8241801:8242351: | pnut | 4531 |
| chr2L:4371351:4372301: | Traf4 | 9427 |

| | | |
|---|---|---|
| chrX:16441901:16442201: | rngo | 469 |
| chr3R:18171851:18173151: | wrd | 4095 |
| chr3L:19909951:19910451: | Prp3 | 0 |
| chrX:5754951:5755901: | CG15771 | 0 |
| chr2R:12025051:12025551: | otk2 | 0 |
| chr3L:17421251:17421901: | blot | 0 |
| chrX:6861351:6862051: | CG43736 | 1837 |
| chr3L:14775651:14776051: | mop | 78 |
| chr2L:21899001:21900001: | CG11629 | 713 |
| chrX:6279051:6279901: | Ubi-p5E | 0 |
| chrX:13396551:13397101: | CG1673 | 0 |
| chr2R:9552601:9553501: | CG1888 | 4467 |
| chrX:6668801:6669551: | CG3198 | 4422 |
| chr2R:24525551:24526201: | Brca2 | 0 |
| chr2R:13223501:13224101: | Fsn | 1043 |
| chr3L:12515601:12516051: | tral | 543 |
| chr3L:16023001:16023701: | Notum | 0 |
| chr3L:1334801:1336601: | CG2211 | 0 |
| chrX:2136651:2137351: | ph-p | 1020 |
| chr3L:18839901:18840951: | Indy | 5316 |
| chrX:2135351:2135851: | ph-p | 2520 |
| chr3L:15336151:15337001: | Toll-6 | 0 |
| chr3L:18834551:18835051: | Indy | 11216 |
| chrX:3686901:3689651: | Mnt | 15101 |
| chr2R:15675101:15676951: | fus | 0 |
| chr2R:11603151:11603951: | tou | 12094 |
| chr2R:11604451:11605601: | tou | 10444 |
| chr3R:10886501:10887401: | Leash | 962 |
| chr3R:10889001:10889751: | Adk3 | 0 |
| chr2L:19164651:19166251: | CG17568 | 15451 |
| chrX:13089651:13090451: | hep | 0 |
| chrX:13739351:13740001: | CG11151 | 0 |
| chrX:13087501:13087951: | hep | 2483 |
| chr2L:107701:108351: | Sam-S | 799 |
| chr3R:10355951:10356951: | jumu | 5770 |
| chr2L:7976701:7977151: | CG7231 | 6795 |
| chr2L:8949651:8950701: | C1GalTA | 515 |
| chr2L:8951001:8951901: | C1GalTA | 0 |
| chr2L:18844801:18845451: | CG17321 | 0 |
| chr3R:13644551:13645601: | sqd | 703 |
| chr3R:12719601:12720301: | Men | 2141 |
| chr3L:11214401:11216151: | CG42671 | 0 |
| chrX:5516751:5518701: | CG12730 | 144 |
| chr3R:9229001:9230751: | pum | 6931 |
| chrX:6981401:6982301: | bou | 4185 |

| | | |
|---|---|---|
| chrX:6983501:6984201: | bou | 2285 |
| chr2R:11198501:11199251: | Syx6 | 20288 |
| chr2R:11196851:11197701: | Syx6 | 21838 |
| chr3L:3789101:3789551: | CG32266 | 1253 |
| chr3L:3787101:3788251: | CG32266 | 2553 |
| chr3R:20829001:20830601: | AdSS | 0 |
| chr2R:18972851:18973551: | Topors | 240 |
| chrX:4686101:4686651: | Pp2C1 | 252 |
| chr2L:4028101:4029351: | ed | 2026 |
| chr3R:18658501:18659151: | CG31224 | 158 |
| chr3L:4692001:4693001: | RhoGEF64C | 0 |
| chr3L:21049201:21049951: | siz | 14667 |
| chrX:12660051:12660901: | Tis11 | 637 |
| chrX:8060051:8061701: | mys | 0 |
| chr2L:4375051:4375501: | CG17612 | 7234 |
| chrX:11805051:11805951: | Hsc70-3 | 1166 |
| chr2R:16104051:16104851: | Rho1 | 1365 |
| chr3L:5654201:5654851: | lin-28 | 0 |
| chr3R:8658651:8659801: | bel | 960 |
| chrX:4684201:4684801: | Pp2C1 | 2102 |
| chr2L:4820201:4821751: | CG15628 | 23 |
| chr2R:10540201:10543351: | lola | 0 |
| chrX:16943351:16944701: | RhoGAP15B | 62 |
| chr2R:23203801:23204701: | CG34371 | 0 |
| chr3L:22720301:22721401: | mael | 0 |
| chrX:6354051:6354651: | kdn | 3423 |
| chr2R:13219101:13219651: | CG4630 | 0 |
| chrX:5885351:5886151: | CG16721 | 5483 |
| chr3R:14277651:14279601: | trx | 7302 |
| chr2L:3805401:3806751: | Reph | 1857 |
| chrX:17170401:17171151: | baz | 10396 |
| chr3L:18890401:18891401: | CG3961 | 0 |
| chr3L:13113651:13114551: | trn | 0 |
| chrX:7959051:7959551: | CG15330 | 2988 |
| chrX:10930451:10931751: | Myo10A | 0 |
| chr2R:21480451:21481201: | Sdc | 0 |
| chr2R:16110451:16111501: | Asph | 121 |
| chr3R:22673201:22674501: | Gclm | 12009 |
| chr3R:21198851:21199451: | CG5862 | 15628 |
| chr3L:1485551:1486151: | stet | 3396 |
| chr3L:19908101:19909401: | Su | 9 |
| chr3R:18167751:18169351: | wrd | 0 |
| chr3R:14723501:14724351: | kibra | 0 |
| chr2R:14755651:14756251: | CG10151 | 37 |
| chr3R:30270651:30271801: | hdc | 6131 |

| | | |
|---|---|---|
| chrX:6985651:6987651: | bou | 0 |
| chr3R:11763301:11764301: | Lk6 | 156 |
| chr2R:16338751:16339301: | Vha44 | 2115 |
| chr2R:18288401:18289251: | CG14502 | 0 |
| chrX:4690751:4691251: | ctp | 2952 |
| chr2R:13038601:13039201: | CG13323 | 8055 |
| chr2R:6195801:6196751: | Ptr | 0 |
| chr2L:21753501:21754151: | step | 3315 |
| chrX:13278201:13279151: | HDAC4 | 6481 |
| chr2R:13565851:13566301: | CG42807 | 6458 |
| chrX:7095901:7096351: | Sxl | 1702 |
| chr2R:18805951:18806601: | MFS14 | 2225 |
| chr3L:14780951:14781551: | mop | 4823 |
| chr3R:9235951:9237801: | pum | 0 |
| chr2R:22403151:22404001: | CG6044 | 176 |
| chr2R:13826001:13827101: | CG6329 | 0 |
| chr3L:14201001:14201501: | saturn | 1822 |
| chrX:19866001:19867201: | Hers | 0 |
| chrX:6866051:6866501: | CG14434 | 693 |
| chr2L:4031051:4031551: | ed | 0 |
| chr2R:15931101:15932051: | bdg | 0 |
| chrX:3861101:3861551: | CG2930 | 7422 |
| chr3R:27281101:27282151: | mrt | 510 |
| chrX:3136151:3137051: | N | 1282 |
| chrX:5086201:5087751: | rg | 443 |
| chr2R:16102651:16103751: | Rho1 | 0 |
| chr2R:10417801:10418751: | Prx2540-2 | 0 |
| chrX:9466251:9466951: | CG44815 | 20040 |
| chrX:8051301:8053701: | fs | 7735 |
| chr2R:22798201:22798701: | CG42260 | 0 |
| chrX:3691301:3692801: | Mnt | 19501 |
| chrX:12661351:12662001: | Tis11 | 1937 |
| chrX:6352151:6353601: | kdn | 1523 |
| chr2R:9547951:9548601: | CG1888 | 0 |
| chr3R:11542851:11543601: | dpr4 | 0 |
| chr2L:9607851:9608601: | gcm2 | 0 |
| chr2L:16496401:16496951: | Tpr2 | 4862 |
| chr2R:15372851:15373551: | Pms2 | 1332 |
| chr2R:19141501:19143401: | ena | 0 |
| chr3R:24046501:24048401: | Rox8 | 0 |
| chr2R:14306901:14308451: | Shroom | 0 |
| chr3L:21046951:21048401: | siz | 12417 |
| chr2R:20831601:20832151: | sktl | 3877 |
| chr3L:15146601:15147401: | FucTA | 0 |
| chr3L:1746601:1747151: | CG13921 | 0 |

| | | |
|---|---|---|
| chrX:5886601:5887451: | CG16721 | 6733 |
| chr2R:14981651:14982751: | pcs | 389 |
| chr3R:25646951:25648301: | jigr1 | 9109 |
| chr2L:5981701:5983801: | eIF4A | 0 |
| chr2R:11597501:11598201: | tou | 17844 |
| chr2R:8236901:8238151: | pnut | 0 |
| chr3R:23345351:23348151: | DNApol-epsilon255 | 0 |
| chr2R:17097051:17098101: | GstS1 | 1400 |
| chrX:7966901:7968351: | sws | 0 |
| chrX:11806901:11807801: | Hsc70-3 | 0 |
| chr3R:18007601:18008051: | DNaseII | 11324 |
| chr2R:9291951:9293101: | Drep2 | 96 |
| chrX:7096951:7097701: | Sxl | 352 |
| chr3L:4102401:4103001: | CG14995 | 255 |
| chr2R:22796701:22798001: | CG42260 | 288 |
| chrX:13276901:13278001: | HDAC4 | 7631 |
| chrX:3682401:3682951: | Mnt | 10601 |
| chr2L:7887051:7887901: | CG7102 | 0 |
| chr2R:14432051:14432551: | phyl | 0 |
| chrX:18647101:18647501: | Rip11 | 147 |
| chrX:9042101:9043001: | Nost | 11897 |
| chr2R:11202101:11203601: | Syx6 | 15938 |
| chr3R:14722101:14722851: | kibra | 1410 |
| chr3L:16572251:16572701: | Mipp1 | 633 |
| chr3R:5672301:5673051: | Atg17 | 2589 |
| chr3R:9226201:9227651: | pum | 10031 |
| chr3R:23887001:23887601: | Pli | 2451 |
| chr3L:19906801:19907551: | Su | 1859 |
| chr2R:22401801:22402551: | babos | 0 |
| chr3R:26041601:26042501: | gro | 0 |
| chr3R:28325951:28327501: | larp | 8939 |
| chrX:1232501:1233201: | CG3638 | 3483 |
| chr2L:2752551:2756001: | Pgk | 0 |
| chr3R:30272551:30273501: | hdc | 4431 |
| chr3L:347551:348151: | mth | 595 |
| chrX:5746651:5747351: | IntS6 | 0 |
| chr3R:9052651:9053551: | Kdm2 | 157 |
| chr2L:8957651:8958651: | CG31886 | 400 |
| chr2L:16485501:16487301: | dac | 0 |
| chr3R:14817701:14818451: | CG14857 | 15929 |
| chr2R:10427751:10431001: | RanBPM | 0 |
| chr3L:22402751:22403301: | Ten-m | 4562 |
| chr2R:17592751:17593751: | Smurf | 0 |
| chr2L:5542751:5543351: | Lam | 272 |
| chr3R:18912751:18913201: | CG14291 | 0 |

| | | |
|---|---|---|
| chr3R:16452801:16453451: | Pak3 | 370 |
| chrX:6867851:6868601: | CG14434 | 658 |
| chr2R:21712851:21715001: | CG30403 | 1239 |
| chr2L:160651:162101: | spen | 1620 |
| chr2R:22685701:22687101: | Ppa | 1886 |
| chrX:3137901:3138401: | N | 3032 |
| chr3R:11758301:11762051: | Lk6 | 2406 |
| chr2R:12452951:12454501: | Lac | 0 |
| chr2R:5772951:5773501: | ZnT41F | 9357 |
| chr3R:18298051:18298701: | l | 178 |
| chr3L:16408051:16409251: | fax | 2577 |
| chr2R:20595851:20596901: | HnRNP-K | 8301 |
| chr2L:19161351:19161901: | CG17568 | 19801 |
| chr3L:22723101:22723551: | CG14451 | 1011 |
| chrX:18243201:18244151: | upd2 | 0 |
| chr2L:20638201:20638801: | CG31680 | 1320 |
| chrX:2336301:2336751: | Ilp6 | 1644 |
| chrX:12663251:12663751: | Tis11 | 3837 |
| chr3R:11773251:11775651: | l | 3477 |
| chr3R:22133301:22133901: | SKIP | 0 |
| chrX:6283401:6283901: | CG11700 | 0 |
| chr3L:17658451:17659451: | CG7484 | 3835 |
| chr3L:20398501:20399001: | trbl | 2608 |
| chr3L:17988551:17990351: | CG32192 | 49966 |
| chr3L:9465751:9466401: | CG44838 | 4702 |
| chr2R:8145751:8146401: | kermit | 10253 |
| chr2R:9198601:9198951: | unpg | 3397 |
| chrX:10738651:10739601: | ras | 4901 |
| chrX:6988651:6989451: | ogre | 1867 |
| chr3R:6810451:6811301: | Dfd | 18616 |
| chr2R:16335601:16336301: | Hmgs | 0 |
| chrX:17710651:17711301: | CG42684 | 508 |
| chrX:1233701:1235751: | CG3638 | 933 |
| chr3R:18913701:18914401: | Xrp1 | 567 |
| chr3R:19163701:19165151: | sqz | 0 |
| chr3R:21059801:21061251: | Mvl | 0 |
| chr2L:14489851:14491201: | noc | 0 |
| chr2R:19143801:19144451: | ena | 2170 |
| chr2L:11968801:11969351: | CG16965 | 12240 |
| chr3R:24295451:24296151: | crb | 374 |
| chr2R:22933851:22934701: | CG3788 | 0 |
| chr2R:16585351:16586101: | RpLP2 | 28 |
| chrX:10473901:10474701: | nocte | 0 |
| chr2L:15498951:15499951: | lace | 0 |
| chrX:4693951:4694551: | ctp | 6152 |

| | | |
|---|---|---|
| chr3L:20695151:20696001: | kni | 0 |
| chr2R:19275201:19276001: | rib | 5124 |
| chrX:17834001:17834501: | CG6398 | 3174 |
| chr3R:8669001:8669751: | p | 933 |
| chrX:17159951:17160951: | baz | 0 |
| chr2R:13829051:13829651: | CG6329 | 2681 |
| chr2L:7984051:7984651: | Snoo | 0 |
| chrX:9684101:9687151: | nej | 0 |
| chr3L:14634101:14635001: | dlp | 1501 |
| chr3R:16249901:16250851: | bor | 13264 |
| chr3R:9054151:9054901: | Kdm2 | 1657 |
| chrX:3694151:3696151: | Mnt | 22351 |
| chr2R:8064801:8065801: | CG12769 | 0 |
| chrX:16780201:16780751: | if | 2683 |
| chr2R:13035051:13035751: | CG13323 | 11505 |
| chr3R:30450151:30450751: | Osi23 | 5106 |
| chr3L:20784451:20785751: | CG11399 | 446 |
| chrX:11154251:11155401: | Dlic | 0 |
| chr3L:15514251:15514901: | CG13454 | 2155 |
| chr3R:28634301:28634951: | CG10011 | 148 |
| chr3L:5664351:5665101: | CG46320 | 231 |
| chrX:514351:514751: | arg | 4917 |
| chr2R:21364351:21364901: | CG17974 | 13774 |
| chr2L:15264401:15265051: | yuri | 0 |
| chr2L:11804401:11805201: | crol | 4237 |
| chrX:8050051:8050551: | fs | 10885 |
| chrX:10209451:10210151: | CG15309 | 2223 |
| chrX:7969451:7970001: | sws | 1216 |
| chr2R:8648751:8650501: | ptc | 0 |
| chr2R:14970051:14970501: | ckn | 6760 |
| chr3R:28339501:28340101: | Gfat2 | 0 |
| chrX:3679851:3680451: | Mnt | 8051 |
| chrX:12649601:12650351: | CkIalpha | 3275 |
| chr3R:28324751:28325351: | larp | 11089 |
| chrX:18864301:18865351: | CG34401 | 352 |
| chr3R:30794651:30795101: | wts | 11518 |
| chr2R:7779401:7780251: | CG45093 | 2386 |
| chr3R:9239751:9240301: | D1 | 353 |
| chr3L:12804051:12805201: | Atg1 | 0 |
| chr2L:5879201:5880201: | rau | 0 |
| chrX:18634651:18635201: | wgn | 0 |
| chr2R:12019601:12020201: | otk | 0 |
| chr2R:19423451:19425201: | CalpA | 0 |
| chrX:8674651:8675201: | CG12772 | 1862 |
| chr2L:2454251:2455101: | dpp | 25880 |

| | | |
|---|---|---|
| chr3L:15329251:15330101: | Toll-6 | 6591 |
| chrX:11494901:11495551: | Drak | 1767 |
| chr3R:25049901:25050451: | CG11791 | 286 |
| chr3R:9394251:9395051: | FER | 0 |
| chr3R:13074501:13075051: | CG43063 | 11173 |
| chrX:12664951:12665651: | Tis11 | 5537 |
| chr2L:21104951:21105551: | CG46307 | 384 |
| chrX:19199951:19200351: | CG7990 | 1628 |
| chr3L:14205051:14205551: | saturn | 1729 |
| chr2R:16210051:16210901: | Shark | 7168 |
| chr2L:14234051:14234901: | Dyrk2 | 0 |
| chr3L:16410101:16412051: | fax | 0 |
| chr3L:19559301:19559851: | CG9300 | 6882 |
| chrX:2333851:2334851: | Ilp6 | 0 |
| chr2R:8724151:8724801: | gcl | 0 |
| chrX:5744201:5744801: | IntS6 | 2453 |
| chr3L:14785251:14785851: | bmm | 561 |
| chr2R:21545251:21547351: | CG30283 | 4679 |
| chr2L:11970251:11971651: | CG16965 | 9940 |
| chr3L:5665301:5665901: | CG46320 | 1181 |
| chr3L:6964101:6964651: | sgl | 107 |
| chr2L:7530351:7531351: | Obp28a | 32987 |
| chr3R:12369101:12369601: | GstD1 | 0 |
| chr2R:24743901:24744601: | Usp15-31 | 153 |
| chr3L:20620401:20622501: | knrl | 0 |
| chr3R:19165451:19166251: | sqz | 1241 |
| chrX:19753951:19754501: | Pmp70 | 3182 |
| chrX:4678001:4679501: | fzr | 36 |
| chr2R:11592151:11594451: | tou | 21594 |
| chr2R:9278651:9279451: | Pkn | 5646 |
| chrX:12753801:12754451: | ade5 | 6166 |
| chr3R:30465551:30466301: | PH4alphaEFB | 0 |
| chr3L:16645551:16646051: | Abl | 1831 |
| chr2L:13165551:13166151: | DnaJ-H | 0 |
| chr2R:10100551:10101951: | 14-3-3zeta | 875 |
| chr3R:23248751:23249401: | CG4467 | 0 |
| chr2R:16583151:16584401: | RpLP2 | 1728 |
| chr3R:11948851:11949351: | Cad87A | 0 |
| chr2R:8665651:8666601: | Acsl | 0 |
| chr2R:20835701:20836801: | sktl | 0 |
| chr2R:10798451:10799201: | CG33144 | 0 |
| chr3L:8978601:8979201: | CG5644 | 679 |
| chrX:2635851:2636401: | sgg | 1900 |
| chr3L:3225851:3226701: | CG11505 | 1014 |
| chrX:17650851:17652751: | RhoGAPp190 | 0 |

| | | |
|---|---|---|
| chrX:5417751:5419051: | spoon | 1816 |
| chr3L:10225951:10226401: | scramb1 | 712 |
| chr3L:18846001:18846651: | Indy | 0 |
| chr3L:18621101:18621651: | not | 478 |
| chr2L:19157851:19158851: | l | 22619 |
| chr3L:19081151:19081901: | Mkp3 | 4525 |
| chr3R:25588201:25588801: | CG4582 | 5023 |
| chrX:3676501:3678801: | Mnt | 4701 |
| chr2L:14233201:14233751: | Dyrk2 | 375 |
| chrX:21033201:21033751: | CG1529 | 1252 |
| chr3L:20401251:20402151: | trbl | 0 |
| chr2L:11806251:11809601: | crol | 0 |
| chr3R:9056301:9058251: | Kdm2 | 3807 |
| chrX:10211351:10212351: | CG15309 | 23 |
| chrX:12666351:12666751: | Tis11 | 6937 |
| chr3L:18148051:18148551: | CG7330 | 372 |
| chr3L:17972251:17973501: | CG5290 | 59687 |
| chr3R:11351501:11353801: | KP78a | 0 |
| chr3L:20507901:20508451: | CG5078 | 1870 |
| chr2R:21056551:21057151: | shg | 46 |
| chr3R:19321601:19322651: | Dl | 3566 |
| chr2R:8647201:8648351: | ptc | 1298 |
| chr2R:18156651:18157501: | lolal | 691 |
| chrX:5661951:5663301: | CG42265 | 0 |
| chr3R:16247801:16248301: | bor | 15814 |
| chrX:3696701:3697351: | Rala | 22284 |
| chr2L:5546801:5547501: | Oscillin | 0 |
| chrX:2636801:2637501: | sgg | 2850 |
| chr2L:3631801:3633201: | CG15418 | 10098 |
| chrX:12006601:12008151: | CG1806 | 0 |
| chrX:10741851:10742801: | ras | 1701 |
| chrX:13746851:13747851: | Grip91 | 2169 |
| chrX:3332451:3333101: | CG10793 | 16050 |
| chr3R:17716901:17718001: | osa | 349 |
| chr3R:19147351:19148051: | Ppcs | 0 |
| chrX:3141951:3142651: | N | 7082 |
| chr2R:17672501:17673001: | P32 | 0 |
| chr3L:18167001:18167501: | CG13699 | 4480 |
| chr3L:9461651:9462951: | CG44838 | 602 |
| chr3R:22666351:22667901: | Gclm | 18609 |
| chr3R:18396651:18397851: | Vti1b | 0 |
| chr2R:8667151:8669601: | Acsl | 1339 |
| chr2R:10432151:10432851: | RanBPM | 2444 |
| chr3R:9512151:9513151: | Ras85D | 0 |
| chr2L:2862151:2862801: | CG2991 | 4988 |

| | | |
|---|---|---|
| chr2R:8232101:8232701: | dpn | 0 |
| chr3L:1577301:1577851: | CG13917 | 603 |
| chrX:6656851:6657601: | Cdc7 | 2311 |
| chr3L:13227451:13229051: | caps | 0 |
| chr2L:17382501:17383251: | Lrch | 864 |
| chr3L:22407501:22408651: | Ten-m | 0 |
| chr2L:11066751:11067451: | Samuel | 0 |
| chrX:12750201:12752451: | CG4004 | 5990 |
| chr3L:11832551:11833351: | CycA | 227 |
| chr3L:1882551:1883601: | CG13937 | 0 |
| chrX:15712601:15713451: | CG8184 | 0 |
| chr3R:26036251:26037351: | E | 0 |
| chr3L:21276651:21277301: | AcCoAS | 2942 |
| chr2R:19292701:19293551: | Tab2 | 173 |
| chr3L:22727701:22728401: | CG11367 | 0 |
| chr3R:6966551:6967251: | Antp | 31977 |
| chr3R:25207751:25208701: | Cad96Ca | 0 |
| chr2L:12421701:12422201: | vir-1 | 1238 |
| chrX:17291251:17292201: | CG8661 | 6752 |
| chrX:19850651:19852151: | pico | 0 |
| chr3R:14737851:14739501: | eff | 1818 |
| chr3L:1861251:1862051: | Rap1 | 188 |
| chrX:16091351:16092051: | Tob | 0 |
| chr3L:2466451:2467051: | CG16762 | 8563 |
| chr3R:10132951:10134001: | tws | 5771 |
| chr3R:24031151:24032001: | KrT95D | 0 |
| chrX:15076001:15077001: | Lsd-2 | 0 |
| chr2R:8083051:8083801: | slv | 0 |
| chrX:9666351:9666901: | CG3106 | 3449 |
| chr2R:17851251:17851901: | CG30323 | 1394 |
| chrX:15143101:15143901: | RpL37a | 3735 |
| chr2R:7488151:7490901: | wech | 0 |
| chrX:16798151:16798651: | CG9132 | 2059 |
| chr2R:20110551:20111801: | 18w | 0 |
| chrX:17178201:17179151: | baz | 18196 |
| chr2R:8060301:8061751: | CG12769 | 3803 |
| chr3R:15973251:15974351: | Hel89B | 0 |
| chr2L:7418251:7419101: | chm | 5835 |
| chr3R:8105801:8106701: | puc | 470 |
| chr3L:17563301:17564351: | CycT | 5514 |
| chr3L:4263401:4263851: | CG1273 | 0 |
| chr3L:17971051:17971551: | CG5290 | 58487 |
| chr2L:2450551:2451501: | dpp | 22180 |
| chrX:6266051:6266451: | schlank | 777 |
| chr3R:15665651:15666451: | pxb | 0 |

| | | |
|---|---|---|
| chr3R:29214101:29216301: | eEF1gamma | 3871 |
| chrX:19750851:19751301: | Pmp70 | 82 |
| chr3R:31058701:31060051: | dco | 1159 |
| chrX:21053751:21054901: | bves | 0 |
| chrX:3845201:3846201: | CG2901 | 17005 |
| chr2L:14508801:14509351: | CG33648 | 14961 |
| chr3L:15815351:15816151: | fwe | 259 |
| chr2R:19668851:19669951: | hrg | 0 |
| chr2R:23658851:23659801: | Pde8 | 1285 |
| chrX:6533901:6534651: | dx | 0 |
| chr2R:16579201:16581051: | Cdk4 | 0 |
| chrX:2125451:2126001: | ph-d | 2166 |
| chr3R:9220201:9221001: | pum | 16681 |
| chr3R:24630001:24630851: | Wsck | 0 |
| chr3L:7225201:7225851: | CG14826 | 8661 |
| chr2R:14964201:14965851: | ckn | 910 |
| chr2R:13449151:13450201: | cnn | 274 |
| chr2R:15329151:15329651: | CG8160 | 881 |
| chrX:5094201:5094851: | rg | 8443 |
| chrX:1899301:1899801: | arm | 845 |
| chr3R:10035151:10035601: | Mical | 7294 |
| chrX:2254851:2255601: | CG3091 | 655 |
| chr2R:24389251:24390601: | mAChR-A | 0 |
| chr3L:1349451:1349901: | 312 | 1215 |
| chr3L:18439651:18440451: | skl | 0 |
| chr3R:11779551:11780951: | l | 424 |
| chr3R:8333651:8335351: | Poxm | 0 |
| chr3R:15664551:15665351: | pxb | 832 |
| chr2L:8534651:8536201: | CG17834 | 5511 |
| chr3R:25183301:25185301: | dan | 0 |
| chr3R:11604401:11605301: | CG6791 | 14256 |
| chr2L:19154551:19155301: | l | 19319 |
| chrX:13624701:13625851: | NFAT | 0 |
| chr3L:7344751:7345501: | Cln7 | 0 |
| chr2L:3789451:3790151: | bark | 2285 |
| chr3L:2489851:2490401: | CG1146 | 1048 |
| chr3R:10108951:10110001: | Fmr1 | 120 |
| chr3L:10660051:10660701: | CG32066 | 312 |
| chrX:9544301:9544901: | CG32700 | 5937 |
| chr2R:8670101:8670951: | Acsl | 4289 |
| chr3L:1885101:1885951: | Tmhs | 0 |
| chrX:12470151:12470751: | CG42258 | 6624 |
| chr3R:14714051:14714801: | kibra | 9460 |
| chrX:16089151:16089751: | Tob | 1744 |
| chr2L:6339001:6339751: | Cpr | 166 |

| | | |
|---|---|---|
| chr3L:17995251:17995901: | CG32192 | 44416 |
| chr3R:7973951:7974601: | CD98hc | 3827 |
| chr2R:10237951:10239601: | Hdc | 0 |
| chr2R:21645401:21647051: | CG10082 | 0 |
| chr3R:9033551:9034551: | hyx | 4209 |
| chr3R:22898451:22899551: | klg | 0 |
| chr3L:17245501:17246101: | CG7707 | 2066 |
| chr3R:15280501:15281651: | Tm1 | 0 |
| chr3L:2228301:2229451: | Oseg2 | 195 |
| chr2L:19820551:19821001: | CG13962 | 6020 |
| chrX:8041001:8044401: | CG2233 | 3485 |
| chrX:19505651:19506351: | Pfrx | 10147 |
| chrX:8420651:8421751: | lawc | 8994 |
| chr3R:18670701:18671351: | CG31230 | 328 |
| chr3R:24218501:24219251: | CG17786 | 0 |
| chr3L:17968451:17969251: | CG5290 | 55887 |
| chr2L:2885751:2887401: | lilli | 0 |
| chr3L:6985801:6986251: | CG43439 | 1216 |
| chr3L:7910801:7911301: | pbl | 653 |
| chr2L:430801:431401: | ex | 0 |
| chr2R:11615801:11616551: | tou | 0 |
| chr2L:16718451:16719151: | Cyt-c-p | 781 |
| chr3R:19325951:19326951: | Dl | 0 |
| chrX:2122751:2124001: | ph-d | 0 |
| chr2R:22678201:22678951: | RYBP | 7615 |
| chr3L:19086051:19087501: | Mkp3 | 0 |
| chr2L:8416051:8416901: | Akap200 | 362 |
| chr2L:15478051:15478901: | sna | 0 |
| chr3R:6893151:6893901: | ftz | 28828 |
| chr2R:18673051:18673901: | edl | 0 |
| chrX:521251:524001: | elav | 0 |
| chr2L:7991251:7992251: | pes | 1933 |
| chrX:12471251:12472951: | CG42258 | 7724 |
| chr3R:7973201:7973701: | CD98hc | 4727 |
| chr2R:20587451:20588701: | HnRNP-K | 0 |
| chr3R:29876301:29877351: | CG31038 | 0 |
| chr2R:10078001:10078601: | egr | 0 |
| chr2R:9566401:9567551: | CG1809 | 4764 |
| chrX:21056501:21057101: | bves | 2539 |
| chrX:20411501:20412251: | CG15459 | 1330 |
| chr3R:22041501:22043201: | how | 0 |
| chr3L:15588001:15588451: | CG7656 | 0 |
| chr2L:11516551:11517101: | Dlg5 | 0 |
| chr3R:25996651:25997651: | E | 0 |
| chr2R:10551651:10552451: | psq | 5437 |

| | | |
|---|---|---|
| chrX:11816701:11819151: | rudhira | 0 |
| chr3L:18822551:18823251: | Cat | 0 |
| chrX:18297901:18298251: | upd1 | 11040 |
| chrX:7307301:7308201: | brk | 0 |
| chr3R:25472001:25473201: | Fur1 | 0 |
| chr2R:16187201:16188201: | clu | 618 |
| chrX:7336801:7337551: | CBP | 0 |
| chr3L:19894951:19898151: | Su | 11259 |
| chr2R:14962101:14963151: | aPKC | 0 |
| chr3L:9137401:9138101: | nwk | 0 |
| chr3L:12146901:12147351: | Nrx-IV | 383 |
| chr3R:24092501:24093051: | Syx1A | 12712 |
| chr3R:16242151:16243051: | tara | 16501 |
| chr3R:21186601:21187951: | Snmp1 | 18397 |
| chr3R:9247051:9247601: | CG8420 | 403 |
| chrX:6997051:6999051: | Inx2 | 0 |
| chr3R:25637251:25637901: | jigr1 | 0 |
| chrX:16087351:16087901: | Tob | 3594 |
| chr2R:5782101:5783101: | ZnT41F | 0 |
| chr3L:13476901:13477801: | stv | 0 |
| chr2L:10967101:10967801: | CG33129 | 2474 |
| chr2R:7492201:7492651: | Coop | 0 |
| chr3L:19292201:19293651: | Gbs-76A | 0 |
| chr2L:4997251:4998051: | Rtnl1 | 11669 |
| chrX:2642251:2643151: | sgg | 8300 |
| chrX:3671901:3672701: | Mnt | 101 |
| chrX:17812101:17812701: | CG12986 | 6984 |
| chrX:3451051:3452701: | CG12535 | 2109 |
| chr2R:21522051:21522651: | Egfr | 0 |
| chrX:17657351:17659501: | beta-Spec | 0 |
| chr3R:31222401:31222951: | CG12054 | 1700 |
| chr2L:13512401:13514651: | CG33640 | 13190 |
| chr3R:9417401:9419901: | ps | 0 |
| chrX:11787101:11787551: | Karl | 0 |
| chr3R:21641001:21642451: | E2f1 | 18420 |
| chrX:9692551:9693201: | btd | 985 |
| chrX:13412601:13413401: | Set2 | 0 |
| chr3L:11196801:11197351: | GlcAT-P | 185 |
| chr2R:17027651:17028751: | RhoGEF2 | 462 |
| chr2R:16607651:16608201: | CG5065 | 0 |
| chr2R:22911451:22912301: | CG30187 | 6489 |
| chr2R:10437701:10438551: | Galphao | 0 |
| chr3L:11707701:11708351: | CG11658 | 202 |
| chr2R:13940751:13942251: | shot | 0 |
| chr2R:24986651:24987251: | emp | 582 |

| | | |
|---|---|---|
| chr2L:2887801:2889701: | lilli | 1873 |
| chr2R:24736451:24737151: | Mid1 | 0 |
| chr3R:7102851:7103951: | e | 8917 |
| chr2R:20106101:20107101: | 18w | 4410 |
| chr2R:24767901:24768601: | GstE12 | 0 |
| chrX:7305951:7307051: | brk | 888 |
| chr3R:18922951:18924751: | Mpc1 | 3908 |
| chr2L:9447951:9450151: | CG33723 | 7212 |
| chr2L:3606301:3607001: | odd | 0 |
| chr3R:30803001:30804301: | wts | 2318 |
| chr3R:20753001:20753651: | Syp | 3078 |
| chr3L:14968051:14969701: | Tom | 0 |
| chr2R:24543051:24543901: | ITP | 5631 |
| chr2L:19426251:19426901: | Pax | 0 |
| chr3L:21843101:21843901: | CG14563 | 716 |
| chr3R:18573101:18574201: | fray | 11 |
| chr2L:16715551:16716851: | CG31808 | 0 |
| chr3R:16240951:16241801: | tara | 15301 |
| chr3L:22851101:22851801: | CG32462 | 1711 |
| chr2R:20960601:20961751: | hbn | 0 |
| chr3L:6495751:6496701: | sfl | 72 |
| chrX:15593351:15594501: | CG9220 | 0 |
| chr2R:17458351:17459001: | CG14478 | 689 |
| chrX:2283401:2283901: | Vml | 7580 |
| chr2R:14768401:14769451: | Spred | 0 |
| chr2L:7993401:7994901: | pes | 0 |
| chr2L:3538451:3540401: | drm | 0 |
| chrX:9693501:9694401: | btd | 0 |
| chrX:16085801:16086451: | CG8958 | 2570 |
| chrX:2325501:2326401: | CG14050 | 0 |
| chrX:12673601:12674451: | Tis11 | 14187 |
| chrX:17733651:17734201: | CG42684 | 23508 |
| chr3R:14020451:14021301: | Cyp6d5 | 7927 |
| chr3R:29063701:29065251: | CG11873 | 662 |
| chrX:12640551:12641251: | Tomosyn | 8947 |
| chr3R:31724501:31726251: | ttk | 10635 |
| chr2L:4998751:4999301: | Rtnl1 | 10419 |
| chr2R:10553751:10554551: | NA-psq | 3337 |
| chrX:9163901:9164451: | mei-P26 | 647 |
| chr3R:22368901:22370151: | PyK | 1366 |
| chr3R:22043951:22044401: | how | 1761 |
| chr3L:19929001:19929651: | RhoGDI | 259 |
| chr3R:15250251:15250801: | CG42404 | 158 |
| chr3L:17999251:18000251: | CG32192 | 40066 |
| chr3R:21640051:21640701: | E2f1 | 20170 |

| | | |
|---|---|---|
| chr3L:16845101:16845651: | Zcchc7 | 0 |
| chrX:5899351:5900951: | Act5C | 0 |
| chr3L:11084401:11085101: | JIL-1 | 8092 |
| chr3L:14754701:14760551: | CG42507 | 0 |
| chrX:7944751:7945551: | l | 1357 |
| chr3R:13814351:13815551: | tal-1A | 1243 |
| chr3R:18274851:18275551: | eIF1A | 187 |
| chr3R:16274451:16275151: | gish | 2000 |
| chr3L:10664451:10665851: | simj | 0 |
| chr3L:17964651:17965451: | CG5290 | 52087 |
| chrX:7709251:7710451: | CHES-1-like | 0 |
| chrX:369551:370251: | ac | 0 |
| chr2R:18309551:18310151: | Tango8 | 1671 |
| chr3L:15619551:15620301: | CG7372 | 0 |
| chr3L:19294651:19295351: | fal | 88 |
| chr3L:9004701:9005401: | Doc3 | 0 |
| chr2R:13939501:13940251: | shot | 1859 |
| chr2L:8419751:8420651: | Akap200 | 4062 |
| chr3R:10139751:10140501: | tws | 0 |
| chrX:1209251:1210101: | CG11382 | 0 |
| chr3L:17569901:17570401: | CycT | 12114 |
| chr3R:23334301:23335001: | pnt | 11166 |
| chr3L:13904251:13905001: | dysc | 1016 |
| chr2L:9874451:9875001: | nAChRalpha6 | 11249 |
| chr2L:14688901:14689951: | osp | 0 |
| chr3L:3060051:3060851: | CG16753 | 4973 |
| chr2R:10555051:10555751: | psq | 2137 |
| chrX:11370101:11370601: | dlg1 | 437 |
| chr3R:14624301:14624851: | put | 852 |
| chr2L:5000251:5001501: | Rtnl1 | 8219 |
| chr3L:18138951:18139701: | geko | 0 |
| chr3R:13063751:13064651: | sim | 5997 |
| chrX:5770401:5771251: | CG3097 | 0 |
| chrX:10823301:10824451: | Imp | 0 |
| chrX:6338701:6339401: | CG3842 | 940 |
| chr2R:14958351:14959401: | aPKC | 3568 |
| chr3R:30805601:30806251: | wts | 368 |
| chrX:2323801:2324301: | CG14050 | 1916 |
| chr2L:19423301:19424301: | CG16771 | 0 |
| chrX:2155751:2156451: | wapl | 2532 |
| chr3R:8680751:8680951: | CG33325 | 718 |
| chr3R:14055851:14056401: | foxo | 544 |
| chr3R:15285851:15287301: | CG45218 | 3498 |
| chr3L:1463501:1464101: | rho | 0 |
| chr3L:4233601:4234101: | ImpL2 | 2046 |

| | | |
|---|---|---|
| chr3R:21638251:21639101: | InR | 18931 |
| chr2R:16220951:16221551: | mrj | 242 |
| chr3R:13976001:13976601: | Orc2 | 11117 |
| chrX:12596001:12597001: | Sec16 | 0 |
| chr3L:1586201:1587051: | CG13917 | 9503 |
| chr2R:20996201:20998851: | CG10543 | 0 |
| chr2L:8277401:8278751: | CG8086 | 19900 |
| chr2R:24891251:24891651: | NaCP60E | 0 |
| chrX:8271251:8271851: | Corp | 0 |
| chr2L:3476351:3477001: | Thor | 1433 |
| chr3L:2896351:2896801: | Shab | 1335 |
| chr2L:4301351:4301901: | tutl | 18204 |
| chrX:5636351:5638101: | CG42492 | 10386 |
| chr3L:10666351:10667051: | simj | 1613 |
| chr3L:21807901:21808601: | eg | 102 |
| chr3L:14177251:14178551: | D | 69 |
| chrX:5901451:5902401: | Act5C | 591 |
| chr2L:8541501:8544651: | Sema1a | 0 |
| chr3R:13812451:13813451: | tal-1A | 0 |
| chr2R:23666551:23667351: | Pde8 | 8985 |
| chr3R:31621601:31622401: | kek6 | 0 |
| chr3R:14056601:14058001: | foxo | 0 |
| chr3L:3336651:3337301: | kst | 0 |
| chrX:18131651:18132151: | CG43133 | 1474 |
| chr3R:14017301:14018301: | Cyp6d5 | 10927 |
| chr3L:13236801:13237351: | caps | 8110 |
| chrX:12437651:12438151: | Pkcdelta | 5428 |
| chr2R:20582001:20583101: | qsm | 490 |
| chr3L:13667051:13668101: | bru3 | 135299 |
| chr2L:5906901:5907801: | bchs | 0 |
| chr3L:18867401:18868051: | Dysb | 0 |
| chrX:10822001:10823051: | Imp | 1226 |
| chr3L:492351:493001: | CG34267 | 7450 |
| chr2R:14332001:14332651: | CG8613 | 19580 |
| chr3R:11977051:11977501: | CG10005 | 0 |
| chr2R:17462051:17462701: | CG14478 | 4389 |
| chr2L:5997101:5998301: | lid | 1200 |
| chr2L:867151:868801: | aru | 0 |
| chr2R:21497151:21497901: | Fkbp14 | 28 |
| chr3L:14972201:14972801: | Brd | 0 |
| chr3R:21637051:21637651: | InR | 17731 |
| chr2L:8162351:8163351: | Piezo | 0 |
| chr3L:12832351:12832851: | CG10960 | 10255 |
| chrX:7704551:7707601: | CHES-1-like | 1784 |
| chrX:17396951:17397601: | B-H1 | 0 |

| | | |
|---|---|---|
| chr3R:7107401:7110351: | Mlp84B | 7381 |
| chr3R:16071901:16072501: | CG10185 | 4308 |
| chrX:16967501:16968001: | Ubr1 | 678 |
| chr3R:16277551:16278101: | gish | 5100 |
| chrX:11372601:11373301: | dlg1 | 2937 |
| chrX:1991651:1992351: | PIG-K | 17013 |
| chr3L:11046351:11047201: | CG43693 | 4170 |
| chr3L:1546201:1547201: | Pcyt1 | 110 |
| chrX:19782851:19783501: | zld | 47 |
| chr2R:25061501:25062101: | gsb | 0 |
| chrX:17737901:17739501: | CG42684 | 27758 |
| chr2L:826051:827051: | dock | 83 |
| chr2L:3477951:3478751: | Thor | 0 |
| chrX:11822951:11823651: | rudhira | 6054 |
| chrX:15483051:15483601: | CG6340 | 678 |
| chr3R:17055951:17056901: | Dad | 1945 |
| chr2R:19411051:19411851: | hts | 13052 |
| chr3L:9373301:9374251: | Hsp22 | 371 |
| chr2R:23978351:23978951: | Sox14 | 0 |
| chr2R:10558501:10559201: | NA-psq | 614 |
| chr3L:16043551:16044201: | Diap1 | 6833 |
| chr2L:128601:129651: | CG3164 | 1140 |
| chrX:3835301:3836351: | CG2901 | 7105 |
| chr3R:14748651:14749851: | smp-30 | 1395 |
| chr2L:3888651:3889151: | capu | 13709 |
| chr3R:11568701:11569551: | Mrp4 | 0 |
| chr3R:23278701:23279701: | orb | 1155 |
| chrX:7299551:7301251: | brk | 6688 |
| chr2R:9978751:9979551: | eve | 0 |
| chr3R:30765301:30766201: | zfh1 | 0 |
| chrX:6543851:6544301: | CG34417 | 3081 |
| chrX:18290601:18291101: | upd3 | 13369 |
| chrX:9578951:9579401: | LPCAT | 586 |
| chr3R:6998951:6999801: | Antp | 0 |
| chr2R:16129051:16130701: | spin | 4071 |
| chr3L:15235151:15235901: | Tollo | 0 |
| chr2R:8174101:8175151: | LRP1 | 0 |
| chrX:5904151:5904901: | CG4020 | 845 |
| chr3R:29069151:29070301: | CG11873 | 6112 |
| chr2R:11175201:11175601: | CG13229 | 2996 |
| chr3R:15094401:15095701: | CG6966 | 0 |
| chr2R:10559451:10560201: | NA-psq | 1564 |
| chr3L:14219501:14220151: | CG7768 | 1298 |
| chr3R:13364101:13365451: | Dic1 | 0 |
| chrX:10224551:10225201: | CG34408 | 4442 |

| | | |
|---|---|---|
| chrX:15484601:15485101: | Gmap | 1821 |
| chr2L:8424651:8425351: | grk | 8247 |
| chr2L:6939651:6940251: | CG18304 | 2563 |
| chr2R:18514701:18515251: | CG43066 | 128 |
| chr2R:13539551:13540201: | Vmat | 5301 |
| chr3R:16279851:16282001: | gish | 7400 |
| chr2R:25214901:25215601: | Kr | 11010 |
| chrX:20489601:20490051: | CG15461 | 3422 |
| chr3L:16659951:16661651: | Baldspot | 0 |
| chr2R:14954251:14955001: | aPKC | 7968 |
| chr2R:12903951:12904851: | NAT1 | 11143 |
| chr3R:11980151:11980751: | CG18547 | 0 |
| chr3L:19045201:19045851: | nkd | 0 |
| chr2R:11475201:11476101: | inv | 737 |
| chr3R:30763901:30764751: | zfh1 | 1175 |
| chr3L:5633951:5634751: | Blimp-1 | 2969 |
| chr3L:15234151:15234751: | Tollo | 868 |
| chr3L:18005251:18005751: | CG32192 | 34566 |
| chr3L:4430251:4430851: | nAChRbeta1 | 903 |
| chr2L:3825301:3825901: | slp1 | 0 |
| chr3R:17053351:17054551: | Dad | 0 |
| chr3R:7000451:7001451: | Antp | 1224 |
| chr2R:19455501:19457501: | mei-W68 | 0 |
| chr2L:17735501:17736051: | CG43231 | 30821 |
| chr3R:10863851:10864451: | CG6574 | 864 |
| chr2R:16834051:16834451: | CG34460 | 7180 |
| chrX:10865551:10866001: | CG2157 | 3283 |
| chr3R:15290651:15292251: | CG45218 | 0 |
| chr3R:8094001:8094301: | CG7900 | 0 |
| chrX:5593801:5594251: | Vsx1 | 4 |
| chr3L:20493501:20494201: | ZnT77C | 720 |
| chrX:19297851:19299201: | CG8034 | 0 |
| chr3R:4452101:4454101: | CG31522 | 8782 |
| chr2L:7188351:7189051: | CG4496 | 959 |
| chr3R:21632301:21634001: | InR | 12981 |
| chr2R:16031001:16032151: | SP2353 | 0 |
| chr2R:11173351:11173951: | CG13229 | 1146 |
| chr2L:12678051:12678901: | CG15485 | 8338 |
| chr3L:7156101:7157351: | corn | 0 |
| chr2R:11626151:11627101: | Egm | 7129 |
| chr2L:19548051:19548751: | Tep4 | 0 |
| chr2R:6168151:6168751: | EcR | 336 |
| chrX:4507851:4508701: | CG3556 | 21530 |
| chr2R:18171301:18171951: | Dgp-1 | 656 |
| chr2R:10227801:10228651: | CG46321 | 0 |

| | | |
|---|---|---|
| chrX:6642951:6643601: | CG14443 | 8993 |
| chr3R:19656401:19656951: | CG6184 | 915 |
| chr2R:9711401:9712951: | dap | 0 |
| chr3R:18931501:18932201: | CG42613 | 2306 |
| chr3L:16046551:16047751: | Diap1 | 3283 |
| chr2R:22382601:22383401: | CG13506 | 0 |
| chrX:19886601:19887701: | amn | 2359 |
| chrX:13111701:13112551: | CG43313 | 0 |
| chr3R:18196701:18199501: | Ssdp | 94 |
| chr2R:10781651:10783251: | mthl13 | 8208 |
| chr3L:9126901:9128251: | bol | 6154 |
| chrX:2317351:2318101: | CG14050 | 8116 |
| chrX:7937551:7938101: | Ubr3 | 1656 |
| chr2L:5296901:5297401: | vri | 7958 |
| chrX:7702301:7703001: | CHES-1-like | 6384 |
| chr2R:6631201:6632951: | Vha16-1 | 139 |
| chr2R:24727151:24727951: | Mid1 | 8706 |
| chr2R:21502051:21503351: | MESK2 | 0 |
| chr3L:9377151:9377951: | Hsp26 | 0 |
| chr3L:15532301:15532701: | CG43248 | 829 |
| chr3L:4227151:4227601: | ImpL2 | 8546 |
| chr3L:14977401:14978051: | CG3349 | 1569 |
| chrX:6331851:6332551: | CG42240 | 1010 |
| chr2R:19629951:19632551: | sm | 0 |
| chr3L:17577451:17578251: | CycT | 19664 |
| chr2L:15467051:15467501: | sna | 10759 |
| chrX:8657051:8657501: | oc | 6371 |
| chr3R:24767501:24769501: | CG13631 | 0 |
| chr3L:21024651:21027451: | skd | 0 |
| chr3L:6716901:6717451: | ple | 2074 |
| chr3L:1592601:1593551: | CG12104 | 9123 |
| chrX:2292651:2294151: | CG2865 | 0 |
| chr3R:17680301:17682251: | alt | 264 |
| chrX:10816751:10817251: | Imp | 7026 |
| chr2L:4346651:4347151: | CG15429 | 5824 |
| chr3R:6791351:6792101: | Dfd | 0 |
| chr3R:26020751:26022001: | E | 0 |
| chrX:19788001:19789101: | zld | 4454 |
| chr3R:13983051:13984251: | Orc2 | 18167 |
| chr3R:9428051:9428851: | ps | 10112 |
| chr2R:21193101:21193701: | Lapsyn | 0 |
| chr2R:21250201:21251701: | ASPP | 0 |
| chr3R:18583451:18585001: | qin | 6026 |
| chr2R:20511051:20511501: | CG13426 | 1668 |
| chrX:9583501:9586151: | Hex-A | 0 |

| | | |
|---|---|---|
| chrX:9650051:9651401: | Ptpmeg2 | 7744 |
| chrX:8225901:8226401: | CG2147 | 14283 |
| chrX:9283601:9284801: | lz | 0 |
| chrX:11828601:11829101: | CG15741 | 8925 |
| chr2L:2955751:2956351: | Rbp9 | 990 |
| chr3R:29073701:29074701: | CG11873 | 10662 |
| chr2L:6078701:6079551: | Kr-h1 | 3052 |
| chr3R:21630351:21631251: | InR | 11031 |
| chr3L:7263751:7264251: | CG14830 | 0 |
| chr2L:8708751:8709601: | Hnf4 | 10 |
| chr3L:19938801:19939401: | Ac76E | 0 |
| chr3R:29253801:29254251: | stg | 1549 |
| chr3L:5629851:5631151: | Blimp-1 | 0 |
| chr3L:4133851:4134501: | ens | 49 |
| chrX:9173851:9174401: | mei-P26 | 10597 |
| chrX:6330151:6331101: | CG42240 | 0 |
| chr2R:13015401:13016101: | Su | 18901 |
| chrX:10815501:10816101: | Imp | 8176 |
| chr2L:7495201:7496051: | Ziz | 256 |
| chrX:590251:591051: | vnd | 8131 |
| chr3L:14600551:14601051: | HGTX | 1012 |
| chrX:18285651:18286001: | upd3 | 8419 |
| chr3R:20574001:20575001: | TFAM | 2009 |
| chr3L:10674001:10675351: | simj | 9263 |
| chr3R:10614051:10615251: | hth | 24317 |
| chr3L:9124901:9125851: | bol | 8554 |
| chr3L:539151:540551: | klar | 63 |
| chrX:19889151:19889751: | amn | 309 |
| chr3L:4380201:4380801: | DopEcR | 0 |
| chr3R:21094201:21094851: | RhoGAP93B | 2976 |
| chrX:18534201:18534851: | CG15047 | 5519 |
| chrX:15675101:15675751: | CG12708 | 0 |
| chr3R:24259251:24260951: | beta-PheRS | 6573 |
| chr2R:10224751:10225701: | CG46321 | 2256 |
| chr3L:21800301:21800701: | eg | 8002 |
| chr2L:7855101:7855701: | CG14535 | 0 |
| chr3R:12244301:12244851: | Cyp313a3 | 5036 |
| chr3L:13949351:13949901: | CG32137 | 10102 |
| chr3L:15534351:15535701: | CrebA | 0 |
| chrX:2294401:2295001: | Raf | 465 |
| chr3L:12489851:12490501: | CG10660 | 175 |
| chrX:2469501:2470501: | boi | 0 |
| chr2R:15349301:15350301: | unc-5 | 8830 |
| chr2L:12459701:12460451: | CG5418 | 6460 |
| chr3L:3069751:3071001: | CG32486 | 0 |

| | | |
|---|---|---|
| chr2R:9929451:9930201: | Etf-QO | 18559 |
| chr3L:16049901:16051201: | Diap1 | 0 |
| chrX:5339901:5340501: | SK | 0 |
| chr2L:16299301:16300051: | CG17328 | 0 |
| chrX:10174751:10175051: | CG43902 | 16318 |
| chrX:22854951:22855701: | fog | 0 |
| chrX:23044951:23045651: | CG12061 | 8886 |
| chr2L:16519951:16521801: | CG5953 | 11076 |
| chrX:5194551:5195001: | CG5062 | 12183 |
| chr3L:14599451:14599951: | HGTX | 0 |
| chrX:4940101:4940651: | Ptp4E | 974 |
| chr3L:9019101:9019851: | Doc2 | 0 |
| chr3R:21348801:21349851: | CG16791 | 0 |
| chr2R:6220151:6221251: | tomboy40 | 4383 |
| chr3R:29255151:29255701: | stg | 99 |
| chrX:19220151:19221251: | CG7884 | 0 |
| chrX:19604201:19604801: | CG14207 | 398 |
| chr2L:248251:249751: | kis | 1072 |
| chr3R:18585251:18586051: | qin | 4976 |
| chr3L:18640301:18641451: | MYPT-75D | 14343 |
| chr2R:21075301:21076351: | Treh | 1284 |
| chr3R:25515301:25516351: | msi | 0 |
| chrX:10814051:10814651: | Imp | 9626 |
| chr3L:18185351:18187301: | hid | 0 |
| chr3R:27580401:27582051: | wdb | 190 |
| chr2L:3594101:3594551: | odd | 12205 |
| chr2L:22134051:22134551: | CG42748 | 2831 |
| chr3R:26010501:26010951: | E | 0 |
| chrX:2295601:2297801: | Raf | 136 |
| chr2R:15890651:15891351: | CG33463 | 5076 |
| chr2R:21248751:21249301: | ASPP | 1849 |
| chr2R:18413301:18414301: | imd | 0 |
| chrX:9118551:9119251: | AP-1gamma | 9861 |
| chr3R:14065751:14066101: | foxo | 8807 |
| chr3R:31712751:31714201: | ttk | 0 |
| chr3L:10675801:10676401: | simj | 11063 |
| chr2L:12460851:12461851: | CG5418 | 7610 |
| chrX:14218051:14219101: | l | 6595 |
| chrX:22855901:22856451: | fog | 708 |
| chr3L:15535901:15536351: | CrebA | 507 |
| chr3R:23228201:23229001: | cnc | 1491 |
| chr3L:6942751:6943901: | tow | 0 |
| chr3L:21023251:21023901: | skd | 3533 |
| chr2L:9863501:9863901: | nAChRalpha6 | 22349 |
| chr2R:6628101:6628801: | CG33919 | 3525 |

| | | |
|---|---|---|
| chr3L:11096201:11097051: | CG33947 | 0 |
| chr3R:21096351:21097401: | RhoGAP93B | 426 |
| chr3R:29457101:29458601: | CG2321 | 30664 |
| chr3R:20291951:20293551: | CG4367 | 11403 |
| chr3R:24272551:24273501: | Orct2 | 0 |
| chr2L:13877851:13878501: | Smg5 | 26160 |
| chrX:13641501:13642401: | NFAT | 16757 |
| chr3R:10616551:10617601: | hth | 21967 |
| chrX:16607801:16608401: | mbt | 2284 |
| chrX:5052651:5053351: | ovo | 4379 |
| chr3R:20007751:20008351: | Hs6st | 161 |
| chr3L:1366751:1367401: | ru | 3227 |
| chr3L:3757551:3758201: | ppk27 | 2351 |
| chr3L:16051801:16053251: | Mbs | 0 |
| chr3R:23921801:23923001: | sba | 0 |
| chr3R:11371801:11372901: | KP78a | 18671 |
| chr2L:3772751:3773151: | bowl | 1052 |
| chrX:12546101:12548151: | Cpr11B | 4973 |
| chrX:3161851:3163351: | CG18508 | 9789 |
| chr2L:10756901:10757501: | CG17124 | 577 |
| chr3R:30912051:30913001: | Ptx1 | 0 |
| chr3R:10387001:10387501: | cwo | 757 |
| chr2R:7352301:7352901: | Dscam1 | 28998 |
| chr2L:6647101:6647801: | CG31635 | 1313 |
| chrX:18282301:18282851: | upd3 | 5069 |
| chr3L:7866901:7867851: | Pdp1 | 0 |
| chr3R:11312301:11312801: | pros | 15679 |
| chr3R:30532151:30532751: | tmod | 0 |
| chr2R:25052301:25052751: | gsb-n | 0 |
| chrX:7695551:7697701: | CHES-1-like | 11684 |
| chr2R:13592351:13593401: | CG45088 | 0 |
| chr3R:13051251:13052601: | pic | 0 |
| chr2R:8272451:8273201: | CG30371 | 3118 |
| chr2L:246301:247501: | kis | 3322 |
| chr3L:16451901:16452401: | dsx-c73A | 0 |
| chr3R:14696701:14697401: | Meltrin | 0 |
| chrX:3372601:3373451: | Myc | 0 |
| chr3L:11242601:11243351: | Ir68a | 6384 |
| chr2L:19136451:19137251: | l | 1219 |
| chrX:19662751:19663351: | CG14223 | 620 |
| chr3R:23226451:23227201: | cnc | 0 |
| chr2R:13925151:13927201: | shot | 14909 |
| chr3R:9532801:9534101: | by | 0 |
| chrX:15162801:15163401: | CG9095 | 0 |
| chr2L:9581601:9582151: | gcm | 0 |

| | | |
|---|---|---|
| chr3L:542851:544801: | Hipk | 0 |
| chr3R:25517851:25518601: | msi | 1734 |
| chr2L:3771201:3772101: | bowl | 0 |
| chrX:8560601:8562101: | Trf4-1 | 280 |
| chrX:20481501:20482101: | CG15461 | 4079 |
| chr3R:10387901:10388501: | cwo | 0 |
| chr2R:14457901:14458751: | Oaz | 22003 |
| chr3R:21626151:21627051: | InR | 6831 |
| chr3R:29257951:29258551: | CG45544 | 1467 |
| chr3R:21856001:21857001: | glec | 0 |
| chr2L:3836351:3837001: | slp2 | 0 |
| chrX:19353001:19354051: | kek5 | 0 |
| chr2R:13191251:13191951: | seq | 1166 |
| chr3R:17775351:17776951: | TyrR | 12164 |
| chrX:13251451:13251951: | mew | 6539 |
| chr2R:11633051:11633901: | CG9005 | 7040 |
| chrX:15333051:15333801: | HDAC6 | 0 |
| chr3R:10236301:10236851: | CG42795 | 10612 |
| chr2R:9988151:9988851: | TER94 | 308 |
| chr3R:15993151:15993701: | srp | 7000 |
| chr3L:15988201:15988751: | Hip14 | 0 |
| chr3R:28858251:28859701: | Inx3 | 0 |
| chrX:2658301:2658951: | HLH3B | 22772 |
| chr3R:29555851:29556601: | Dr | 0 |
| chrX:3718401:3719801: | Rala | 0 |
| chr2R:12958401:12959301: | CG13321 | 5290 |
| chr3L:20823401:20823901: | Mst77F | 5375 |
| chr3R:6848451:6849201: | Scr | 780 |
| chr3R:29618451:29619351: | CG1907 | 0 |
| chrX:7930101:7931501: | mahe | 0 |
| chr2L:3425851:3426501: | pgant2 | 218 |
| chrX:8650451:8651351: | oc | 0 |
| chr3R:30818651:30819301: | CG15544 | 8518 |
| chr2R:10123651:10125051: | CG46319 | 0 |
| chr3R:11648701:11649251: | Csk | 1455 |
| chrX:2410701:2411251: | CG12496 | 0 |
| chr2R:15895751:15896251: | CG33463 | 176 |
| chrX:10170651:10171251: | CG43902 | 12218 |
| chr3L:16164051:16165101: | CG5151 | 0 |
| chr3L:22835151:22835901: | jim | 0 |
| chr3R:10079101:10079801: | CG5361 | 3729 |
| chr2L:19134701:19135851: | l | 0 |
| chr2R:25114151:25114701: | CG30430 | 14731 |
| chr2R:13189001:13190751: | seq | 0 |
| chr2L:20890151:20890701: | CG14401 | 20298 |

| | | |
|---|---|---|
| chr3L:12434301:12434851: | toe | 1634 |
| chr3R:24894401:24895301: | CG11069 | 0 |
| chr2L:11045051:11045551: | CG18666 | 10257 |
| chr3R:21459801:21460501: | lbe | 13265 |
| chr3R:9534501:9535051: | by | 1491 |
| chr2L:1649551:1650351: | chinmo | 907 |
| chrX:10619601:10620701: | CG43347 | 0 |
| chr2R:23924901:23925301: | CG2812 | 0 |
| chr3L:3249701:3250751: | CG43389 | 1687 |
| chr3L:15924701:15925251: | sff | 0 |
| chrX:8624751:8625301: | Caf1-180 | 18448 |
| chr2R:24369551:24370151: | CG13578 | 7680 |
| chr2R:10219351:10220101: | CG46321 | 7856 |
| chr3R:16224401:16225101: | tara | 550 |
| chr3R:6849901:6850251: | Scr | 0 |
| chr2R:11634951:11635551: | CG9005 | 5390 |
| chr2R:9519001:9520001: | brp | 15237 |
| chr2L:5943401:5945001: | Gpdh | 0 |
| chr3R:30820001:30820501: | CG15544 | 7318 |
| chr3L:9694251:9694901: | CG6767 | 1799 |
| chr2L:14385101:14385651: | ppk | 3930 |
| chr3L:9505101:9506201: | path | 0 |
| chr2L:7245151:7245901: | Ndae1 | 3442 |
| chrX:6700201:6700951: | CG14441 | 694 |
| chr3R:10089151:10089751: | CG5361 | 13779 |
| chr3L:17619201:17619751: | Eip74EF | 0 |
| chr2L:3655251:3658551: | for | 0 |
| chrX:12925301:12926601: | fne | 7825 |
| chrX:9495351:9497801: | mgl | 2328 |
| chrX:2300351:2301801: | Raf | 4886 |
| chr3R:21853501:21854601: | glec | 1665 |
| chr2R:12118851:12119601: | jeb | 0 |
| chrX:3099101:3099601: | CG4116 | 31540 |
| chr2R:13923951:13924601: | shot | 17509 |
| chrX:8090401:8091651: | UbcE2H | 2413 |
| chr2L:8010451:8011051: | Glyat | 0 |
| chr2R:15010501:15011151: | hbs | 245 |
| chr2R:25225651:25227401: | Kr | 0 |
| chr2L:1650701:1651451: | chinmo | 0 |
| chr3R:28860751:28862001: | Inx3 | 1255 |
| chr3R:4360751:4361351: | CG1090 | 869 |
| chr3L:11245801:11246651: | NA-scyl | 5656 |
| chr3R:5700851:5701601: | CG2104 | 7566 |
| chrX:10808501:10809051: | Imp | 15226 |
| chr2L:16525951:16526751: | CG5953 | 6126 |

| | | |
|---|---|---|
| chr2R:17475951:17476751: | insb | 0 |
| chr2R:17716001:17717951: | CG6424 | 1395 |
| chr2R:13991051:13992751: | mam | 0 |
| chr2R:20943251:20943851: | Act57B | 170 |
| chr3L:15223651:15223851: | Tollo | 11768 |
| chrX:16068151:16068801: | dpr18 | 4414 |
| chr3L:9692301:9693801: | CG6767 | 2899 |
| chr2L:5306301:5307001: | CG14024 | 7894 |
| chr3R:31636351:31636801: | kek6 | 14216 |
| chr2R:5141401:5141901: | Atf6 | 2832 |
| chr3R:9436401:9437001: | ps | 18462 |
| chr3L:18016451:18017101: | CG32192 | 23216 |
| chrX:5046401:5048501: | ovo | 0 |
| chrX:16476501:16477001: | eIF4H1 | 27837 |
| chr3L:10681551:10682351: | CG11811 | 8673 |
| chr3R:5086501:5088401: | corto | 0 |
| chr3R:4206601:4207201: | aux | 4582 |
| chr2R:18652601:18653351: | SP2637 | 4871 |
| chrX:7690501:7693251: | CHES-1-like | 16134 |
| chr2R:11231751:11233251: | metro | 0 |
| chr2L:5237001:5238151: | TpnC25D | 20301 |
| chr3R:13011851:13013351: | CtBP | 203 |
| chrX:8091951:8092901: | UbcE2H | 1163 |
| chr3R:8792351:8793001: | Ir85a | 1683 |
| chr2R:19467001:19468701: | mei-W68 | 10832 |
| chr2R:8862251:8862951: | RyR | 1888 |
| chr2R:11637051:11637951: | CG9005 | 2990 |
| chr3R:10622051:10622551: | hth | 17017 |
| chr2L:10457351:10457901: | LManII | 0 |
| chr3L:16807101:16808101: | CG9674 | 0 |
| chr3R:29287251:29287851: | CG14506 | 9329 |
| chr3L:377151:378601: | CG13884 | 10715 |
| chrX:14211151:14212701: | l | 0 |
| chr3L:12892301:12892801: | CG14118 | 0 |
| chr2R:5142301:5142701: | Atf6 | 2032 |
| chr3R:8697301:8698451: | hb | 0 |
| chr2R:15252301:15252851: | scb | 3518 |
| chr2L:12986751:12987651: | Vha68-3 | 7966 |
| chr3R:10087001:10087651: | CG5361 | 11629 |
| chr3L:11247451:11248501: | NA-scyl | 3806 |
| chr2R:21202501:21203201: | Glycogenin | 2019 |
| chr2R:6156101:6157401: | EcR | 11686 |
| chr3R:5716651:5717301: | cas | 0 |
| chr2L:8717751:8718551: | Hnf4 | 8141 |
| chr3R:21621601:21622151: | InR | 2281 |

| | | |
|---|---|---|
| chr3L:4071451:4072151: | wit | 0 |
| chr2R:18406651:18407101: | GstE7 | 0 |
| chrX:3722901:3723501: | Tlk | 2895 |
| chr2R:10056351:10057051: | Def | 1776 |
| chrX:2305601:2306951: | Raf | 10136 |
| chr2L:17336251:17336851: | CG45691 | 15540 |
| chr2L:6088201:6088951: | CG45075 | 2461 |
| chrX:16751151:16751751: | goe | 159 |
| chr3R:23221101:23221751: | cnc | 4960 |
| chr3L:9763251:9764101: | CG8177 | 0 |
| chr2R:9056201:9056651: | CG13743 | 3957 |
| chr3L:14448351:14449101: | CG3919 | 37726 |
| chrX:1975751:1976601: | PIG-K | 32763 |
| chr2R:7418401:7419551: | so | 0 |
| chrX:8213701:8216501: | CG2147 | 24183 |
| chr3R:11391051:11391501: | mRpL40 | 16417 |
| chr3R:17653501:17654351: | CG14322 | 5823 |
| chrX:2303501:2304701: | Raf | 8036 |
| chrX:5918551:5919301: | CG3726 | 1455 |
| chr3R:13683651:13684951: | flfl | 0 |
| chr2R:14715601:14716301: | mspo | 1980 |
| chr2L:18483851:18484951: | CG10283 | 0 |
| chr2R:17718901:17719901: | CG6424 | 0 |
| chr2L:9928901:9929601: | CG42367 | 979 |
| chr2R:22728951:22729801: | jbug | 10658 |
| chr2L:18629001:18629651: | MESR3 | 11746 |
| chr3R:19889801:19890951: | CG4733 | 0 |
| chr3R:18049051:18050401: | htl | 2583 |
| chrX:2049101:2050701: | moody | 0 |
| chr3R:4610351:4610851: | 5-HT2A | 2019 |
| chr2L:7249151:7249951: | Ndae1 | 0 |
| chr3L:10254151:10254751: | Or67c | 9446 |
| chrX:7285351:7285801: | CG2059 | 557 |
| chrX:654251:655951: | Hmt4-20 | 82 |
| chr3R:23220101:23220651: | cnc | 6060 |
| chrX:2664401:2664851: | HLH3B | 16872 |
| chrX:13319451:13320101: | Jafrac1 | 9723 |
| chr2L:21794451:21795801: | CG31612 | 0 |
| chrX:19153201:19155451: | RhoGAP18B | 11631 |
| chr2R:19394901:19395451: | SdhA | 520 |
| chrX:11389551:11390401: | CG15196 | 19546 |
| chr3R:13794351:13795401: | Dip-B | 12064 |
| chr2R:13004901:13005301: | Su | 8401 |
| chr3L:13884751:13885201: | CG13737 | 5758 |
| chr3R:11389601:11390151: | mRpL40 | 17767 |

| | | |
|---|---|---|
| chrX:17788951:17790151: | OdsH | 859 |
| chrX:12929851:12930401: | fne | 12375 |
| chr3L:15104001:15105101: | CG45071 | 781 |
| chr3L:21014301:21015101: | skd | 12333 |
| chr3R:10849201:10850051: | CG42394 | 0 |
| chr3R:16934101:16935051: | Abd-B | 37185 |
| chr2L:13059951:13060751: | CG9932 | 0 |
| chr3R:4609351:4610001: | 5-HT2A | 2869 |
| chr2L:12854401:12855001: | kek1 | 31615 |
| chrX:10240001:10240601: | Ipod | 6922 |
| chrX:5579401:5579951: | Vsx1 | 14304 |
| chr3R:23314001:23314851: | bb8 | 23813 |
| chr3R:23295151:23295951: | bb8 | 4963 |
| chr3R:30600301:30601051: | CG2246 | 438 |
| chr2L:8103951:8104601: | mon2 | 19409 |
| chrX:9638901:9639551: | Ptpmeg2 | 19594 |
| chr2R:24964201:24964551: | CG2765 | 525 |
| chr2R:10213851:10214501: | CG46321 | 13456 |
| chrX:2588651:2589501: | egh | 149 |
| chr3L:17590501:17591201: | Eip74EF | 28124 |
| chr3R:21617951:21619451: | InR | 0 |
| chr3R:31168401:31169451: | stops | 0 |
| chr2R:9230751:9231701: | Cyp4p2 | 5811 |
| chr3R:11655801:11656401: | CG42327 | 0 |
| chrX:19360901:19361651: | kek5 | 7721 |
| chr2R:14713351:14714051: | mspo | 0 |
| chr2L:17750951:17751751: | CadN | 26589 |
| chr3R:15923451:15924001: | CG34276 | 1628 |
| chr3L:3081051:3081651: | CG11486 | 9919 |
| chr3L:12692651:12693901: | mirr | 0 |
| chr2L:18881151:18881751: | tup | 0 |
| chr3R:16933301:16933751: | Abd-B | 38485 |
| chrX:1623101:1623601: | dor | 41451 |
| chr2L:20821401:20822051: | CheB38c | 535 |
| chrX:13126451:13126851: | CG32638 | 1943 |
| chr3R:11302551:11303351: | pros | 25129 |
| chr3L:11251701:11252601: | scyl | 0 |
| chr2R:24712551:24713251: | ST6Gal | 18166 |
| chr2L:377601:378151: | al | 0 |
| chrX:3381901:3382551: | Myc | 8743 |
| chr3R:4807551:4808051: | CG1129 | 3393 |
| chr3R:8027001:8027901: | ImpE3 | 1645 |
| chr3L:18022051:18023251: | CG32192 | 17066 |
| chr2L:3662051:3662801: | Drgx | 0 |
| chr2L:16285651:16287901: | crp | 0 |

| | | |
|---|---|---|
| chr3R:29087151:29089151: | alpha-Man-Ib | 23671 |
| chrX:18832251:18832801: | Pvf1 | 0 |
| chr2L:17752201:17753151: | CadN | 25189 |
| chrX:13322251:13322751: | Jafrac1 | 7073 |
| chrX:10801051:10802601: | sesB | 14094 |
| chr2L:66351:67601: | dbr | 0 |
| chr3L:17611801:17612551: | Eip74EF | 6774 |
| chr2R:5992501:5993101: | Src42A | 10658 |
| chr2R:5726701:5727401: | ap | 124 |
| chr2L:16532651:16533751: | CG5953 | 0 |
| chr3R:4805901:4807251: | CG1129 | 4193 |
| chrX:10546501:10547201: | CG15296 | 19390 |
| chr2R:12886651:12887151: | vg | 2020 |
| chr3L:21010751:21012151: | skd | 15283 |
| chr3R:21616101:21617101: | InR | 2220 |
| chr3L:3827901:3828701: | Rdh | 2453 |
| chr3R:17665801:17667001: | CG7523 | 3894 |
| chr2R:10210701:10211951: | CG34221 | 10991 |
| chr3L:5726351:5726851: | CG44521 | 13021 |
| chr2R:24793201:24793751: | Reg-5 | 0 |
| chr3R:16051101:16051751: | msps | 301 |
| chr3R:15919651:15921701: | Fbxl7 | 0 |
| chr3R:23173401:23174251: | CG46310 | 0 |
| chrX:10628401:10629351: | X11Lbeta | 665 |
| chr2R:12111051:12111551: | jeb | 7789 |
| chr2L:12055651:12056551: | CG34164 | 0 |
| chr3R:9543451:9543951: | mura | 7855 |
| chr2L:3580401:3581401: | sob | 0 |
| chr2R:19178651:19179501: | hppy | 0 |
| chr3R:14853651:14854401: | CG3610 | 8609 |
| chrX:8640801:8641251: | oc | 9430 |
| chr2R:22270351:22271251: | CG5819 | 22149 |
| chr2R:11493801:11494351: | inv | 19337 |
| chr3R:17118851:17119551: | CG34278 | 914 |
| chr2L:16534001:16534601: | CG5953 | 1125 |
| chrX:9515351:9515951: | CG34028 | 3828 |
| chr2L:17754051:17754601: | CadN | 23739 |
| chr3L:7854301:7855901: | Pdp1 | 11468 |
| chr3R:28585251:28585851: | fkh | 0 |
| chrX:12699201:12699951: | CG15725 | 5903 |
| chr3R:11254251:11254951: | CG5214 | 9950 |
| chrX:6709251:6709701: | CG14441 | 9744 |
| chrX:17124801:17125701: | CG5004 | 17220 |
| chrX:9599301:9600301: | Gga | 729 |
| chrX:12720051:12720551: | CG15725 | 14198 |

| | | |
|---|---|---|
| chr3L:21007201:21010401: | CG12984 | 15529 |
| chr2L:16284151:16285351: | crp | 2379 |
| chr3L:18064601:18065301: | Eip75B | 0 |
| chrX:15834351:15835251: | Chc | 239 |
| chr2R:12884301:12885201: | vg | 0 |
| chr3R:13874851:13875601: | E5 | 0 |
| chr2L:16534851:16536301: | CG5953 | 1975 |
| chr2L:19764451:19765101: | fbp | 316 |
| chrX:8100001:8100701: | CG2258 | 1394 |
| chr3L:22910051:22910751: | CG12768 | 0 |
| chr3L:20348201:20349901: | eRF1 | 0 |
| chr3R:9359351:9359851: | CG18473 | 3454 |
| chr3R:23175151:23175701: | CG46310 | 1279 |
| chrX:8548901:8549801: | IntS4 | 8980 |
| chr3R:24699151:24699801: | CG46316 | 12979 |
| chrX:15345201:15345851: | CG6227 | 33 |
| chr2R:14814251:14814701: | CG10202 | 5286 |
| chrX:8295351:8296151: | CG15343 | 21546 |
| chr3L:15098551:15099601: | CTPsyn | 638 |
| chr2R:6235401:6236501: | Pld | 5503 |
| chrX:5560451:5561151: | Vsx2 | 269 |
| chrX:395501:396351: | sc | 0 |
| chrX:9190551:9191151: | CG12115 | 0 |
| chr2R:17638951:17639401: | CG10936 | 0 |
| chr2R:14000601:14001401: | mam | 9399 |
| chr2L:19397851:19399351: | CG17544 | 0 |
| chrX:13130651:13131801: | MFS10 | 1634 |
| chr3R:29818601:29819251: | neo | 0 |
| chr3R:10638651:10639251: | hth | 317 |
| chrX:3175751:3176651: | dnc | 0 |
| chrX:17345801:17346501: | B-H2 | 31221 |
| chr3L:4363251:4364151: | Cip4 | 0 |
| chr2L:9567401:9569101: | CG3838 | 0 |
| chrX:17870951:17871551: | CG6769 | 9950 |
| chr2L:8478301:8478951: | CheA29a | 953 |
| chr3R:4433351:4433951: | eIF3a | 0 |
| chr3R:16403401:16403951: | ss | 0 |
| chr2R:18846051:18846651: | CG15099 | 0 |
| chrX:19718301:19718851: | CG12531 | 72 |
| chr3R:18742501:18743801: | Mekk1 | 0 |
| chr3R:4802301:4803751: | CG1129 | 7693 |
| chr3R:14598051:14598751: | CG14853 | 10891 |
| chr2R:6612851:6613651: | CG33919 | 10926 |
| chr3L:18063151:18063651: | Eip75B | 1045 |
| chr3R:22626451:22627401: | mRpL45 | 7960 |

| | | |
|---|---|---|
| chr3L:7781501:7782201: | Clk | 5899 |
| chr2R:20932951:20933401: | Act57B | 10620 |
| chr3R:21111601:21113301: | Rab11 | 194 |
| chrX:5561601:5562151: | Vsx2 | 1419 |
| chr3L:18792651:18793301: | ftz-f1 | 8008 |
| chr2L:21141701:21142251: | Dap160 | 96 |
| chrX:13237601:13238151: | CG32639 | 1036 |
| chr2L:19361901:19363451: | dnt | 0 |
| chr3R:20782401:20783051: | Takl1 | 9801 |
| chr3R:13877001:13877501: | E5 | 2086 |
| chr3L:22822351:22822951: | jim | 12243 |
| chrX:13657101:13657451: | Lig4 | 10360 |
| chrX:20352301:20352851: | CG1504 | 21930 |
| chrX:10247301:10247851: | Ipod | 14222 |
| chrX:2387451:2388551: | PsGEF | 0 |
| chr2R:15816701:15817501: | Zasp52 | 0 |
| chr2R:12996001:12997501: | Su | 0 |
| chr3L:4061901:4062451: | dib | 0 |
| chr2L:21051901:21052451: | CG42238 | 0 |
| chr2L:5107551:5108751: | Msp300 | 6675 |
| chr2L:1662601:1663601: | chinmo | 11344 |
| chrX:6231701:6232301: | CG14445 | 5014 |
| chr3L:6217901:6218951: | LanA | 0 |
| chrX:12937901:12938751: | hec | 18711 |
| chr3R:21446001:21447001: | lbe | 0 |
| chr2R:18583051:18583801: | GEFmeso | 27976 |
| chr3R:27635951:27636901: | Mtl | 0 |
| chr2L:18795901:18796851: | ham | 2996 |
| chr3L:1919551:1921551: | Dbx | 0 |
| chrX:9305251:9306451: | c12 | 12772 |
| chr3R:4910751:4911401: | Cdep | 200 |
| chr3R:5413601:5414651: | CG14669 | 0 |
| chr2L:12655601:12656301: | pdm2 | 864 |
| chrX:16148851:16149701: | disco-r | 0 |
| chr2R:14798851:14799951: | kn | 7110 |
| chr3R:5263951:5265251: | Tim17a2 | 1260 |
| chrX:13784051:13784601: | inaE | 338 |
| chr2L:9205001:9205901: | tai | 38224 |
| chrX:3294651:3295901: | CG10793 | 20501 |
| chr2R:6609951:6610801: | CG15233 | 8974 |
| chr3R:18739901:18740701: | Mekk1 | 2893 |
| chr3L:18029301:18030001: | CG32192 | 10316 |
| chr2L:16279851:16280601: | crp | 7129 |
| chr3L:6789501:6790351: | vvl | 0 |
| chr2L:12974651:12975451: | Vha68-2 | 330 |

| | | |
|---|---|---|
| chr3R:20594551:20595201: | bon | 1413 |
| chrX:1963701:1965251: | Hr4 | 21762 |
| chrX:20444851:20445351: | CG42578 | 1020 |
| chr2R:15424851:15425401: | CG30472 | 713 |
| chr3R:4599501:4600101: | 5-HT2A | 12769 |
| chr3R:18244351:18245051: | 14-3-3epsilon | 1821 |
| chr3R:27149951:27151001: | CG6051 | 68 |
| chr3R:10170101:10170901: | Best1 | 0 |
| chrX:13234101:13234851: | CG32639 | 1715 |
| chrX:16150301:16151151: | disco-r | 1325 |
| chr3R:9550401:9551901: | mura | 0 |
| chr3R:24565401:24566151: | REPTOR | 16713 |
| chr2R:17813801:17814351: | grh | 12670 |
| chr2R:11650751:11651701: | CG9003 | 0 |
| chr3R:9352201:9354151: | CG43675 | 1344 |
| chr2L:368701:369151: | al | 8961 |
| chr3L:880851:881401: | CG13907 | 612 |
| chrX:1275901:1276851: | Atf3 | 618 |
| chr3L:9400951:9402601: | eIF4E1 | 0 |
| chrX:4411201:4411851: | bi | 1005 |
| chr3L:7381201:7381851: | smid | 0 |
| chr3L:9678001:9678701: | fry | 402 |
| chr2R:14927901:14928601: | Kank | 0 |
| chr3L:12467851:12468551: | eyg | 0 |
| chr3R:21152401:21153501: | cDIP | 7591 |
| chr2R:14006501:14007551: | mam | 15299 |
| chr3L:9422701:9423451: | aay | 0 |
| chr3R:4797501:4798351: | CG17387 | 12780 |
| chr3L:20441651:20442201: | CG5910 | 218 |
| chr3L:18787701:18788301: | ftz-f1 | 13008 |
| chr3L:881701:882451: | CG13907 | 0 |
| chr2L:6667651:6668251: | Galt | 0 |
| chrX:3292601:3293151: | CG10793 | 23251 |
| chr3R:7037001:7038101: | Sodh-1 | 14234 |
| chr3R:5072401:5073001: | corto | 14071 |
| chrX:15352301:15352751: | CG6227 | 7133 |
| chr3R:23722551:23724251: | CG10365 | 0 |
| chr2R:23697601:23698251: | mRpL43 | 0 |
| chrX:19682651:19683851: | Ubqn | 0 |
| chr2L:12482651:12484701: | t-cup | 12632 |
| chr3R:17566601:17567301: | Hmx | 7672 |
| chr3L:20442701:20443451: | CG5910 | 283 |
| chr2R:22136551:22137251: | a | 0 |
| chr3R:18121301:18122251: | CG14316 | 29869 |
| chr3L:7947751:7948401: | exex | 0 |

| | | |
|---|---|---|
| chr3R:21604651:21607201: | InR | 12120 |
| chr3L:12081601:12082151: | Sema5c | 0 |
| chr2R:14007951:14008801: | mam | 16749 |
| chrX:17767951:17768851: | unc-4 | 0 |
| chr2R:14908051:14908651: | Cyp317a1 | 9740 |
| chr3R:18063151:18063701: | htl | 10168 |
| chr2L:21851451:21851801: | tsh | 22859 |
| chr2L:11221151:11221701: | ab | 10474 |
| chrX:17878301:17879101: | CG12985 | 4617 |
| chr3L:11118401:11119201: | FoxK | 1582 |
| chr2R:7885751:7886501: | Dgk | 0 |
| chr3R:5518601:5519151: | Itp-r83A | 1485 |
| chr2R:6605701:6606351: | CG15233 | 4724 |
| chr3R:8190551:8191351: | grn | 9108 |
| chr3R:10535601:10536301: | Cyp12e1 | 100813 |
| chr2L:2798701:2799401: | Syt1 | 578 |
| chrX:10783701:10786401: | Ant2 | 506 |
| chrX:5363901:5364601: | SK | 23666 |
| chr3L:9675451:9676051: | fry | 3052 |
| chrX:13663951:13664451: | Lig4 | 3360 |
| chr2R:10485201:10485901: | pre-lola-G | 0 |
| chr3L:21879151:21881051: | mub | 34364 |
| chr3L:15090101:15090751: | sstn | 3968 |
| chr3R:19875001:19875651: | CG4562 | 27 |
| chr2R:11144901:11145501: | CG13231 | 788 |
| chr3R:25094651:25095051: | fd96Cb | 0 |
| chr2L:12044501:12045301: | CG6785 | 0 |
| chr2R:12979701:12981501: | Psc | 0 |
| chr2R:19228801:19230201: | cora | 0 |
| chrX:9624351:9625201: | CG42797 | 15406 |
| chr3L:2824601:2825151: | pgant6 | 42347 |
| chr2L:13549951:13551201: | kuz | 0 |
| chr2R:9875001:9875501: | CG46338 | 0 |
| chr2R:15600051:15600851: | CG42524 | 13003 |
| chrX:6224301:6224851: | sqh | 289 |
| chr3R:31688951:31689751: | CG11550 | 9072 |
| chr2L:220401:224651: | CG13693 | 7836 |
| chrX:3740451:3741301: | Tlk | 20445 |
| chr3R:16918801:16919501: | Abd-B | 52735 |
| chr2L:16545501:16546151: | CG42389 | 486 |
| chr2L:8735551:8736151: | raw | 4509 |
| chr2L:7827801:7829401: | mts | 152 |
| chr3R:24588701:24589401: | mld | 3049 |
| chr2R:7530651:7531301: | Drat | 1680 |
| chrX:2083451:2084251: | Unc-76 | 2842 |

| | | |
|---|---|---|
| chr2R:8018701:8019201: | CG14762 | 4947 |
| chr3L:4160801:4161551: | nab | 0 |
| chr2L:8030901:8032251: | Cka | 181 |
| chr2L:21150951:21151601: | del | 2406 |
| chrX:19545951:19546951: | CG32532 | 531 |
| chr3L:2823301:2823951: | pgant6 | 41047 |
| chr2L:8088151:8088851: | Bsg | 4732 |
| chr3L:18398151:18398851: | rpr | 0 |
| chrX:15356201:15357001: | Pp1-13C | 4060 |
| chrX:9623351:9623751: | CG42797 | 14406 |
| chr2R:14481251:14482251: | PRAS40 | 65 |
| chr3L:12056251:12056751: | rols | 8108 |
| chr2R:21101351:21102351: | ktub | 0 |
| chr3R:22732801:22733551: | Takl2 | 300 |
| chrX:1941551:1942201: | Hr4 | 0 |
| chr2L:11447001:11448301: | salm | 1388 |
| chr3R:24571701:24573101: | REPTOR | 9763 |
| chr2R:22241751:22244301: | dve | 0 |
| chrX:3741801:3743051: | Tlk | 21795 |
| chr2R:11397101:11398151: | sprt | 0 |
| chr3R:8187651:8188151: | grn | 6208 |
| chrX:16216401:16218051: | disco | 0 |
| chr3R:5612201:5612901: | Rga | 211 |
| chr3R:29377101:29377651: | Ptp99A | 0 |
| chrX:16497151:16497751: | Cnx14D | 27664 |
| chr3R:30617251:30617751: | CG44954 | 4155 |
| chrX:18587201:18587701: | CG15040 | 29888 |
| chr3R:16502451:16502951: | CG42342 | 0 |
| chr2R:10477601:10478751: | whd | 0 |
| chr2R:11657601:11658401: | CG13198 | 2704 |
| chr3R:11156701:11157251: | Ugt86Da | 0 |
| chr3R:13901451:13902101: | ems | 0 |
| chrX:21261651:21262101: | CG1722 | 10504 |
| chr3R:24586351:24587051: | mld | 699 |
| chr2L:16548001:16548701: | CG5968 | 1258 |
| chrX:1943101:1943801: | Hr4 | 1162 |
| chr2L:11445401:11446801: | salm | 0 |
| chrX:15633251:15633901: | sog | 0 |
| chr2R:22256201:22256701: | dve | 12239 |
| chrX:19548401:19548901: | CG32532 | 920 |
| chr3L:2820901:2821551: | pgant6 | 38647 |
| chr3R:23201001:23201451: | fzo | 2583 |
| chr2R:12984401:12986301: | Psc | 3408 |
| chr3L:12913751:12914351: | CG32115 | 106 |
| chr3L:11123801:11124501: | NaPi-III | 454 |

| | | |
|---|---|---|
| chr3L:11010001:11011101: | klu | 814 |
| chr3R:21433951:21434901: | lbl | 0 |
| chr3R:11269051:11269501: | CG5214 | 24750 |
| chrX:3744301:3745201: | Tlk | 24295 |
| chr2L:14409301:14410101: | elB | 0 |
| chr3L:15085001:15085501: | Or71a | 6555 |
| chr3L:11764901:11765501: | CG6024 | 681 |
| chr2R:15734651:15735801: | CG30089 | 7271 |
| chr2R:13619651:13620701: | CG17716 | 2343 |
| chr2L:8739801:8740751: | raw | 0 |
| chrX:6804151:6805101: | pod1 | 468 |
| chrX:19689901:19690651: | CG14227 | 954 |
| chr2L:16268901:16270051: | CG4935 | 6761 |
| chr2R:22859951:22860751: | inaD | 1158 |
| chr2L:8084601:8085001: | Bsg | 1182 |
| chr2R:24813401:24815001: | Dll | 0 |
| chr3R:21829201:21829901: | Gr93a | 2602 |
| chr3L:13629201:13629901: | CG10710 | 109597 |
| chr3L:11008551:11009801: | klu | 0 |
| chr3R:18849101:18849701: | CG18208 | 0 |
| chr2R:17804001:17804651: | grh | 2870 |
| chr2L:5658301:5659551: | DIP-theta | 0 |
| chr2R:21899051:21899501: | PpN58A | 16519 |
| chrX:17885501:17887301: | mnb | 0 |
| chrX:8533801:8534451: | PIP82 | 4104 |
| chrX:15028951:15029401: | be | 4516 |
| chr2R:6131701:6134351: | CG14589 | 2952 |
| chrX:3190651:3191451: | dnc | 14212 |
| chrX:18370701:18371851: | CrebB | 4345 |
| chr3R:17558451:17559251: | Hmx | 0 |
| chrX:18930801:18931401: | CG43759 | 467 |
| chr2L:8083251:8084151: | Bsg | 0 |
| chr3R:21598401:21599151: | InR | 20170 |
| chr2L:9175901:9177251: | tai | 9124 |
| chr2R:10596151:10596651: | CG11883 | 21011 |
| chr2L:18782501:18783751: | ham | 9155 |
| chr2L:16267801:16268601: | CG4935 | 5661 |
| chr2L:7701401:7701951: | Tep2 | 0 |
| chr2R:19373051:19373501: | CG9416 | 201 |
| chr3R:25137451:25138351: | danr | 0 |
| chr2R:18232001:18233201: | fj | 0 |
| chr2R:22867601:22868101: | fd59A | 58 |
| chr3R:17557301:17558101: | Hmx | 829 |
| chr3L:18946901:18947451: | CG32204 | 864 |
| chr2R:24342551:24343051: | bs | 0 |

| | | |
|---|---|---|
| chr2R:13897451:13898051: | CG16935 | 35450 |
| chrX:10262001:10262551: | Hk | 934 |
| chr2R:13501751:13502951: | drk | 0 |
| chr3R:9340651:9342901: | CG45050 | 436 |
| chrX:20982451:20982901: | CG11666 | 1785 |
| chr2R:17800851:17802801: | grh | 0 |
| chr3L:4167201:4167851: | mas | 0 |
| chr2L:4221551:4222751: | ft | 0 |
| chrX:16332351:16333901: | Dsp1 | 0 |
| chr2R:11662401:11663251: | Tret1-1 | 2517 |
| chr3L:2182051:2182551: | CG13810 | 3379 |
| chr2R:13622501:13623351: | CG17716 | 0 |
| chr3R:10826551:10827301: | Cad86C | 0 |
| chr3R:6402701:6403351: | Dmtn | 2544 |
| chr3R:31662751:31663251: | CG1815 | 6214 |
| chrX:1947751:1948651: | Hr4 | 5812 |
| chr3R:8181251:8182201: | grn | 0 |
| chrX:1951501:1952151: | Hr4 | 9562 |
| chr3R:19096501:19097051: | CG6040 | 0 |
| chr2R:20915901:20917051: | Rx | 0 |
| chr3R:11281401:11282051: | CG5214 | 37100 |
| chr3R:17556151:17556751: | Hmx | 2179 |
| chr2R:18229501:18231701: | fj | 1060 |
| chr3L:3963401:3964651: | CG12605 | 120 |
| chr3R:21948501:21949301: | Eip93F | 0 |
| chr3R:30880951:30881451: | CG12071 | 0 |
| chrX:3748601:3749401: | Tlk | 28595 |
| chrX:20693601:20694601: | run | 2932 |
| chrX:4423651:4425501: | bi | 10796 |
| chr3L:12328701:12329251: | Lmx1a | 0 |
| chr2L:478751:481251: | cbt | 0 |
| chr3R:12278801:12279551: | Sfp87B | 5676 |
| chr2R:17335651:17336151: | CG10939 | 58442 |
| chr3R:30099101:30099701: | CG45072 | 23562 |
| chrX:18374451:18375001: | por | 4853 |
| chrX:12395001:12395501: | CG32651 | 12976 |
| chrX:13219951:13220401: | REG | 7627 |
| chr2L:6114951:6115551: | CG31642 | 2961 |
| chr3L:15079101:15079951: | Or71a | 655 |
| chr3R:6405051:6405901: | Dmtn | 0 |
| chr3L:12580051:12580751: | ara | 0 |
| chr2R:18355101:18358051: | sbb | 0 |
| chr2R:20913551:20914801: | Rx | 1952 |
| chr3L:768951:769801: | hng3 | 13463 |
| chr2R:11665251:11665901: | Tret1-1 | 0 |

| | | |
|---|---|---|
| chrX:15023601:15024651: | be | 0 |
| chr3R:16025501:16026351: | pnr | 0 |
| chr3R:12280501:12281351: | Sfp87B | 7376 |
| chr2L:10353751:10354301: | CG34367 | 0 |
| chr2L:17565701:17566251: | CG7094 | 9937 |
| chr2R:15613151:15614051: | CG42524 | 0 |
| chrX:15811451:15813801: | CG8509 | 1897 |
| chrX:4596251:4597051: | peb | 20513 |
| chr2R:11666451:11667051: | Tret1-1 | 684 |
| chrX:4426551:4427451: | bi | 13696 |
| chr3R:30851701:30852551: | tll | 0 |
| chr3R:6406701:6408101: | gpp | 0 |
| chr3R:14346701:14347451: | HEATR2 | 14335 |
| chr3L:17486751:17487901: | Oatp74D | 0 |
| chr3R:18722451:18723001: | CG34283 | 0 |
| chr3R:21557001:21558301: | slou | 0 |
| chr2R:18377301:18377801: | CG43202 | 4043 |
| chr2R:25257251:25257751: | CG9380 | 1934 |
| chr3R:16702401:16704101: | Ubx | 30325 |
| chr3R:13147501:13148101: | 2mit | 7586 |
| chr3L:22132551:22133251: | olf413 | 2839 |
| chr3R:5282651:5283601: | CG31538 | 4817 |
| chr3L:21202701:21203251: | CG33056 | 0 |
| chrX:6601301:6602251: | pigs | 1240 |
| chr3L:18771351:18772201: | Atg3 | 22665 |
| chr2L:8587801:8588251: | Sema1a | 45655 |
| chr2R:23713051:23713951: | Sesn | 0 |
| chr2R:8798051:8798651: | sns | 0 |
| chr3L:14131351:14131901: | Sox21b | 28 |
| chr2R:12261101:12261851: | Cam | 1704 |
| chr2L:12365751:12366601: | CG17010 | 9311 |
| chr3R:19860801:19861501: | Ire1 | 0 |
| chr2R:10025101:10026451: | Pka-R2 | 0 |
| chr2L:15425701:15426451: | wor | 122 |
| chr3L:4624351:4626451: | Src64B | 0 |
| chr2R:23579951:23581351: | apt | 15061 |
| chr3L:445701:446301: | CG13891 | 11796 |
| chrX:9743801:9744701: | Sp1 | 14154 |
| chr2R:6258801:6259401: | Pld | 16798 |
| chr3R:21558851:21559351: | slou | 1670 |
| chr3R:8413851:8414351: | Obp85a | 6246 |
| chr3L:22134001:22136501: | olf413 | 0 |
| chrX:13464201:13464801: | CG34324 | 6539 |
| chrX:17484301:17484851: | CG8568 | 31759 |
| chr2L:3095001:3095551: | PpD6 | 14121 |

| | | |
|---|---|---|
| chr2R:14024451:14025101: | CG18371 | 3519 |
| chrX:7444951:7445501: | CG11368 | 6708 |
| chr3L:7804501:7805851: | CG32369 | 3974 |
| chr2R:10159601:10160351: | mlt | 671 |
| chr3R:21589901:21590351: | CG15498 | 20753 |
| chr3R:14574751:14575301: | CG7987 | 586 |
| chrX:6599301:6600301: | pigs | 0 |
| chr3R:20254601:20255101: | cic | 1832 |
| chr3L:6903751:6905001: | Prat2 | 9875 |
| chr3R:31810001:31810501: | CG11576 | 0 |
| chr3L:3149051:3149801: | BtbVII | 406 |
| chr3R:6410201:6411201: | gpp | 3347 |
| chr3R:19225251:19226701: | vib | 0 |
| chr3L:19635251:19636051: | wnd | 0 |
| chr3R:30386601:30389701: | Fer1HCH | 0 |
| chr2R:16270301:16270801: | Khc | 1170 |
| chr2R:12259051:12259651: | Cam | 0 |
| chrX:13805351:13806301: | CG11095 | 5383 |
| chrX:840451:841401: | CG14635 | 3422 |
| chr3R:8895551:8896201: | pyd | 35695 |
| chr3R:17943501:17944401: | cpo | 23670 |
| chr3R:20252651:20254401: | cic | 0 |
| chrX:12085601:12086401: | CG15734 | 624 |
| chr3R:26797751:26799301: | Tl | 0 |
| chr3R:9865851:9866601: | Teh1 | 1519 |
| chr2L:523051:524001: | lwr | 18579 |
| chr3L:15021001:15021451: | CG32148 | 2041 |
| chr2R:20896101:20896651: | otp | 6763 |
| chr3R:21561151:21562151: | slou | 3970 |
| chr3L:2808051:2808751: | pgant6 | 25797 |
| chr3R:11886901:11888551: | mthl5 | 0 |
| chr2L:16422401:16423501: | Idgf1 | 21777 |
| chr2L:348001:348401: | Ent1 | 9473 |
| chr2L:6786601:6788151: | nrv1 | 0 |
| chrX:686701:687851: | sdk | 0 |
| chr2L:486801:487351: | cbt | 7114 |
| chrX:4617001:4617851: | peb | 0 |
| chr2R:20907151:20907751: | CG9235 | 1953 |
| chr2L:522051:522751: | lwr | 19829 |
| chr3L:8512251:8513101: | CG6765 | 0 |
| chr3L:9546751:9547401: | CG42673 | 0 |
| chr2L:20561351:20562401: | CG44270 | 30286 |
| chrX:18946501:18947301: | CG43759 | 16167 |
| chr3R:23962701:23963351: | CG31140 | 0 |
| chr3R:19241651:19242151: | unc79 | 3979 |

| | | |
|---|---|---|
| chr3L:270701:272151: | miple2 | 0 |
| chr3L:22787851:22788451: | CG14448 | 0 |
| chr3R:30337951:30338751: | CG34300 | 31312 |
| chr2L:21827951:21829151: | tsh | 0 |
| chr3R:5823151:5823701: | CG2023 | 831 |
| chr3R:14108151:14108951: | CG12402 | 0 |
| chr2R:14498201:14498651: | Cpsf160 | 4837 |
| chr3L:13993251:13993851: | CG17364 | 0 |
| chrX:19393351:19393851: | kek5 | 40171 |
| chr3R:9573401:9574051: | CG9399 | 160 |
| chr3R:25328501:25329551: | CHKov1 | 0 |
| chr2R:14139651:14141351: | Prosap | 0 |
| chr3L:7808651:7810201: | CG32369 | 0 |
| chr3R:31310551:31311251: | CG34347 | 0 |
| chr3L:18765601:18766251: | Atg3 | 16915 |
| chr3R:12448801:12450051: | CG18549 | 0 |
| chr3L:6813851:6814301: | vvl | 23694 |
| chr2R:6120251:6120951: | CG14589 | 16352 |
| chr2L:12504101:12505651: | t-cup | 6769 |
| chr3L:10289201:10289951: | Or67d | 15998 |
| chrX:9749551:9750101: | Sp1 | 19904 |
| chr3R:8899551:8900151: | pyd | 31745 |
| chr2L:7809451:7810301: | cdc14 | 402 |
| chr3R:16898501:16900201: | abd-A | 68453 |
| chr3L:18539651:18540151: | CG32198 | 12676 |
| chr2R:11529301:11530051: | en | 1092 |
| chr3R:23965001:23965701: | CG31140 | 1822 |
| chr2L:490101:491201: | cbt | 10414 |
| chrX:12090251:12090951: | CG15734 | 3227 |
| chr3L:5840301:5840851: | vn | 4813 |
| chrX:15803851:15804651: | sd | 0 |
| chr3R:18480551:18481101: | fru | 64485 |
| chr3R:23818551:23819351: | eIF4G2 | 0 |
| chr3L:8913651:8914251: | mfr | 981 |
| chrX:5008551:5009051: | CG12680 | 11111 |
| chr2R:23572901:23573951: | apt | 8011 |
| chr2R:14031151:14031751: | CG18371 | 2532 |
| chr2R:11527601:11528751: | en | 0 |
| chr2L:517901:518701: | lwr | 23879 |
| chr3R:9157751:9158651: | CG11997 | 18866 |
| chr2R:19356351:19357401: | Rgk1 | 0 |
| chrX:8517951:8518551: | Nrg | 579 |
| chr2R:18611501:18612201: | GEFmeso | 0 |
| chr2R:12252501:12253251: | CG13168 | 741 |
| chr2L:7121851:7122401: | Pvf3 | 35294 |

| | | |
|---|---|---|
| chr2L:6791901:6792501: | nrv1 | 5041 |
| chr2L:9781951:9782601: | IP3K1 | 0 |
| chr3R:9872151:9872901: | Glut4EF | 1780 |
| chrX:16562151:16562751: | Pp2B-14D | 2136 |
| chr2L:15332301:15334201: | esg | 0 |
| chr2L:12507401:12508301: | t-cup | 10069 |
| chr3R:7162401:7163151: | sas | 0 |
| chr3R:21581501:21582451: | CG15498 | 12353 |
| chrX:8757601:8758301: | CG1789 | 37082 |
| chr3L:14272701:14273351: | NA-fz | 992 |
| chr2L:16251501:16252201: | CG46309 | 0 |
| chr3L:5975701:5977001: | CG10479 | 0 |
| chr2L:516151:517001: | lwr | 25579 |
| chr2L:5460901:5461801: | mid | 0 |
| chrX:4611201:4611801: | peb | 5763 |
| chr3R:9873301:9875201: | Glut4EF | 0 |
| chr3L:14550701:14551651: | Gbs-70E | 0 |
| chrX:276001:276401: | CG3777 | 3302 |
| chr2L:6793601:6794951: | nrv2 | 3911 |
| chrX:3775701:3781351: | HIP-R | 8618 |
| chr3L:14273751:14274701: | fz | 0 |
| chr3L:18233951:18234851: | CheA75a | 17237 |
| chr2R:6269651:6270201: | Pld | 27648 |
| chr2R:16514751:16515301: | Sema2a | 14574 |
| chrX:1398851:1400101: | CG11448 | 5798 |
| chrX:16559201:16560051: | Pp2B-14D | 0 |
| chr3L:22945051:22945851: | Mes2 | 0 |
| chr3R:16715501:16716301: | Ubx | 18125 |
| chr3R:21807851:21809101: | CASK | 13573 |
| chrX:3545951:3546651: | ppk8 | 11161 |
| chr2L:11581151:11582201: | kek2 | 0 |
| chr3L:19833151:19833651: | trc | 653 |
| chr3L:10567801:10568301: | CG6527 | 8720 |
| chrX:15007701:15008251: | be | 16185 |
| chr3R:29782251:29782801: | fig | 1800 |
| chr3L:18756951:18757801: | Atg3 | 8265 |
| chr2R:20327251:20328101: | CG8929 | 0 |
| chr3R:21297151:21297651: | SIFaR | 3154 |
| chr3R:21295601:21296851: | SIFaR | 1604 |
| chr3L:3983151:3984151: | scrt | 0 |
| chr2R:10618201:10618851: | CG11883 | 540 |
| chr3L:22948251:22948901: | Mes2 | 2595 |
| chr2L:6536051:6536601: | eya | 10375 |
| chr3R:16718451:16719051: | Ubx | 15375 |
| chrX:8510751:8511201: | Nrg | 6172 |

| | | |
|---|---|---|
| chrX:10120651:10121201: | CG12645 | 32411 |
| chr3L:7978851:7980401: | nmo | 0 |
| chrX:2545251:2546051: | trol | 0 |
| chr2L:510301:511051: | cbt | 30614 |
| chr3R:6424001:6424601: | gpp | 17147 |
| chr3L:20310251:20310901: | polo | 598 |
| chr3R:4565251:4565801: | CG34357 | 7185 |
| chr2L:5799301:5800601: | CG9171 | 0 |
| chr2L:5805001:5805651: | CG11034 | 0 |
| chr2R:15135101:15135601: | Rpb12 | 9918 |
| chr2R:6574651:6575601: | CG15233 | 25377 |
| chrX:6189501:6189951: | Ca-alpha1T | 2532 |
| chr3L:17149601:17150401: | Rbp6 | 87811 |
| chr3R:16719601:16720501: | Ubx | 13925 |
| chr3L:11294601:11295251: | CG6175 | 668 |
| chr3L:748201:749701: | emc | 0 |
| chr3R:27194001:27194601: | Ser | 0 |
| chrX:5965501:5966101: | Grip | 178 |
| chrX:16525751:16526601: | Cnx14D | 337 |
| chr2R:14040851:14041651: | CG18371 | 12232 |
| chr2L:6833451:6834101: | sens-2 | 11518 |
| chr2L:12617801:12619001: | Ref2 | 27355 |
| chrX:3773051:3773901: | HIP-R | 16068 |
| chr4:613001:613651: | Eph | 2318 |
| chr3L:6256451:6257301: | Best2 | 0 |
| chr3L:5276501:5278501: | shep | 0 |
| chr2R:11366501:11367801: | CG13204 | 0 |
| chr2R:8326701:8327651: | pdm3 | 0 |
| chr2L:4197101:4197851: | CG3714 | 220 |
| chr2R:6037451:6037851: | SCAP | 1064 |
| chr2R:18061801:18062351: | CG34386 | 64 |
| chr2R:21871501:21872151: | Fili | 0 |
| chrX:14923201:14923801: | CG9503 | 0 |
| chr2L:11358301:11359101: | salr | 0 |
| chr3R:21289001:21289651: | SIFaR | 4347 |
| chr3L:7434951:7435751: | Rac2 | 0 |
| chrX:11713501:11715351: | Amun | 214 |
| chr3R:29774051:29775101: | fig | 5351 |
| chrX:14748901:14750051: | NetB | 0 |
| chr3R:14369951:14370601: | NK7 | 2350 |
| chr2R:7699151:7699751: | CG30496 | 0 |
| chr3L:17740251:17740851: | CG7460 | 0 |
| chr2R:8985401:8986001: | CG8213 | 0 |
| chr2L:12520601:12521751: | t-cup | 23269 |
| chrX:18023451:18024251: | CG6867 | 0 |

| | | |
|---|---|---|
| chrX:1387901:1389001: | CG11448 | 4053 |
| chr3L:21916001:21916801: | SLC22A | 17903 |
| chr2L:9521051:9523451: | Oatp30B | 0 |
| chr3L:10306201:10306701: | Or67d | 32998 |
| chr3R:29771601:29773701: | kay | 5607 |
| chr3L:14108001:14108651: | Sox21a | 0 |
| chr3R:16726401:16727001: | Ubx | 7425 |
| chr2R:9376401:9377101: | CG13954 | 12366 |
| chr3L:10736751:10737851: | CG43245 | 34074 |
| chrX:17506801:17507901: | CG43658 | 53279 |
| chr2R:11107351:11107951: | luna | 8427 |
| chr2R:6102301:6102901: | Ars2 | 16677 |
| chr3L:6262501:6263301: | ImpL3 | 0 |
| chr3R:12467501:12468401: | Hsp70Ba | 0 |
| chr3R:19836601:19837301: | bnl | 0 |
| chr2R:14526351:14527301: | ttv | 85 |
| chr3L:10307801:10308651: | Or67d | 34598 |
| chr3L:2151551:2152151: | zormin | 0 |
| chr3R:14373051:14374001: | NK7 | 101 |
| chr2L:17621101:17621651: | CG15147 | 4730 |
| chr2R:8566001:8566401: | Cyp6a14 | 801 |
| chr3R:4738651:4741101: | Nep2 | 6774 |
| chr3L:603751:604301: | CG13893 | 0 |
| chr4:788801:789301: | MED26 | 0 |
| chr3R:8955251:8956001: | CG11966 | 7036 |
| chr3L:12609051:12609901: | caup | 0 |
| chr3L:10739101:10740601: | CG43245 | 31324 |
| chrX:4779101:4779651: | CG2871 | 1727 |
| chr3R:17920301:17920701: | cpo | 470 |
| chr2R:16499501:16500701: | Sema2a | 0 |
| chrX:18404351:18405001: | CG34328 | 4287 |
| chrX:7604351:7605251: | ct | 3897 |
| chr3R:9704301:9705551: | Unc-115a | 0 |
| chr3L:8559701:8560451: | rhea | 14701 |
| chr2L:16234701:16235301: | CG42817 | 3227 |
| chrX:1133801:1134951: | CG3655 | 0 |
| chr3L:738851:739601: | emc | 9799 |
| chrX:6012951:6013851: | mab-21 | 0 |
| chrX:1321551:1322401: | DAAM | 0 |
| chr2R:7692651:7693401: | LRR | 5501 |
| chr3R:9952801:9953301: | Art4 | 32759 |
| chr2R:6287301:6287751: | Pld | 45298 |
| chr2L:16587601:16589001: | CG31815 | 16821 |
| chrX:7607701:7608701: | ct | 447 |
| chr2R:23551201:23552201: | Dcp-1 | 0 |

| | | |
|---|---|---|
| chr3R:16876651:16877151: | abd-A | 46603 |
| chr3L:11596351:11596951: | CG7368 | 251 |
| chr2L:16231001:16231751: | CG42817 | 0 |
| chrX:7226051:7226401: | CG1958 | 40503 |
| chrX:1380601:1381101: | CG32813 | 0 |
| chr2L:12529151:12530201: | bun | 16410 |
| chr3R:16734251:16734901: | Ubx | 0 |
| chr3L:20294601:20295651: | gogo | 0 |
| chrX:18409951:18410501: | CG32549 | 0 |
| chr3L:15699551:15700601: | comm2 | 0 |
| chrX:7224951:7225301: | CG1958 | 41603 |
| chr3L:15729001:15729901: | comm | 572 |
| chr3R:29764001:29764851: | kay | 1144 |
| chrX:7629151:7629701: | ct | 20004 |
| chr3L:19170301:19172401: | CG14082 | 31327 |
| chr3L:15728151:15728801: | comm | 0 |
| chr2R:22592651:22593701: | dnr1 | 0 |
| chrX:11703001:11703501: | CG2444 | 1799 |
| chr3R:25782901:25783451: | CG4960 | 3847 |
| chrX:10102451:10103201: | RhoU | 15925 |
| chr2L:6252251:6253201: | Ddr | 0 |
| chr3R:7186851:7188451: | lap | 0 |
| chr3R:14382151:14383151: | soti | 453 |
| chr2R:13337151:13337801: | CG17048 | 16698 |
| chr3R:12477201:12478101: | Hsp70Ba | 9426 |
| chrX:7626601:7627701: | ct | 17454 |
| chrX:10302301:10302901: | alpha-Man-Ia | 23057 |
| chr3L:3417801:3418801: | sty | 6134 |
| chr3R:4071551:4072051: | CG42402 | 5888 |
| chr2L:1:6251: | CG11023 | 1278 |
| chr3L:20103751:20105651: | CG14186 | 0 |
| chrX:1375201:1375751: | AMPKalpha | 0 |
| chr3L:3419301:3419801: | sty | 5134 |
| chr2L:3354601:3355501: | E23 | 0 |
| chr3L:20949251:20950351: | fng | 0 |
| chr2R:9389801:9390351: | l | 0 |
| chr3L:3420051:3420701: | sty | 4234 |
| chrX:8785701:8786251: | Lim1 | 19553 |
| chr3R:29690851:29691601: | dmrt99B | 990 |
| chrX:7195901:7196501: | CG9650 | 58179 |
| chr3L:8546251:8546901: | rhea | 1251 |
| chr3L:616401:617301: | CG43337 | 5015 |
| chr3L:12627551:12628151: | caup | 17921 |
| chr3L:17757251:17757801: | CG32182 | 311 |
| chr3L:11586551:11587351: | rt | 419 |

| | | |
|---|---|---|
| chr3R:22476551:22477101: | Nrx-1 | 8040 |
| chrX:6000901:6001651: | CG4766 | 105 |
| chr2R:22513751:22515101: | CG4610 | 21555 |
| chrX:17535701:17536151: | CG43658 | 25029 |
| chrX:10308851:10309501: | alpha-Man-Ia | 16457 |
| chr3R:30160301:30160851: | CG11498 | 0 |
| chr3R:15374251:15374801: | Rbp | 0 |
| chr3L:10325001:10326001: | Or67d | 51798 |
| chr2R:24293801:24294301: | CG3394 | 0 |
| chr3R:19740701:19741251: | GluClalpha | 9987 |
| chr2L:16601201:16602101: | CG31815 | 3721 |
| chr2L:701401:702501: | ds | 12482 |
| chr3L:17872851:17873401: | CG7408 | 5695 |
| chr3R:23781701:23783051: | mbc | 0 |
| chr4:807551:808201: | MED26 | 18574 |
| chr2R:9440651:9442251: | ced-6 | 2686 |
| chr2L:12542901:12544501: | bun | 2110 |
| chr3R:27441301:27441651: | CG44290 | 358 |
| chr2L:13665851:13666501: | CG18507 | 327 |
| chr3R:15378901:15379551: | h-cup | 3400 |
| chrX:8924401:8925351: | CG43255 | 442 |
| chr2R:12704751:12705401: | CG42663 | 0 |
| chrX:11689501:11690001: | PhKgamma | 5136 |
| chr2L:12545151:12546851: | bun | 0 |
| chrX:14950451:14951501: | pdgy | 2031 |
| chr3L:5881151:5882451: | CG13287 | 0 |
| chr3R:27438351:27438751: | side | 870 |
| chr2R:21842901:21843601: | CG4372 | 4104 |
| chrX:10087901:10088451: | RhoU | 1375 |
| chr3R:16857351:16858151: | abd-A | 27303 |
| chrX:14952051:14952701: | pdgy | 831 |
| chr2L:12586801:12587551: | nub | 74 |
| chr3L:14037801:14038751: | Hsc70Cb | 783 |
| chr2L:5421551:5422101: | H15 | 17272 |
| chrX:11685951:11686801: | bif | 6982 |
| chrX:14953351:14953951: | pdgy | 0 |
| chrX:14970951:14971551: | eag | 15475 |
| chr2L:22018501:22019751: | tio | 0 |
| chr2L:10009751:10010351: | Trp1 | 371 |
| chr3L:629901:630401: | CG43337 | 7586 |
| chr3R:9928401:9929251: | Glut4EF | 53721 |
| chr3L:19793351:19793951: | SREBP | 0 |
| chr2L:1737001:1737701: | CG17646 | 4478 |
| chr2R:7597601:7598151: | scra | 7483 |
| chr3L:19790251:19792251: | Gyc76C | 1318 |

| | | |
|---|---|---|
| chrX:11681651:11682251: | bif | 2682 |
| chr2R:9408751:9409851: | wun | 3612 |
| chr2R:7665501:7666051: | CG2064 | 0 |
| chr2L:22023951:22025151: | tio | 4744 |
| chrX:10324201:10324901: | alpha-Man-Ia | 1057 |
| chr2R:22569501:22570601: | CG11362 | 7195 |
| chr3L:10769401:10769801: | CG43245 | 2124 |
| chrX:10344751:10345401: | CG43740 | 11100 |
| chr2L:714701:715751: | ds | 0 |
| chrX:10325151:10326051: | alpha-Man-Ia | 0 |
| chrX:8805651:8807101: | Lim1 | 0 |
| chr3R:12508651:12509251: | Hsp70Bc | 0 |
| chr3L:707601:708651: | CG13895 | 0 |
| chr2L:21632851:21633601: | Ac3 | 265 |
| chr2L:21221601:21222351: | CG8677 | 290 |
| chr3R:12502001:12502651: | Hsp70Bbb | 0 |
| chr2L:2241951:2242601: | CG4267 | 0 |
| chr3L:2712401:2713001: | Fife | 712 |
| chr2L:16857651:16858251: | CG31809 | 463 |
| chr2R:9426351:9427301: | Pdk | 0 |
| chr3L:5926251:5927051: | CG33523 | 2390 |
| chrX:20916501:20916901: | shakB | 10149 |
| chr2R:9413151:9414651: | wun2 | 0 |
| chr2L:6178451:6179551: | smal | 0 |
| chr3R:16763451:16764551: | CG31275 | 12118 |
| chr3R:27426001:27426501: | side | 10981 |
| chr3R:12505351:12506101: | Hsp70Bb | 0 |
| chr2R:4708951:4709501: | Nipped-B | 19516 |
| chr3L:959251:960151: | CG12038 | 33832 |
| chr3R:27988901:27989551: | betaTub97EF | 518 |
| chrX:900751:901351: | CG11663 | 6270 |
| chrX:11616201:11616851: | Met | 78 |
| chr3L:1208051:1208551: | LysB | 681 |
| chr3L:10517501:10518051: | S-Lap4 | 26641 |
| chr3L:20922101:20922651: | CG10589 | 5175 |
| chr3L:10516351:10517051: | S-Lap4 | 25491 |
| chr2L:5403401:5404051: | H15 | 229 |
| chrX:9809001:9810351: | CG32698 | 25779 |
| chr2R:17264851:17265801: | CG43108 | 6966 |
| chr2R:10675001:10675401: | stan | 1657 |
| chr3R:19498601:19500101: | CG15025 | 7487 |
| chr2L:6186401:6186901: | smal | 7269 |
| chr3L:4541601:4543301: | CG11357 | 0 |
| chrX:11621851:11623201: | Ptp10D | 0 |
| chr3L:2107251:2107951: | sls | 7660 |

| | | |
|---|---|---|
| chr3L:11557151:11557801: | CG7394 | 4195 |
| chr3L:3452751:3453251: | eIF5B | 7726 |
| chrX:17963251:17963851: | CG12672 | 43755 |
| chr4:834101:834851: | Sox102F | 0 |
| chr3R:19495001:19495701: | CG15025 | 3887 |
| chr3R:7749851:7750401: | pyd3 | 0 |
| chr3L:8034801:8035501: | bip1 | 23212 |
| chr2L:4594801:4595501: | HP6 | 16584 |
| chr2L:20494301:20494951: | CG34007 | 79 |
| chrX:2765451:2765901: | Syx4 | 21376 |
| chr2L:21617901:21619151: | CG2201 | 4929 |
| chr3L:11552851:11553901: | CG7394 | 8095 |
| chr2L:21237101:21238301: | Hr39 | 0 |
| chr4:557051:557601: | Thd1 | 253 |
| chr3L:10357801:10358201: | CG12362 | 54617 |
| chr2L:3302951:3303551: | CG9663 | 0 |
| chrX:2769251:2769651: | CG32795 | 20916 |
| chr3R:16829901:16830651: | abd-A | 0 |
| chrX:10059651:10060201: | Gr9a | 1426 |
| chrX:8893601:8895101: | Moe | 3231 |
| chr2L:12288451:12288951: | CG31862 | 49897 |
| chr2L:2221501:2222351: | papi | 754 |
| chrX:8891401:8892301: | Moe | 6031 |
| chr3L:685001:685951: | ebd1 | 0 |
| chr2R:12779901:12780801: | sca | 0 |
| chr2L:15109001:15109551: | CG15269 | 0 |
| chr3L:4975901:4976851: | CG17030 | 3918 |
| chr3L:681901:682851: | CG3386 | 0 |
| chr2R:12742951:12743551: | Mos | 9385 |
| chrX:2795851:2796951: | w | 0 |
| chr3R:19463101:19464701: | CG6255 | 36 |
| chr3R:15419051:15419751: | Atx2 | 10629 |
| chr3R:15420101:15421201: | Atx2 | 11679 |
| chr2L:21250451:21251251: | l | 9917 |
| chrX:2789501:2793401: | CG32795 | 0 |
| chr2R:5261901:5262401: | d4 | 2898 |
| chr2R:6501501:6502051: | jing | 0 |
| chr2L:4614201:4614851: | dpy | 0 |
| chr3L:11534301:11534951: | CG11714 | 11589 |
| chr3L:1176951:1178601: | bab2 | 0 |
| chr3L:8827051:8827951: | dally | 0 |
| chr3R:16811651:16812401: | abd-A | 17648 |
| chr3L:3625801:3626551: | dar1 | 0 |
| chr3L:10810801:10811601: | CG42831 | 3218 |
| chr3L:19731151:19731801: | CG42529 | 2158 |

| | | |
|---|---|---|
| chr2L:2201901:2203801: | CG31668 | 2365 |
| chrX:14082551:14083301: | Ste | 20735 |
| chr3R:14447251:14447651: | natalisin | 28659 |
| chrX:20230801:20232001: | sw | 4381 |
| chr2L:2200201:2200751: | CG34172 | 2290 |
| chr2R:19825401:19826851: | Toll-7 | 0 |
| chr3R:14473001:14473451: | cv-c | 8262 |
| chr2L:2196601:2197451: | CG34172 | 461 |
| chr2R:6827901:6828651: | CG15236 | 0 |
| chr2R:24229651:24230201: | CG30419 | 0 |
| chr3R:14468201:14468701: | cv-c | 13012 |
| chr3L:20217101:20217651: | CG6933 | 2769 |
| chr2L:21582301:21582651: | nompB | 6772 |
| chr3L:5184351:5185551: | Srp54k | 31674 |
| chr3R:26229501:26230501: | CG12290 | 0 |
| chr3L:21644551:21645201: | CG43980 | 0 |
| chrX:14653701:14654401: | NetA | 0 |
| chr3L:20185751:20187051: | Ssk | 1471 |
| chr2L:21572551:21573351: | Lamp1 | 38 |
| chr3R:6157001:6157601: | CG31559 | 0 |
| chr2L:21571501:21572201: | Lamp1 | 1188 |
| chr2L:21570551:21571101: | Lamp1 | 2288 |
| chrX:23205301:23206551: | ATbp | 99299 |
| chr2L:1805701:1806251: | Gr22a | 10785 |
| chr2L:2178251:2179151: | aop | 0 |
| chr4:507151:508851: | zfh2 | 5342 |
| chr3L:21476301:21477351: | croc | 0 |
| chr3L:21479401:21480201: | croc | 2923 |
| chrX:9870101:9870551: | CG45061 | 1151 |
| chrX:23210651:23212101: | ATbp | 104649 |
| chr3L:20198101:20198701: | CG42674 | 0 |
| chr3R:31937401:31938201: | CG2003 | 39920 |
| chrX:23212551:23213801: | ATbp | 106549 |
| chr4:501351:502151: | zfh2 | 0 |
| chrX:23214101:23218701: | ATbp | 108099 |
| chr3L:21485101:21485801: | Neu2 | 4785 |
| chr3L:23066001:23066601: | RpL10 | 28870 |
| chr3L:11493301:11493801: | chrb | 5752 |
| chr2L:15056601:15057601: | ck | 261 |
| chr3R:15476051:15476951: | AdamTS-A | 0 |
| chr3L:140701:141301: | CG6845 | 5132 |
| chr2L:2164601:2165201: | CG15382 | 8849 |
| chr3R:31945551:31946151: | CG2003 | 31970 |
| chrX:9880751:9881501: | CG9689 | 452 |
| chr2L:2163551:2164101: | CG15382 | 7799 |

| | | |
|---|---|---|
| chr3L:10851051:10851651: | NA-tna | 6238 |
| chr2L:2162451:2163301: | CG15382 | 6699 |
| chr3L:11487001:11488001: | chrb | 0 |
| chr3L:10852201:10853201: | NA-tna | 4688 |
| chr2L:2161151:2162051: | CG15382 | 5399 |
| chr3L:10856101:10858101: | tna | 0 |
| chrX:14037901:14038701: | Ste | 8794 |
| chr3L:130751:132801: | Pdk1 | 1302 |
| chr4:907701:908351: | unc-13 | 3980 |
| chr2R:7170751:7171401: | CG11060 | 4485 |
| chr3L:21498801:21499701: | Hr78 | 1308 |
| chr2L:21309351:21309851: | Mondo | 199 |
| chr3L:128201:129801: | Pdk1 | 0 |
| chr2R:6871351:6872901: | coro | 0 |
| chr2L:20281751:20282301: | CG12617 | 12392 |
| chr2L:1062501:1062901: | IA-2 | 14449 |
| chr3L:11428301:11429001: | CG7560 | 3144 |
| chr2L:21540901:21541301: | His3 | 0 |
| chr3L:21509551:21510001: | M6 | 1700 |
| chr3R:7309551:7310201: | rn | 0 |
| chr3L:21604401:21604851: | S1P | 12525 |
| chr3L:23090151:23090701: | RpL10 | 4770 |
| chr2L:21534401:21534701: | His3 | 0 |
| chr3L:3542251:3543901: | Eip63E | 26501 |
| chrX:23242851:23243501: | ATbp | 136849 |
| chr3L:3544801:3545501: | Eip63E | 29051 |
| chrX:9905001:9905751: | CG1986 | 5375 |
| chr2L:21529501:21529901: | His3 | 0 |
| chr2R:6885551:6886101: | Spn42De | 0 |
| chr3L:21598051:21598951: | TfAP-2 | 11224 |
| chrX:23246201:23246901: | ATbp | 140199 |
| chr2R:7150351:7150901: | pk | 0 |
| chrX:23249601:23250301: | ATbp | 143599 |
| chr2L:21524701:21525051: | His3 | 0 |
| chr4:931501:932101: | eIF4G1 | 787 |
| chr3L:1102101:1103001: | bab1 | 1013 |
| chr4:932501:933051: | eIF4G1 | 1787 |
| chr2R:7146751:7147351: | Spn43Aa | 341 |
| chrX:23253001:23253901: | ATbp | 146999 |
| chr4:933951:935001: | mGluR | 1818 |
| chr2L:2130051:2131001: | GlyP | 0 |
| chr2L:21519851:21520201: | His3 | 0 |
| chrX:20158801:20159701: | Rab35 | 171 |
| chr3L:1098451:1099201: | bab1 | 1888 |
| chr4:936301:936901: | mGluR | 0 |

| | | |
|---|---|---|
| chr4:457501:458001: | bip2 | 8827 |
| chr2L:16972601:16973101: | beat-IIIb | 0 |
| chr3L:21584501:21585251: | TfAP-2 | 1577 |
| chr2L:21514801:21515151: | His3 | 0 |
| chr2L:20310351:20311451: | spir | 0 |
| chr4:453801:454451: | CaMKI | 8298 |
| chr2L:15013351:15013951: | GABA-B-R1 | 2380 |
| chr2L:21509751:21510101: | His3 | 0 |
| chrX:13986451:13987101: | mamo | 3426 |
| chr3R:6002901:6003301: | Rm62 | 5278 |
| chrX:13994551:13995551: | ben | 794 |
| chr2L:21504701:21505051: | His3 | 0 |
| chr2L:22505101:22505651: | RpL5 | 31233 |
| chr2L:15745301:15746501: | CycE | 1649 |
| chr2L:15747201:15748501: | CycE | 0 |
| chr3R:6007501:6008151: | Rm62 | 428 |
| chr2L:21499601:21500101: | His3 | 0 |
| chrX:23278001:23281101: | ATbp | 171999 |
| chr2L:14995401:14996051: | mol | 1508 |
| chr2L:21494601:21494951: | His3 | 0 |
| chrX:23282001:23284301: | ATbp | 175999 |
| chr4:431851:432401: | lgs | 11510 |
| chrX:21634001:21634601: | DIP1 | 3854 |
| chr2L:21489551:21489901: | His3 | 0 |
| chr2L:1885951:1886851: | Wdr62 | 1330 |
| chr2L:15762351:15762951: | Gli | 0 |
| chr2L:1118751:1119651: | CG14340 | 5811 |
| chr2L:21484351:21484851: | His3 | 0 |
| chr3R:32015201:32016001: | heph | 0 |
| chr2R:7106301:7106801: | esn | 19986 |
| chr2L:16049551:16050151: | beat-Ia | 0 |
| chr2L:21479351:21479751: | His3 | 0 |
| chr2L:21474301:21474701: | His3 | 0 |
| chrX:66601:67401: | CG17636 | 59313 |
| chr2L:21469251:21469701: | His3 | 0 |
| chr4:404251:404701: | dati | 8647 |
| chr2L:21464251:21464651: | His3 | 0 |
| chr2R:5401601:5402101: | laccase2 | 11643 |
| chr2L:22549751:22550251: | CG17493 | 0 |
| chr2L:21459151:21459601: | His3 | 0 |
| chr3L:40001:40951: | CG43149 | 13800 |
| chr2L:21454051:21454551: | His3 | 0 |
| chr2L:21439851:21440351: | His3 | 0 |
| chr2L:21435001:21435451: | His3 | 0 |
| chr2L:21419801:21420251: | His3 | 0 |

| | | |
|---|---|---|
| chr2L:21430151:21430401: | His3 | 0 |
| chr2L:21424801:21425451: | His3 | 0 |
| chr3R:32071001:32072401: | Map205 | 2561 |
| chr3R:26536751:26537451: | scrib | 402 |
| chr2R:7049401:7050001: | lbm | 164 |
| chr2R:6992501:6993101: | CG30159 | 0 |
| chr2L:1954301:1955151: | CG33673 | 7574 |
| chrX:6551:7351: | CG17636 | 119363 |
| chr2L:1970201:1971101: | erm | 0 |
| chr2L:2013751:2014351: | CG4238 | 3893 |
| chr2L:1971351:1971851: | erm | 990 |
| chr2L:1972451:1973351: | Der-1 | 606 |
| chr3R:3788401:3789601: | CG41128 | 38039 |
| chrX:22438751:22439301: | CG14621 | 4926 |
| chr4:306401:307051: | Syt7 | 0 |
| chr4:303351:304001: | Syt7 | 2480 |
| chr4:302451:303101: | Syt7 | 3380 |
| chr4:295251:295951: | Syt7 | 10530 |
| chr4:294451:295001: | Syt7 | 11480 |
| chr2R:4169201:4169801: | RNASEK | 18326 |
| chrX:23430601:23433801: | ATbp | 324599 |
| chr2L:15911601:15912701: | CG13244 | 4187 |
| chrX:23438701:23440101: | ATbp | 332699 |
| chrX:23440301:23440901: | ATbp | 334299 |
| chrX:22382651:22383701: | CG14621 | 60526 |
| chr4:207401:207951: | CG31998 | 1457 |
| chr4:183351:184001: | Arl4 | 3963 |
| chrX:23535051:23538701: | ATbp | 429049 |
| chr4:1257851:1258501: | Cadps | 27015 |
| chr4:1260001:1260401: | Cadps | 29165 |
| chrX:22260201:22260401: | CG41562 | 8576 |
| chrX:22258251:22259301: | CG41562 | 6626 |
| chr4:1268401:1269001: | Cadps | 37565 |
| chr4:1274751:1275151: | Cadps | 43915 |
| chr3R:3588801:3589801: | Parp | 1748 |
| chr4:1294901:1295201: | Cadps | 64065 |
| chr4:1318551:1319451: | Cadps | 87715 |
| chr4:53601:54201: | ci | 2840 |
| chr4:52701:53251: | ci | 3790 |
| chr2R:3934351:3934801: | uex | 14624 |
| chr4:41301:41901: | PlexB | 1877 |
| chr4:33701:34201: | PlexB | 9577 |
| chr3L:23582601:23583151: | AGO3 | 28009 |
| chr3R:3344651:3345251: | spok | 21842 |
| chr2R:3750401:3751051: | RYa | 103056 |

| | | |
|---|---|---|
| chr2L:23093101:23093501: | Spf45 | 24716 |
| chr2L:23096951:23097851: | Spf45 | 28566 |
| chr2L:23099601:23101451: | Spf45 | 31216 |
| chr2R:3731151:3731701: | RYa | 83806 |
| chr2L:23184951:23185401: | CG12567 | 10468 |
| chr2L:23186151:23186551: | CG12567 | 11668 |
| chr2L:23189351:23189851: | CG12567 | 14868 |
| chr3R:3169651:3169851: | Pzl | 93731 |
| chr2L:23512601:23513851: | Tim23 | 199024 |
| chr2R:3311851:3312301: | CG41242 | 52101 |
| chr3L:24210051:24210701: | nvd | 114165 |
| chr3L:24286851:24287401: | nvd | 37465 |
| chr2R:3062801:3063451: | dpr21 | 3048 |
| chr2R:3062001:3062451: | dpr21 | 4048 |
| chr3L:24392301:24392801: | nvd | 67436 |
| chr3R:2568551:2569051: | Pzl | 694531 |
| chr2R:2904501:2905001: | CG41378 | 57794 |
| chr3R:2449851:2450251: | Pzl | 813331 |
| chr2R:2841851:2842401: | CG41378 | 4307 |
| chr3L:24608801:24610351: | mRpS5 | 71566 |
| chr3L:24610651:24611301: | mRpS5 | 73416 |
| chr3L:24611651:24612201: | mRpS5 | 74416 |
| chr3L:24614101:24614651: | mRpS5 | 76866 |
| chr3L:24618601:24619201: | mRpS5 | 81366 |
| chr3L:24622751:24624001: | mRpS5 | 85516 |
| chr3L:24626901:24627351: | mRpS5 | 89666 |
| chr2R:2805151:2805801: | CG41378 | 40907 |
| chr3L:24656601:24657001: | scro | 67161 |
| chr3L:24674301:24674951: | scro | 49211 |
| chr3L:24684851:24685251: | scro | 38911 |
| chr3R:2116601:2117101: | Pzl | 1146481 |
| chr3L:25085851:25086451: | Set1 | 21143 |
| chr2R:2260251:2260751: | CG17684 | 1364 |
| chr3L:25804051:25804701: | FASN3 | 41273 |
| chr3L:25805301:25805951: | FASN3 | 42523 |
| chr3L:25874401:25875001: | FASN3 | 111623 |
| chr2R:1528651:1529301: | Maf1 | 1557 |
| chr2R:1524101:1524801: | Maf1 | 2294 |
| chr2R:1512451:1513051: | Maf1 | 14044 |
| chr3R:951951:952401: | Myo81F | 384876 |
| chr2R:1233401:1234601: | CG40191 | 29539 |
| chr2R:870051:870601: | CG46302 | 15238 |
| chr2R:748751:749101: | CG46302 | 105713 |
| chr2R:601751:602351: | CG46302 | 252463 |
| chr2R:553601:554001: | CG45781 | 226834 |

| | | |
|---|---|---|
| chr2R:552751:553301: | CG45781 | 225984 |
| chr3L:27007451:27011201: | CG40228 | 72705 |
| chr3L:27068551:27073601: | CG40228 | 10305 |
| chr3L:27075001:27075451: | CG40228 | 8455 |
| chr3L:27075951:27077551: | CG40228 | 6355 |
| chr3L:27079151:27079601: | CG40228 | 4305 |
| chr3L:27081451:27082251: | CG40228 | 1655 |
| chr3L:27129901:27130501: | vtd | 6024 |
| chr2R:318751:319351: | CG45781 | 7417 |
| chr3L:27289601:27290101: | Dbp80 | 10286 |
| chr3L:27356801:27357501: | Dbp80 | 77486 |
| chr2R:1151:2701: | CG45781 | 324067 |

| **Coordinates_Mitotic_Only** | **nearst_gene** | **distance_nearest_gene** |
|---|---|---|
| chr2L:11541951:11542301: | CG43438 | 5807 |
| chr2L:11542801:11543101: | CG43438 | 6657 |
| chr2L:11544601:11545051: | CG43438 | 8457 |
| chr2L:16141301:16141401: | CG34168 | 6030 |
| chr2L:16146051:16146251: | CG34168 | 10780 |
| chr2L:23101701:23102201: | Spf45 | 33316 |
| chr2L:23103701:23104001: | Spf45 | 35316 |
| chr2L:23118851:23119051: | Spf45 | 50466 |
| chr2L:23123801:23123901: | CG12567 | 50583 |
| chr2L:23231001:23231301: | CG12567 | 56518 |
| chr2L:23231701:23231851: | CG12567 | 57218 |
| chr2L:23374101:23374401: | Tim23 | 60524 |
| chr2R:1515751:1515851: | Maf1 | 11244 |
| chr2R:1521451:1521851: | Maf1 | 5244 |
| chr2R:1522201:1522651: | Maf1 | 4444 |
| chr2R:15227701:15227851: | Hex-C | 7373 |
| chr2R:1523051:1523701: | Maf1 | 3394 |
| chr2R:1720001:1720101: | CG41520 | 70941 |
| chr2R:1809451:1810151: | CG41520 | 160391 |
| chr2R:1826051:1826151: | CG41520 | 176991 |
| chr2R:2088601:2089201: | CG17684 | 172914 |
| chr2R:2090501:2090751: | CG17684 | 171364 |
| chr2R:2165551:2165751: | CG17684 | 96364 |
| chr2R:2165951:2166201: | CG17684 | 95914 |
| chr2R:2168401:2168651: | CG17684 | 93464 |
| chr2R:24701701:24702001: | ST6Gal | 7316 |
| chr2R:3492051:3492251: | CG41242 | 127650 |
| chr2R:3509001:3509501: | RYa | 137845 |
| chr2R:3511201:3511401: | RYa | 135945 |
| chr2R:3511801:3512101: | RYa | 135245 |
| chr2R:3683751:3684201: | RYa | 36406 |

| | | |
|---|---|---|
| chr2R:3763351:3763551: | RYa | 116006 |
| chr2R:3885051:3885201: | uex | 64224 |
| chr2R:3888901:3889001: | uex | 60424 |
| chr2R:3889901:3890051: | uex | 59374 |
| chr2R:4863851:4864451: | CG14464 | 34606 |
| chr2R:4864651:4865351: | CG14464 | 35406 |
| chr2R:4865651:4866951: | CG14464 | 36406 |
| chr2R:4867201:4867501: | CG14464 | 37956 |
| chr2R:4867701:4868601: | CG14464 | 38456 |
| chr2R:583501:583801: | CG45781 | 256734 |
| chr2R:586101:586751: | CG45781 | 259334 |
| chr2R:5901:6251: | CG45781 | 320517 |
| chr2R:6551:7451: | CG45781 | 319317 |
| chr3L:23112901:23113751: | CG33217 | 12274 |
| chr3L:23114051:23114301: | CG33217 | 11724 |
| chr3L:23114551:23114901: | CG33217 | 11124 |
| chr3L:23115651:23115951: | CG33217 | 10074 |
| chr3L:23116151:23117301: | CG33217 | 8724 |
| chr3L:23571701:23572151: | AGO3 | 17109 |
| chr3L:24173451:24173751: | Snap25 | 99272 |
| chr3L:24455151:24455251: | mRpS5 | 81985 |
| chr3L:24593151:24595451: | mRpS5 | 55916 |
| chr3L:24595951:24596301: | mRpS5 | 58716 |
| chr3L:24597001:24597201: | mRpS5 | 59766 |
| chr3L:24602251:24603301: | mRpS5 | 65016 |
| chr3L:24604351:24604751: | mRpS5 | 67116 |
| chr3L:24605451:24606001: | mRpS5 | 68216 |
| chr3L:24606601:24608601: | mRpS5 | 69366 |
| chr3L:24612451:24612701: | mRpS5 | 75216 |
| chr3L:24613251:24613401: | mRpS5 | 76016 |
| chr3L:24615001:24615201: | mRpS5 | 77766 |
| chr3L:24615651:24616451: | mRpS5 | 78416 |
| chr3L:24616651:24616751: | mRpS5 | 79416 |
| chr3L:24619451:24619901: | mRpS5 | 82216 |
| chr3L:24620101:24621001: | mRpS5 | 82866 |
| chr3L:24622001:24622401: | mRpS5 | 84766 |
| chr3L:24624851:24625551: | mRpS5 | 87616 |
| chr3L:24626301:24626451: | mRpS5 | 89066 |
| chr3L:24628001:24628601: | mRpS5 | 90766 |
| chr3L:24629051:24629901: | mRpS5 | 91816 |
| chr3L:24631451:24632201: | scro | 91961 |
| chr3L:24632901:24633051: | scro | 91111 |
| chr3L:24633351:24633751: | scro | 90411 |
| chr3L:24634501:24637051: | scro | 87111 |
| chr3L:24637251:24637451: | scro | 86711 |

| | | |
|---|---|---|
| chr3L:24640351:24640551: | scro | 83611 |
| chr3L:24644551:24644751: | scro | 79411 |
| chr3L:24647901:24648401: | scro | 75761 |
| chr3L:24657301:24657501: | scro | 66661 |
| chr3L:24657701:24657801: | scro | 66361 |
| chr3L:24658551:24658701: | scro | 65461 |
| chr3L:24672401:24672701: | scro | 51461 |
| chr3L:24673051:24673301: | scro | 50861 |
| chr3L:24683351:24683701: | scro | 40461 |
| chr3L:25131101:25131551: | UQCR-11 | 10393 |
| chr3L:25132751:25133201: | UQCR-11 | 12043 |
| chr3L:25133451:25135051: | UQCR-11 | 12743 |
| chr3L:25963101:25963301: | CG40178 | 50811 |
| chr3L:26878251:26879051: | CG40228 | 204855 |
| chr3L:26879251:26879401: | CG40228 | 204505 |
| chr3L:26879751:26880101: | CG40228 | 203805 |
| chr3L:26880951:26881051: | CG40228 | 202855 |
| chr3L:26881551:26882051: | CG40228 | 201855 |
| chr3L:26936901:26937351: | CG40228 | 146555 |
| chr3L:26938051:26938751: | CG40228 | 145155 |
| chr3L:26938951:26939351: | CG40228 | 144555 |
| chr3L:26939651:26940151: | CG40228 | 143755 |
| chr3L:26940801:26940901: | CG40228 | 143005 |
| chr3L:26942851:26943201: | CG40228 | 140705 |
| chr3L:26944001:26944401: | CG40228 | 139505 |
| chr3L:26945001:26945151: | CG40228 | 138755 |
| chr3L:26946851:26947201: | CG40228 | 136705 |
| chr3L:26949451:26950251: | CG40228 | 133655 |
| chr3L:27006601:27007051: | CG40228 | 76855 |
| chr3L:27074151:27074501: | CG40228 | 9405 |
| chr3L:27078101:27078651: | CG40228 | 5255 |
| chr3L:28057551:28058101: | CG17514 | 44953 |
| chr3L:28073051:28073201: | CG17514 | 60453 |
| chr3L:619801:620001: | CG43337 | 2315 |
| chr3R:123151:123301: | Myo81F | 443775 |
| chr3R:1379251:1379351: | Myo81F | 812176 |
| chr3R:1441401:1441551: | Myo81F | 874326 |
| chr3R:1620401:1620551: | Myo81F | 1053326 |
| chr3R:172801:173051: | Myo81F | 394025 |
| chr3R:183701:183901: | Myo81F | 383175 |
| chr3R:2260401:2261051: | Pzl | 1002531 |
| chr3R:2262001:2262151: | Pzl | 1001431 |
| chr3R:2263151:2263851: | Pzl | 999731 |
| chr3R:2264301:2264851: | Pzl | 998731 |
| chr3R:2449101:2449251: | Pzl | 814331 |

| | | |
|---|---|---|
| chr3R:253851:253951: | Myo81F | 313125 |
| chr3R:267751:267951: | Myo81F | 299125 |
| chr3R:27551:27751: | Myo81F | 539325 |
| chr3R:2969651:2969751: | Pzl | 293831 |
| chr3R:3033351:3033801: | Pzl | 229781 |
| chr3R:316101:316301: | Myo81F | 250775 |
| chr3R:324951:325201: | Myo81F | 241875 |
| chr3R:3366401:3366701: | spok | 43592 |
| chr3R:3394651:3394751: | spok | 71842 |
| chr3R:3834551:3834901: | CG41099 | 11828 |
| chr3R:3836351:3836501: | CG41099 | 10228 |
| chr3R:471551:471951: | Myo81F | 95125 |
| chr3R:728251:728601: | Myo81F | 161176 |
| chr4:1277651:1277901: | Cadps | 46815 |
| chr4:1279901:1280001: | Cadps | 49065 |
| chr4:1288251:1288501: | Cadps | 57415 |
| chr4:1296001:1296201: | Cadps | 65165 |
| chr4:593501:593651: | CG1909 | 10394 |
| chr4:594401:595051: | CG1909 | 11294 |
| chr4:77901:78001: | pan | 8576 |
| chrM:101:851: | | -1 |
| chrM:10451:13451: | | -1 |
| chrM:1251:6251: | | -1 |
| chrM:14051:14651: | | -1 |
| chrM:6551:7801: | | -1 |
| chrM:8251:9151: | | -1 |
| chrX:10007301:10007501: | CG33557 | 1709 |
| chrX:10048901:10049001: | Yp2 | 3635 |
| chrX:10292001:10292251: | CG12643 | 26739 |
| chrX:10340151:10340401: | CG43740 | 6500 |
| chrX:10488951:10489251: | CG32687 | 1747 |
| chrX:1074251:1074451: | CG14629 | 22727 |
| chrX:12046851:12047851: | Ten-a | 3154 |
| chrX:12124501:12124951: | CG2750 | 17302 |
| chrX:12545351:12545651: | Cpr11B | 7473 |
| chrX:12795101:12795401: | CG3775 | 6260 |
| chrX:12796151:12796401: | CG3775 | 5260 |
| chrX:12797701:12797801: | CG3775 | 3860 |
| chrX:202851:203051: | CG3038 | 42805 |
| chrX:20380901:20381101: | CG12679 | 4380 |
| chrX:21807851:21808251: | DIP1 | 177704 |
| chrX:21808651:21809151: | DIP1 | 178504 |
| chrX:21810351:21811001: | DIP1 | 180204 |
| chrX:21957101:21957201: | CG40813 | 185901 |
| chrX:21958801:21958901: | CG40813 | 184201 |

| | | |
|---|---|---|
| chrX:21973351:21973501: | CG40813 | 169601 |
| chrX:22259701:22259951: | CG41562 | 8076 |
| chrX:22273901:22274251: | CG41562 | 22276 |
| chrX:22274901:22275301: | CG41562 | 23276 |
| chrX:22277251:22277351: | CG41562 | 25626 |
| chrX:22278151:22278551: | CG41562 | 26526 |
| chrX:22280801:22281001: | CG41562 | 29176 |
| chrX:22315001:22315101: | CG41562 | 63376 |
| chrX:22354351:22354551: | CG14621 | 89676 |
| chrX:22362751:22362951: | CG14621 | 81276 |
| chrX:22363301:22363801: | CG14621 | 80426 |
| chrX:22369951:22370301: | CG14621 | 73926 |
| chrX:22370551:22370651: | CG14621 | 73576 |
| chrX:22370951:22371301: | CG14621 | 72926 |
| chrX:22374101:22374401: | CG14621 | 69826 |
| chrX:22384201:22384351: | CG14621 | 59876 |
| chrX:22385851:22386051: | CG14621 | 58176 |
| chrX:23219251:23219501: | ATbp | 113249 |
| chrX:23223401:23223701: | ATbp | 117399 |
| chrX:23224351:23225051: | ATbp | 118349 |
| chrX:23225401:23225751: | ATbp | 119399 |
| chrX:23225951:23228501: | ATbp | 119949 |
| chrX:23231801:23232351: | ATbp | 125799 |
| chrX:23234401:23234551: | ATbp | 128399 |
| chrX:23234751:23235001: | ATbp | 128749 |
| chrX:23235251:23235501: | ATbp | 129249 |
| chrX:23239651:23240201: | ATbp | 133649 |
| chrX:23240401:23241251: | ATbp | 134399 |
| chrX:23256601:23257051: | ATbp | 150599 |
| chrX:23261351:23262101: | ATbp | 155349 |
| chrX:23270801:23271201: | ATbp | 164799 |
| chrX:23272551:23272901: | ATbp | 166549 |
| chrX:23273751:23274501: | ATbp | 167749 |
| chrX:23290151:23290551: | ATbp | 184149 |
| chrX:23290951:23291551: | ATbp | 184949 |
| chrX:23291751:23292001: | ATbp | 185749 |
| chrX:23327301:23327551: | ATbp | 221299 |
| chrX:23343601:23343751: | ATbp | 237599 |
| chrX:23344751:23344851: | ATbp | 238749 |
| chrX:23352901:23353001: | ATbp | 246899 |
| chrX:23354651:23355101: | ATbp | 248649 |
| chrX:23358251:23358501: | ATbp | 252249 |
| chrX:23367851:23368301: | ATbp | 261849 |
| chrX:23371701:23371801: | ATbp | 265699 |
| chrX:23375851:23376051: | ATbp | 269849 |

| | | |
|---|---|---|
| chrX:23377051:23377501: | ATbp | 271049 |
| chrX:23381101:23381201: | ATbp | 275099 |
| chrX:23383201:23383601: | ATbp | 277199 |
| chrX:23385501:23385601: | ATbp | 279499 |
| chrX:23385801:23385901: | ATbp | 279799 |
| chrX:23388051:23388251: | ATbp | 282049 |
| chrX:23388501:23388701: | ATbp | 282499 |
| chrX:23389301:23389401: | ATbp | 283299 |
| chrX:23391751:23391901: | ATbp | 285749 |
| chrX:23398001:23398351: | ATbp | 291999 |
| chrX:23406801:23407551: | ATbp | 300799 |
| chrX:23412301:23412751: | ATbp | 306299 |
| chrX:23414651:23415101: | ATbp | 308649 |
| chrX:23421201:23421301: | ATbp | 315199 |
| chrX:23425901:23426051: | ATbp | 319899 |
| chrX:23441701:23442101: | ATbp | 335699 |
| chrX:23455401:23455701: | ATbp | 349399 |
| chrX:23455951:23457451: | ATbp | 349949 |
| chrX:23472151:23472551: | ATbp | 366149 |
| chrX:23472951:23473501: | ATbp | 366949 |
| chrX:23486301:23486601: | ATbp | 380299 |
| chrX:23526051:23526351: | ATbp | 420049 |
| chrX:6206701:6207501: | vanin-like | 4143 |
| chrX:7026951:7027301: | CG4593 | 13065 |
| chrX:7028601:7029001: | CG14427 | 11602 |
| chrX:7029451:7030001: | CG14427 | 10602 |
| chrX:8661201:8661451: | oc | 10521 |
| chrX:9033951:9034301: | Nost | 3747 |
| chrX:9092951:9093351: | CCT2 | 3439 |
| chrX:9203951:9204301: | su | 2132 |
| chrX:9235151:9235501: | Ir8a | 1084 |
| chrX:9374201:9374651: | BCL7-like | 17568 |
| chrX:9396801:9397201: | BCL7-like | 40168 |
| chrX:9461701:9462051: | CG44815 | 15490 |
| chrX:9534801:9535001: | CG34026 | 984 |
| chrX:9556851:9557301: | CG15317 | 308 |
| chrX:9597451:9598051: | Gga | 2979 |
| chrX:9624001:9624101: | CG42797 | 15056 |
| chrX:9625451:9625551: | CG42797 | 16506 |
| chrX:9625801:9625901: | CG42797 | 16856 |
| chrX:970951:971201: | CG3690 | 19052 |
| chrX:9711001:9711251: | btd | 16816 |
| chrX:9726301:9727101: | Sp1 | 2547 |
| chrX:9919651:9920101: | CG1986 | 8526 |
| chrX:9956151:9956351: | CG1791 | 8527 |

| | | |
|---|---|---|
| chrX:9956651:9956801: | CG1791 | 8077 |
| chrY:1020151:1020301: | ARY | 317722 |
| chrY:1027451:1027551: | ARY | 325022 |
| chrY:1028801:1028901: | ARY | 326372 |
| chrY:1631451:1631551: | Ppr-Y | 5023 |
| chrY:226401:226801: | PRY | 34382 |
| chrY:227801:228351: | PRY | 35782 |
| chrY:2610201:2610301: | WDY | 336780 |
| chrY:2633501:2634151: | WDY | 360080 |
| chrY:2654151:2654401: | FDY | 367888 |
| chrY:322801:324101: | kl-3 | 12280 |
| chrY:3302901:3303051: | CG45766 | 1075 |
| chrY:3560401:3560551: | ORY | 1194 |
| chrY:653051:653251: | ARY | 49179 |
| chrY:816301:816401: | ARY | 113872 |
| chrY:884601:884951: | ARY | 182172 |
| chrY:890151:890351: | ARY | 187722 |
| chrY:937851:938051: | ARY | 235422 |
| chrY:955551:955751: | ARY | 253122 |
| chrY:956351:956551: | ARY | 253922 |
| chrY:958501:958651: | ARY | 256072 |
| chrY:974451:975551: | ARY | 272022 |
| chrY:988251:988551: | ARY | 285822 |
| chrY:989951:990051: | ARY | 287522 |
| chrY:991851:991951: | ARY | 289422 |
| chrY:993651:993751: | ARY | 291222 |
| chrY:993951:994401: | ARY | 291522 |
