## Supplementary Table 4 for "The control of transcriptional memory by stable mitotic bookmarking"

| Experiment | Stage | Reference |
|---|---|---|
| TSS | - | GCF_000001215.4 (Release_6) |
| GAF-ChIP-seq | 2-4 hr AEL, WT embryos | *Koenecke et al. 2017* |
| H3K27ac-ChIP-seq | 2-4 hr AEL, tol10b embryos | *Koenecke et al. 2016* |
| H3K27ac-ChIP-seq | 2-4 hr AEL, gd7 embryos | *Koenecke et al. 2016* |
| Enhancer reporter transgenes | Embryos | *Kvon et al. 2014* |
| Hi-C | nc12, nc13, nc14 | *Hug et al. 2017* |
| Insulators | Embryos | *Negre et al. 2010* |
| ATAC-seq | nc8, 9, 10…13 every 3min | *Blythe & Wieschaus 2016* |
| ATAC-seq degrad-FP | 2–2.5 hr AEL | *Gaskill et al. 2021* |
| H3K27ac | nc14 | *Li et al. 2014* |
| H3K27me3 | nc14 | *Li et al. 2014* |
| RNA-seq | nc14 | *Lott et al. 2011* |
